# Substrate scope and catalytic mechanism of α, β-epoxyketone synthase EpnF illuminated by *in situ* esterase-mediated deprotection

**DOI:** 10.64898/2025.12.23.696313

**Authors:** Callum J. Bullock, Marlene L. Rothe, Joshua W. Cartwright, Munro Passmore, Alex J. Mullins, Fabrizio Alberti, Józef R. Lewandowski, Lona M. Alkhalaf, Gregory L. Challis

## Abstract

α, β-Epoxyketones are an important class of bacterial natural products with wide-ranging potential applications in oncology, immunology, and infectious disease. Their clinical potential derives from the α, β-epoxyketone pharmacophore, which covalently modifies the *N*-terminal catalytic threonine residue of proteasome β-subunits with high selectivity. Although a synthetic α, β-epoxyketone (Carfilzomib) is approved for clinical use, stereo-controlled synthesis of the pharmacophore remains challenging, involving either multiple steps and energy-intensive processes, or unsustainable reagents. Unusual flavoenzymes catalyze the assembly of the pharmacophore in α, β-epoxyketone biosynthesis. A detailed understanding of the substrate scope and catalytic mechanism of these enzymes has thus far been limited by the intrinsic instability of their α- (di)methyl-β-ketoacid substrates. Here, we report the development and application of an esterase-mediated unmasking strategy for *in-situ* generation of these substrates from the corresponding methyl esters. Using this approach, we demonstrate that EpnF, the epoxyketone synthase involved in eponemycin / TMC-86A biosynthesis, tolerates a broad range of synthetic substrate analogs, including several with *N*-terminal protecting groups widely used in peptide synthesis. These findings establish that EpnF has the potential to be developed into a useful biocatalyst for the chemoenzymatic synthesis of dipeptidyl epoxyketone precursors of clinically approved drugs and drug candidates. To elucidate the molecular basis for catalysis of α, β-epoxyketone formation by EpnF, substrate docking and molecular dynamics simulations were performed on a well-validated AlphaFold model, providing support for a previously proposed decarboxylation-dehydrogenation-monooxygenation mechanism. Site-directed mutagenesis and LC–MS analysis validated the proposed roles of key active-site residues in substrate positioning and catalysis of epoxide formation. Collectively, these results demonstrate that EpnF and related enzymes belong to a new class of internal flavoprotein monooxygenases and provide a foundation for developing epoxyketone synthases into useful biocatalysts for the sustainable synthesis of high-value α,β-epoxyketones.

## INTRODUCTION

Microbial natural products are a vast and chemically diverse group of specialized metabolites that have profoundly influenced modern drug discovery and development.^1^ Investigating the biosynthetic mechanisms for such metabolites not only illuminates the chemical ingenuity of specialized metabolism but also discovers a wealth of biocatalysts capable of performing synthetically challenging transformations with high regio/stereo-control and broad substrate scope.

One important class of microbial specialized metabolites are hybrids of nonribosomal peptides and polyketides. Molecules belonging to this class have important applications in the treatment of various forms of cancer. Examples include the DNA strand–cleaving agent bleomycin **1**, the potent histone deacetylase inhibitor romidepisn (FK228; **2**), and the microtubule-stabilizer epothilone, which formed the basis for development of ixabepilone **3** (Fig. 1A).^2–4^

**Figure 1.**
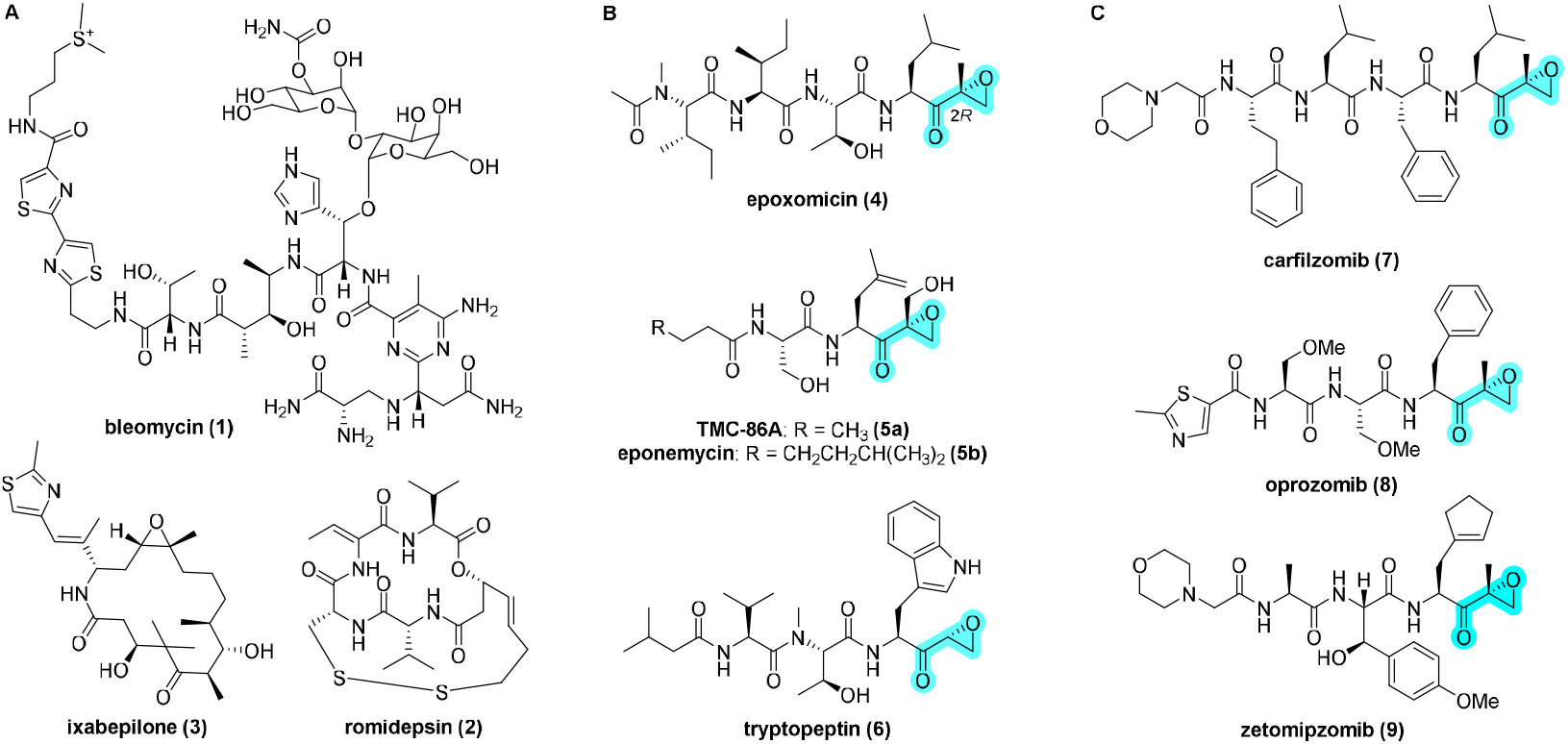
(**A**) Examples of Hybrid nonribosomal peptide-polyketide bacterial specialized metabolites used to treat cancer. Ixabepilone is a semisynthetic derivative of epothilone B. (**B**) Representative members of the peptidyl epoxyketone class of proteasome-inhibiting bacterial natural products. (**C**) Examples of synthetic peptidyl epoxyketone proteasome-inhibiting drugs and drug candidates inspired by natural products. Carfilzomib and oprozomib are being used and developed, respectively, for the treatment of multiple myeloma. Zetomipzomib has undergone a phase IIa trial for autoimmune hepatitis. The epoxyketone pharmacophore of the proteasome-inhibiting natural products and drugs is highlighted in cyan.

Among the most mechanistically intriguing anticancer natural products are the peptidyl α,β-epoxyketone proteasome inhibitors, exemplified by epoxomicin **4**, TMC-86a **5a**/ eponemycin **5b** and tryptopeptin A **6**(Fig. 1B).^5–7^ These metabolites feature a variable di, tri, or tetrapeptide fused to a C-terminal α,β-epoxyketone pharmacophore, which is an irreversible and highly selective covalent inhibitor of proteasomes.^8,9^

The proteasome is a multi-subunit N-terminal nucleophile (Ntn) hydrolase complex central to regulated protein degradation in eukaryotic cells. It mediates the proteolytic degradation of ubiquitinylated substrates using a catalytic *N*-terminal threonine residue within each β-subunit.^10^

Epoxyketone-mediated proteasome inhibition proceeds through formation of a dual covalent adduct with the *N*-terminal threonine of the β5 subunit.^11^ Nucleophilic addition of the threonine hydroxyl group to the carbonyl group of the epoxyketone is followed by intramolecular attack of the epoxide β-carbon by the threonine α-amino group, yielding a stable oxazepano adduct that is catalytically inactive.^12^ This dual functionalisation mechanism, unique among electrophilic pharmacophores, confers exceptional selectivity for the proteasome over other non-Ntn proteases, underpinning the clinical efficacy of this natural product class.^8^

The translation of the peptidyl epoxyketone scaffold into therapeutic agents has been particularly impactful in oncology.^13^ Exploration of synthetic analogs of epoxomicin **4** led to the discovery of carfilzomib **7**, which received FDA approval for the treatment of multiple myeloma in 2012 (Fig. 1C).^14^ Further development resulted in the orally bioavailable epoxyketone Oprozomib **8**, which has been evaluated in phase Ib/II clinical trials for treatment of multiple myeloma (Fig. 1C).^15,16^ Epoxyketones have also gained attention for their potential to treat autoimmune disorders and parasitic diseases.^17–19^ A phase IIa trial of Zetomipzomib (KZR-616; **9**) in patients with autoimmune hepaptitis was completed in late 2024 (Fig. 1C).

Not with standing these advances, the stereoselective synthesis of α, β-epoxyketones is non-trivial. Current industrial-scale syntheses involve multistep procedures employing complex chiral catalysts and precise temperature control.^20–22^ This has motivated the search for alternative routes to the α, β-epoxyketone pharmacophore, employing greener and more sustainable catalysts and near-ambient temperatures.

Kaysser, Moore and co-workers reported the genetic basis for the biosynthesis of epoxomicin **4** and eponemycin **5b** in *Goodfellowia coeruleoviolacea* ATCC 53904 and *Streptomyces hygroscopicus* ATCC 53709, respectively,^23^ although sequencing errors hampered subsequent attempts to elucidate the roles played by individual biosynthetic genes in the assembly of the latter.^24, 25^ We identified a cryptic gene cluster in *Streptomyces chromofuscus* ATCC49982 that directs the biosynthesis of TMC-86A **5a**,^26^ which is structurally closely related to eponemycin (Fig. 1B). This gene cluster shows a very high degree of similarity to the eponemycin biosynthetic gene cluster (BGC) in *S. hygroscopicus*.

The *epnF* gene was proposed, based on co-expression with *epnG* and *epnH*, which encode a bimodular nonribosomal peptide synthetase (NRPS) and a monomodular polyketide synthase (Fig. 2), to encode the key epoxyketone forming enzyme.^27^ Using synthetic putative substrates, we directly demonstrated that EpnF, an unusual multifunctional flavoenzyme, catalyzes the conversion of leucine-containing analog of α-dimethyl-β-ketoacid **10a** to the corresponding analog of epoxyketone **10b** (Fig. 2).^26^ We also showed that the cytochrome P450 (CYP) TmcI catalyzes hydroxylation of the methyl group appended to the epoxyketone in the leucine-containing analog of **10b** (Fig. 2).^25. 26^

**Figure 2.**
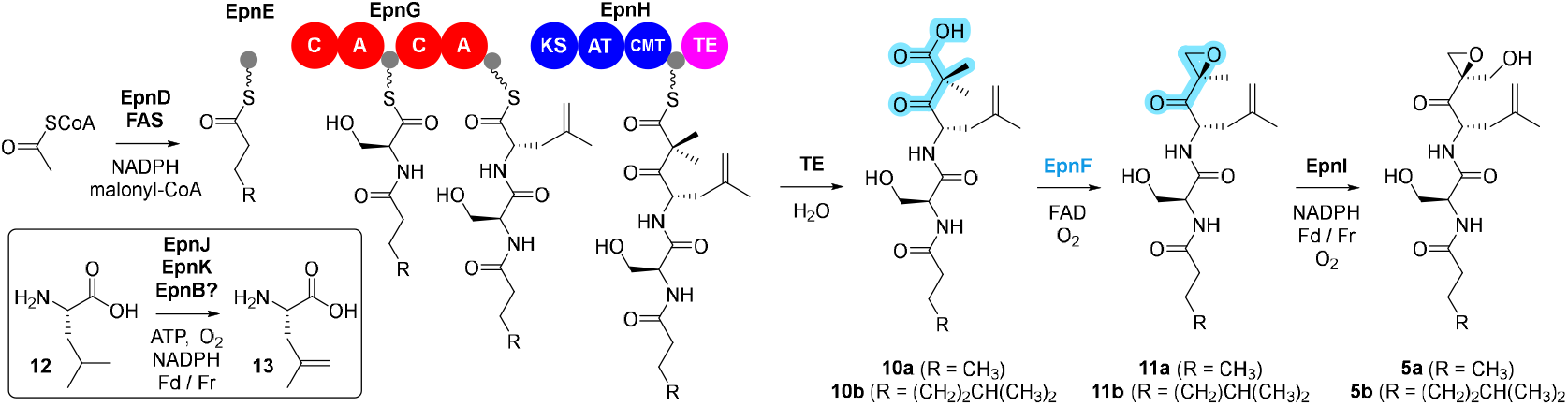
Pathway for the biosynthesis of TMC-86a **5a** and eponemycin **5b** in *Streptomyces hygroscopicus* ATCC 53709. EpnF-catalyzed conversion of the α-dimethyl-β-ketoacid of key intermediate **10a** to the corresponding epoxyketone in **11a** is highlighted in cyan. This and the catalytic function of EpnI otholog TmcI were demonstrated experimentally using a synthetic analog of **10a** containing a leucine residue in place of the 4,5-dehydrolecuine residue.

We subsequently developed improved vectors for direct BGC capture from genomic DNA using transformation-associated recombination (TAR) in yeast.^25^ This enabled us to capture the eponemycin and TMC-86A BGCs from genomic DNA of *S. hygroscopicus* and *S. chromofuscus*, respectively.^25^ Expression of these gene clusters in *S. albidolflavus* J1704 (formerly known as *S. albus*) showed that the *S. hygroscopicus* BGC directs the production of TMC-86A, in addition to eponemycin and several congeners.^25^ By exploiting the extremely high efficiency of homologous recombination in yeast, we developed a platform for parallelized BGC editing and applied it to elucidating the functions of several further genes in the eponemycin BGC.^25^ These experiments (i) confirmed that the TmcI homolog encoded by *epnI* catalyses hydroxylation of the methyl group appended to the epoxyketone; (ii) showed that *epnD* and *epnE*, encoding homologs of FabH (a ketosynthase initiating chain assembly in bacterial fatty acid biosynthesis) and an acyl carrier protein (ACP), respectively, play key roles in the assembly of the *N*-butanoyl group of TMC-86A **5a**; and (iii) demonstrated that *epnJ* and *epnK*, encoding a monomodular NRPS and a CYP, respectively, are involved in the conversion of Lleucine to L-4,5-dehydroleucine, which is subsequently incorporated into eponemycin **5b** / TMC-86A **5a** by EpnG (Fig. 2).

EpnF shows sequence similarity to acyl-CoA dehydrogenases (ACADs), flavoenzymes that catalyze α, β-desaturation of acyl-CoA thioesters in mitochondria.^28^ This proceeds via deprotonation of the acyl-CoA thioester α-carbon and concomitant transfer of hydride from the β-carbon to the flavin cofactor. Crystallographic and mutagenesis studies have identified the side-chain carboxylate group of a conserved glutamate residue (E376 in medium-chain ACADs) as the general base responsible for α-carbon deprotonation.^29,30^

By analogy, the EpnF-catalyzed reaction is proposed to proceed via decarboxylation of the *N*-acyl-dipeptidyl β-ketoacid to generate an enolate which collapses with transfer of hydride from the β-carbon of one of the branching methyl groups to the bound FAD cofactor.^26^ The resulting FADH2 then reacts with dioxygen to form the corresponding peroxide. This undergoes Michael addition to the β-carbon of the enone created by the hydride transfer and 1, 3-elimination of C4a-hydroxyflavin generates the epoxyketone, consistent with the proposed mechanism for epoxidation of electron-deficient alkenes by flavin monooxygenases.^31^ Elimination of water from C4a-hydroxyflavin regenerates FAD.

The use of natural product biosynthetic enzymes for production of high-value chemicals is gaining traction in the chemical industry, due to their sustainability and ability to catalyze highly chemo, regio and stereo-selective reactions under mild conditions.^32^ Biocatalysts have been employed in the synthesis of various anthropogenic chemicals, including (i) the HIV drug islatravir, which was assembled using only five engineered enzymes, more than halving the number of steps previously required;^33^ and (ii) the proton pump inhibitor esomeprazole and several statins, which are assembled using engineered Baeyer-Villiger monooxygenases.^34,35^ The ever decreasing cost of DNA synthesis is accelerating the discovery and development of novel biocatalysts, expanding the toolbox for chemoenzymatic approaches to the synthesis of high value chemicals.^36^ Consequently, the biocatalysis sector is expected to be worth $14.9 billion by 2027.^37^

A detailed understanding of the substrate scope and catalytic mechanism of EpnF and related epoxyketone synthases is important for assessing their potential to be developed into useful biocatalysts for enantiocontrolled epoxyketone assembly. To date, this has been hindered by the chemical instability of the α-dimethyl-β-ketoacid moiety, leading to the conclusion that EpnF has a relatively modest substrate scope.^38^ This moiety undergoes rapid decarboxylation to the corresponding ketone, which cannot be converted by the enzyme into epoxidized products.^26^

Here, we report the development and application of an enzymatic unmasking strategy to generate α-dimethyl-β-ketoacid substrates of EpnF *in-situ*. This enabled us to probe a wide range of synthetic substrate analogs with variations in the *N*-acyl cap and both side chains of the dipeptidyl moiety. Our results demonstrate that EpnF is able to convert a wide range of dipeptidyl-dimethyl-β-ketoacids to the corresponding epoxyketones, including compounds with readily removable *N*-terminal protecting groups, which could be employed in the synthesis of products with longer peptidyl chains. Substrate / intermediate docking and molecular dynamics (MD) simulations on an AlphaFold model provided insights into the catalytic mechanism of EpnF and identified active residues likely to be important for substrate positioning and/or catalysis. The role played by these residues was LC-MS analysis of epoxyketone product / enone intermediate formation using an analog of the native substrates probed via a combination of site directed mutagenesis and generated by enzymatic unmasking *in situ*.

## RESULTS AND DISCUSSION

### Development of *in-situ* substrate unmasking assay

To circumvent difficulties caused by the instability of the αdimethyl-β-ketoacid moiety, we investigated producing it *in situ* via hydrolysis of the corresponding methyl ester using commercially available pig liver esterase (PLE). Analog **17** of the α-dimethyl-β-ketoacid intermediate **11b** in eponemycin biosynthesis was synthesized to explore this. Condensation of *N*-Boc-L-leucine **14** with methyl potassium malonate (MPM) using carbonyl diimidazole (CDI) and magnesium chloride afforded the corresponding β-keto ester, which was α-dimethylated with methyl iodide and potassium carbonate to yield **15**. The *t*-butoxycarbonyl (Boc) group was removed from **15** using hydrochloric acid and the resulting amine hydrochloride salt was coupled with *N*Boc-L-serine using HATU and Hünig’s base to afford **16**. Removal of the Boc group from **16** using hydrochloric acid followed by *N*-acylation with heptanoic acid using HATU and Hünig’s base gave **17**. (Scheme 1).

**Scheme 1.**
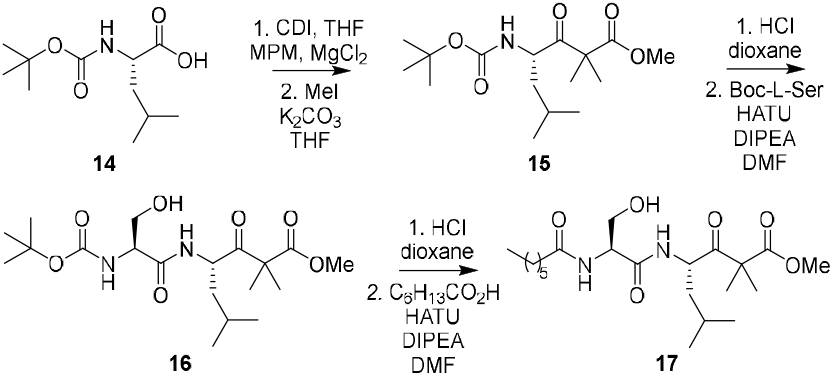
Synthesis of α-dimethyl-β-keto ester **17**, a stabilised analog of eponemycin biosynthetic intermediate **10b**.

EpnF was overproduced in *E. coli* as an N-terminal octahistidine fusion and purified as described previously.^26^ α-Dimethyl-β-keto ester **17** was incubated with PLE and FAD in the presence and absence of EpnF at 30 °C for 3 hours. The reactions were quenched by addition of acetonitrile containing 0.1% formic acid and analyzed by liquid chromatography-mass spectrometry (LC-MS). Species with *m/z* values corresponding to both the α-dimethyl-β-keto-acid **18**, resulting from PLE-catalyzed hydrolysis of the methyl ester, and isopropyl ketone **19**, the spontaneous decarboxylation product of **17**, were formed in the reaction lacking EpnF. In the reaction containing EpnF, the formation of α-dimethylβ-keto-acid **18** and isopropyl ketone **19** was strongly suppressed, and a new product appeared with an *m/z* corresponding to epoxyketone **20** (Fig. 3). These data show that EpnF converts the majority of α-dimethyl-β-ketoacid **18** generated by PLE-catalyzed hydrolysis of methyl ester **17** to epoxyketone **20**, confirming the suitability of the *in-situ* unmasking strategy for assessing the substrate scope of EpnF.

**Figure 3.**
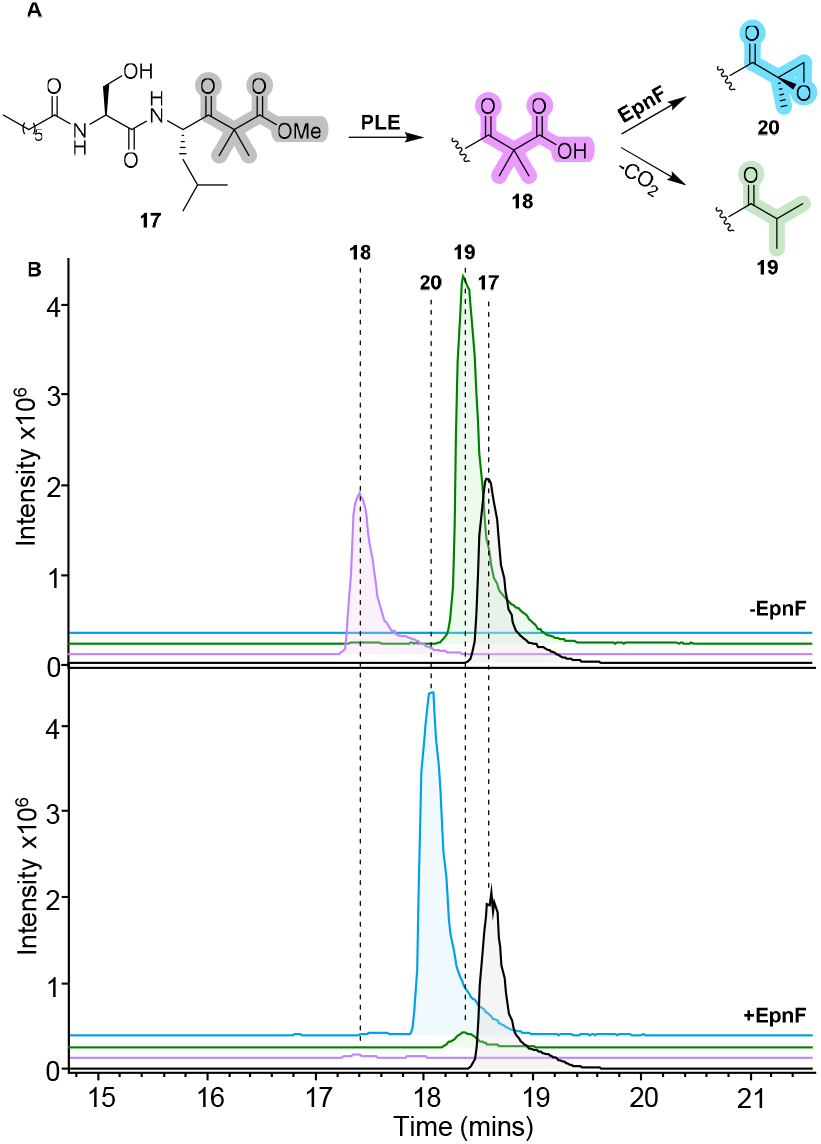
(**A**) Concept of *in-situ* substrate unmasking assay. The unstable α-dimethyl-β-keto-acid **18** is generated via slow hydrolysis of the corresponding methyl ester **17** by pig liver esterase (PLE). This enables EpnF to convert the majority of **18** to the epoxyketone **20**, minimizing spontaneous decarboxylation to form the dead-end shunt product **19. (B**) LC-MS chromatograms from reactions employing PLE to unmask **17** in the absence (top) and presence (bottom) of EpnF.

### Exploration of EpnF substrate scope

To probe the substrate tolerance of EpnF, we synthesized an analog library of **17** with variations in the amino acid side chains and *N*-acyl group (Fig. 4). Analogs with alterations to the amino acid side chains were synthesized by preparing various γ-amino-α-dimethyl-β-keto methyl esters, via condensation of the corresponding Boc-protected amino acids with MPM, followed by dimethylation and deprotection. These were then coupled with *N*-butanoyl-L-amino acids (Scheme 2), derived from the hydrochloride salts of the corresponding amino acid methyl esters, via *N*-acylation and ester hydrolysis. Analogs bearing *N*-terminal *n*-butyl, acetyl, and pivaloyl groups were prepared via the route used for the synthesis of **17** (Scheme 1), by substituting the acid used for *N*-acylation in the final step.

**Scheme 2.**
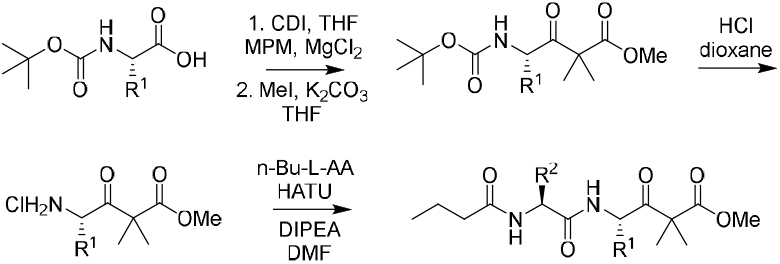
Synthesis of methyl ester pro-substrates of bio-synthetic intermediate **10a** with altered amino acid side chains.

**Figure 4.**
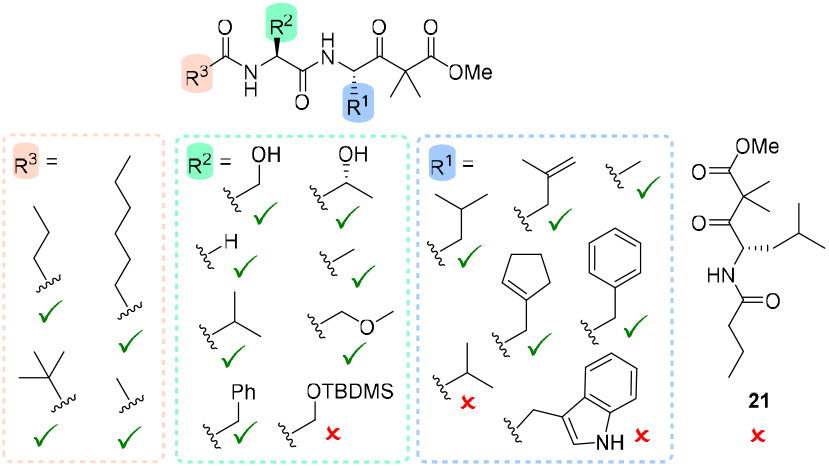
Overview of analogs of **17** synthesized and examined for turnover to the corresponding epoxyketones in the *in-situ* unmasking assay. A tick / cross indicates whether epoxyketone formation was observed for the analog containing the indicated structural change. A mono-peptidyl analog **21** with the Ser residue omitted and R^3^ = n-propyl was not turned over to the corresponding epoxyketone.

The ability of EpnF to turnover these analogs was evaluated using the *in-situ* substrate unmasking assay. This showed that the enzyme tolerates analogs with a wide range of variations in R^2^ (Fig. 4 and S1), corresponding to the side chain of the L-Ser residue in the native substrates.

This was surprising given that all naturally occurring epoxyketones, except for carmaphycins, have a β-hydroxy group in the side chain corresponding to R^2^. No turnover was observed for the analog with the hydroxymethyl side chain protected as a TBS ether, indicating that very large R^2^ substituents cannot be accommodated by the active site of the enzyme (Fig. 4 and S1).

A similar pattern was observed for analogs with variations to R^1^, corresponding to the side chain of the L-4, 5-dehydro-Leu residue in the native substrates (Fig. 4 and S2). Methyl, benzyl, and (cyclopentenyl)methyl groups, corresponding to the side chains in naturally occurring clarepoxcin, and synthetic oprozomib and zetomipzomib, respectively, were all tolerated. Analogs with an indolylmethyl group, corresponding to the side chain of tryptopeptin, and an isopropyl group were not converted to epoxyketones, however, indicating the EpnF active site places some limits on the size of R^1^ groups that can be accommodated. This is consistent with the very low levels of tryptopeptin **6** relative to congeners lacking the epoxyketone observed in native producers and heterologous expression hosts.^39^

In addition to the *N*-butanoyl group incorporated into intermediate **10a** in the biosynthesis of TMC-86A,^25^ and the *N*heptanoyl group in analog **17** used to develop the *in-situ* substrate unmasking assay, *N*-pivaloyl and *N*-acetyl groups were also tolerated by EpnF (Fig. 4 and S3). This suggests that the nature of the *N*-acyl group does not place stringent limits on the enzyme’s substrate scope.

Collectively, these data indicate that significant variation in R^1^, R^2^ and the *N*-acyl group can all be tolerated by EpnF, suggesting that the dipeptidyl backbone is the minimal epitope for recognition by the enzyme’s active site. We thus synthesized a further analog containing a single *N*-acylated leucine residue **21** and the *N*-butanoyl group of intermediate **10a** in TMC-86A biosynthesis. EpnF was unable to convert this to the corresponding epoxyketone (Fig. 4 and S4), consistent with this hypothesis.

Inspired by the broad tolerance of EpnF to various *N*-acyl groups, we synthesized further analogs of **17** bearing *N*-terminal Boc, Alloc, Fmoc, and Cbz groups. This enabled us to examine the feasibility of a chemoenzymatic approach to dipeptidyl epoxyketones that can be deprotected and *N*-acylated with a wide range of peptidyl (and other) moieties. All four of these analogs were converted to the corresponding epoxyketones (Fig. 5A and S5).

**Figure 5.**
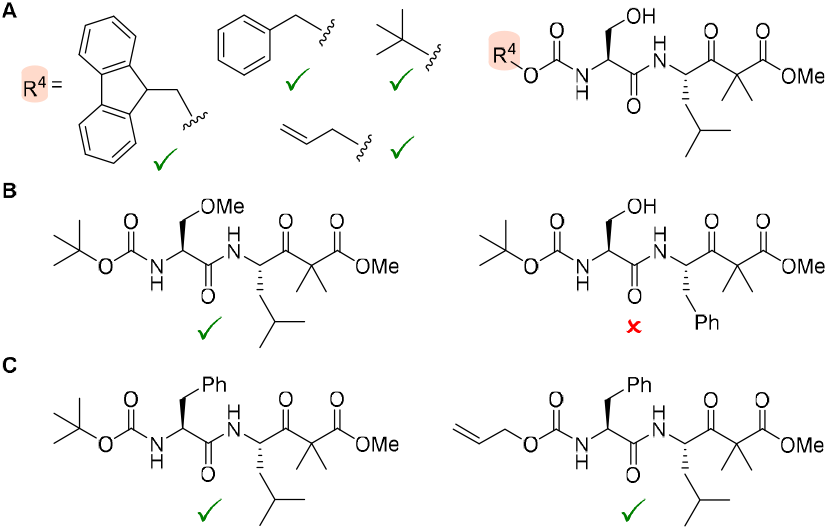
(**A**) Turnover of analogs of **17** with the *N*-heptanoyl group substituted by *N*-terminal protecting groups commonly employed in peptide synthesis to the corresponding epoxyketones. (**B**) Turnover of oprozomib-related *N*-protected dipeptidyl-α-dimethyl-β-ketoacid methyl esters to the corresponding epoxyketones. (**C**) Turnover of carfilzomib-related *N*-protected dipeptidyl-α-dimethyl-βketoacid methyl esters to the corresponding epoxyketones. In all cases the standard in-situ unmasking assay was used.

Encouraged by these results, we next examined *N*-protected analogs of **17** with R^1^ and/or R^2^ groups corresponding to those incorporated into equivalent positions of oprozomib and carfilzomib (Fig. 5). Although a Boc-protected analog with R^2^ = methoxymethyl was converted to the corresponding epoxyketone, no turnover of an analog bearing a Boc group with R^1^ = benzyl was observed (Fig. 5B and S6). Thus, a chemoenzymatic approach to oprozomib may require directed evolution of EpnF to further broaden its substrate scope. On the other hand, both Boc- and Alloc-protected analogs with R^2^= benzyl and R^1^ = isobutyl were converted to the corresponding epoxyketones by EpnF (Fig. 5C and S7). Turnover of the Boc-protected analog is particularly noteworthy, because it is known that Boc groups can be efficiently removed from related compounds without detriment to the epoxyketone.^21^ These results indicate that it should be feasible to develop a chemoenzymatic synthesis of carfilzomib.

Overall, our data show that EpnF is a promising biocatalyst for epoxyketone synthesis with a broad substrate scope. To provide a basis for rational engineering to further expand substrate tolerance, we turned our attention to developing a better understanding of the structure and catalytic mechanism of EpnF.

### Sequence and structural comparison to ACADs and related enzymes

Although epoxyketone synthases show sequence similarity to ACADs, the structural commonalities and differences between these two enzyme families remain to be explored. The catalytic mechanism of ACADs is proposed to involve deprotonation of the acyl-CoA substrate at the α-carbon by a conserved active site Glu residue and concomitant hydride transfer from the β-carbon to the oxidized flavin cofactor (Fig. 6A).^29, 30^

**Figure 6.**
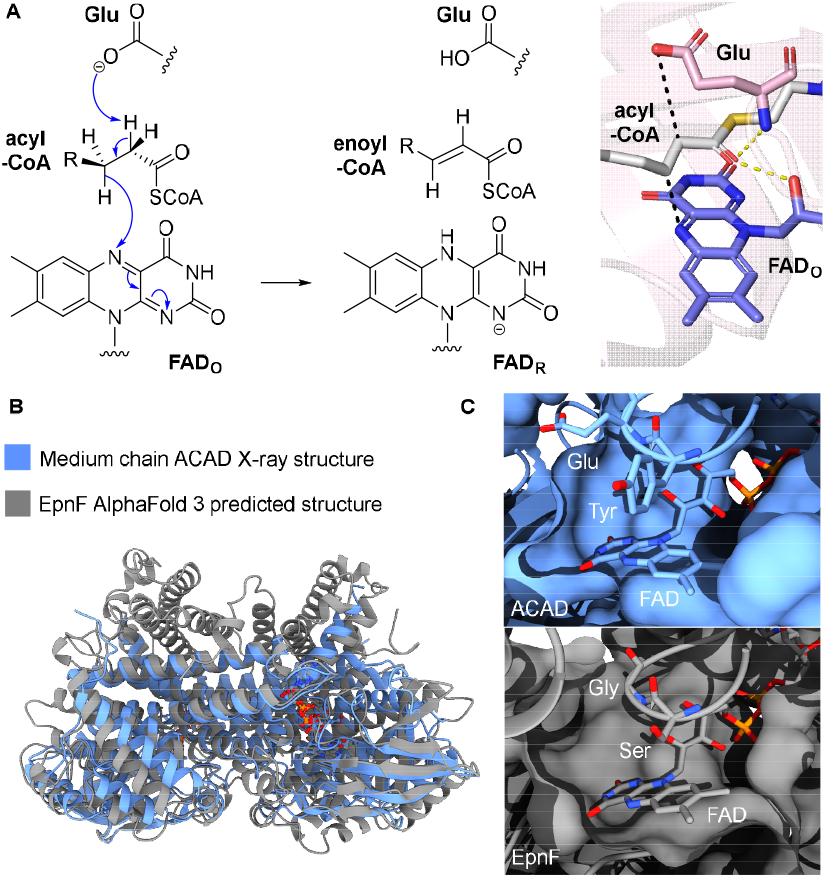
(**A**) Proposed catalytic mechanism of ACADs, involving concomitant deprotonation of the α-carbon of the acyl-CoA substrate by a conserved active site Glu residue and hydride transfer from the β-carbon to N5 of the oxidized flavin cofactor (FAD_O_), resulting in the formation of an enoyl-CoA thioester and reduced flavin (FAD_R_). The right panel shows the X-ray crystal structure of an MCAD with acyl-CoA substrate bound (PDB ID: 3MDE). (**B**) Overlay of the AlphaFold3 model of EpnF and the X-ray crystal structure of the medium chain ACAD. (**C**) Comparison of the active site architecture of the medium chain ACAD with that predicted by AlphaFold3 for EpnF, highlighting the flavin cofactor in both and the Tyr-Glu dyad in the former, which is replaced by a conserved Ser-Gly dyad in the latter.

Acyl-CoA oxidases (ACOs) catalyze analogous reactions to ACADs but in peroxisomes rather than mitochondria and use dioxygen as the terminal electron acceptor instead of an electron transferring flavoprotein.^40^ Given that the epoxyketone synthase catalytic mechanism is proposed to involve transfer of electrons from FADH_2_ to dioxygen,^26^ one might expect these enzymes to be closely related to ACOs. Phylogenetic analysis shows that ACADs segregate according to substrate chain length specificity,^41^ whereas ACOs form a single clade, and that epoxyketone synthases are related most closely to bacterial very long-chain ACADs (Fig. SX).

Attempts to obtain diffraction quality crystals of EpnF were unsuccessful. Thus, to gain insight into the structural differences between epoxyketone synthases and ACADs, we created AlphaFold models of the EpnF homodimer (previously shown by gel filtration to be the oligomerization state in solution) and shown to overlay well with the X-ray crystal structures of ACADs (Fig. 6B and S16). Notably, the conserved active site Tyr and Glu residues in ACADs are substituted by Ser and Gly residues, respectively, in EpnF (Fig. 6C).^29, 30^

### Investigation of EpnF catalytic mechanism via substrate docking and MD simulations

We previously hypothesized that epoxyketone synthases catalyze decarboxylative hydride transfer from the β-carbon of their α-(di)methyl-β-keto-acid substrates to the oxidized flavin cofactor (Fig. 7).^26^ The resulting enone intermediate is proposed to be epoxidized by a C4a-flavin peroxide, arising from the reaction of reduced flavin with dioxygen, via a Michael addition/1, 3-elimination cascade (Fig. 7).^26^

**Figure 7.**
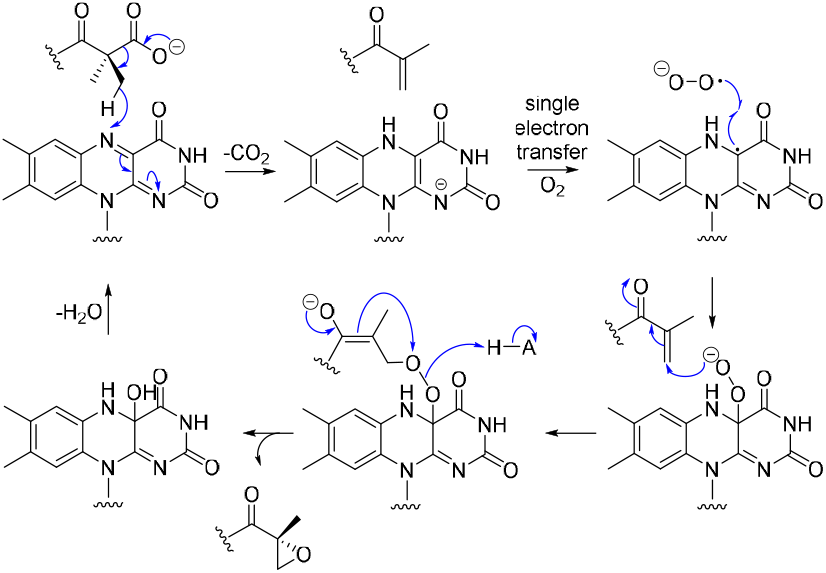
Proposed catalytic mechanism of EpnF.^26^

Species with molecular formulae corresponding to the proposed intermediate enone are observed in UHPLC-ESI-Q-ToF-MS analyses of EpnF-catalyzed epoxyketone formation. An authentic standard of the enone corresponding to an analog of **17** with R^1^= methyl and R^3^ = n-propyl was synthesized and found to have the same retention time as the putative enone intermediate in the corresponding enzymatic reaction (Fig. S8). However, this enone was not converted to an epoxyketone when incubated with EpnF containing a reduced flavin cofactor, indicating it cannot reenter the catalytic cycle once released from the active site.

To explore the proposed catalytic mechanism of EpnF further, we conducted a 100 ns molecular dynamics (MD) simulation on an AlphaFold 2 model of the enzyme into which FAD and substrate **10b** had been manually docked such that the *pro-S* and *pro-R* methyl groups were positioned a similar distance from the flavin. In the simulation, The *pro-S* methyl group quickly moves towards the flavin cofactor and remains in close proximity to it throughout the simulation (Fig. 8A). This is likely because the ribityl hydroxyl groups of the cofactor donate hydrogen bonds to the carboxyl group of **10b**, stabilizing the conformation that positions the *pro-S* methyl group closest to the flavin (Fig. 8B). Thus, we propose hydride is transferred from the *pro-S* methyl group of **10b** to the flavin cofactor in EpnF. Substitution of the conserved Glu residue serving as the general base in ACADs with Gly in EpnF enables the carboxyl group in **10b** to be oriented such that decarboxylation and hydride transfer can be coupled (Fig. 8B). This mirrors the catalytic mechanism of medium-chain ACADs, which has been shown by kinetic isotope effect studies to involve synchronous α- and β-C–H bond fission.^42^ However, we cannot rule out a transient enolate intermediate in the EpnF-catalyzed reaction.

**Figure 8.**
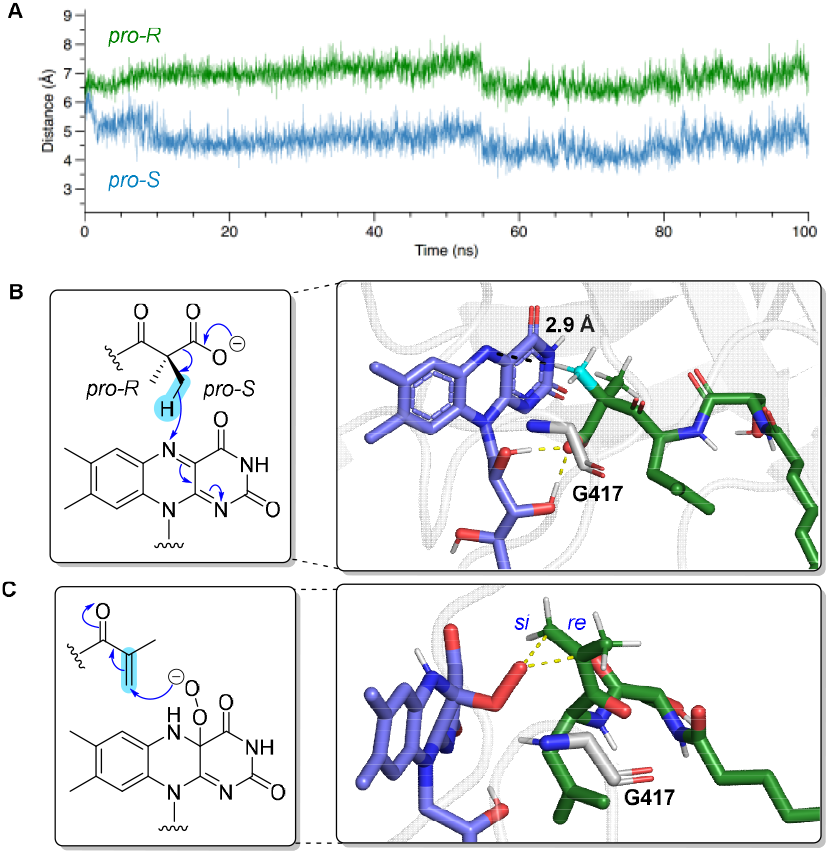
(**A**) Plot of distance between the FAD N5 atom and the carbon atoms of the *pro-S* and *pro-R* methyl groups over the course of the simulation. The mean values were 4.67 Å and 6.89 Å, respectively. (**B**) Concerted decarboxylative hydride transfer mechanism proposed for EpnF (left panel) and a representative frame from the corresponding MD simulation with the carboxyl group of the substrate hydrogen bonded to the ribityl hydroxyl groups of the cofactor and the *pro-S* methyl group positioned close to N5 of the flavin (right panel). (**C**) Left panel: proposed Michael addition of the C4a-peroxyflavin, arising from the reaction of dioxygen with reduced flavin (generated by hydride transfer from the substrate), to the observed enone intermediate in epoxyketone formation. Right panel: A representative frame from the corresponding MD simulation showing the distal oxygen atom of the C4a-peroxyflavin positioned over the *si*-face of the enone intermediate.

One interesting question is whether N1 of FAD gets protonated during the decarboxylative hydride transfer reaction. MD simulations of EpnF containing FADH− showed no suitable proton donors in the vicinity of the N1 atom. Instead, several residues able to donate hydrogen bonds appear to stabilize the negative charge on N1 by creating an oxyanion hole-like environment (Fig. 17).

Having established the feasibility of the enone intermediate in our previously proposed mechanism,^26^ we turned our attention to epoxidation of this intermediate by C4a-peroxy-flavin (proposed to result from reaction of dioxygen with the reduced flavin cofactor). The enone intermediate derived from **10b** and C4a-peroxy-FAD were manually modelled into the EpnF active site, and a 100 ns MD simulation was conducted. The β-carbon of the enone remained near the distal (negatively charged) oxygen atom of the peroxyflavin throughout the simulation, consistent with the previously proposed Michael addition / 1, 3-elimination mechanism for epoxide formation (Fig. 8C).^26^ Significantly, the distal oxygen atom is positioned over the *si*-face of the enone C=C (Fig. 8C), consistent with the 2*R* epoxide stereochemistry in epoxyketone natural products (Fig. 1B).^5-7^

The MD simulations indicate that both formation of the enone intermediate and its epoxidation by C4a-peroxyflavin are feasible steps in epoxyketone formation. Thus, we determined which residues engage in persistent polar contacts with the substrate and enone intermediate during the simulations. The carboxylate group of the substrate accepts hydrogen bonds from the ribityl hydroxyl groups of FAD, analogous to hydrogen bonding observed between the oxygen atom of the thioester group and the ribityl hydroxyl groups in ACADs.^30^ In addition, the guanidinium group in the side chain of the Arg283 residue donates hydrogen bonds to the oxygen atoms of the keto group and the N-acyl group of the substrate (Fig 9A). Similarly, the guanidinium group in the side chain of the Arg283 residue donates hydrogen bonds to the oxygen atoms in the same two functional groups of the enone intermediate (Fig. 9B). Additional hydrogen bonds are formed between the side chain hydroxyl groups of Ser158 and Ser416 and the carbonyl group of the L-serine residue in the enone intermediate and the distal oxygen atom of the C4a-peroxy-flavin, respectively (Fig. 9B).

**Figure 9.**
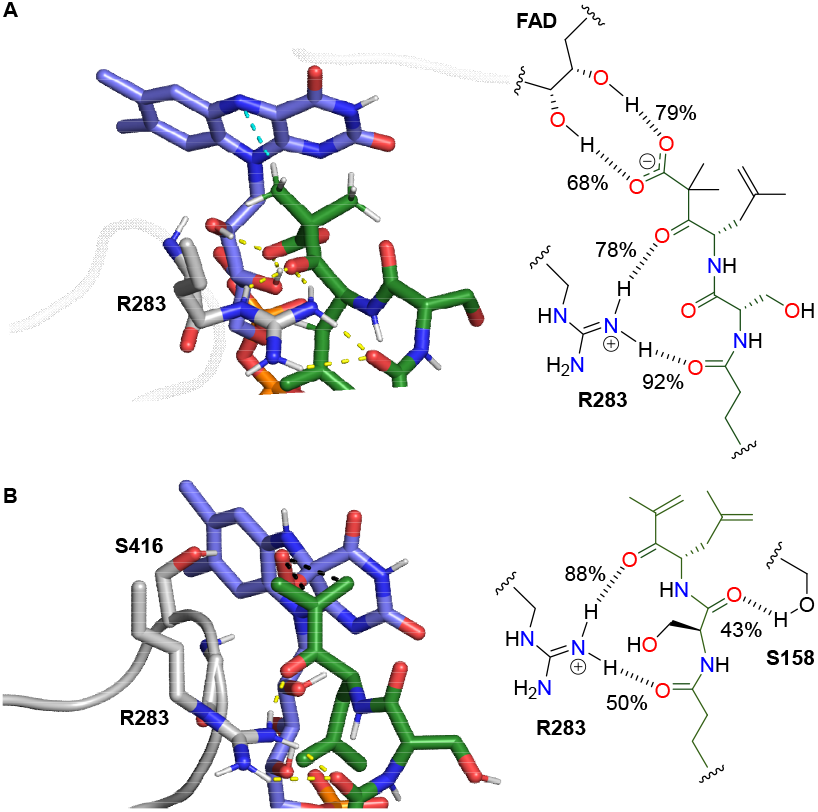
(**A**) Left panel: Frame from MD simulation of FAD and substrate **10b** bound in the active site of EpnF, showing the Arg283 side chain and the ribityl hydroxyl groups of the cofactor donating hydrogen bonds to two of the carbonyl groups and the carboxyl group, respectively, in the substrate. Right panel: schematic highlighting these key polar contacts and showing the percentage of frames containing them. (**B**) Left panel: Frame from MD simulation of 4a-per-oxy-FAD and the enone intermediate derived from **10b** bound in the active site of EpnF, showing the Arg283 and Ser158 side chains donating hydrogen bonds to three of the carbonyl groups in the intermediate and the Ser416 side chain donating a hydrogen bond to the distal oxygen atom of the 4a-peroxyflavin. Right panel: schematic highlighting the key polar contacts between the enone intermediate and the side chains of Arg283 and Ser158 and showing the per-centage of frames containing them.

**Figure 10.**
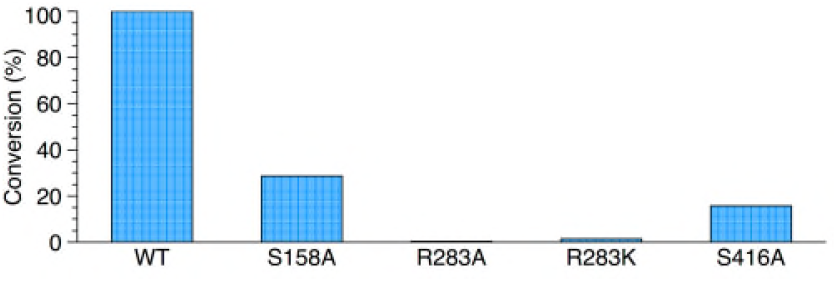
Percentage epoxyketone formation relative to wild type (WT) by S158, R283, and S416 mutants of EpnF.

### Mutagenesis of residues implicated in substrate binding and catalysis

To experimentally investigate the roles played by EpnF residues implicated in hydrogen bonding to the substrate / C4a-peroxy-flavin distal oxygen atom by the MD simulations, we used site-directed mutagenesis. R283A, R283K, S416A, and S158A mutants were created and overproduced in *E. coli* as soluble recombinant proteins that were purified using Ni^2+^-IMAC (Fig. S19, ). The purified proteins were confirmed to be correctly folded using CD spectroscopy and to contain the desired mutation by UHPLC-ESI-Q-ToF-MS analysis (Fig. S20 and S21).

Turnover of an analog of **17** bearing a Cbz group in place of the heptanoyl group by the mutants was assessed using the *in-situ* substrate unmasking assay. Epoxyketone formation was strongly suppressed in reactions containing either the R283A or R283K mutant, consistent with R283 playing a key role in binding of the substrate and orienting it relative to the flavin cofactor, in addition to stabilization of transient negative charge on the oxygen atom of the keto group during the decarboxylative desaturation and epoxidation reactions. Significant suppression of epoxyketone formation was also observed in assays containing the S158A and S416A mutants, consistent with the involvement of these residues in binding the enone intermediate and orientation of the peroxy group for Michael addition to the enone, respectively. Interestingly, LC-MS analyses also revealed a significant increase in formation of the enone intermediate relative to the epoxyketone in the mutants (Fig. S22 and S23), providing further evidence that these residues contribute primarily to catalysis of the epoxidation reaction.

In addition to the essential role played by EpnF-like enzymes in peptidyl-epoxyketone biosynthesis, homologs have been reported to participate in the assembly of other natural product classes, such as the matlystatins and threopeptin. In matlystatin and threopeptin biosynthesis, the EpnF homologs MatG and SnaO have been proposed to catalyze the conversion of α-methyl-β-ketoacids to the corresponding enones (Fig. 11A).^43, 44^ Interestingly, in both MatG and SnoA, the residue corresponding to Ser416 of EpnF is mutated to Ala (Fig.11B), supporting the hypothesis that this Ser residue helps to orient the flavin-bound peroxy group for Michael addition to the enone intermediate.

**Figure 11.**
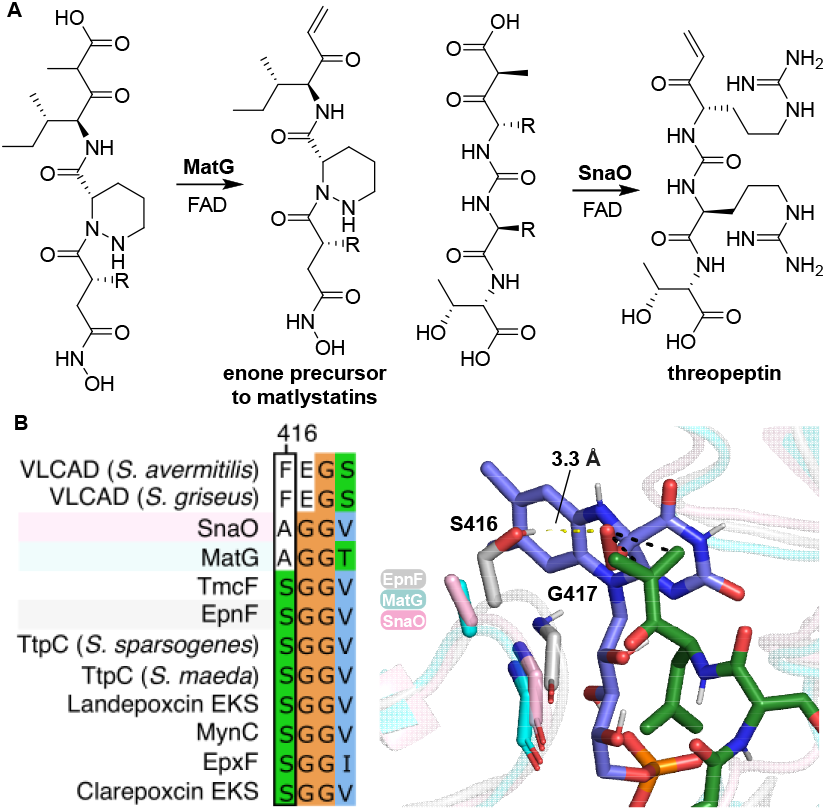
(**A**) Role played by EpnF homologs MatG and SnaO in the biosynthesis of the matlystatins and threopeptin, respectively. (**B**) Left panel: alignment of active site residues in ACADs, and epoketone synthases with MatG and SnaO. Residue 416 of EpnF is Ser in all epoxyketone synthases identified to date, whereas in MatG and SnaO the corresponding residue is Ala. Right panel: overlay of a frame from the MD simulation of the EpnF AlphaFold model with C4a-peroxy-FAD and the enone intermediate derived from **10b** bound in the active site with AlphaFold models of MatG and SnaO. In EpnF, S416 donates a hydrogen bond to the distal oxygen atom of the peroxy-flavin, stabilizing it and orienting it for Michael addition to the enone intermediate. Mutation of this residue to Ala in MatG and SnaO prevents the enone intermediates from being converted to the corresponding epoxyketones, presumably because C4a-peroxy-FAD undergoes rapid collapse to hydrogen peroxide and the oxidized flavin.

## CONCLUSIONS

The results reported here provide detailed insight into the substrate scope and catalytic mechanism of the α,βepoxyketone synthase EpnF. A major obstacle to such studies is the intrinsic instability of the native β-ketoacid substrates, which readily undergo spontaneous decarboxylation. By developing an esterase-mediated *in-situ* unmasking strategy, we were able to employ chemically stable α(di)methyl-β-ketoacid methyl esters to directly probe the substrate scope of EpnF. This revealed that EpnF is able to convert a wide range of *N*-acyl-dipeptidyl-α-dimethyl-βketo acids to the corresponding epoxyketones, demonstrating a broader substrate tolerance than previously inferred from studies relying on unstable β-ketoacids.^38^ Only compounds with much larger side chains than those incorporated into the dipeptidyl moiety of native substrates, or in which this moiety is replaced with a single amino acid, failed to turnover. The ability of EpnF to accept substrates bearing *N*-terminal protecting groups commonly employed in peptide synthesis (Boc, Cbz, Fmoc, and Alloc), including a Boc-protected progenitor of the *C*-terminal dipeptidylepoxyketone fragment of Carfilzomib, is particularly noteable. Overall, our observations suggest EpnF and related epoxyketone synthases show promise for development into useful biocatalysts that could be employed for chemoenzymatic synthesis of structurally diverse epoxyketone proteasome inhibitors.

Sequence comparison of EpnF with canonical ACADs indicated that the conserved active site Glu residue in the latter is replaced by glycine in epoxyketone synthases. This appears to allow α-(di)methyl-β-keto acids to bind in the active site, enabling decarboxylative transfer of hydride from one of the α-methyl groups to the flavin cofactor. Molecular dynamics simulations of on an AlphaFold 2-derived model of EpnF with flavin and an eponemycin biosynthetic intermediate bound support this notion and indicate a strong preference for hydride transfer from the *pro-S* methyl group. This is testable hypothesis that could be probed in future experiments via stereospecific ^2^H or ^13^C-labeling of one of the α-methyl groups. The enone intermediate arising from decarboxylative hydride transfer to the flavin cofactor was experimentally observed by LC-MS in incubations of EpnF with its native substrates and analogs, and in one case the identity of this intermediate was confirmed by comparison with synthetic standard. Additional simulations were conducted with C4a-peroxy-FAD, produced by reaction of the FADH_2_ resulting from hydride transfer to the flavin with dioxygen, and the enone intermediate bound in the EpnF active site. These were consistent with epoxidation of the *si* face of the enone via Michael addition of the distal oxygen atom of the C4a-peroxy-flavin and subsequent 1, 3-elimination of C4a-hydroxyflavin from the resulting enolate. This gives rise to the *2R* epoxide stereochemistry observed in all naturally occurring peptidyl epoxyketones.

The Arg283, Ser158, and Ser416 residues of EpnF predicted by the MD simulations to engage in persistent polar interactions with either the substrate and/or enone intermediate, were shown by site-directed mutagenesis to be important for epoxyketone formation. All catalytic activity was strongly suppressed in R283A and R238K mutants, whereas epoxyketone production was supressed and accumulation of the enone intermediate was elevated in S158A and S416A mutants. These data are consistent with (i) Arg283 playing a key role in binding the substrate to the active site, (ii) stabilisation of the C4a-flavin-peroxide intermediate and orientation for enone epoxidation via hydrogen bonding to the Ser416 side chain, and (iii) correct positioning of the enone intermediate for epoxidation via hydrogen bonding to the Ser158 side chain. The observation that the residue corresponding to Ser416 in MatG and SnaO, homologs of EpnF proposed to convert α-methyl-β-keto acid intermediates in matlystatin and threopeptin biosynthesis, respectively, to the corresponding enones, is mutated to Ala provides further support for the proposed role of Ser416. While this manuscript was in preparation, a few weeks after we completed all of our experimental work and data analysis,^45^ the X-ray crystal structure of EpxF, a homolog of EpnF involved in epoxomicin biosynthesis, was reported.^46^ An overlay of this structure with our Alphafold2 model of EpnF containing manually placed FAD shows excellent agreement (Fig. S24), further validating our use of this model as a starting point for the MD simulations. Using an indirect proteasome inhibition-based assay, it was shown that mutation of the Arg273 and Ser406 residues in EpxF (corresponding to Arg283 and Ser416 in EpnF) to Ala strongly suppresses epoxyketone formation. Further studies employing labelled substrates and ^13^C NMR spectroscopy revealed that the R273A mutant loses all catalytic activity, whereas the S406A mutant is still able to produce the enone intermediate. These observations are consistent with the catalytic roles we propose for the corresponding residues in EpnF. Biomimetic studies by the same authors, employing simplified FAD and substrate analogues,^46^ were fully consistent with the mechanism for EpnF-catalyzed epoxyketone formation we originally proposed,^26^ and have further characterised here.

In conclusion, EpnF is the founding member of a new family of internal flavoprotein monooxygenases, responsible for the introduction of epoxyketone and enone functional groups into diverse metabolites. The ability to distinguish enzymes belonging to this family from conventional ACADs based on a pair of key active site residues, and to use the nature of one of these residues to further predict whether a particular enzyme primarily catalyzes epoxyketone or enone formation, will greatly facilitate more accurate functional annotation of ACAD-like enzymes encoded by BGCs and genome mining approaches for the targeted discovery of novel enone and epoxyketone-containing specialized metabolites.

## ASSOCIATED CONTENT

### Supporting Information

Description of methods used, and supplementary figures and tables. This material is available free of charge via the Internet at http://pubs.acs.org..

## AUTHOR INFORMATION

### Notes

The authors declare the following competing financial interests: G.L.C. is a non-executive director, shareholder, and consultant of Erebagen Ltd.

## ACKNOWLEDGMENTS

We thank Dr Nikola Chmel for assistance with measuring CD spectra. CJB and J.C.W, were supported by the BBSRC Midlands Integrative Biosciences Doctoral Training Partnership (grant refs BB/J014532/1 and BB/T00746X/1). M.L.R. was supported by the EPSRC Doctoral Training Centre in Synthetic Biology (grant ref EP/L016494/1). M.P. and A.J.M. were supported by a grant (ref App39520) and a Discovery Fellowship (grant ref BB/X010619/1), respectively, from the BBSRC.

