## Supporting Information for "Substrate scope and catalytic mechanism of α, β-epoxyketone synthase EpnF illuminated by *in situ* esterase-mediated deprotection"

#### Supplementary figures and tables

### 1 Chemical, Biochemical, Biophysical and Computational Methods

#### 1.1 Chemical procedures

##### 1.1.1 Chemical materials and instrumentation

All commercially available chemicals, reagents, and solvents were purchased from Acros Organic, Fischer Scientific, Sigma Aldrich, Fluorochem or VWR and used without further purification. All aqueous solutions were prepared with deionised water. Ambient temperature refers to laboratory temperatures of 20-22 °C, 0 °C refers to an ice water bath, and heating of experiments was achieved with thermostatically controlled oil baths. Thin-layer chromatography analysis (TLC) was conducted using aluminium-backed plates pre-coated with Merck silica gel 60 F254 and compounds were visualised using UV radiation and/or potassium permanganate stain (10 g/L $KMnO_4$, 50 g/L $K_2CO_3$, 0.625 g/L NaOH). Concentration of samples in *vacuo* was achieved by evaporation of solvents on a BUCHI Rotavapor R-200 or R-210 connected to a BUCHI Vacuum Pump V-700. Silica column chromatography was performed on 40-63 μM, 40-60 Å silica gel (Sigma Aldrich).

NMR spectra were recorded on Bruker Advance AV-300 and HD-400 MHz spectrometers. Chemical shifts are reported in parts per million (ppm) referenced from CDCl3 (δH: 7.26 ppm and δC: 77.16 ppm) or MeOD (δH: 3.31 ppm and δC: 49.0 ppm) and assignments were made with the aid of COSY, HMBC, and HSQC spectra. Coupling constants (*J*) are rounded to the nearest 0.5 Hertz (Hz) and multiplicities are given as multiplet (m), singlet (s), doublet (d), triplet (t), quartet (q), quintet (quint.), sextet (sext.), septet (sept.), or combinations thereof.

High resolution mass spectra of small molecules were recorded using UHPLC- ESI-Q-TOF-MS using a Bruker Compact instrument coupled to Dionex Ultimate 3000 HPLC fitted with an Zorbax Eclipse Plus C18 column reverse phase column (100 x 2.1 mm, 1.8 μm), 30 °C).

#### 1.2 General synthetic procedures

###### 1.2.1.1 General procedure 1 – EDC mediated coupling

Procedure adapted from Zabala *et al*. Under an atmosphere of argon, $Et_3N$ (1.2 equiv.) was added to a solution of amino acid methyl ester (1 equiv.) in $CH_2Cl_2$ (0.2 mM) on ice. After 10 minutes, EDC hydrochloride (2.0 equiv.) and butyric acid (1.2 equiv.) was added, the reaction mixture was allowed to warm to ambient temperature and left to stir overnight. The reaction was quenched by addition of 1 M aqueous HCl, the layers were separated, and the aqueous layer extracted three times with EtOAc. The combined organic layers were dried with $MgSO_4$, filtered, and concentrated *in vacuo*. The resulting residue was purified by silica gel chromatography to give the desired product.[1]

###### 1.2.1.2 General procedure 2 – LiOH/$H_2O_2$ hydrolysis

Procedure adapted from Zabala *et al*. To a stirred solution of substrate (1 equiv.) in THF (2:1 THF:$H_2O$) was added 30% (w/w) hydrogen peroxide (0.22 equiv.) and 1 M lithium hydroxide solution (3 equiv.) dropwise. The THF was removed *in vacuo* after 2 hours. The remaining aqueous phase was washed twice with diethyl ether, acidified to pH 1 with 1 M hydrochloric acid, and sodium chloride was added until the solution was saturated. The solution was then extracted three times with EtOAc, and the combined organic phases dried over $MgSO_4$. The solvent was removed *in vacuo* to yield the desired product.[1]

###### 1.2.1.3 General procedure 3 – Methyl potassium malonate condensation

Procedure adapted from Ričko *et al*. To a stirred solution of CDI (1.5 equiv.) in anhydrous THF (0.2 mM), *N*-Boc-L-amino acid (1.0 equiv.) was added under argon and stirred at ambient temperature. After 2 h, $MgCl_2$ (1.5 equiv.) and methyl potassium malonate (1.5 equiv.) were added and the solution stirred for 24 h at ambient temperature. Solvent was removed in *vacuo* and the residue partitioned between EtOAc and 1 M HCl. After collecting the organic phase, the aqueous phase was extracted three times with EtOAc. The combined organic phases were washed twice with 5% (*w/v*) $NaHCO_3$ and brine then dried with $MgSO_4$ before solvent was removed in *vacuo*.[2]

###### 1.2.1.4 General procedure 4 – Methyl iodide dimethylation

Procedure adapted from Pettit *et al*. To a solution of β-keto-ester (1.0 equiv.) in THF (0.2 mM), $K_2CO_3$ (10 equiv.) and freshly distilled methyl iodide (5.0 equiv.) were added under an argon atmosphere. The reaction was stirred overnight before additional $K_2CO_3$ (5.0 equiv.), and methyl iodide (2.5 equiv.) were added. The reaction was stirred overnight, quenched with $H_2O$ (10 mL) and THF was removed *in vacuo*. The aqueous layer was extracted three times with EtOAc, and the combined organic layers were washed with brine twice, dried over $MgSO_4$, filtered, and concentrated *in vacuo*. The crude residue was purified by silica gel chromatography to give the desired product.[3]

###### 1.2.1.5 General procedure 5 – Boc deprotection

Procedure adapted from Han *et al*. α-dimethyl-β-keto-ester was added to a flask under argon then anhydrous 4 M HCl in dioxane (6 equiv.) was added and the reaction was stirred at ambient temperature. The solvent was removed *in vacuo* after 2 hours and residual dioxane was co-evaporated with hexane to provide the product.[4]

##### 1.2.1.6 General procedure 6 – HATU coupling

HATU (2 equiv.) was added to a solution of carboxylic acid (1.0 equiv.), in DMF (0.2 mM) under argon on ice and stirred for 2 minutes. Separately, the amine was dissolved in DMF (0.2 mM) and the amine solution and DIPEA (3 equiv.) were simultaneously transferred to the reaction flask dropwise. The resulting solution allowed to warm to ambient temperature overnight before diluting with sat. $NaHCO_3$. The aqueous phase was extracted three times with EtOAc and the combined organic layers were washed twice with 10% (w/v) citric acid, 5% (w/v) LiCl and brine solutions, dried over $MgSO_4$ and concentrated *in vacuo*. The crude residue was purified by silica gel chromatography to yield the desired product.

#### 1.2.2 Chemical synthesis of butyryl amino acids

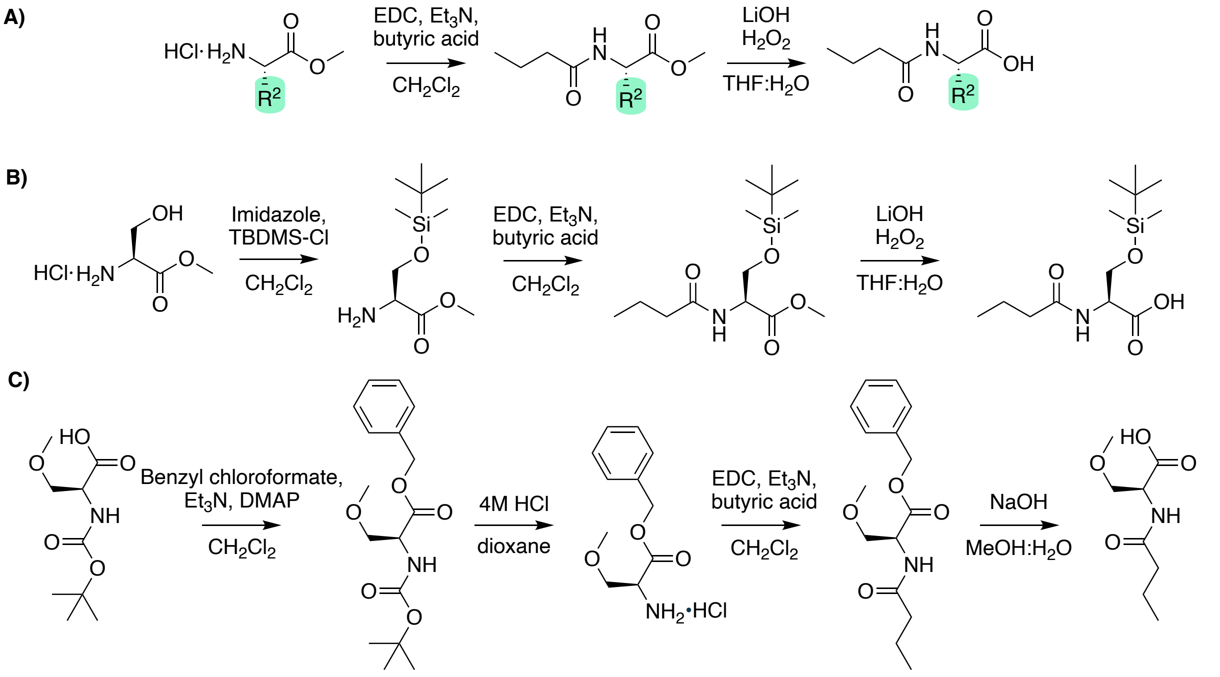

*Scheme S1.1: Synthetic routes described In this section: A) synthesis of butyryl-L-amino acids, B) synthesis of TBDMS-protected butyryl-L-serine, C) synthesis of butyryl-O-methyl-L-serine.*

##### Methyl butyryl-L-serinate

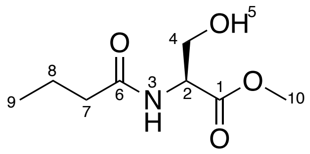

Prepared as per general procedure 1 – with methyl L-serinate hydrochloride (1.7 g, 10.9 mmol, 1 equiv.), purified by silica gel chromatography (EtOAc) yielding the product as a clear oil (1.537 g, 8.1 mmol, 74%). Characterisation data are in agreement with previous reports.

$\delta_H$ (400 MHz, Chloroform-*d*) 0.96 (3H, t, *J* 7.5, H9), 1.68 (2H, sext., *J* 7.5, H8), 2.25 (2H, t, *J* 7.5, H7), 3.79 (3H, s, H10), 3.87 – 3.94 (1H, m, H4), 3.95 – 4.07 (1H, m, H4), 4.68 (1H, dt, *J* 7.5, 4.0), 6.45 (1H, s, H3).
$\delta_C$ (101 MHz, Chloroform-*d*) 13.8 (C9), 19.1 (C8), 38.5 (C7), 52.9 (C10), 54.8 (C2), 63.7 (C4), 171.2 (C1), 173.8 (C6).
HRMS (ESI): Calc. for $[M+Na]^+$ $C_8H_{15}NO_4Na^+$ = 212.0893, Obs. = 212.0896.

**butyryl-L-serine**

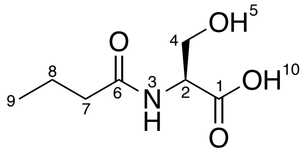

Prepared as per general procedure 2 – with methyl butyryl-L-serinate (1.285 g, 6.7 mmol, 1 equiv.), yielding the final product as a white waxy solid (0.589 g, 3.3 mmol, 50%). Characterisation data are in agreement with previous reports.
$\delta_H$ (400 MHz, Methanol-*d*$_4$) 0.97 (3H, t, *J* 7.5, H9), 1.66 (2H, sext., *J* 7.5, H8), 2.26 (2H, t, *J* 7.5, H7), 3.82 (1H, dd, *J* 11.0, 4.0, H4), 3.90 (1H, dd, *J* 11.0, 5.0, H4), 4.50 (1H, t, *J* 4.5, H2).
$\delta_C$ (101 MHz, Methanol-*d*$_4$) 14.0 C9), 20.2 (C8), 38.7 (C7), 56.0 (C2), 62.9 (C4), 173.5 (C1), 176.2 (C6).
HRMS (ESI): Calc. for $[M+Na]^+$ $C_7H_{13}NO_4Na^+$ = 198.0737, Obs. = 198.0741.

**Methyl butyryl-L-alaninate**

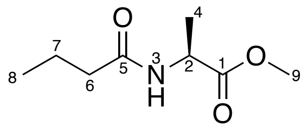

Prepared as per general procedure 1 – with methyl L-alaninate hydrochloride (1.5 g, 10.7 mmol, 1 equiv.), purified by silica gel chromatography (3:2 EtOAc:hexane) yielding the final product as a clear oil (1.715 g, 9.2 mmol, 86%).
$\delta_H$ (400 MHz, Chloroform-*d*) 0.95 (3H, t, *J* 7.5, H8), 1.40 (3H, d, *J* 7.0, H4), 1.67 (2H, sext., *J* 7.5, H7), 2.19 (2H, t, *J* 7.5, H6), 3.75 (3H, s, H9), 4.61 (1H, quint., *J* 7.0, H2), 6.01 (1H, br. s, H3).
$\delta_C$ (101 MHz, Chloroform-*d*) 13.8 (C8), 18.8 (C4), 19.1 (C7), 38.6 (6), 48.0 (2), 52.6 (C9), 172.6 (C5), 173.9 (C1).
HRMS (ESI): Calc. for $[M+Na]^+$ $C_8H_{15}NO_3Na$ $^+$ = 196.0944, Obs. = 196.0955.

**Butyryl-L-alanine**

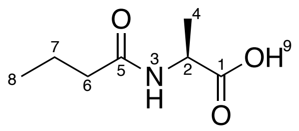

Prepared as per general procedure 2 – with methyl butyryl-L-alaninate (1.575 g, 9.0 mmol, 1 equiv.), yielding the final product as a white waxy solid (1.294 g, 8.1 mmol, 89%). Characterisation data are in agreement with previous reports.[5]

$\delta_H$ (400 MHz, Chloroform-*d*) 0.95 (3H, t, *J* 7.5, H8), 1.46 (3H, d, *J* 7.0, H4), 1.67 (2H, sext., *J* 7.5, H7), 2.23 (2H, t, *J* 7.5, H6), 4.58 (1H, quint., *J* 7.0, H2), 6.26 (1H, d, *J* 7.0, H3).

$\delta_C$ (101 MHz, Chloroform-*d*) 13.7 (C9), 18.2 (C4), 19.1 (C7), 38.4 (C6), 48.4 (C2), 174.1 (C1), 176.0 (C6).

LRMS (ESI): Calc. for $[M-H]^-$ $C_7H_{12}NO_3^-$ = 158.1, Obs. = 158.1.

**Methyl butyryl-L-threoninate**

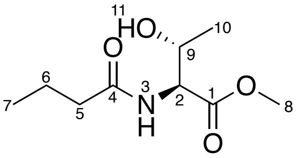

Prepared as per general procedure 1 – with methyl L-threoninate hydrochloride (1.0 g, 5.9 mmol, 1 equiv.), purified by silica gel chromatography (9:1 EtOAc:hexane) yielding the desired product as a white solid (0.715 g, 3.5 mmol, 60%).

$\delta_H$ (400 MHz, Methanol-$d_4$) 0.97 (3H, t, *J* 7.5, H7), 1.17 (3H, d, *J* 6.5, H10), 1.67 (2H, sext., *J* 7.5, H6), 2.29 (2H, t, *J* 7.5, H5), 3.74 (3H, s, H8), 4.11 – 4.38 (1H, m, H9), 4.46 (1H, d, *J* 3.0, H2).

$\delta_C$ (101 MHz, Methanol-$d_4$) 14.0 (C7), 20.3 (C6), 20.3 (C10), 38.7 (C5), 52.7 (C8), 59.2 (C2), 68.3 (C9), 172.6 (C1), 176.6 (C4).

HRMS (ESI): Calc. for $[M+H]^+$ $C_9H_{18}NO_4^+$ = 204.1230, Obs. = 204.1231.

**Butyryl-L-threonine**

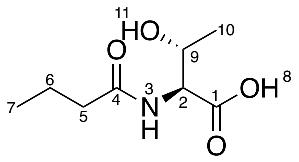

Prepared as per general procedure 2 – with methyl butyryl-L-threoninate (0.7 g, 3.4 mmol, 1 equiv.), yielding the final product as a white solid (0.248 g, 1.3 mmol, 38%). Characterisation data are in agreement with previous reports.[5]

$\delta_H$ (400 MHz, Methanol-$d_4$) 0.98 (3H, t, *J* 7.5, H7), 1.18 (3H, d, *J* 6.5, H10), 1.67 (2H, sext., *J* 7.5, H6), 2.30 (2H, t, *J* 7.5, H5), 4.31 (1H, qd, *J* 6.5, 3.0, H9), 4.43 (1H, d, *J* 3.0, H2).

δ$_C$ (101 MHz, Methanol-$d_4$) 14.0 (C7), 20.3 (C6), 20.5 (C10), 38.8 (C5), 59.0 (C2), 68.4 (C9), 173.8 (C1), 176.5 (C4).

LRMS (ESI): Calc. for $[M-H]^-$ $C_8H_{14}NO_4^-$ = 188.1, Obs. = 188.1.

**methyl *O*-(*tert*-butyldimethylsilyl)-L-serinate**

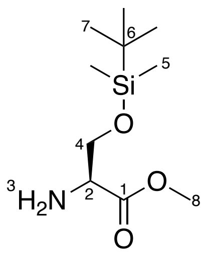

Procedure adapted from Drescher *et al.* Imidazole (2.66 g, 39.1 mmol, 3.0 equiv.) then TBDMS-Cl (2.72 g, 18.0 mmol, 1.4 equiv.) were added to a solution of L-serine methyl ester hydrochloride (2.0 g, 12.8 mmol, 1 equiv.) in $CH_2Cl_2$ (100 mL) and stirred at ambient temperature overnight. The mixture was diluted with sat. $NH_4Cl$ solution (50 mL), the organic layer was separated, and the aqueous layer was extracted with $CH_2Cl_2$ (3 × 50 mL). The combined organic layers were dried over $MgSO_4$ and concentrated *in vacuo*. The crude residue was purified by silica gel chromatography (EtOAc) to give the product as a clear oil (2.76 g, 11.9 mmol, 93%).[6] Characterisation data are in agreement with previous reports.[6]

δ$_H$ (400 MHz, Methanol-$d_4$) 0.06 (3H, s, H5), 0.07 (3H, s, H5), 0.89 (9H, s, H7), 3.51 (1H, t, *J* 4.0, H2), 3.72 (3H, s, H8), 3.79 (1H, dd, *J* 10.0, 4.0, H4), 3.96 (1H, dd, *J* 10.0, 4.0, H4).

δ$_C$ (101 MHz, Methanol-$d_4$) -5.5 (C5), -5.4 (C5), 19.1 (C6), 26.2 (C7), 52.5 (C8), 57.1 (C2), 66.3 (C4), 175.4 (C1).

LRMS (ESI): Calc. for $[M+H]^+$ $C_{10}H_{24}NO_3Si^+$ = 234.2, Obs. = 234.1.

**Methyl *O*-(tert-butyldimethylsilyl)-*N*-butyryl-L-serinate**

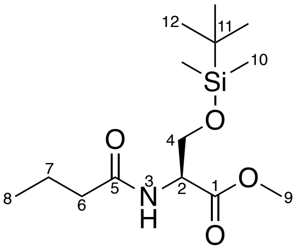

Prepared as per general procedure 1 – with methyl *O*-(*tert*-butyldimethylsilyl)-L-serinate (1.3 g, 5.9 mmol, 1 equiv.), purified by silica gel chromatography (hexane:EtOAc 4:1) yielding the final product as a clear oil (1.534 g, 5.1 mmol, 91%).

δ$_H$ (400 MHz, Chloroform-*d*) 0.01 (3H, s, H10), 0.02 (3H, s, H10), 0.85 (9H, s, H12), 0.96 (3H, t, *J* 7.5, H8), 1.68 (2H, sext., *J* 7.5, H7), 2.23 (2H, t, *J* 7.5, H6), 3.74 (3H, s, H9), 3.81 (1H, dd,

*J* 10.0, 3.0, H4), 4.05 (1H, dd, *J* 10.0, 3.0, H4), 4.68 (1H, dt, *J* 8.0, 3.0, H2), 6.25 (1H, d, *J* 8.0, H3).
$\delta_C$ (101 MHz, Chloroform-*d*) -5.5 (C10), 13.8 (C8), 18.3 (C11), 19.1 (C7), 25.8 (C12), 38.6 (C6), 52.5 (C9), 54.2 (C2), 63.7 (C4), 171.2 (C1), 172.8 (C5).
HRMS (ESI): Calc. for $[M+Na]^+$ $C_{14}H_{29}NO_4SiNa^+$ = 326.1758, Obs. = 326.1767.

**_O_-(tert-butyldimethylsilyl)-_N_-butyryl-L-serine**

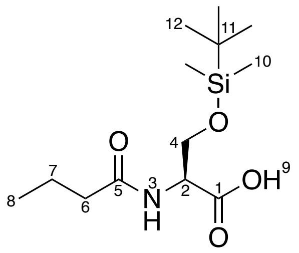

Prepared as per general procedure 2 – with methyl *O*-(tert-butyldimethylsilyl)-*N*-butyryl-L-serinate (0.2 g, 6.6 mmol, 1 equiv.) yielding the product as a white waxy solid (0.165 g, 5.7 mmol, 86%).
$\delta_H$ (400 MHz, Chloroform-*d*) 0.05 (3H, s, H10), 0.05 (3H, s, H10), 0.87 (9H, s, H12), 0.96 (3H, t, *J* 7.5, H8), 1.68 (2H, sext., *J* 7.5, H7), 3.83 (1H, dd, *J* 10.0, 4.0, H4), 4.12 (1H, dd, *J* 10.0, 3.0, H4), 4.64 – 4.73 (1H, m, H2), 6.38 (1H, d, *J* 8.0, H3).
$\delta_C$ (101 MHz, Chloroform-*d*) -5.5 (C10), -5.4 (C10), 13.8 (C8), 18.3 (C11), 19.1 (C7), 25.8 (C12), 38.5 (C6), 54.1 (C2), 63.3 (C4), 173.8 (C5), 174.3 (C1).
HRMS (ESI): Calc. for $[M+Na]^+$ $C_{13}H_{27}NO_4SiNa^+$ = 312.1602, Obs. = 312.1597.

**Methyl butyryl-L-valinate**

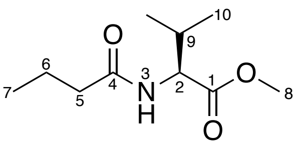

Prepared as per general procedure 1 – with methyl L-valinate hydrochloride (1 g, 6.0 mmol, 1 equiv.), purified by silica gel chromatography (1:1 hexane:EtOAc), yielding the final product as a clear oil (0.767 g, 3.8 mmol, 64%).
$\delta_H$ (400 MHz, Chloroform-*d*) 0.70–1.03 (9H, m, H7 & H10), 1.63 (2H, sext., *J* 7.5, H6), 2.03–2.13 (1H, m, H9), 2.17 (2H, t, *J* 7.5, H5), 3.68 (3H, s, H8), 4.53 (1H, dd, *J* 9.0, 5.0, H2), 6.08 (1H, d, *J* 9.0, H3).
$\delta_C$ (101 MHz, Chloroform-*d*) 13.7 (C7), 17.9 (C10), 19.0 (C10), 19.2 (C6), 31.3 (C9), 38.6 (C5), 52.1 (C8), 56.9 (C2), 172.8 (C1), 173.0 (C4).
HRMS (ESI): Calc. for $[M+Na]^+$ $C_{10}H_{19}NO_3Na^+$ = 224.1257, Obs. = 224.1257.

**Butyryl-L-valine**

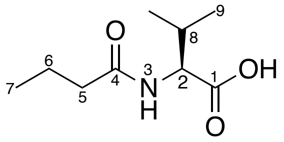

Prepared as per general procedure 2 – with methyl butyryl-L-valinate (0.767 g, 3.8 mmol, 1 equiv.), yielding the final product as a white powder (0.567 g, 3.0 mmol, 79%). Characterisation data are in agreement with previous reports.[5]

$\delta_H$ (400 MHz, Methanol-$d_4$) 0.91 – 1.02 (9H, m, H7 & H9), 1.65 (2H, sext., *J* 7.5, H6), 2.13-2.21 (1H, m, H8), 2.25 (2H, t, *J* 7.5, H5), 4.33 (1H, d, *J* 6.0, H2).

$\delta_C$ (101 MHz, Methanol-$d_4$) 14.0 (C7), 18.4 (C9), 19.6 (C9), 20.4 (C6), 31.6 (C8), 38.6 (C5), 59.0 (C2), 175.0 (C1), 176.4 (C4).

LRMS (ESI): Calc. for $[M-H]^-$ $C_9H_{16}NO_3^-$ = 186.1, Obs. = 186.1.

**Methyl butyryl-L-phenylalaninate**

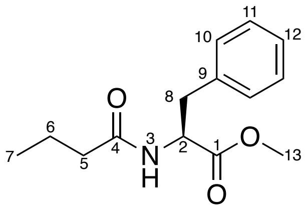

Prepared as per general procedure 1 – with methyl L-phenylalaninate (1.0 g, 4.6 mmol, 1 equiv.), purified by silica gel chromatography to yield the final product as a clear oil (1.0 g, 4.0 mmol, 87%). Characterisation data are in agreement with previous reports.[7]

$\delta_H$ (400 MHz, Chloroform-*d*) 0.95 (3H, t, *J* 7.5, H7), 1.66 (2H, sext., *J* 7.5, H6), 2.19 (2H, t, *J* 7.5, H5), 3.12 (1H, dd, *J* 14.0, 6.0, H8), 3.20 (1H, dd, *J* 14.0, 6.0, H8), 3.76 (3H, s, H13), 4.95 (1H, dt, *J* 8.0, 6.0, H2), 5.99 (1H, d, *J* 8.0, H3).

$\delta_C$ (101 MHz, Chloroform-*d*) 13.8 (C7), 19.1 (C6), 38.0 (C8), 38.5 (C5), 52.4 (C13), 53.0 (C2), 127.2 (C10), 128.6 (C11), 129.3 (C12), 136.0 (C9), 172.3 (4), 172.6 (C1).

LRMS (ESI): Calc. for $[M+Na]^+$ $C_{14}H_{16}NO_3Na^+$ = 272.1, Obs. = 272.1.

**Butyryl-L-phenylalanine**

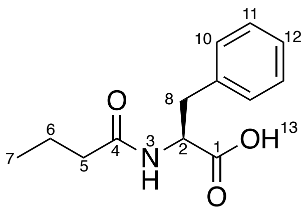

Prepared as per general procedure 2 – with methyl butyryl-L-phenylalaninate (0.8 g, 3.2 mmol, 1 equiv.), yielding the final product as a white waxy solid (0.634 g, 2.70 mmol, 84%). Characterisation data are in agreement with previous reports.[7]

$\delta_H$ (400 MHz, Methanol-$d_4$) 0.82 (3H, t, *J* 7.5, H7), 1.52 (2H, sext., *J* 7.5, H6), 2.12 (2H, t, *J* 7.5, H5), 2.93 (1H, dd, *J* 14.0, 9.5, H8), 3.22 (1H, dd, *J* 14.0, 5.0, H8), 4.68 (1H, dd, *J* 9.6, 5.0, H2), 7.16 – 7.30 (5H, m, H10, H11 & H12).

$\delta_C$ (101 MHz, Methanol-$d_4$) 13.8 (C7), 20.2 (C6), 38.4 (C8), 38.6 (C5), 54.9 (C2), 127.7 (C10), 129.4 (C11), 130.2 (C12), 138.6 (C9), 174.8 (C1), 176.0 (C4).

LRMS (ESI): Calc. for $[M-H]^-$ $C_{13}H_{16}NO_3^-$ = 234.1, Obs. = 234.1.

**Methyl butyrylglycinate**

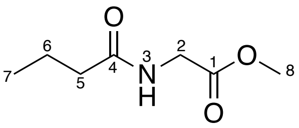

Prepared as per general procedure 1 – with methyl glycinate hydrochloride (0.5 g, 3.9 mmol, 1 equiv.), purified by silica gel chromatography (7:3 EtOAc:hexane) yielding the final product as a clear oil (0.429 g, 2.7 mmol, 68%).

$\delta_H$ (400 MHz, Chloroform-*d*) 0.90 (3H, t, *J* 7.5, H7), 1.62 (2H, sext., *J* 7.5, H6), 2.18 (2H, t, *J* 7.5, H5), 3.70 (3H, s, H8), 3.98 (2H, d, *J* 5.0, H2), 6.31 (1H, br. s, H3).

$\delta_C$ (101 MHz, Chloroform-*d*) 13.7 (C7), 19.0 (C6), 38.2 (C5), 41.1 (C2), 52.3 (C8), 170.6 (C1), 173.4 (C4).

HRMS (ESI): Calc for $[M+Na]^+$ $C_7H_{13}NO_3Na^+$ = 182.0788, Obs. = 182.0789.

**Butyrylglycine**

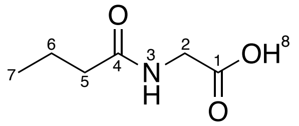

Prepared as per general procedure 2 – with methyl butyrylglycinate (0.16 g, 1.0 mmol, 1 equiv.), yielding the final product as a clear oil (0.079 g, 0.5 mmol, 55%). Characterisation data are in agreement with previous reports.[8]

$\delta_H$ (400 MHz, Methanol-$d_4$) 0.96 (3H, t, *J* 7.5, H7), 1.65 (2H, sext., *J* 7.5, H6), 2.23 (2H, t, *J* 7.5, H5), 3.89 (2H, s, H2).

$\delta_C$ (101 MHz, Methanol-$d_4$) 13.9 (C7), 20.2 (C6), 38.7 (C5), 41.7 (C2), 173.1 (C1), 176.6 (C4).

LRMS (ESI): Calc. for $[M-H]^-$ $C_6H_{10}NO_3^-$ = 144.1, Obs. = 144.1.

**Benzyl *N*-(*tert*-butoxycarbonyl)-*O*-methyl-L-serinate**

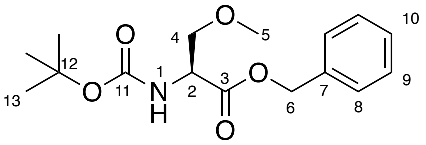

Procedure adapted from Zhou *et al*. Under an atmosphere of argon, $Et_3N$ (0.45 g, 4.4 mmol, 1.3 equiv.) and DMAP (0.041 g, 0.44 mmol, 0.1 equiv.) were added to a solution of *N*-(*tert*-butoxycarbonyl)-*O*-methyl-L-serine (0.75 g 3.4 mmol, 1 equiv.) in $CH_2Cl_2$ (50 mL). The resulting solution was cooled to 0 °C, and benzyl chloroformate (0.76 g, 4.4 mmol, 1.3 equiv.) was added dropwise. The reaction was allowed to warm to ambient temperature and then diluted with brine (25 mL) after 3 hours. The layers were separated, and the aqueous layer was extracted with $CH_2Cl_2$ (3 × 25 mL). The organic layers were combined and dried over $MgSO_4$, filtered and organic solvent was removed *in vacuo*. The crude residue was purified by silica gel chromatography (7:3 hexane:EtOAc) to yield the product as a white solid (0.655 g, 2.1 mmol, 62%).[9]

$\delta_H$ (400 MHz, Methanol-*d*$_4$) 1.43 (9H, s, H13), 3.27 (3H, s, H5), 3.59 (1H, dd, *J* 10.0, 4.0, H4), 3.73 (1H, dd, *J* 10.0, 5.0, H4), 4.39 (1H, t, *J* 4.5, H2), 5.10 (1H, d, *J* 12.5, H6), 5.22 (1H, d, *J* 12.5, H6), 7.17 – 7.60 (5H, m, H8, H9 & H10).

$\delta_C$ (101 MHz, Methanol-*d*$_4$) 28.7 (C13), 55.4 (C2), 59.3 (C5), 67.8 (C6), 73.0 (C4), 80.6 (C12), 129.0 (C8), 129.1 (C9), 129.4 (C10), 137.1 (C7), 157.6 (C11), 171.9 (C3).

HRMS (ESI): Calc. for $[M+Na]^+$ $C_{16}H_{23}NO_5Na^+$ = 332.1468, Obs. = 332.1468.

**Benzyl *O*-methyl-L-serinate hydrochloride**

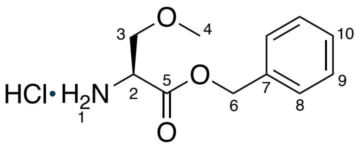

Prepared as per general procedure 5 – with benzyl *N*-(*tert*-butoxycarbonyl)-*O*-methyl-L-serinate (0.55 g, 1.8 mmol, 1 equiv.) yielding the desired product as a white powder (0.429 g, 1.7 mmol, 98%).

$\delta_H$ (400 MHz, Methanol-*d*$_4$) 3.36 (3H, s, H4), 3.80 (1H, dd, *J* 10.5, 3.0, H3), 3.88 (1H, dd, *J* 10.5, 4.5, H3), 4.33 (1H, t, *J* 4.0, H2), 5.26 (1H, d, *J* 12.0, H6), 5.35 (1H, d, *J* 12.0, H6), 7.31 – 7.46 (5H, m, H7, H8 and H9).

$\delta_C$ (101 MHz, Methanol-*d*$_4$) 54.5 (C2), 59.6 (C4), 69.1 (C6), 70.5 (C3), 129.5 (C8), 129.7 (C9), 129.7 (C10), 136.4 (C7), 168.5 (C5).

HRMS (ESI): Calc. for $[M+Na]^+$ $C_{11}H_{16}NO_3Na^+$ = 210.1125, Obs. = 210.1129.

**Benzyl *N*-butyryl-*O*-methyl-L-serinate**

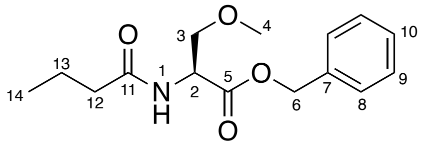

Procedure adapted from Li *et al*. To a solution of butyric acid (0.14 mL, 1.6 mmol, 1 equiv.) in $CH_2Cl_2$ (10 mL), EDC hydrochloride (0.388 g, 2.0 mmol, 1.3 equiv.) and DMAP (0.038 g, 0.3 mmol, 0.2 equiv.) were added under argon at 0 °C and stirred for 10 minutes. Benzyl *O*-methyl-L-serinate (0.42 g, 1.7 mmol, 1.1 equiv.) and $Et_3N$ (0.33 mL, 2.3 mmol, 1.5 equiv.), were added successively and the reaction was allowed to warm to ambient temperature overnight. The reaction was quenched with 1 M HCl (15 mL), and the organic layer was separated. The aqueous layer was extracted with EtOAc (3 x 15 mL) then the combined organic layers were washed with 5% (w/v) $NaHCO_3$ (45 mL) and brine (45 mL). The organic solvent was removed *in vacuo* and the crude residue purified by recrystallisation in hexane, yielding a white powder (0.25 g, 0.9 mmol, 52%).[10]

$\delta_H$ (400 MHz, Methanol-*d*$_4$) 0.91 (3H, t, *J* 7.5, H14), 1.60 (2H, sext., *J* 7.5, H13), 2.21 (2H, t, *J* 7.5, H12), 3.28 (3H, s, H4), 3.59 (1H, dd, *J* 10.0, 4.0, H3), 3.75 (1H, dd, *J* 10.0, 5.0, H3), 4.65 (1H, t, *J* 4.5, H2), 5.11 (1H, d, *J* 12.5, H6), 5.20 (1H, d, *J* 12.5, H6), 7.25 – 7.35 (5H, m, H8, H9 & H10).

$\delta_C$ (101 MHz, Methanol-*d*$_4$) 13.9 (C14), 20.3 (C13), 38.5 (C12), 54.2 (C2), 59.3 (C4), 68.0 (C6), 72.8 (C3), 129.1 (C10), 129.3 (C9), 129.5 (C8), 137.2 (C7), 171.4 (C5), 176.3 (C11).

HRMS (ESI): Calc. for $[M+H]^+$ $C_{15}H_{22}NO_4^+$ = 280.1543, Obs. = 280.1544.

***N*-butyryl-*O*-methyl-L-serine**

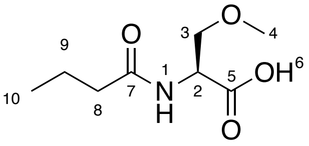

Procedure adapted from Ogawa *et al*. Benzyl *N*-butyryl-*O*-methyl-L-serinate (0.1 g, 0.36 mmol, 1 equiv.) was dissolved in MeOH (1.1 mL) and 1 M NaOH (1.1 mL, 3 equiv.) was added dropwise and the reaction stirred for 15 minutes. Solvent was removed *in vacuo* and the remaining solution washed with diethyl ether (2 x 5 mL). The aqueous layer was acidified to pH 1 with 1 M HCl, saturated with NaCl and extracted with EtOAc (3 x 5 mL). The organics were dried over $MgSO_4$, filtered and solvent removed *in vacuo* yielding the product as a clear oil.[11] (0.032 g, 0.169 mmol, 47%).

$\delta_H$ (400 MHz, Methanol-*d*$_4$) 0.96 (3H, t, *J* 7.5, H10), 1.65 (2H, sext., *J* 7.5, H9), 2.25 (2H, t, *J* 7.5, H8), 3.35 (3H, s, H4), 3.63 (1H, dd, *J* 10.0, 4.0, H3), 3.78 (1H, dd, *J* 10.0, 5.0, H3), 4.60 (1H, dd, *J* 5.0, 3.5, H2).

$\delta_C$ (101 MHz, Methanol-$d_4$) 13.9 (C10), 20.2 (C9), 38.5 (C8), 53.9 (C2), 59.3 (C4), 73.0 (C3), 173.2 (C7), 176.2 (C5).

LRMS (ESI): Calc. for $[M-H]^-$ $C_8H_{14}NO_4^-$ = 188.1, Obs. = 188.1.

##### 1.2.3 Chemical synthesis of amino α-dimethyl-β-keto-ester fragments

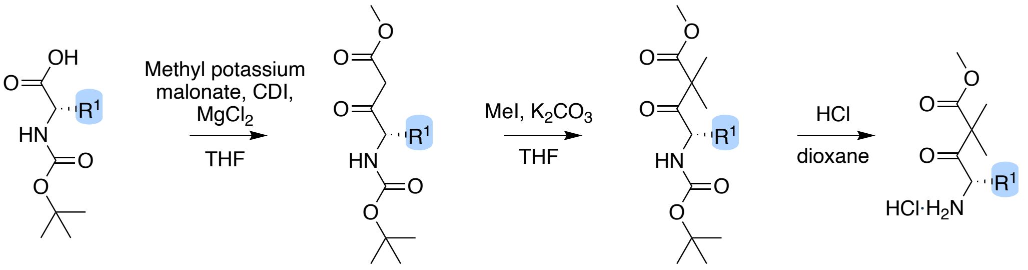

*Scheme S1.2: Synthetic route to amino α-dimethyl-β-keto-ester fragments.*

**Methyl (*S*)-4-((*tert*-butoxycarbonyl)amino)-6-methyl-3-oxoheptanoate**

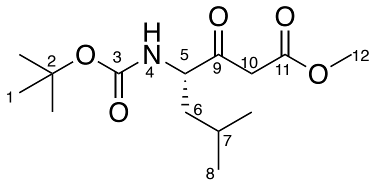

Prepared as per general procedure 3 - with (*tert*-butoxycarbonyl)-L-leucine (3.0 g, 12.9 mmol, 1 equiv.), purified by silica gel chromatography (4:1 Hexane:EtOAc) yielding the final product as a pale-yellow oil (2.578 g, 8.9 mmol, 69.1%).

$\delta_H$ (400 MHz, Chloroform-$d$) 0.92 (6H, d, $J$ 6.5, H8), 1.32 – 1.39 (1H, m, H6), 1.42 (9H, s, H1), 1.54 – 1.64 (1H, m, H6), 1.64 – 1.76 (1H, m, H7), 3.42 – 3.66 (2H, m, H10), 3.71 (3H, s, H12), 4.26 – 4.39 (1H, m, H5), 4.95 (1H, d, $J$ 8.5, H4).

$\delta_C$ (101 MHz, Chloroform-$d$) 21.7 (C8), 23.3 (C8), 24.9 (C7), 28.4 (C1), 39.9 (C6), 46.1 (C10), 52.5 (C12), 58.3 (C5), 80.2 (C2), 155.7 (C3), 167.6 (C11), 203.0 (C9).

HRMS (ESI): Calc. for $[M+Na]^+$ $C_{14}H_{25}NO_5Na^+$ = 310.1625, Obs. = 310.1620.

**methyl (*S*)-4-((*tert*-butoxycarbonyl)amino)-2,2,6-trimethyl-3-oxoheptanoate (15)**

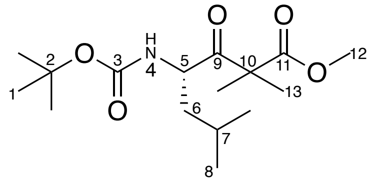

Prepared as per general procedure 4 – with methyl (*S*)-4-((*tert*-butoxycarbonyl)amino)-6-methyl-3-oxoheptanoate (1.6 g, 5.6 mmol, 1 equiv.), purified by silica gel chromatography (7:3 hexane:$Et_2O$) yielding the final product as a clear oil (1.435 g, 4.5 mmol, 82%).

$\delta_H$ (400 MHz, Chloroform-*d*) 0.89 (3H, d, *J* 6.5, H8), 0.92 (3H, d, *J* 6.5, H8), 1.16 – 1.78 (17H, m, H1, H6 & H13), 1.58 – 1.74 (1H, m, H7), 3.69 (3H, s, H12), 4.61 (1H, td, *J* 10.0, 3.5, H5), 4.79 (1H, d, *J* 10.0, H4).

$\delta_C$ (101 MHz, Chloroform-*d*) 21.5 (C8), 22.1 (C13), 22.4 (C13), 23.6 (C8), 24.8 (C7), 28.4 (C1), 42.0 (C6), 52.6 (C12), 53.9 (C5), 79.8 (C2), 155.1 (C3), 173.6 (C11), 208.8 (C9).

HRMS (ESI): Calc. for $[M+Na]^+$ $C_{16}H_{29}NO_5Na^+$ = 338.1938, Obs. = 338.1937.

**methyl (*S*)-4-amino-2,2,6-trimethyl-3-oxoheptanoate hydrochloride**

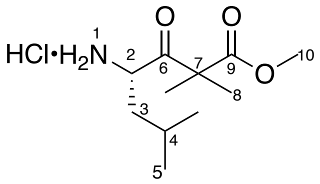

Prepared as per general procedure 5 – with Methyl (*S*)-4-((tert-butoxycarbonyl)amino)-2,2,6-trimethyl-3-oxoheptanoate (1.780 g, 5.6 mmol, 1 equiv.) yielding the final product as a white powder (1.420 g, 5.6 mmol, 100%). Characterisation data are in agreement with previous reports.

$\delta_H$ (400 MHz, Chloroform-*d*) 0.95 (3H, d, *J* 6.5, H5), 0.98 (3H, d, *J* 6.5, H5), 1.33 – 1.46 (4H, m, H3 & H8), 1.49 (3H, s, H8), 1.73 – 1.92 (1H, m, H3), 2.09 – 2.15 (1H, m, H4), 3.72 (3H, s, H10), 4.50 (1H, d, *J* 11.0, H2), 8.73 (1H, s, H1).

$\delta_C$ (101 MHz, Chloroform-*d*) 21.0 (C5), 22.8 (C8), 23.0 (C8), 23.7 (C5), 24.5 (C4), 39.5 (C3), 53.0 (C10), 54.4 (C7), 54.8 (C2), 173.1 (C9), 203.8 (C6).

HRMS (ESI): Calc. for $[M+H]^+$ $C_{11}H_{22}NO_3^+$ = 216.1594, Obs. = 216.1592.

**methyl (*S*)-4-((*tert*-butoxycarbonyl)amino)-3-oxo-5-phenylpentanoate**

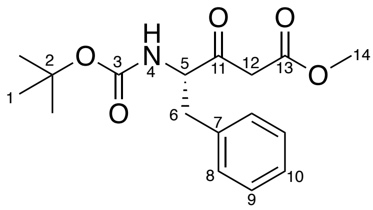

Prepared as per general procedure 3 – with (*tert*-butoxycarbonyl)-L-phenylalanine (0.493 g, 1.9 mmol, 1 equiv.), purified by silica gel chromatography (3:2 hexane:EtOAc) yielding the final product as a white powder (0.363 g, 1.1 mmol, 61%).

$\delta_H$ (400 MHz, Chloroform-*d*) 1.43 (9H, s, H1), 3.00 (1H, dd, *J* 14.0, 7.5, H6), 3.18 (1H, dd, *J* 14.0, 6.0, H6), 3.42 – 3.62 (2H, m, H12), 3.74 (3H, s, H14), 4.59 (1H, q, *J* 7.5, H5), 5.05 (1H, d, *J* 8.0, H4), 7.13 – 7.26 (2H, m, H8), 7.24 – 7.49 (3H, m, H9 & H10).

$\delta_C$ (100 MHz, Chloroform-*d*) 28.4 (C1), 37.1 (C6), 46.8 (C12), 52.5 (C14), 60.6 (C5), 80.4 (C2), 127.2 (C8), 128.9 (C10), 129.4 (C9), 136.2 (C7), 155.4 (C3), 167.4 (C13), 202.0 (C11).

HRMS (ESI): Calc. for $[M+Na]^+$ $C_{17}H_{23}NO_5Na^+$ = 344.1468, Obs. = 344.1468.

**methyl (*S*)-4-((*tert*-butoxycarbonyl)amino)-2,2-dimethyl-3-oxo-5-phenylpentanoate**

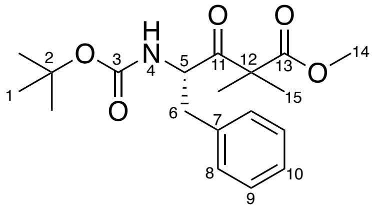

Prepared as per general procedure 4 – with Methyl (*S*)-4-((tert-butoxycarbonyl)amino)-3-oxo-5-phenylpentanoate (0.8 g, 2.5 mmol, 1 equiv), purified by silica gel chromatography yielding the final product as a white powder (0.748 g, 2.1 mmol, 86%).

$\delta_H$ (400 MHz, Chloroform-*d*) 1.28 – 1.49 (15H, m, H1 & H15), 2.85 (1H, dd, *J* 14.0, 8.0, H6), 3.16 (1H, dd, *J* 14.0, 5.5, H6), 3.73 (3H, s, H14), 4.82 (1H, d, *J* 10.0, H4), 4.92 (1H, q, *J* 8.0, H5), 7.22 (2H, d, *J* 7.5, H8), 7.24 – 7.38 (3H, m, H9 & H10).

$\delta_C$ (101 MHz, Chloroform-*d*) 21.8 (C15), 22.2 (C15), 28.3 (C1), 38.6 (C6), 52.6 (C14), 54.9 (C12) 56.5 (C5), 80.0 (C2), 126.9 (C8), 128.6 (C10), 129.6 (C9), 136.6 (C7), 154.7 (C3), 173.6 (C13), 207.6 (C11).

HRMS (ESI): Calc. for $[M+Na]^+$ $C_{19}H_{27}NO_5Na^+$ = 372.1781, Obs. = 372.1785.

**Methyl (S)-4-amino-2,2-dimethyl-3-oxo-5-phenylpentanoate hydrochloride**

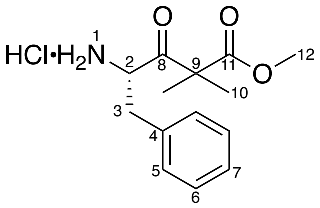

Prepared as per general procedure 5 – with methyl (*S*)-4-((tert-butoxycarbonyl)amino)-2,2-dimethyl-3-oxo-5-phenylpentanoate (0.7 g, 2.0 mmol, 1 equiv.), yielding the final product as a white powder (0.56 g, 1.94 mmol, 97%).

$\delta_H$ (400 MHz, Methanol-*d*$_4$) 1.38 (3H, s, H10), 1.49 (3H, s, H10), 2.87 (1H, dd, *J* 14.5, 9.0, H3), 3.28 – 3.38 (1H, m, H3), 3.79 (3H, s, H12), 4.78 (1H, dd, *J* 9.0, 5.0, H2), 7.30 (2H, d, *J* 7.0, H5), 7.33 – 7.45 (3H, m, H6 & H7).

$\delta_C$ (101 MHz, Methanol-*d*$_4$) 22.5 (C10), 22.6 (C10), 37.7 (C3), 53.6 (C12), 56.2 (C9), 57.7 (C2), 129.1 (C7), 130.3 (C5), 130.6 (C6), 135.1 (C4), 174.7 (C11), 205.8 (C8).

HRMS (ESI): Calc. for $[M+H]^+$ $C_{14}H_{20}NO_3^+$ = 250.1438, Obs. = 250.1446.

**Methyl (*S*)-4-((tert-butoxycarbonyl)amino)-6-methyl-3-oxohept-6-enoate**

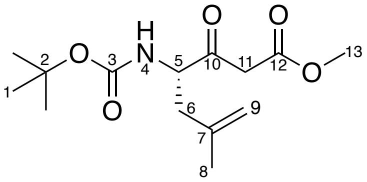

Prepared as per general procedure 3 – with (*tert*-butoxycarbonyl)-4,5-L-dehydro-leucine dicyclohexylammonium salt (0.50 g, 1.2 mmol, 1 equiv.), purified by silica gel chromatography (1:1 hexane:EtOAc) yielding the final product as a white powder (0.233 g, 0.8 mmol, 67%).

$\delta_H$ (400 MHz, Chloroform-*d*) 1.43 (9H, s, H1), 1.75 (3H, s, H8), 2.27 (1H, dd, *J* 14.0, 9.5, H6), 2.56 (1H, dd, *J* 14.5, 5.0, H6), 3.45 – 3.69 (2H, m, H11), 3.74 (3H, s, H13), 4.38 (1H, q, *J* 8.0, H5), 4.78 (1H, s, H9), 4.89 (1H, s, H9).

$\delta_C$ (101 MHz, Chloroform-*d*) 22.0 (C8), 28.4 (C1), 39.2 (C6), 46.3 (C11), 52.5 (C13), 57.8 (C5), 80.5 (C2), 114.9 (C9), 140.6 (C7), 155.6 (C3), 167.6 (C12), 202.5 (C10).

HRMS (ESI): Calc. for $[M+Na]^+$ $C_{14}H_{23}NO_5Na^+$ = 308.1468, Obs. = 308.1466.

**Methyl (*S*)-4-((tert-butoxycarbonyl)amino)-2,2,6-trimethyl-3-oxohept-6-enoate**

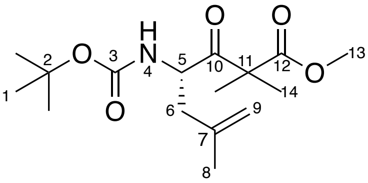

Prepared as per general procedure 4 – with Methyl (*S*)-4-((tert-butoxycarbonyl)amino)-6-methyl-3-oxohept-6-enoate (0.547 g, 1.9 mmol, 1 equiv.), purified by silica gel chromatography (4:1 hexane:EtOAc) yielding the final product as a white powder (0.328 g, 1.0 mmol, 55%).

$\delta_H$ (400 MHz, Chloroform-*d*) 1.26 – 1.55 (15H, m, H1 & H14), 1.73 (3H, s, H8), 2.11 (1H, dd, *J* 14.5, 9.5, H6), 2.44 (1H, d, *J* 14.0, H6), 3.72 (3H, s, H13), 4.68 – 4.70 (1H, m, H5), 4.73 (1H, s, H9), 4.82 (1H, s, H9).

$\delta_C$ (101 MHz, Chloroform-*d*) 22.0 (C8), 22.1 (C14), 22.5 (C14), 28.3 (C1), 41.0 (C6), 52.7 (C13), 53.9 (C5), 54.8 (C11), 79.9 (C2), 114.5 (C9), 140.9 (C7), 155.1 (C3), 173.8 (C12), 208.1 (C10).

HRMS (ESI): Calc. for $[M+Na]^+$ $C_{16}H_{27}NO_5Na^+$ = 336.1781, Obs. = 336.1777.

**Methyl (*S*)-4-amino-2,2,6-trimethyl-3-oxohept-6-enoate hydrochloride**

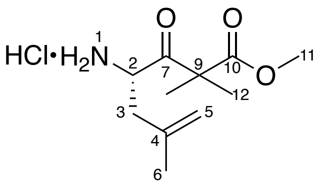

Prepared as per general procedure 5 – with Methyl (*S*)-4-((tert-butoxycarbonyl)amino)-2,2,6-trimethyl-3-oxohept-6-enoate (0.297 g, 0.9 mmol, 1 equiv.), yielding the final product as a white powder (0.232 g, 0.9 mmol, 97%).

$\delta_H$ (400 MHz, Chloroform-*d*) 1.46 (3H, s, H12), 1.55 (3H, s, H12), 1.79 (3H, s, H6), 2.47 – 2.69 (2H, m, H3), 3.75 (3H, s, H11), 4.64 (1H, s, H2), 5.04 (1H, s, H5), 5.11 (1H, s, H5).

$\delta_C$ (101 MHz, Chloroform-*d*) 22.1 (C6), 22.8 (C12), 22.9 (C12), 38.7 (C3), 53.1 (C11), 53.9 (C2), 54.9 (C9), 118.4 (C5), 137.2 (C4), 173.1 (C10), 203.6 (C7).

HRMS (ESI): Calc. for $[M+H]^+$ $C_{11}H_{20}NO_3^+$ = 214.1438, Obs. = 214.1438.

**Methyl (*S*)-4-((tert-butoxycarbonyl)amino)-5-methyl-3-oxohexanoate**

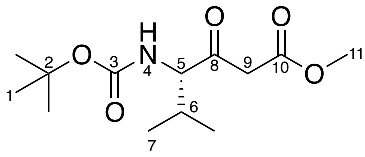

Prepared as per general procedure 3 – with (*tert*-butoxycarbonyl)-L-valine (1.0 g, 4.6 mmol, 1 equiv.), purified by silica gel chromatography (7:3 hexane:EtOAc) yielding the final product as a clear oil (0.91 g, 3.3 mmol, 72%). Characterisation data are in agreement with previous reports.[12]

$\delta_H$ (400 MHz, Chloroform-*d*) 0.80 (3H, d, *J* 7.0, H7), 0.99 (3H, d, *J* 7.0, H7), 1.42 (9H, s, H1), 2.15-2.27 (1H, m, H6), 3.48-4.58 (2H, m, H9), 3.71 (3H, s, H11), 4.29 (1H, dd, *J* 9.0, 4.5, H5), 5.07 (1H, d, *J* 9.0, H4).

$\delta_C$ (101 MHz, Chloroform-*d*) 16.7 (C7), 19.8 (C7), 28.3 (C1), 29.5 (C6), 46.9 (C9), 52.4 (C11), 64.4 (C5), 80.0 (C2), 155.8 (C3), 167.2 (C10), 202.1 (C8).

LRMS (ESI): Calc. for $[M+H]^+$ $C_{13}H_{24}NO_5^+$ = 274.2, Obs. = 274.2.

**Methyl (*S*)-4-((tert-butoxycarbonyl)amino)-2,2,5-trimethyl-3-oxohexanoate**

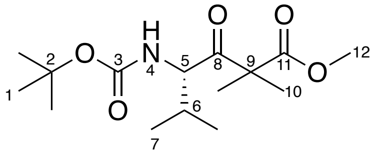

Prepared as per general procedure 4 – with Methyl (*S*)-4-((tert-butoxycarbonyl)amino)-5-methyl-3-oxohexanoate (0.35 g, 1.3 mmol, 1 equiv.), purified by silica gel chromatography yielding the final product as a clear oil (0.312 g, 1.0 mmol, 81%).

$\delta_H$ (400 MHz, Chloroform-*d*) 0.77 (3H, d, *J* 7.0, H7), 0.91 (3H, d, *J* 7.0, H7), 1.38 (3H, s, H10), 1.40 (3H, s, H10), 1.42 (9H, s, H1), 2.05-2.16 (1H, m, H6), 3.70 (3H, s, H12), 4.51 (1H, dd, *J* 10.0, 4.5, H5), 4.83 (1H, d, *J* 10.0, H4).

$\delta_C$ (101 MHz, Chloroform-*d*) 16.3 (C7), 20.0 (C7), 22.2 (C10), 22.6 (C10), 28.4 (C1), 29.9 (C6), 52.6 (C12), 60.3 (C5), 79.9 (C2), 155.5 (C3), 173.6 (C11), 207.7 (C8).

HRMS (ESI): Calc for $[M+Na]^+$ $C_{15}H_{27}NO_5Na^+$ = 324.1781, Obs. = 324.1777.

**Methyl (*S*)-4-amino-2,2,5-trimethyl-3-oxohexanoate hydrochloride**

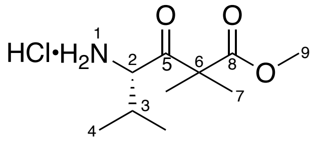

Prepared as per general procedure 5 – with Methyl (*S*)-4-((tert-butoxycarbonyl)amino)-2,2,5-trimethyl-3-oxohexanoate (0.5 g, 1.6 mmol, 1 equiv.), yielding the final product as a white powder (0.383 g, 1.6 mmol, 97%).

$\delta_H$ (400 MHz, Chloroform-*d*) 0.97 (3H, d, *J* 7.0, H4), 1.27 (3H, d, *J* 7.0, H4), 1.45 (3H, s, H7), 1.52 (3H, s, H7), 2.21 (1H, quint., *J* 7.0, H3), 3.73 (3H, s, H9), 4.45 (1H, s, H2), 8.67 (2H, br. s, H1).

$\delta_C$ (101 MHz, Chloroform-*d*) 15.5 (C4), 20.6 (C4), 23.0 (C7), 23.1 (C7), 28.6 (C3), 53.0 (C9), 54.8 (C6), 60.7 (C2), 173.0 (C8), 202.9 (C5).

HRMS (ESI): Calc. for $[M+H]^+$ $C_{10}H_{20}NO_3^+$ = 202.1438, Obs. = 202.1435.

**Methyl (*S*)-4-((tert-butoxycarbonyl)amino)-3-oxopentanoate**

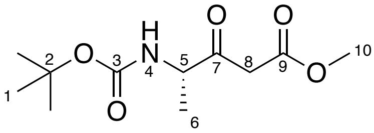

Prepared as per general procedure 3 – with (*tert*-butoxycarbonyl)-L-alanine (1 g, 5.3 mmol, 1 equiv.), purified by silica gel chromatography (hexane:EtOAc 2:1) yielding the final product as a white powder (0.488 g, 1.9 mmol, 38%).

$\delta_H$ (400 MHz, Chloroform-*d*) 1.34 (3H, d, *J* 7.0, H6), 1.43 (9H, s, H1), 3.44 – 3.66 (2H, m, H8), 3.73 (3H, s, H10), 4.36 (1H, quint., *J* 7.0, H5), 4.92 – 5.31 (1H, m, H4).

$\delta_C$ (101 MHz, Chloroform-*d*) 17.1 (C6), 28.4 (C1), 45.7 (C8), 52.6 (C10), 55.5 (C5), 80.3 (C2), 155.2 (C3), 167.5 (C9), 202.4 (C7).

HRMS (ESI): Calc. for $[M+Na]^+$ $C_{11}H_{19}NO_5Na^+$ = 268.1155, Obs. = 268.1157.

**Methyl (*S*)-4-((tert-butoxycarbonyl)amino)-2,2-dimethyl-3-oxopentanoate**

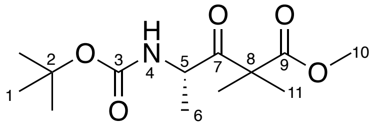

Prepared as per general procedure 4 – with Methyl (*S*)-4-((tert-butoxycarbonyl)amino)-3-oxopentanoate (0.4 g, 1.6 mmol, 1 equiv.), purified by silica gel chromatography yielding the final product as a white powder (0.349 g, 1.2 mmol, 78%).

$\delta_H$ (400 MHz, Chloroform-*d*) 1.24 (1H, d, *J* 7.0, H6), 1.40 (15H, m, H1 & H11), 3.71 (3H, s, H10), 4.63 (1H, quint., *J* 7.0, H5), 4.98 (1H, d, *J* 9.0, H4).
$\delta_C$ (101 MHz, Chloroform-*d*) 19.1 (C6), 22.2 (C11), 22.4 (C11), 28.4 (C1), 51.5 (C5), 52.7 (C5), 79.9 (C2), 154.8 (C3), 173.6 (C9), 208.5 (C7).
HRMS (ESI): Calc. for $[M+Na]^+$ $C_{13}H_{23}NO_5Na^+$ = 296.1468, Obs. = 296.1465.

**Methyl (*S*)-4-amino-2,2-dimethyl-3-oxopentanoate hydrochloride**

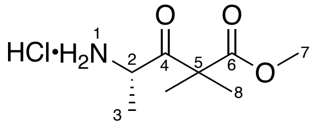

Prepared as per general procedure 5 – with Methyl (*S*)-4-((tert-butoxycarbonyl)amino)-2,2-dimethyl-3-oxopentanoate (0.3 g, 1.1 mmol, 1 equiv.), yielding the final product as a white powder (0.224 g, 1.1 mmol, 97%). Characterisation data are in agreement with previous reports.
$\delta_H$ (400 MHz, Chloroform-*d*) 1.44 (3H, s, H8), 1.52 (3H, s, H8), 1.62 (3H, d, *J* 7.0, H3), 3.74 (3H, s, H7), 4.68 (1H, q, *J* 7.0, H2), 8.60 (2H, br. s, H1).
$\delta_C$ (101 MHz, Chloroform-*d*) 16.7 (C3), 22.4 (C8), 23.0 (C8), 52.7 (C2), 53.1 (C7), 54.2 (C5), 173.0 (C6), 204.7 (C4).
HRMS (ESI): Calc. for $[M+Na]^+$ $C_8H_{15}NO_3Na^+$ = 196.0944, Obs. = 196.0976.

**Methyl (S)-4-((tert-butoxycarbonyl)amino)-5-(1 H-indol-3-yl)-3-oxopentanoate**

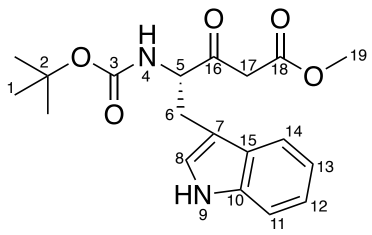

Prepared as per general procedure 3 with *N*-Boc-L-tryptophan (0.605 g, 1.99 mmol, 1 equiv.), yielding the desired product as an orange oil (0.657 g, 1.83 mmol, 91%).
$\delta_H$ (400 MHz, Chloroform-*d*) 1.41 (9H, s, H1), 3.47 (2H, m, H17), 3.25 (2H, m, H6), 3.66 (3H, s, H19), 4.67 (1H, m, H5), 5.13 (1H, br. d, *J* 7.0, H4), 7.03 (1H, s, H8), 7.14 (3H, t, *J* 9.5, H11), 7.12 (1H, t, *J* 7.5, H14), 7.36 (1H, d, *J* 8.0, H13), 7.62 (1H, d, *J* 7.5, H12), 8.15 (1H, br. s, H9).
$\delta_C$ (101 MHz, Chloroform-*d*) 27.0 (C6), 28.4 (C1), 46.9 (C17), 52.5 (C19), 60.0 (C5), 79.9 (C2), 109.7 (C7), 111.4 (C14), 118.9 (C11), 120.0 (C12), 122.5 (C13), 123.1 (C8), 127.1 (C15), 136.2 (C10), 167.0 (C18), 201.1 (C16).
HRMS (ESI): Calc. for $[M+Na]^+$ $C_{19}H_{24}N_2O_5Na^+$ = 383.1577, Obs. = 383.1586.

**Methyl (*S*)-4-((tert-butoxycarbonyl)amino)-5-(1H-indol-3-yl)-2,2-dimethyl-3-oxopentanoate**

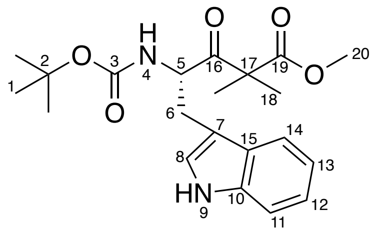

Prepared as per general procedure 4 – with methyl (*S*)-4-((*tert*-butoxycarbonyl)amino)-5-(1*H*-indol-3-yl)-3-oxopentanoate (0.648 g, 1.8 mmol, 1 equiv.), yielding the desired product as yellow solid (0.616 g, 1.59 mmol 88%).

$\delta_H$ (400 MHz, Chloroform-*d*) 1.30 (3H, s, H18), 1.36 (12H, s, H1, H18), 3.07 (1H, dd, *J* 7.0, 15.0 Hz, H6), 3.21 (1H, dd, *J* 6.0, 15.0 Hz, H6), 3.57 (3H, s, H20), 4.82 (1H, br. d, H4), 4.96 (1H, m, H5) 7.01 (1H, s, H8), 7.16 (2H, m, H12, H13), 7.34 (1H, d, *J* 8.0, H11), 7.60 (1H, d, *J* 8.0, H14), 8.06 (1H, br. s, H9).

$\delta_C$ (101 MHz, Chloroform-*d*) 21.9 (C18), 22.3 (C18), 28.2 (C6), 28.4 (C1), 52.5 (C20), 54.8 (C17), 55.9 (C5), 80.2 (C2), 110.4 (C7), 111.2 (C14), 118.9 (C11), 119.8 (C12), 120.1 (C10), 122.3 (C13), 123.1 (C8), 135.7 (15), 173.1 (C19), 207.9 (C16).

HRMS (ESI): Calc. for $[M+Na]^+$ $C_{21}H_{28}N_2O_5Na^+$ = 411.1890, Obs. = 411.1891.

**methyl (*S*)-4-amino-5-(1*H*-indol-3-yl)-2,2-dimethyl-3-oxopentanoate**

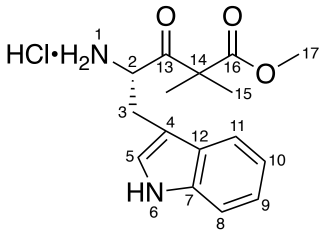

Prepared as per general procedure 5 – with methyl (*S*)-4-((*tert*-butoxycarbonyl)amino)-5-(1*H*-indol-3-yl)-2,2-dimethyl-3-oxopentanoate (0.756 g, 1.95 mmol, 1 equiv.), yielding the final product as a brown powder (0.632 g, 1.95 mmol, 99%).

$\delta_H$ (400 MHz, Chloroform-*d*) 1.44 (3H, s, H15), 1.53 (3H, s, H15), 2.99 (1H, m, H3). 3.27 (1H, m, H3), 3.71 (3H, s, H17), 4.68 (1H, br. s, H2), 7.10 (3H, m, H5, H9, H10), 7.29 (2H, m, H8, H11), 8.07 (3H, br. s, H6, H1).

$\delta_C$ (101 MHz, Chloroform-*d*) 22.7 (C15), 22.9 (C15), 26.6 (C3), 53.3 (C17), 55.0 (C14), 55.9 (C2), 105.5 (C4), 112.2 C11), 118.1 (C8), 120.5 (C9), 122.0 (C10), 126.8 (C5), 128.1 (C12), 136.5 (C7), 173.2 (C16), 204.4 (C13).

LRMS (ESI): Calc. for $[M+H]^+$ $C_{16}H_{21}N_2O_3^+$ = 289.2, Obs. = 289.2.

**methyl (*S*)-2-((*tert*-butoxycarbonyl)amino)-3-(cyclopent-1-en-1-yl)propanoate**

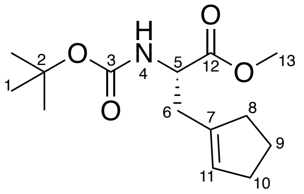

To a solution of cyclopentanone (**48**) (0.80 mL, 8.8 mmol) in anhydrous DCM (30 mL) was added Na2CO3 (1.41 g, 13.3 mmol, 1.5 equiv.) and triflic anhydride (5.0 g, 17.72 mmol, 2 equiv.) dropwise at – 10 °C under an argon atmosphere. The cooling bath was removed, and the reaction stirred overnight before it was quenched with water (30 mL) and extracted with DCM (3 × 30 mL). The combined organic layers were washed with brine (90 mL), dried with MgSO4, filtered, and concentrated in *vacuo* to provide crude cyclopent-1-en-1-yl trifluoromethanesulfonate as black liquid (1.05 g, 4.86 mmol, 55%) which was used in the next step without further purification.

To a solution of zinc dust (1.60 g, 24.3 mmol, 5 equiv.) in DMF (6 mL) was added TMSCl (0.62 mL, 4.9 mmol, 1 equiv.) dropwise and the mixture was stirred at room temperature for 1 hour. The upper clear layer was removed, the bottom layer was washed with DMF (2 × 10 mL), cooled to 0 °C and *N*-Boc-β-iodoalanine-OMe (1.61 g, 4.90 mmol, 1 equiv.), cyclopent-1-en-1-yl trifluoromethanesulfonate (1.05 g, 4.90 mmol), and Pd(dppf)Cl2 (0.671 g, 0.970 mmol, 0.02 equiv.) was added. The reaction mixture was stirred at room temperature overnight, quenched with brine (10 mL) and extracted with EtOAc (3 × 10 mL). The combined organic layers were washed with brine (30 mL), dried with MgSO4, filtered, and concentrated in *vacuo*. The crude product was purified by silica gel chromatography (EtOAc/hexane, 1:99 to 1:9) to yield the product as a yellow oil (0.898 g, 3.33 mmol, 69%).
$\delta_H$ (400 MHz, Chloroform-*d*) 1.44 (9H, s, H1), 1.86 (2H, quint., *J* 7.0, H9), 2.22 (2H, m, H8), 2.29 (2H, m, H10), 2.54 (2H, m, H6), 3.72 (3H, m, H13), 4.41 (1H, m, H5), 4.94 (1H, br d, *J* 6.0, H4), 5.46 (1H, s, H11).
$\delta_C$ (101 MHz, Chloroform-*d*) 23.7 (C9), 28.5 (C1), 32.6 (C10), 34.4 (C6), 34.9 (C8), 52.2 (C5), 53.4 (C13), 80.8 (C2), 128.5 (C11), 139.7 (C7), 173.3 (C12).
HRMS (ESI): Calc. for $[M+Na]^+$ $C_{14}H_{23}NO_4Na^+$ = 292.1519, Obs. = 292.1515.

**(*S*)-2-((*tert*-butoxycarbonyl)amino)-3-(cyclopent-1-en-1-yl)propanoic acid**

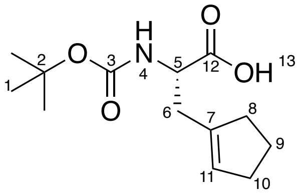

To a solution of methyl (*S*)-2-((*tert*-butoxycarbonyl)amino)-3-(cyclopent-1-en-1-yl)propanoate (0.338 g, 1.25 mmol) in water/methanol (2:1, 10 mL) was added LiOH (0.0901 g, 3.76 mmol, 3 equiv.) and the reaction was stirred at room temperature overnight. Methanol was removed *in vacuo*, the aqueous layer washed with DCM (10 mL), acidified with HCl (1 M, aqueous) to pH 3, and extracted with DCM (3 × 10 mL). The combined organic layers were dried with MgSO4, filtered, and concentrated in *vacuo* to provide the product **51** as opaque oil (0.320 g, 1.25 mmol, 100%).

$\delta_H$ (400 MHz, Chloroform-*d*) 1.44 (9H, s, H1), 1.87 (2H, quint., *J* 6.5, H9), 2.23 (2H, m, H8), 2.30 (2H, m, H10), 2.54 (2H, m, H6), 4.41 (1H, m, H5), 4.94 (1H, br. d, *J* 6.0, H4), 5.50 (1H, s, H11).

$\delta_C$ (101 MHz, Chloroform-*d*) 23.6 (C9), 28.4 (C1), 32.6 (C10), 33.9 (C6), 34.8 (C8), 52.1 (C5), 80.4 (C2), 128.7 (C11), 138.8 (C7), 177.5 (C12).

HRMS (ESI): Calc. for $[M-H]^-$ $C_{13}H_{20}NO_4^+$ = 254.1398, Obs. = 254.1401.

**methyl (*S*)-4-((*tert*-butoxycarbonyl)amino)-5-(cyclopent-1-en-1-yl)-3-oxopentanoate**

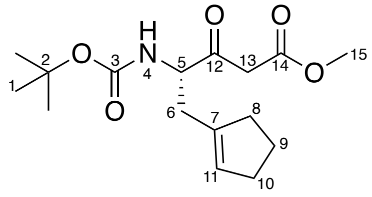

Prepared as per general procedure 3 with methyl (*S*)-4-((*tert*-butoxycarbonyl)amino)-5-(cyclopent-1-en-1-yl)-3-oxopentanoate (0.297 g, 1.17 mmol, 1 equiv.), yielding the desired product as a yellow oil (0.321 g, 1.03 mmol, 88%).

$\delta_H$ (400 MHz, Chloroform-*d*) 1.43 (9H, s, H1), 1.86 (2H, quint., *J* 7.0, H9), 2.22 (2H, m, H8), 2.30 (2H, m, H10), 2.52 (2H, m, H6), 3.50-3.60 (2H, m, H13), 3.73 (3H, s, H15), 4.41 (1H, m, H5), 4.96 (1H, br. s, H4), 5.47 (1H, s, H11).

$\delta_C$ (101 MHz, Chloroform-*d*) 23.6 (C9), 28.4 (C1), 32.6 (C10), 32.8 (C6), 34.9 (C8), 46.3 (C13), 52.6 (C15), 58.3 (C5), 80.4 (C2), 128.7 (C11), 139.0 (C7), 167.6 (C14), 202.4 (C12).

HRMS (ESI): Calc. for $[M+Na]^+$ $C_{16}H_{25}NO_5Na^+$ = 334.1625, Obs. = 334.1620.

**methyl (*S*)-4-((*tert*-butoxycarbonyl)amino)-5-(cyclopent-1-en-1-yl)-2,2-dimethyl-3-oxopentanoate**

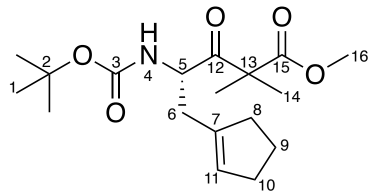

Prepared as per general procedure 4 – with methyl (*S*)-4-((*tert*-butoxycarbonyl)amino)-5-(cyclopent-1-en-1-yl)-3-oxopentanoate (0.316 g, 1.02 mmol, 1 equiv.), yielding the desired product as yellow oil (0.328 g, 0.966 mmol 95%).

$\delta_H$ (400 MHz, Chloroform-*d*) 1.40 (15H, s, H1 & H14), 1.84 (2H, quint., *J* 7.0, H9), 2.21 (2H, m, H8), 2.30 (2H, m, H10), 2.45 (2H, m, H6), 3.72 (3H, s, H16), 4.70 (1H, br. s, H5), 5.44 (1H, s, H11).

$\delta_C$ (101 MHz, Chloroform-*d*) 22.1 (C14), 22.5 (C14), 23.7 (C9), 28.4 (C1), 32.7 (C10), 34.6 (C6), 34.9 (C8), 52.7 (C16), 54.3 (C5), 79.9 (C2), 128.2 (C11), 134.7 (C14), 139.0 (C7), 173.8 (C15), 208.1 (C12).

HRMS (ESI): Calc. for $[M+Na]^+$ $C_{18}H_{29}NO_5Na^+$ = 362.1938, Obs. = 362.1934.

**methyl (*S*)-4-amino-5-(cyclopent-1-en-1-yl)-2,2-dimethyl-3-oxopentanoate**

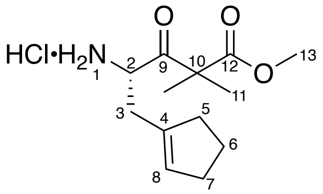

Prepared as per general procedure 5 – with Methyl (*S*)-4-((tert-butoxycarbonyl)amino)-2,2,5-trimethyl-3-oxohexanoate (0.311 g, 0.916 mmol, 1 equiv.), yielding the final product as a brown oil (0.252 g, 0.915 mmol, 99%).

$\delta_H$ (400 MHz, Chloroform-*d*) 1.46 (3H, s, H11), 1.54 (3H, s, H11), 1.90 (2H, m, H6), 2.18 (2H, m, H5), 2.34 (2H, m, H7), 2.68 (2H, m, H3), 3.76 (3H, s, H13), 4.61 (1H, br. s, H2), 5.80 (1H, s, H8).

$\delta_C$ (101 MHz, Chloroform-*d*) 22.8 (C11), 23.0 (C11), 23.5 (C6), 32.3 (C7), 32.9 (C3), 34.8 (C5), 53.2 (C13), 54.4 (C2), 132.9 (C8), 134.7 (C11), 136.0 (C4), 172.8 (C12), 203.9 (C9).

HRMS (ESI): Calc. for $[M+H]^+$ $C_{13}H_{22}NO_3^+$ = 240.1594, Obs. = 240.1588.

##### 1.2.4 Chemical synthesis of functionalised butyryl α-dimethyl-β-keto-ester substrates

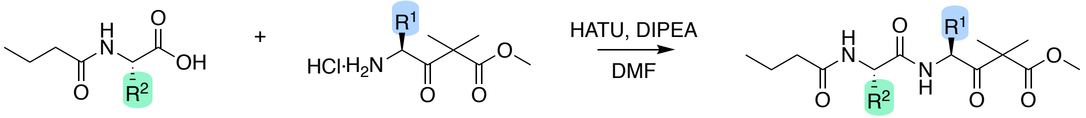

*Scheme S1.3: General synthetic scheme for synthesis of butyryl α-dimethyl-β-keto-ester substrates.*

**Methyl (*S*)-4-((*S*)-2-butyramido-3-hydroxypropanamido)-2,2,6-trimethyl-3-oxoheptanoate (22)**

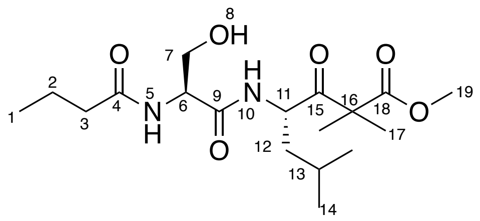

Prepared as per general procedure 6 - with butyryl-L-serine (0.12 g, 0.68 mmol, 1 equiv.), purified by silica gel chromatography (1:1 to 9:1 EtOAc:hexane) yielding the product as a white powder (0.748 g, 0.20 mmol, 29%).

$\delta_H$ (400 MHz, Methanol-$d_4$) 0.78 – 1.08 (9H, m, H1 & H14), 1.36 (3H, s, H17), 1.39 – 1.44 (4H, s, H12 & H17), 1.52 (1H, td, *J* 12.5, 4.0, H12), 1.58 – 1.77 (3H, m, H2 & H13), 2.24 (2H, t, *J* 7.5, H3), 3.55 – 3.82 (5H, m, H7 & H19), 4.41 (1H, t, *J* 5.5, H6), 4.95 (1H, dd, *J* 11.0, 3.5, H11).

$\delta_C$ (101 MHz, Methanol-$d_4$) 14.0 (C1), 20.2 (C2), 21.7 (C14), 22.6 (C17), 22.6 (C17), 23.8 (C14), 25.7 (C13), 38.7 (C3), 42.0 (C12), 53.2 (C19), 53.6 (C11), 56.1 (C16), 56.6 (C6), 62.9 (C7), 172.0 (C9), 175.0 (C18), 176.3 (C4), 208.5 (C15).

HRMS (ESI): Calc. for $[M+Na]^+$ $C_{18}H_{32}N_2O_6Na^+$ = 395.2153, Obs. = 395.2147.

**Methyl (*S*)-4-((*S*)-2-butyramidopropanamido)-2,2,6-trimethyl-3-oxoheptanoate (23)**

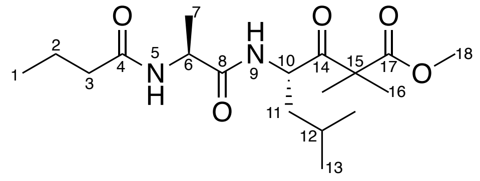

Prepared as per general procedure 6 - with butyryl-L-alanine (0.050 g, 0.31 mmol, 1 equiv.), purified by silica gel chromatography (1:1 to 9:1 EtOAc:hexane) yielding the product as a white powder (0.084 g, 0.24 mmol, 75%).

$\delta_H$ (400 MHz, Methanol-$d_4$) 0.88 – 0.98 (9H, m, H1 & H13), 1.29 (3H, d, *J* 6.5, H7), 1.36 (3H, s, H16), 1.39 (3H, s, H16), 1.41 – 1.44 (1H, m, H11), 1.51 (1H, td, *J* 12.5, 11.0, 4.0, H11), 1.57 – 1.71 (3H, m, H2 & H12), 2.19 (2H, t, *J* 7.0, H3), 3.72 (3H, s, H18), 4.32 (1H, q, *J* 7.0, H6), 4.91 (1H, dd, *J* 11.0, 3.5, H10).

$\delta_C$ (101 MHz, Methanol-$d_4$) 14.0 (C1), 17.8 (C7), 20.3 (C2), 21.7 (C13), 22.6 (C16), 22.7 (C16), 23.9 (C13), 25.7 (C12), 38.6 (C3), 41.9 (C11), 50.1 (C6), 53.1 (C18), 53.7 (C10), 56.1 (C15), 174.5 (C8), 174.9 (C18), 175.9 (C4), 208.5 (C14).

HRMS (ESI): Calc. for $[M+H]^+$ $C_{18}H_{33}N_2O_5^+$ = 357.2384, Obs. = 357.2383.

**Methyl (*S*)-4-((2*S*,3*R*)-2-butyramido-3-hydroxybutanamido)-2,2,6-trimethyl-3-oxoheptanoate (24)**

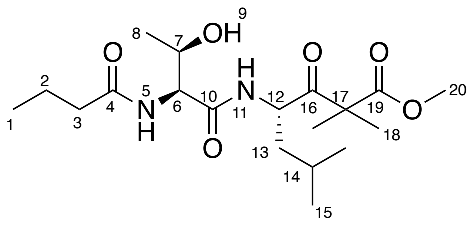

Prepared as per general procedure 6 - with butyryl-L-threonine (0.05 g, 0.26 mmol, 1 equiv.), purified by silica gel chromatography (4:1 EtOAc:hexane) yielding the product as a white powder (0.034 g, 0.09 mmol, 28%).

$\delta_H$ (400 MHz, Methanol-$d_4$) 0.84 – 1.04 (9H, m, H1 & H15), 1.16 (3H, d, *J* 6.5, H8), 1.32 – 1.39 (4H, m, H13 & H18), 1.41 (3H, s, H18), 1.46 – 1.59 (1H, m, H13), 1.58 – 1.75 (1H, m, H14), 2.27 (2H, t, *J* 7.5, H3), 3.72 (3H, s, H20), 4.05 (1H, quint., *J* 6.0, H7), 4.29 (1H, d, *J* 4.5, H6), 4.97 (1H, dd, *J* 11.0, 3.0, H12).

$\delta_C$ (101 MHz, Methanol-$d_4$) 14.0 (C1), 20.3 (C2), 20.3 (C8), 21.6 (C15), 22.6 (C18), 22.6 (C18), 23.8 (C15), 25.7 (C14), 38.8 (C3), 42.0 (C13), 53.2 (C20), 53.5 (C12), 56.2 (C17), 60.0 (C6), 68.3 (C7), 172.2 (C4), 174.9 (C19), 176.3 (C10), 208.6 (C16).

HRMS (ESI): Calc. for $[M+H]^+$ $C_{19}H_{35}N_2O_6^+$ = 387.2490, Obs. = 387.2492.

**Methyl (*S*)-4-((*S*)-3-((tert-butyldimethylsilyl)oxy)-2-butyramidopropanamido)-2,2,6-trimethyl-3-oxoheptanoate (25)**

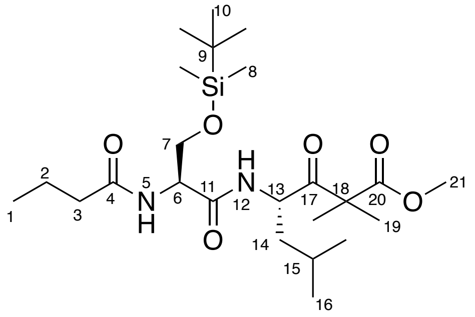

Prepared as per general procedure 6 - with *O*-(tert-butyldimethylsilyl)-*N*-butyryl-L-serine (0.15 g, 0.52 mmol, 1 equiv.), purified by silica gel chromatography (1:1 EtOAc:hexane) yielding the product as a white powder (0.141 g, 0.290 mmol, 56%).

$\delta_H$ (400 MHz, Methanol-$d_4$) 0.09 (6H, s, H8), 0.90 (9H, s, H10), 0.91 – 0.97 (9H, m, H1 & H16), 1.36 (3H, s, H19), 1.38 – 1.43 (4H, m, H14 & H19), 1.51 (1H, ddd, *J* 14.5, 11.0, 4.0, H14), 1.58 – 1.72 (3H, m, H2 & H15), 2.24 (2H, t, *J* 7.5, H3), 3.72 (3H, s, H21), 3.75 – 3.94 (2H, m, H7), 4.45 (1H, t, *J* 6.0, H6), 4.97 (1H, dd, *J* 11.0, 3.0, H13)

$\delta_C$ (101 MHz, Methanol-$d_4$) -5.4 (C8), -5.3 (C8), 14.0 (C1), 19.2 (C9), 20.3 (C2), 21.7 (C16), 22.6 (C19), 22.7 (C19), 23.8 (C16), 25.7 (C15), 26.4 (C10), 38.7 (C3), 42.1 (C14), 53.1 (C21),

53.5 (C13), 56.2 (C18), 56.4 (C6), 64.2 (C7), 171.6 (C11), 174.9 (C20), 176.0 (C4), 208.3 (C17).

HRMS (ESI): Calc. for $[M+Na]^+$ $C_{24}H_{46}N_2O_6SiNa^+$ = 509.3017, Obs. = 509.3012.

**Methyl (*S*)-4-(2-butyramidoacetamido)-2,2,6-trimethyl-3-oxoheptanoate (26)**

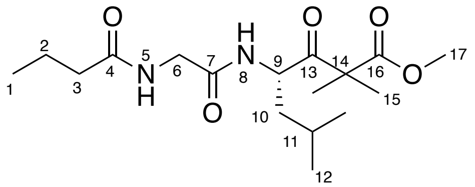

Prepared as per general procedure 6 - with butyrylglycine (0.085 g, 0.56 mmol, 1 equiv.), purified by silica gel chromatography (1:1 EtOAc:hexane) yielding the product as a yellow oil (0.060 g, 0.18 mmol, 30%).

$\delta_H$ (400 MHz, Methanol-*d*$_4$) 0.82 – 1.08 (9H, m, H1 & H12), 1.36 (3H, s, H15), 1.40 (3H, s, H15), 1.43 – 1.51 (2H, m, H10), 1.56 – 1.76 (3H, m, H2 & H10), 2.23 (2H, t, *J* 7.5, H3), 3.71 (3H, s, H17), 3.78 (1H, d, *J* 16.5, H6), 3.86 (1H, d, *J* 16.5, H6), 4.94 (1H, dd, *J* 11.0, 3.5, H9).

$\delta_C$ (101 MHz, Methanol-*d*$_4$) 14.0 (C1), 20.1 (C2), 21.7 (C12), 22.6 (C15), 23.8 (C12), 25.8 (C11), 38.7 (C3), 41.9 (C10), 43.2 (C6), 53.1 (C17), 53.7 (C9), 56.1 (C14), 171.0 (C7), 174.9 (C17), 176.5 (C4), 208.5 (C13).

HRMS (ESI): Calc. for $[M+H]^+$ $C_{17}H_{31}N_2O_5^+$ = 343.2227, Obs. = 343.2229.

**methyl (*S*)-4-((*S*)-2-butyramido-3-methylbutanamido)-2,2,6-trimethyl-3-oxoheptanoate (27)**

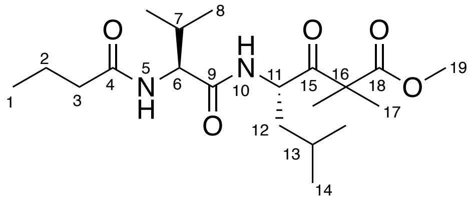

Prepared as per general procedure 6 - with butyryl-*L*-valine (0.075 g, 0.40 mmol, 1 equiv.), purified by silica gel chromatography (3:2 hexane:EtOAc) yielding the product as a white powder (0.034 g, 0.09 mmol, 28%).

$\delta_H$ (400 MHz, Methanol-*d*$_4$) 0.86 – 1.00 (15H, m, H1, H8 & H14), 1.35-1.39 (4H, m, H12 & 17), 1.41 (3H, s, H17), 1.52 (1H, ddd, *J* 15.0, 3.5, H12), 1.58 – 1.70 (3H, m, H2 & H13), 2.01 (1H, sept, *J* 6.5, H7), 2.22 (2H, t, *J* 7.5, H3), 3.72 (3H, s, H19), 4.12 (1H, d, *J* 8.0, H6), 4.95 (1H, d, *J* 11.0, H11).

$\delta_C$ (101 MHz, Methanol-*d*$_4$) 14.0 (C1), 18.9 (C8), 19.7 (C8), 20.4 (C2), 21.5 (C14), 22.6 (C17), 22.7 (C17), 23.8 (C14), 25.8 (C13), 31.6 (7), 38.7 (C5), 41.9 (C12), 53.1 (C19), 53.3 (C11), 56.3 (C16), 60.2 (C6), 173.2 (C9), 174.8 (19), 176.1 (C14), 208.7 (C15).

HRMS (ESI): Calc for $[M+H]^+$ $C_{20}H_{37}N_2O_5^+$ = 385.2697, Obs. = 385.2692.

**Methyl (S)-4-((S)-2-butyramido-3-phenylpropanamido)-2,2,6-trimethyl-3-oxoheptanoate (28)**

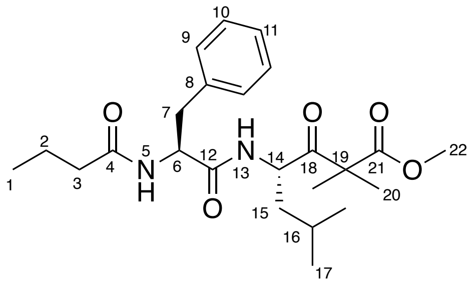

Prepared as per general procedure 6 - with butyryl-L-phenylalanine (0.12 g, 5.1 mmol, 1 equiv.), purified by silica gel chromatography (1:1 EtOAc:hexane) yielding the product as a white powder (0.105 g, 2.43 mmol, 48%).

$\delta_H$ (400 MHz, Methanol-$d_4$) 0.80 (3H, t, *J* 7.5, H1), 0.87 – 0.96 (6H, m, H17), 1.33 (3H, s, H20), 1.35 (3H, s, H20), 1.40 (1H, dt, *J* 11.0, 3.5, H15), 1.45 – 1.55 (3H, m, H2 & H15), 1.59 – 1.70 (1H, m, H16), 2.11 (2H, t, *J* 7.5, H3), 2.82 (1H, dd, *J* 14.0, 9.5, H7), 3.09 (1H, dd, *J* 14.0, 5.5, H7), 3.71 (3H, s, H22), 4.64 (1H, dd, *J* 9.5, 5.5, H6), 4.92 (1H, dd, *J* 11.0, 3.0, H14), 7.13 – 7.32 (5H, m, H9, H10 & H11).

$\delta_C$ (101 MHz, Methanol-$d_4$) 13.8 (C1), 20.3 (C2), 21.6 (C17), 22.5 (C20), 22.7 (C20), 23.8 (C17), 25.7 (C16), 38.6 (C7), 38.7 (C3), 42.0 (C15), 53.1 (C22), 53.5 (C14), 55.6 (C6), 56.1 (C19), 127.7 (C11), 129.4 (C10), 130.3 (C9), 138.4 (C8), 173.1 (C12), 174.9 (C21), 175.9 (C4), 208.4 (C18).

HRMS (ESI): Calc. $[M+H]^+$ $C_{24}H_{37}N_2O_5^+$ = 433.2697, Obs. = 433.2692.

**Methyl (*S*)-4-((*S*)-2-butyramido-3-methoxypropanamido)-2,2,6-trimethyl-3-oxoheptanoate (29)**

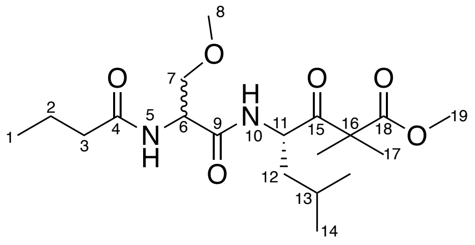

Prepared as per general procedure 6 - with *N*-butyryl-*O*-methyl-L-serine (0.028 g, 0.148 mmol, 1 equiv.), purified by silica gel chromatography (7:3 EtOAc:hexane) yielding the product as a white powder as a 2.3:1.0 ratio of diastereomers. (0.037 g, 0.094 mmol, 64%).

HRMS (ESI): Calc. for $[M+H]^+$ $C_{19}H_{35}N_2O_6^+$ = 387.2490, Obs. = 387.2490.

Major diastereomer

δH (400 MHz, Methanol-$d_4$) 0.80 – 1.04 (9 H, m, H1 & H14), 1.36 (3H, s, H17), 1.38 – 1.41 (4H, s, H12 & H17), 1.52 (1H, ddd, *J* 14.5, 11.0, 3.5, H12), 1.57 – 1.72 (3H, m, H2 & H13), 2.23 (2H, t, *J* 7.5, H3), 3.32 (3H, s, H8), 3.49 – 3.60 (2H, m, H7), 3.70 (3H, s, H19), 3.71 (2H, s), 4.52 (1H, t, *J* 5.5, H6), 4.95 (1H, dd, *J* 11.0, 3.0, H11).

δC (101 MHz, Methanol-$d_4$) 14.0 (C1), 20.2 (C2), 21.7 (C14), 22.5 (C17), 22.6 (C17), 23.8 (C14), 25.6 (C13), 38.6 (C3), 42.1 (C12), 53.1 (C19), 53.5 (C11), 54.6 (C6), 56.2 (C16), 59.2 (C8), 72.9 (C7), 171.7 (C8), 174.9 (C18), 176.2 (C4), 208.4 (C15).

Minor diastereomer

δH (400 MHz, Methanol-$d_4$) The only $^1$H NMR signal with a different chemical shift for this diastereomer is 3.70 (3H, s, H19).

δC (101 MHz, Methanol-d4) 14.0 (C1), 20.2 (C2), 21.6 (C14), 22.5 (C17), 22.6 (C17), 23.8 (C14). 25.7 (C13), 38.6 (C3), 42.0 (C12), 53.0 (C19), 53.5 (C11), 54.4 (C6), 56.2 (C16), 59.2 (C8), 73.1 (C7), 171.5 (C8), 174.9 (C18), 176.1 (C4), 208.3 (C15).

**Methyl (S)-4-((S)-2-butyramido-3-hydroxypropanamido)-2,2,6-trimethyl-3-oxohept-6-enoate (30)**

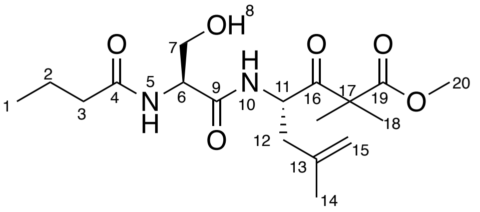

Prepared as per general procedure 6 - with butyryl-L-serine (0.055 g, 0.314 mmol, 1 equiv.), purified by silica gel chromatography (1:1 EtOAc:hexane) yielding the product as a white powder (0.029 g, 0.078 mmol, 25%).

δH (400 MHz, Methanol-$d_4$) 0.95 (3H, t, *J* 7.5, H1), 1.37 (3H, s, H18), 1.41 (3H, s, H18), 1.64 (2H, sext., *J* 7.5, H2), 1.72 (3H, s, H14), 2.17 – 2.29 (3H, m, H3 & H14), 2.44 (1H, dd, *J* 14.5, 4.0, H12), 3.60 – 3.85 (5H, m, H7 & H20), 4.42 (1H, t, *J* 6.0, H7), 4.77 (1H, d, *J* 19.5, H15), 5.05 (1H, dd, *J* 10.0, 4.0, H11).

δC (101 MHz, Methanol-$d_4$) 14.0 (C1), 20.2 (C2), 22.2 (C14), 22.4 (C18), 22.6 (C18), 38.8 (C3), 41.1 (C12), 53.2 (C20), 53.4 (C11), 56.2 (C17), 56.4 (C6), 62.9 (C7), 114.7 (C15), 142.0 (C13), 171.8 (C9), 175.0 (C19), 176.2 (C4), 207.8 (C16).

HRMS (ESI): Calc. for $[M+H]^+$ $C_{18}H_{31}N_2O_6^+$ = 371.2177, Obs. = 371.2173.

**Methyl (*S*)-4-((*S*)-2-butyramido-3-hydroxypropanamido)-2,2-dimethyl-3-oxo-5-phenylpentanoate (31)**

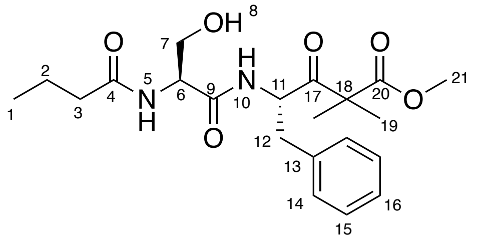

Prepared as per general procedure 6 - with butyryl-L-serine (0.15 g, 0.86 mmol, 1 equiv.), purified by silica gel chromatography (7:3 EtOAc:hexane) yielding the product as a pale-yellow powder (0.125 g, 3.08 mmol, 36%).

$\delta_H$ (400 MHz, Methanol-$d_4$) 0.95 (3H, t, *J* 7.5, H1), 1.29 (3H, s, H19), 1.34 (3H, s, H19), 1.64 (2H, sext., *J* 7.5, H2), 2.22 (2H, t, *J* 7.5, H3), 2.80 (1H, dd, *J* 14.0, 8.5, H12), 3.11 (1H, dd, *J* 14.0, 6.0, H12), 3.56 – 3.85 (5H, m, H7 & H21), 4.38 (1H, t, *J* 5.5, H6), 5.13 (1H, dd, *J* 8.5, 6.0, H11), 7.11 – 7.24 (3H, m, H14 & H16), 7.23 – 7.36 (2H, m, H15).

$\delta_C$ (101 MHz, Methanol-$d_4$) 14.0 (C1), 20.2 (C2), 22.1 (C19), 22.4 (C19), 38.7 (C3), 38.9 (C12), 53.2 (C21), 56.2 (C11), 56.4 (C11), 56.5 (C6), 62.9 (C7), 127.8 (C14), 129.4 (C16), 130.6 (C15), 138.2 (C13), 171.7 (C9), 174.9 (C20), 176.1 (C4), 207.4 (C17).

HRMS (ESI): Calc. for $[M+Na]^+$ $C_{21}H_{30}N_2O_6Na^+$ = 429.1996, Obs. = 429.1999.

**Methyl (*S*)-4-((*S*)-2-butyramido-3-hydroxypropanamido)-2,2,5-trimethyl-3-oxohexanoate (32)**

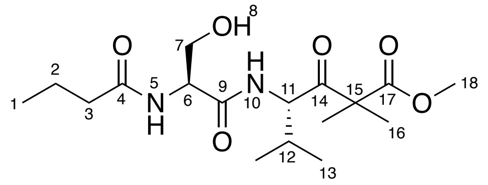

Prepared as per general procedure 6 - with butyryl-L-serine (0.10 g, 0.57 mmol, 1 equiv.), purified by silica gel chromatography (1:1 to 9:1 EtOAc:hexane) yielding the product as a white powder (0.049 g, 0.13 mmol, 22%).

$\delta_H$ (400 MHz, Methanol-$d_4$) 0.85 (3H, d, *J* 6.5, H13), 0.88 (3H, d, *J* 7.0, H13), 0.95 (2H, t, *J* 7.5, H1), 1.37 (3H, s, H16), 1.40 (3H, s, H16), 1.65 (2H, sext., *J* 7.5, H2), 2.10-2.20 (1H, m, H12), 2.25 (2H, t, *J* 7.0, H3), 3.71 (3H, s, H18), 3.73 – 3.84 (2H, m, H7), 4.44 (1H, t, *J* 5.5, H6), 4.81 (1H, d, *J* 6.5, H11).

$\delta_C$ (101 MHz, Methanol-$d_4$) 14.0 (C1), 17.3 (C13), 20.2 (C2), 20.2 (C13), 22.5 (C16), 22.8 (C16), 31.1 (C12), 38.7 (C3), 53.1 (C18), 56.6 (C6), 59.7 (C11), 62.6 (C7), 172.4 (C9), 174.8 (C17), 176.3 (C4), 207.7 (C14).

HRMS (ESI): Calc. for $[M+H]^+$ $C_{17}H_{31}N_2O_6^+$ = 359.2177, Obs. = 359.2176.

**Methyl (S)-4-((S)-2-butyramido-3-hydroxypropanamido)-2,2-dimethyl-3-oxopentanoate (33)**

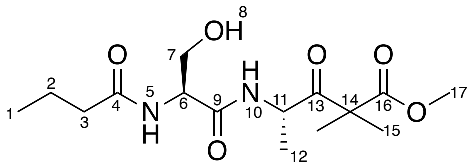

Prepared as per general procedure 6 - with butyryl-L-serine (0.053 g, 0.303 mmol, 1 equiv.), purified by silica gel chromatography (EtOAc) yielding the product as a clear oil (0.013 g, 0.039 mmol, 13%).

$\delta_H$ (400 MHz, Methanol-$d_4$) 0.95 (3H, t, $J$ 7.5, H1), 1.25 – 1.28 (3H, m, H12), 1.38 (3H, s, H15), 1.41 (3H, s, H15), 1.65 (2H, sext., $J$ 7.5, H2), 2.25 (2H, t, $J$ 7.5, H3), 3.68 – 3.77 (5H, m, H7 & H17), 4.40 (1H, t, $J$ 5.5, H6), 4.88 – 4.98 (1H, m, H11).

$\delta_C$ (101 MHz, Methanol-$d_4$) 14.0 (C1), 18.2 (C12), 20.2 (C2), 22.6 (C15), 22.7 (C15), 38.7 (C3), 51.3 (C11), 53.2 (C17) ,56.0 (C4), 56.6, (C6), 63.0 (C7), 171.7 (C8), 175.1 (C17), 176.3 (C4), 208.8 (C13).

HRMS (ESI): Calc. for $[M+Na]^+$ $C_{15}H_{26}N_2O_6Na^+$ = 353.1683, Obs. = 353.1679.

**methyl (*S*)-4-((*S*)-2-butyramido-3-hydroxypropanamido)-5-(1*H*-indol-3-yl)-2,2-dimethyl-3-oxopentanoate (34)**

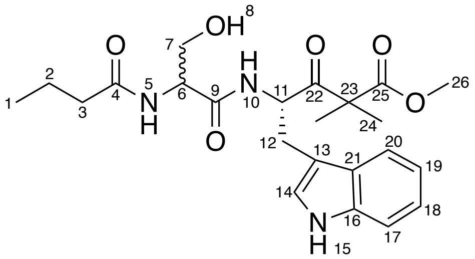

To a solution of butyryl-*L*-serine (0.0701 g, 0.398 mmol) in THF (7 mL) was added NMI (0.10 mL, 1.72 mmol, 3.0 equiv.) and TCFH (0.169 g, 0.603 mmol, 1.5 equiv.) and the solution was stirred for 15 minutes. Methyl (*S*)-4-amino-5-(1*H*-indol-3-yl)-2,2-dimethyl-3-oxopentanoate (0.0.131 g, 0.402 mmol, 1.0 equiv.) was added and the reaction was stirred overnight. The reaction mixture was quenched with 1 M aqueous HCl (10 mL) and THF was removed in *vacuo*. The aqueous layer was extracted with EtOAc (3 × 15 mL), and the combined organic layers were washed with brine (2 × 20 mL). The organic layer was dried with MgSO4, filtered, and concentrated in *vacuo*. The crude product was purified by silica gel chromatography (MeOH/DCM, 1:99 to 1:9) to yield the product as a brown oil as a 2.2:1.0 ratio of diastereomers (0.0462 g, 0.104 mmol, 26%).

Major diastereomer

$\delta_H$ (400 MHz, Methanol-$d_4$) 0.92 (3H, t, *J* 7.5, H1), 1.18 (3H, s, H24), 1.25 (3H, s, H24), 1.63 (2H, m, H2), 2.18 (2H, t, *J* 7.0, H3), 2.97 (1H, dd, *J* 14.5, 6.0, H12), 3.25 (1H, dd, *J* 14.0, 6.0, H12), 3.51 (1H,s, H26), 3.55 (1H, m, H7), 3.67 (1H, m, H7), 4.39 (1H, t, *J* 5.0, H6), 5.24 (1H, t, *J* 17.0, H11), 6.90-7.10 (3H, m, H12, H18 & H19), 7.30 (1H, d, *J* 6.5, H17), 7.58 (1H, d, *J* 7.5, H20).

$\delta_C$ (101 MHz, Methanol-$d_4$) 14.0 (C1), 20.2 (C2), 22.0 (C24), 22.3 (C24), 29.1 (C12), 38.7 (C3), 53.1 (C26), 55.3 (C11), 56.6 (C6), 56.9 (C23), 62.9 (C7), 110.6 (C13), 112.3 (C20), 119.3 (C17), 119.9 (C18), 122.4 (C16), 124.7 (C19), 128.8 (C14), 137.8 (C21), 171.7 (C9), 174.9 (C25), 176.6 (C4), 208.1 (C22).

Minor diastereomer

$\delta_H$ (400 MHz, Methanol-$d_4$) 0.96 (3H, t, *J* 7.5, H1), 1.20 (3H, s, H24), 1.28 (3H, s, H24), 1.63 (2H, m, H2), 2.25 (2H, t, *J* 7.5, H3), 2.97 (1H, dd, *J* 14.5, 6.0, H12), 3.25 (1H, dd, *J* 14.0, 6.0 Hz, H12), 3.51 (1H, s, H26), 3.55 (1H, m, H7), 3.67 (1H, m, H7), 4.39 (1H, t, *J* 5.0, H6), 5.24 (1H, t, *J* 17.0, H11), 6.90-7.10 (3H, m, H12, H18 & H19), 7.29 (1H, d, *J* 6.5, H17), 7.53 (1H, d, *J* 7.5, H20).

$\delta_C$ (101 MHz, Methanol-$d_4$) 14.0 (C1), 20.2 (C2), 22.1 (C24), 22.3 (C24), 29.1 (C12), 38.7 (C3), 53.1 (C26), 55.3 (C11), 56.4 (C6), 56.9 (C23), 62.9 (C7), 110.6 (C13), 112.3 (C20), 119.3 (C17), 120.0 (C18), 122.5 (C16), 124.7 (C19), 128.8 (C14), 137.8 (C21), 171.7 (C9), 174.9 (C25), 176.6 (C4), 208.1 (C22).

HRMS (ESI): Calc. for $[M+Na]^+$ $C_{23}H_{31}N_3O_6Na^+$ = 468.2105, Obs. = 468.2096.

**methyl (*S*)-4-((*S*)-2-butyramido-3-hydroxypropanamido)-5-(cyclopent-1-en-1-yl)-2,2-dimethyl-3-oxopentanoate (35)**

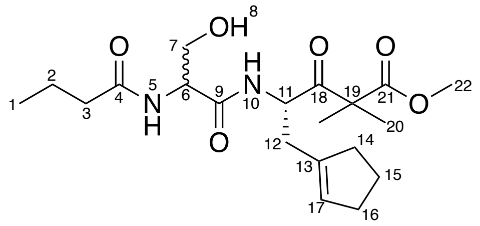

To a solution of butyryl-*L*-serine (0.0384 g, 0.221 mmol) in THF (7 mL) was added NMI (0.05 mL, 0.7 mmol, 3.0 equiv.) and TCFH (0.0923 g, 0.331 mmol, 1.5 equiv.) and the solution was stirred for 15 minutes. Amine **55c** (0.0607 g, 0.221 mmol, 1.0 equiv.) was added and the reaction was stirred overnight. The reaction mixture was quenched with 1 M aqueous HCl (10 mL) and THF was removed in *vacuo*. The aqueous layer was extracted with EtOAc (3 × 15 mL), and the combined organic layers were washed with brine (2 × 20 mL). The organic layer was dried with MgSO4, filtered, and concentrated in *vacuo*. The crude product was purified by silica gel chromatography (MeOH/DCM, 1:99 to 1:9) to yield the product as a brown oil as a 2.1:1.0 mixture of diastereomers (0.0152 g, 0.0383 mmol, 17%).

Major diastereomer

$δ_H$ (400 MHz, Chloroform-*d*) 0.93 (3H, t, *J* 9.5, H1), 1.38 (3H, s, H20), 1.42 (3H, s, H20), 1.64 (2H, quint., *J* 7.0, H2), 1.81 (2H, m, H15), 2.18 (2H, t, *J* 7.0, H3), 2.22 (2H, m, H14), 2.30 (2H, m, H16), 2.53 (2H, m, H12), 3.57 (1H, dd, *J* 19.0, 5.0, H7), 3.72 (3H, s, H22), 4.00 (1H, dd, *J* 12.0, 3.5, H7), 4.40 (1H, br. s, H6), 4.96 (1H, m, H11), 5.38 (1H, s, H17), 6.63 (1H, br. d, *J* 6.5 Hz, H5), 7.14 (1H, br. s, *J* 9.0, H10).

$δ_C$ (101 MHz, Chloroform-*d*) 13.8 (C1), 19.1 (C2), 22.3 (C20), 22.5 (C20), 23.6 (C15), 32.6 (C16), 33.8 (C12), 34.7 (C3), 38.3 (C14), 52.8 (C22), 53.3 (C11), 54.9 (C19), 62.7 (C7), 128.2 (C17), 139.1 (C13), 170.8 (C9), 174.0 (C21), 207.0 (C18).

Minor diastereomer

$δ_H$ (400 MHz, Chloroform-*d*) 0.93 (3H, t, *J* 9.5, H1), 1.20 (3H, s, H20), 1.24 (3H, s, H20), 1.64 (2H, quint., *J* 7.0, H2), 1.81 (2H, m, H15), 2.18 (2H, t, *J* 7.0, H3), 2.22 (2H, m, H14), 2.30 (2H, m, H16), 2.53 (2H, m, H12), 3.54 (1H, dd, *J* 19.0, 5.0, H7), 3.70 (3H, s, H22), 3.97 (1H, dd, *J* 12.0, 3.5, H7), 4.40 (1H, br. s, H6), 4.96 (1H, m, H11), 5.45 (1H, s, H17), 6.86 (1H, br. d, *J* 6.5, H5), 7.14 (1H, br. d, *J* 9.0, H10).

$δ_C$ (101 MHz, Chloroform-*d*) 13.8 (C1), 19.1 (C2), 20.3 (C20), 21.5 (C20), 23.5(C15), 32.6 (C16), 33.8 (C12), 34.7 (C3), 38.4 (C14), 52.8 (C22), 53.1 (C11), 55.0 (C19), 63.0 (C7), 128.7 (C17), 138.9 (C13), 170.5 (C9), 173.8 (C21), 207.0 (C18).

HRMS (ESI): Calc. for $[M+Na]^+$ $C_{20}H_{32}N_2O_6Na^+$ = 419.2153, Obs. = 419.2146.

##### 1.2.5 Synthesis of N-functionalised α-dimethyl-β-keto-ester substrates

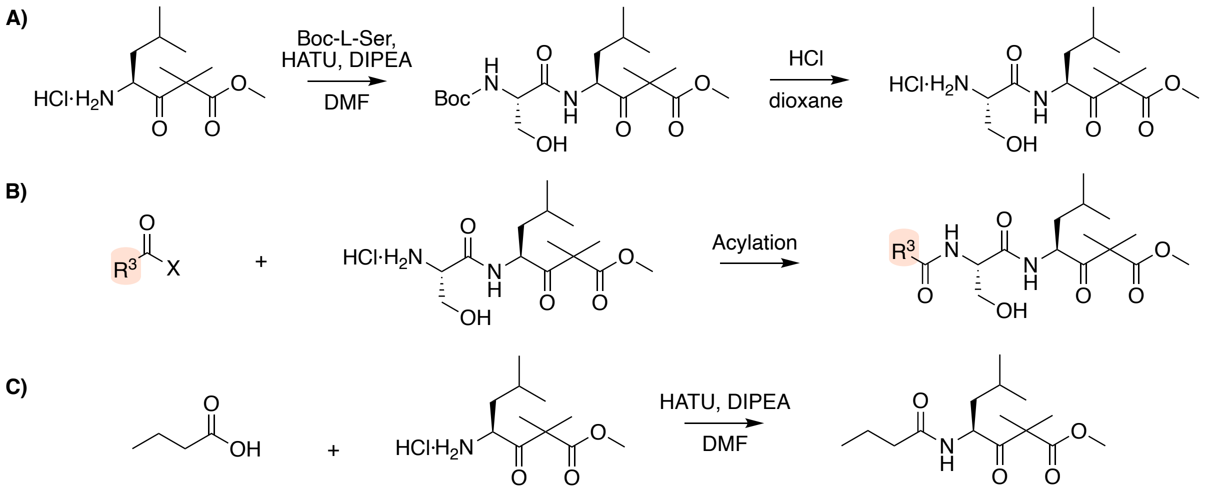

*Scheme S1.4: A) Synthesis of the common amino dipeptide fragment, B) synthesis of $R_3$-functionalised α-dimethyl-β-keto-ester substrates, C) synthesis of truncated substrate.*

**Methyl (*S*)-4-((*S*)-2-((*tert*-butoxycarbonyl)amino)-3-hydroxypropanamido)-2,2,6-trimethyl-3-oxoheptanoate (16)**

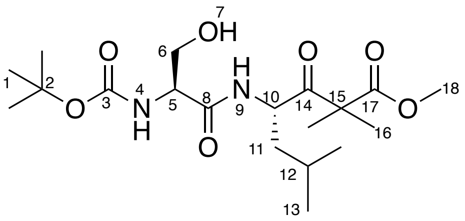

Prepared as per general procedure 6 - with (*tert*-butoxycarbonyl)-L-serine (0.371 g, 1.8 mmol, 1 equiv.), purified by silica gel chromatography (98:2 $CH_2Cl_2$:MeOH) yielding the product as a clear oil (0.267 g, 0.66 mmol, 37%).

$\delta_H$ (400 MHz, Chloroform-*d*) 0.77 – 0.99 (6H, m, H13), 1.39 (3H, s, H16), 1.40 – 1.57 (14H, m, H1, H11 & H16), 1.58 (1H, dq, *J* 13.0, 6.5, H12), 3.62 (1H, dd, *J* 11.0, 5.5, H6), 3.73 (3H, s, H18), 4.02 (1H, dd, *J* 11.5, 3.0, H6), 4.06 – 4.14 (1H, m, H5), 4.94 (1H, q, *J* 8.5, H10), 5.56 (1H, d, *J* 4.0, H4), 6.92 (1H, d, *J* 9.0, H9).

$\delta_C$ (101 MHz, Chloroform-*d*) 21.3 (C13), 22.2 (C16), 22.5 (C16), 23.6 (C13), 24.8 (C12), 28.4 (C1), 41.4 (C11), 52.8 (C18), 52.9 (C10), 54.7 (C5), 55.0 (C15), 62.8 (C6), 80.7 (C2), 156.8 (C3), 171.2 (C8), 173.6 (C17), 207.8 (C14).

HRMS (ESI): Calc for $[M+H]^+$ $C_{19}H_{35}N_2O_7^+$ = 403.2439, Obs. = 403.2437.

**methyl (*S*)-4-((*S*)-3-hydroxy-2-pivalamidopropanamido)-2,2,6-trimethyl-3-oxoheptanoate hydrochloride**

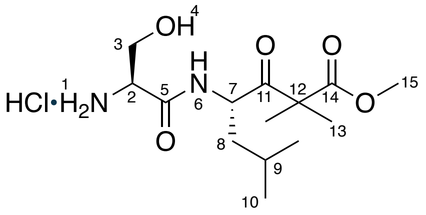

Prepared as per general procedure 5 – with Methyl (*S*)-4-((*S*)-2-((*tert*-butoxycarbonyl)amino)-3-hydroxypropanamido)-2,2,6-trimethyl-3-oxoheptanoate (0.306 g, 0.76 mmol, 1 equiv.), yielding the final product as a white powder (0.247 g, 0.73 mmol, 96%).

$\delta_H$ (400 MHz, Methanol-*d*$_4$) 0.93 (3H, d, *J* 4.0, H10), 0.95 (3H, d, *J* 4.5, H10), 1.38 (3H, s, H13), 1.40 (3H, s, H13), 1.41 – 1.47 (1H, m, H8), 1.53 (1H, ddd, *J* 14.5, 11.0, 4.0, H8), 1.67 (1H, ddt, *J* 13.5, 6.5, 4.0, H9), 3.71 – 3.73 (4H, m, H3 and H15), 3.90 – 4.00 (2H, m, H2 and H3), 4.98 (1H, dd, *J* 11.0, 3.0, H7).

$\delta_C$ (101 MHz, Methanol-*d*$_4$) 21.6 (C10), 22.6 (C13), 22.7 (C13), 23.8 (C10), 25.9 (C9), 41.7 (C12), 53.3 (C15), 53.9 (C7), 56.2 (C2), 61.7 (C3), 167.6 (C5), 174.9 (C11), 208.2 (C11).

HRMS (ESI): Calc. for $[M+H]^+$ $C_{14}H_{27}N_2O_5^+$ = 303.1914, Obs. = 303.1911.

**Methyl (*S*)-4-((*S*)-2-heptanamido-3-hydroxypropanamido)-2,2,6-trimethyl-3-oxoheptanoate (17)**

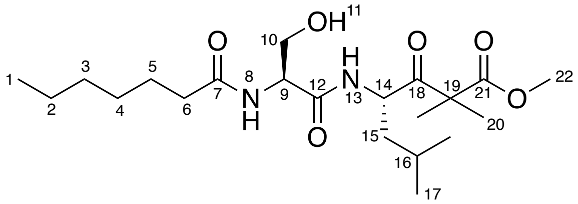

Prepared as per general procedure 6 - with heptanoic acid (0.035 g, 0.292 mmol, 1 equiv.), purified by silica gel chromatography (35:65 EtOAc:hexane) yielding the product as a clear oil (0.041 g, 0.989 mmol, 37%).

$\delta_H$ (400 MHz, Methanol-$d_4$) 0.76 – 1.02 (9H, m, H1 & H17), 1.22 – 1.33 (8H, m, H2, H4 & H5), 1.36 (3H, s, H20), 1.40 (3H, s, H20), 1.42 – 1.44 (1H, m, H15), 1.52 (1H, ddd, *J* 14.5, 11.0, 4.0, H15), 1.56 – 1.72 (3H, m, H6 & H16), 2.26 (2H, t, *J* 7.5, H6), 3.65 – 3.90 (5H, m, H10 & H22), 4.41 (1H, t, *J* 5.5, H9), 4.95 (1H, dd, *J* 11.0, 3.0, H14).

$\delta_C$ (101 MHz, Methanol-$d_4$) 14.4 (C1), 21.7 (C17), 22.5 (C20), 22.6 (C20), 23.6 (C2), 23.8 (C17), 25.7 (C16), 26.8 (C3), 30.0 (C4), 32.7 (C5), 36.9 (C6), 42.0 (C15), 53.2 (C22), 53.6 (C14), 56.1 (C19), 56.6 (C9), 62.8 (C10), 172.0 (C12), 175.0 (C22), 176.4 (C8), 208.5 (C18).

HRMS (ESI): Calc. for $[M+H]^+$ $C_{21}H_{39}N_2O_6^+$ = 415.2803, Obs. = 415.2803.

**Methyl (*S*)-4-((*S*)-2-acetamido-3-hydroxypropanamido)-2,2,6-trimethyl-3-oxoheptanoate (36)**

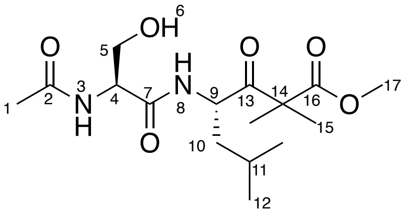

Procedure adapted from Andurkar *et al.* To a solution of methyl (*S*)-4-((*S*)-3-hydroxy-2-pivalamidopropanamido)-2,2,6-trimethyl-3-oxoheptanoate in dry THF (0.15 M, 2.0 mL) on ice was added successively pyridine (0.024 mL, 0.295 mmol, 1 equiv.), DMAP (0.004 g, 0.029 mmol, 0.1 equiv.), and $Ac_2O$ (0.027 mL, 0.295 mmol, 1 equiv.), under argon and the resulting solution was stirred and allowed to warm to ambient temperature for one hour. The reaction was concentrated *in vacuo* and the resulting residue was purified by silica gel chromatography (EtOAc) to yield the product as a white powder (0.048 g, 0.140 mmol, 47%).[13]

$\delta_H$ (400 MHz, Methanol-$d_4$) 0.89 – 0.94 (6H, m, H12), 1.36 (3H, s, H15), 1.40 (3H, s, H15), 1.43 (1H, d, *J* 3.5, H10), 1.52 (1H, ddd, *J* 14.5, 11.0, 4.0, H10), 1.66 (1H, dqd, *J* 13.5, 6.5, 3.0, H11), 2.01 (3H, s, H1), 3.56 – 3.93 (5H, m, H5 & H17), 4.40 (1H, t, *J* 5.5, H4), 4.95 (1H, dd, *J* 11.0, 3.5, H9).

$\delta_C$ (101 MHz, Methanol-$d_4$) 21.7 (H12), 22.5 (C1), 22.6 (C15), 22.7 (C15), 23.8 (C12), 25.7 (C11), 42.0 (C10), 53.2 (C17), 53.7 (C9), 56.1 (C14), 56.8 (C4), 62.9 (C5), 171.9 (C7), 173.5 (C2), 175.0 (C16), 208.5 (C13).

HRMS (ESI): Calc. for $[M+H]^+$ $C_{16}H_{29}N_2O_6^+$ = 345.2020, Obs. = 345.2015.

**Methyl (*S*)-4-((*S*)-3-hydroxy-2-pivalamidopropanamido)-2,2,6-trimethyl-3-oxoheptanoate (37)**

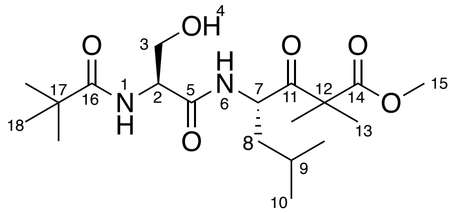

Procedure adapted from Annese *et al*. To a solution of methyl (*S*)-4-((*S*)-3-hydroxy-2-pivalamidopropanamido)-2,2,6-trimethyl-3-oxoheptanoate hydrochloride salt (0.044 g, 0.15 mmol, 1 equiv.) in anhydrous THF (2 mL) under argon at 0 °C, pivaloyl chloride (0.02 mL, 0.17 mmol, 1.1 equiv.) and $Et_3N$ (0.04 mL, 0.29 mmol, 2 equiv.) were added. The reaction was allowed to warm to ambient temperature overnight before the solvent was removed *in vacuo*. The resulting residue was partitioned between d.$H_2O$ (5 mL) and EtOAc (5 ml), the layers were separated and the aqueous layer extracted with EtOAc (2 x 5 mL). The combined aqueous layers were washed with 5% $NaHCO_3$, 1 M HCl and brine, dried over $MgSO_4$, filtered and concentrated *in vacuo*. The crude residue was purified by silica gel chromatography (3:2 EtOAc:hexane) to yield the desired product as a clear oil.[14] (0.023 g, 0.05 mmol, 46%)

$\delta_H$ (400 MHz, Methanol-$d_4$) 0.88 – 0.97 (6H, m, H10), 1.21 (9H, s, H18), 1.36 (3H, s, H13), 1.40 – 1.44 (4H, m, H8 and H13), 1.51 (1H, ddd, *J* 14.5, 11.0, 4.0, H8), 1.59 – 1.70 (1H, m, H9), 3.72 (3H, s, H15), 3.73 – 3.81 (2H, m, H3), 4.40 (1H, t, *J* 5.5, H2), 4.96 (1H, dd, *J* 11.0, 3.0, H7).

$\delta_C$ (101 MHz, Methanol-$d_4$) 21.7 (C10), 22.6 (C13), 22.6 (C13), 23.8 (C10), 25.7 (C9), 27.7 (C18), 39.8 (C17), 42.1 (C8), 53.2 (C15), 53.6 (C7), 56.6 (C7), 62.7 (C3), 172.0 (C5), 175.0 (C14), 181.3 (C16), 208.5 (C11).

HRMS (ESI): Calc. for $[M+H]^+$ $C_{19}H_{35}N_2O_6^+$ = 387.2490, Obs. = 387.2486.

**Methyl (*S*)-4-butyramido-2,2,6-trimethyl-3-oxoheptanoate (21)**

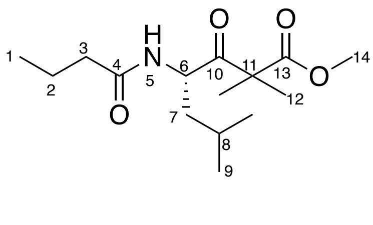

Prepared as per general procedure 6 - with butyric acid (0.032 g, 0.361 mmol, 1 equiv.), purified by silica gel chromatography (2:3 EtOAc:hexane) yielding the product as a yellow oil (0.077 g g, 0.270 mmol, 74%).

$\delta_H$ (400 MHz, Chloroform-*d*) 0.60 – 1.18 (9H, m, H1 & H9), 1.37 (2H, dd, *J* 5.0, 3.5, H7), 1.40 (3H, s, H12), 1.43 (3H, s, H12), 1.55 – 1.70 (3H, m, H2 & H8), 2.14 (1H, t, *J* 7.5, H3), 3.71 (3H, s, H14), 5.05 (1H, td, *J* 9.0, 5.0, H6), 5.73 (1H, d, *J* 9.5, H5).

$\delta_C$ (101 MHz, Chloroform-*d*) 13.8 (C1), 19.2 (C2), 21.5 (C9), 22.3 (C12), 22.4 (C12), 23.7 (C9), 25.0 (C8), 38.7 (C3), 42.1 (C7), 52.1 (C6), 52.8 (C14), 55.2 (C11), 172.2 (C4), 173.5 (C13), 208.9 (C10).

HRMS (ESI): Calc. for $[M+H]^+$ $C_{15}H_{28}NO_4^+$ = 286.2013, Obs. = 286.2012.

##### 1.2.6 Chemical Synthesis of $R^4$-functionalised α-dimethyl-β-keto-ester substrates

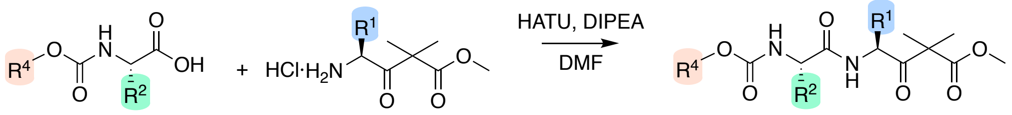

*Scheme S1.5: General synthetic scheme for the synthesis of $R^4$-functionalised dipeptidyl α-dimethyl-β-keto-esters*

**Methyl (*S*)-4-((*S*)-3-hydroxy-2-(2-phenylacetamido)propanamido)-2,2,6-trimethyl-3-oxoheptanoate (38)**

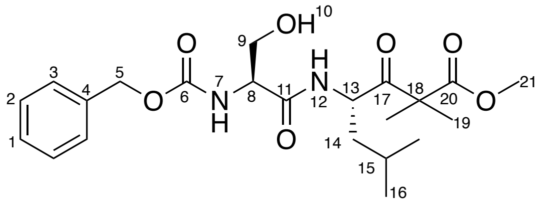

Prepared as per general procedure 6 – with ((benzyloxy)carbonyl)-L-serine (0.087 g, 0.36 mmol, 1 equiv.), purified by silica gel chromatography (98:2 $CH_2Cl_2$:MeOH) yielding the final product as a clear oil (0.034 g, 0.07 mmol, 21%).

$\delta_H$ (400 MHz, Chloroform-*d*) 0.83 (3H, d, *J* 6.5, H16), 0.86 (3H, d, *J* 6.5, H16), 1.35 (3H, H19), 1.36 – 1.39 (5H, m, H14 & H19), 1.52 (1H, sext., *J* 6.5, H15), 3.58 (1H, dd, *J* 6.0, H9), 3.67 (3H, s, H21), 3.96 (1H, d, *J* 11.5, H9), 4.16 (1H, br. s, H8), 4.91 (1H, td, *J* 9.0, 5.5, H13), 5.06 (1H, s, H5), 5.77 (1H, d, *J* 7.5, H7), 6.79 (1H, s, H12), 7.22 – 7.35 (5H, m, H1, H2 & H3).

$\delta_C$ (101 MHz, Chloroform-*d*) 21.4 (C16), 22.3 (C19), 22.5 (C19), 23.5 (C16), 25.0 (C16), 41.3 (C14), 52.9 (C21), 53.1 (C13), 55.1 (C18), 55.4 (C8), 63.0 (C9), 67.4 (C5), 128.2 (C3), 128.4 (C1), 128.7 (C2), 136.1 (C4), 156.6 (C6), 170.5 (C11), 173.6 (C21), 207.9 (C17).

HRMS (ESI): Calc. for $[M+Na]^+$ $C_{22}H_{32}N_2O_7Na^+$ 459.2102, Obs. = 459.2098.

**Methyl (*S*)-4-((*S*)-2-(((allyloxy)carbonyl)amino)-3-hydroxypropanamido)-2,2,6-trimethyl-3-oxoheptanoate (39)**

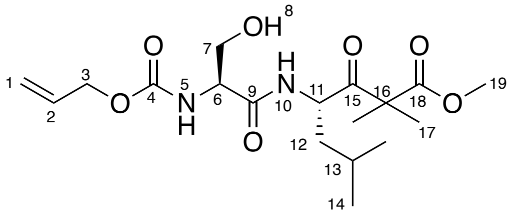

((allyloxy)carbonyl)-L-serine was prepared as per a procedure adapted from Yan *et al*. L-serine (1.0 g, 9.5 mmol, 1 equiv.) and $Na_2CO_3$ (1.0 g, 9.5 mmol, 1 equiv.) were dissolved in d.$H_2O$ (10 ml). Allyl chloroformate (1.0 ml, 9.5 mmol, 1 equiv.) was added dropwise and MeCN (5 mL) were added sequentially and stirred overnight. MeCN was removed *in vacuo,* and the solution was acidified to pH 2 by conc. HCl then extracted with EtOAc (3 x 10 mL), dried over $Na_2SO_4$, filtered and concentrated to yield the desired product as a clear oil. (1.01 g, 5.3 mmol, 56%).

Prepared as per general procedure 6 – with ((allyloxy)carbonyl)-L-serine (0.07 g, 0.37 mmol, 1 equiv.), purified by silica gel chromatography (EtOAc) yielding the final product as a yellow oil (0.083 g, 0.21 mmol, 58%).

$\delta_H$ (400 MHz, Methanol-*d*$_4$) 0.79 – 0.97 (6H, m, H14), 1.36 (3H, s, H17), 1.39 – 1.44 (4H, m, H12 & H17), 1.53 (1H, ddd, *J* 14.5, 11.0, 4.0, H12), 1.65 (1H, dqd, *J* 10.5, 6.5, 4.0, H13), 3.61 – 3.95 (5H, m, H7 & H19), 4.18 (1H, t, *J* 5.5, H6), 4.55 (2H, d, *J* 4.5, H3), 4.96 (1H, dd, *J* 11.0, 3.0, H11), 5.19 (1H, d, *J* 10.5, H1), 5.31 (1H, d, *J* 17.5, H1), 5.94 (1H, ddt, *J* 16.5, 10.5, 5.5, H2).

$\delta_C$ (101 MHz, Methanol-*d*$_4$) 21.7 (C14), 22.6 (C17), 22.6 (C17), 23.8 (C14), 25.7 (C13), 41.9 (C12), 53.2 (C19), 53.6 (C11), 56.2 (C16), 58.4 (C6), 63.1 (C7), 66.7 (C3), 117.7 (C1), 134.2 (C2), 158.2 (C4), 172.3 (C9), 174.9 (C18), 208.5 (C15).

HRMS (ESI): Calc. for $[M+H]^+$ $C_{18}H_{31}N_2O_7^+$ = 387.2126, Obs. = 387.2129.

**Methyl (*S*)-4-((*S*)-2-((((9H-fluoren-9-yl)methoxy)carbonyl)amino)-3-(tert-butoxy)propanamido)-2,2,6-trimethyl-3-oxoheptanoate**

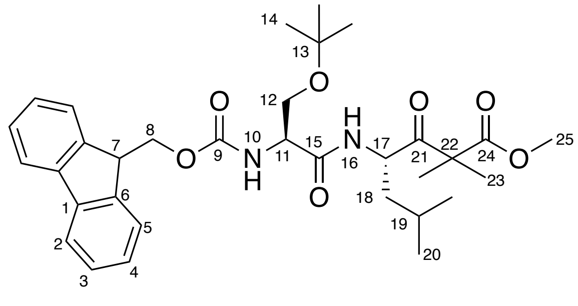

Prepared as per general procedure 6 – with *N*-(((9*H*-fluoren-9-yl)methoxy)carbonyl)-*O*-(*tert*-butyl)-L-serine (0.139 g, 0.361 mmol, 1 equiv.), purified by silica gel chromatography (1:1 EtOAc:hexane) yielding the final product as a white powder (0.198 g, 0.341 mmol, 94%).

$\delta_H$ (400 MHz, Chloroform-*d*) 0.91 (3H, d, *J* 6.5, H20), 0.95 (3H, d, *J* 6.5, H20), 1.21 (9H, s, H14), 1.35 – 1.43 (4H, m, H18 & H23), 1.44 (3H, s, H23), 1.59 – 1.72 (2H, m, H18 & H19), 3.32 (1H, t, *J* 8.5, H12), 3.73 (3H, s, H25), 3.81 (1H, dd, *J* 8.5, 3.5, H12), 4.09 – 4.30 (2H, m, H7 & H11), 4.35 – 4.49 (2H, m, H8), 5.02 (1H, td, *J* 10.0, 3.5, H17), 5.71 (1H, d, *J* 6.5, H16), 7.10 (1H, d, *J* 9.5, H10), 7.32 (2H, t, *J* 7.5, H4), 7.40 (2H, t, *J* 7.5, H3), 7.60 (2H, dd, *J* 7.5, 4.0, H5), 7.77 (2H, d, *J* 7.5, H2).

$\delta_C$ (101 MHz, Chloroform-*d*) 21.5 (C20), 22.3 (C23), 23.7 (C20), 24.8 (C19), 27.4 (C14), 42.1 (C18), 47.3 (C7), 52.4 (C17), 52.8 (C25), 54.4 (C11), 55.2 (C22) 61.8 (C12), 67.3 (C8), 120.1 (C5), 125.2 (C4), 127.2 (C3), 128.4 (C2), 141.4 (C1), 144.0 (C6), 156.0 (C9), 169.7 (C15), 173.5 (C24), 207.6 (C21).

HRMS (ESI): Calc. for $[M+Na]^+$ $C_{33}H_{44}N_2O_7Na^+$ = 603.3041, Obs. = 603.3039.

**Methyl (*S*)-4-((*S*)-2-((((9H-fluoren-9-yl)methoxy)carbonyl)amino)-3-hydroxypropanamido)-2,2,6-trimethyl-3-oxoheptanoate (40)**

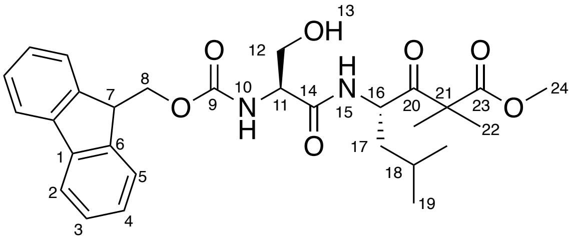

Procedure adapted from Zhang *et al.*[15] To a solution of methyl (*S*)-4-((*S*)-2-((((9H-fluoren-9-yl)methoxy)carbonyl)amino)-3-(tert-butoxy)propanamido)-2,2,6-trimethyl-3-oxoheptanoate (0.1 g, 0.18 mmol, 1 equiv.) in TFA/$CH_2Cl_2$/$H_2O$ (4:1:1, 5 mL) was stirred at rt for 3 h. The solvent was removed *in vacuo* using a sodium hydrogen carbonate trap, and remaining volatiles were co-evaporated with toluene. The resulting residue was purified by silica gel chromatography (1:1 EtOAc:hexane) yielding the final product as a white waxy solid (0.042 g, 0.080 mmol, 42%).

$\delta_H$ (400 MHz, Methanol-*d*$_4$) 0.80 – 1.02 (6H, m, H19), 1.36 (3H, s, H22), 1.37 – 1.44 (4H, m, H17 & H22), 1.51 (1H, ddd, *J* 14.5, 11.0, 4.0, H17), 1.65 (1H, dtd, *J* 14.0, 6.5, 3.5, H18), 3.65 – 3.79 (5H, m, H12 & H24), 4.18 – 4.24 (2H, m, H7 & H11), 4.33 (1H, dd, *J* 10.5, 7.0, H8), 4.40 (1H, dd, *J* 10.5, 7.0, H8), 4.96 (1H, dd, *J* 11.0, 3.5, H16), 7.30 (2H, t, *J* 7.5, H4), 7.38 (2H, t, *J* 7.5, H3), 7.66 (2H, t, *J* 7.0, H5), 7.78 (2H, d, *J* 7.5, H2).

$\delta_C$ (101 MHz, Methanol-*d*$_4$) 21.7 (C19), 22.6 (C22), 22.7 (C22), 23.8 (C19), 25.7 (C18), 42.0 (C17), 48.3 (C7), 53.2 (C24), 53.7 (C16), 56.1 (C21), 58.4 (C11), 63.1 (C12), 68.1 (C8), 120.9

(C5), 126.2 (C4), 126.2 (C3), 128.2 (C2), 142.5 (C1), 145.1 (C6), 158.4 (C9), 172.3 (C14), 174.9 (C23), 208.5 (C20).
HRMS (ESI): Calc. for $[M+H]^+$ $C_{29}H_{37}N_2O_7^+$ = 525.2595, Obs. = 525.2592.

**Methyl (*S*)-4-((*S*)-2-((tert-butoxycarbonyl)amino)-3-methoxypropanamido)-2,2,6-trimethyl-3-oxoheptanoate (41)**

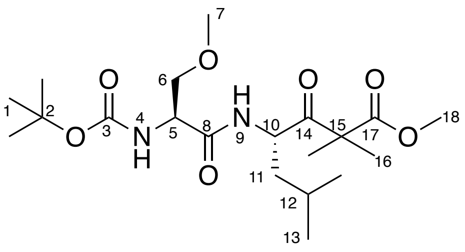

Prepared as per general procedure 6 – with *N*-(*tert*-butoxycarbonyl)-*O*-methyl-L-serine (0.05 g, 0.23 mmol, 1 equiv.), purified by silica gel chromatography (7:3 hexane:EtOAc) yielding the final product as a clear oil (0.076 g, 0.18 mmol, 80%).
$\delta_H$ (400 MHz, Methanol-*d*$_4$) 0.87 – 0.98 (6H, m, H13), 1.35 (1H, s, H16), 1.37 – 1.42 (4H, m, H11 & H16), 1.44 (9H, s, H1), 1.53 (1H, ddd, *J* 14.5, 11.0, 3.5, H11), 1.61 – 1.73 (1H, m, H12), 3.32 (3H, s, H7), 3.47 – 3.64 (2H, m, H6), 3.71 (3H, s, H18), 4.19 (1H, t, *J* 5.0, H6), 4.96 (1H, dd, *J* 11.0, 3.0, H10).
$\delta_C$ (101 MHz, Methanol-*d*$_4$) 21.7 (C13), 22.5 (C16), 22.5 (C16), 23.9 (C13), 25.6 (C12), 28.6 (C1), 42.1 (C11), 53.1 (C18), 53.5 (C10), 56.1 (C5), 59.2 (C7), 73.1 (C6), 82.4 (C2), 158.0 (C3), 172.4 (C8), 174.9 (C17), 208.4 (C14).
HRMS (ESI): Calc. for $[M+H]^+$ $C_{20}H_{37}N_2O_7^+$ = 417.2595, Obs. = 417.2592.

**Methyl (*S*)-4-((*S*)-2-((tert-butoxycarbonyl)amino)-3-hydroxypropanamido)-2,2-dimethyl-3-oxo-5-phenylpentanoate (42)**

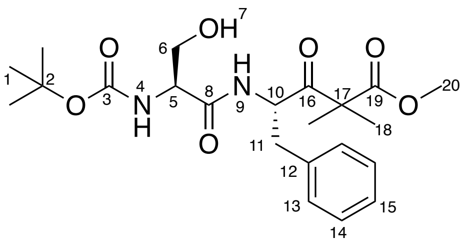

Prepared as per general procedure 6 – with (*tert*-butoxycarbonyl)-L-serine (0.10 g, 0.32 mmol, 1 equiv.), purified by silica gel chromatography (1:1 EtOAc:hexane) yielding the final product as a white powder (0.059 g, 0.14 mmol, 43%).
$\delta_H$ (400 MHz, Chloroform-*d*) 1.32 (3H, s, H18), 1.38 (3H, s, H18), 1.46 (9H, s, H1), 2.77 (1H, dd, *J* 14.0, 9.0, H11), 3.15 (1H, dd, *J* 14.0, 5.5, H11), 3.52 (1H, dd, *J* 11.5, 5.0, H6), 3.70 (3H,

s, H20), 3.91 (1H, d, *J* 11.5, H6), 4.03 (1H, s, H5), 5.20 (1H, td, *J* 9.0, 5.5, H10), 5.34 (1H, d, *J* 7.5, H4), 6.92 (1H, d, *J* 8.5, H9), 7.16 (2H, d, *J* 6.5, H13), 7.18 – 7.32 (3H, m, H14 & H15).
$\delta_C$ (101 MHz, Chloroform-*d*) 21.7 (C18), 22.3 (C18), 28.4 (C1), 38.4 (C6), 52.9 (C20), 55.0 (C10), 55.1 (C5), 62.8 (C6), 80.7 (C2), 127.2 (C13), 128.6 (C15), 129.5 (C14), 136.3 (C12), 156.1 (C3), 170.6 (C8), 173.6 (C19), 206.7 (C16).
HRMS (ESI): Calc. for $[M+H]^+$ $C_{22}H_{33}N_2O_7^+$ = 437.2282, Obs. = 437.2268.

**Methyl (*S*)-4-((*S*)-2-((tert-butoxycarbonyl)amino)-3-methoxypropanamido)-2,2-dimethyl-3-oxo-5-phenylpentanoate (43)**

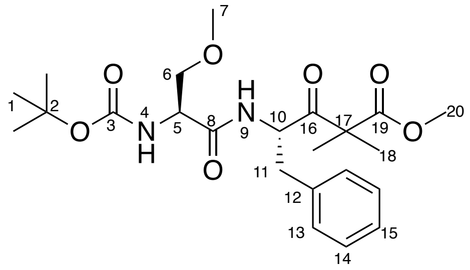

Prepared as per general procedure 6 – with *N*-(*tert*-butoxycarbonyl)-*O*-methyl-L-serine (0.20 g, 0.91 mmol, 1 equiv.), purified by silica gel chromatography (1:1 EtOAc:hexane) yielding the final product as a white powder (0.277 g, 0.61 mmol, 67%).
$\delta_H$ (400 MHz, Chloroform-*d*) 1.31 (3H, s, H18), 1.35 (3H, s, H18), 1.44 (9H, s, H1), 2.83 (1H, dd, *J* 14.0, 8.0, H11), 3.08 (1H, dd, *J* 14.0, 6.0, H11), 3.29 (3H, s, H7), 3.37 (1H, dd, *J* 9.5, 6.5, H6), 3.59 – 3.65 (1H, m, H6), 3.66 (3H, s, H20), 4.05 – 4.15 (1H, m, H5), 5.12 – 5.27 (1H, m, H10), 6.80 (1H, d, *J* 9.5, H9), 7.07 – 7.19 (2H, m, H13), 7.17 – 7.35 (3H, m, H14 & H15).
$\delta_C$ (101 MHz, Chloroform-*d*) 21.9 (C18), 22.1 (C18), 28.4 (C1), 38.4 (C11), 52.8 (C20), 54.8 (C5), 55.2 (C10), 59.0 (C7), 71.7 (C6), 80.5 (C2), 127.6 (C13), 128.6 (C15), 129.6 (C14), 136.2 (C12), 155.5 (C3), 169.7 (C8), 173.4 (C19), 206.9 (C16).
HRMS (ESI): Calc. for $[M+H]^+$ $C_{23}H_{35}N_2O_7^+$ = 451.2439, Obs. = 451.2438.

**Methyl (*S*)-4-((*S*)-2-((tert-butoxycarbonyl)amino)-3-phenylpropanamido)-2,2,6-trimethyl-3-oxoheptanoate (44)**

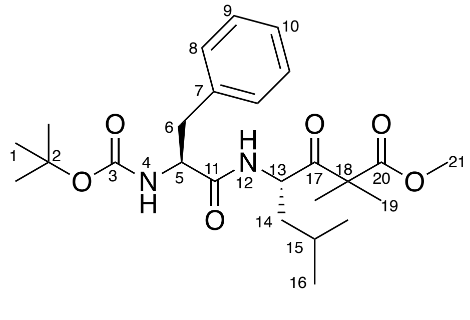

Prepared as per general procedure 6 – with (*tert*-butoxycarbonyl)-L-phenylalanine (0.30 g, 1.1 mmol, 1 equiv.), purified by silica gel chromatography (1:1 EtOAc:hexane) yielding the final product as a white powder (0.425 g, 0.092 mmol, 81%).

$\delta_H$ (400 MHz, Chloroform-*d*) 0.82 (H, d, *J* 6.5, H16), 0.89 (3H, d, *J* 6.5, H16), 1.21 – 1.35 (8H, m, H14 & H10), 1.37 (9H, s, H1), 3.00 (2H, d, *J* 7.0, H6), 3.67 (3H, s, H21), 4.25 (1H, q, *J* 7.0, H5), 4.92 (2H, q, *J* 8.0, H4 & H13), 6.23 (1H, d, *J* 9.0, H12), 7.09 – 7.25 (5H, m, H8, H9 & H10).

$\delta_C$ (101 MHz, Chloroform-*d*) 21.4 (C16), 22.1 (C19), 22.3 (C19), 23.6 (C16), 24.6 (C15), 28.3 (C1), 37.8 (C6), 42.0 (C14), 52.2 (C13), 52.7 (C21), 55.1 (C18), 55.8 (C5), 80.5 (C2), 127.1 (C10), 128.8 (C8), 129.5 (C9), 136.5 (C7), 155.5 (C3), 170.5 (C11), 173.5 (C20), 207.7 (C17).

HRMS (ESI): Calc. $[M+Na]^+$ $C_{25}H_{38}N_2O_6Na^+$ = 485.2622, Obs. = 485.2617.

**methyl (*S*)-4-((*S*)-2-(((allyloxy)carbonyl)amino)-3-phenylpropanamido)-2,2,6-trimethyl-3-oxoheptanoate (45)**

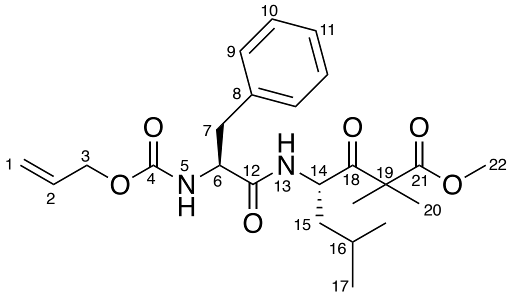

((allyloxy)carbonyl)-L-phenylalanine was prepared according to a procedure adapted from Yan *et al.* L-phenylalanine (1.57 g, 9.5 mmol, 1 equiv.) and $Na_2CO_3$ (1.0 g, 9.5 mmol, 1 equiv.) were dissolved in d.$H_2O$ (10 ml). Allyl chloroformate (1.0 ml, 9.5 mmol, 1 equiv.) was added dropwise and MeCN (5 mL) were added sequentially and stirred overnight. MeCN was removed *in vacuo,* and the solution was acidified to pH 2 by conc. HCl then extracted with EtOAc (3 x 10 mL), dried over $Na_2SO_4$, filtered and concentrated to yield the desired product as a clear oil. (1.92 g, 4.9 mmol, 52%) which was subsequently used without further purification. Following general procedure 6 – with ((allyloxy)carbonyl)-L-phenylalanine (0.30 g, 1.2 mmol, 1 equiv.), purified by silica gel chromatography (1:1 $Et_2O$:hexane) yielding the final product as a white powder (0.131 g, 0.29 mmol, 24%).

$\delta_H$ (400 MHz, Methanol-$d_4$) 0.88 – 0.95 (6H, m, H17), 1.34 (3H, s, H20), 1.36 (3H, s, H20), 1.38 – 1.42 (1H, m, H15), 1.52 (1H, td, *J* 11.0, 5.5, H15), 1.64 (1H, dqd, *J* 13.0, 6.5, 3.5, H16), 2.81 (1H, dd, *J* 14.0, 9.5, H7), 3.07 (1H, dd, *J* 14.0, 5.5, H7), 3.71 (3H, s, H22), 4.37 (1H, dd, *J* 9.5, 5.5, H6), 4.42 – 4.51 (2H, m, H3), 4.91 – 4.99 (1H, m, H14), 5.13 (1H, d, *J* 10.5, H1), 5.21 (1H, d, *J* 16.5, H1), 5.85 (1H, ddt, *J* 16.0, 10.5, 5.5, H2), 7.16 – 7.31 (5H, m, H9, H10 & H11).

δ$_C$ (101 MHz, Methanol-$d_4$) 21.6 (C17), 22.6 (C20), 22.7 (C20), 23.8 (C17), 25.7 (C16), 38.9 (C7), 41.9 (C15), 53.1 (C22), 53.5 (C14), 56.2 (C19), 57.5 (C6), 66.5 (C3), 117.5 (C1), 127.7 (C9), 129.4 (C10), 130.3 (C11), 134.2 (C2), 138.4 (C8), 158.0 (C4), 173.5 (C12), 174.9 (C21), 208.4 (C18).
HRMS (ESI): Calc. for $[M+H]^+$ $C_{24}H_{35}N_2O_6^+$ = 447.2490, Obs. = 447.2490.

##### 1.2.7 Synthesis of α,β-unsaturated ketone authentic standard

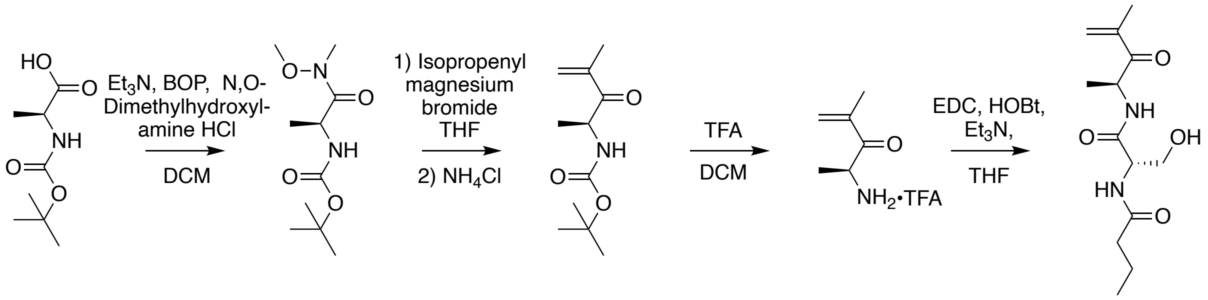

*Scheme S1.6: General synthetic scheme for the synthesis of α,β-unsaturated ketone authentic standard*

***tert*-Butyl (*S*)-(1-(methoxy(methyl)amino)-1-oxopropan-2-yl)carbamate**

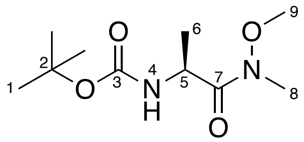

To a stirred solution of *N*-BOC-L-alanine (402 mg, 2.1 mmol) in DCM (9.6 mL) was added O,N-dimethylhydroxylamine hydrochloride (227 mg, 2.3 mmol), triethylamine (620 µL, 4.4 mmol) and BOP reagent (937 mg, 2.1 mmol). Solvent was removed in *vacuo*, and the resulting residue purified by silica gel chromatography (3:2 ethyl acetate:*n*-hexane) to give the desired product **26** as a white solid (266 mg, 1.1 mmol, 54%).[16]
δ$_H$ (500 MHz, Chloroform-*d*) 5.25 (1H, br. d, *J* 7.0, H4), 4.68 (1H, m, H5), 3.77 (3H, s, H9), 3.21 (3H, s, H8), 1.44 (9H, s, H1), 1.31 (3H, d, *J* 7.0, H6).
δ$_C$ (125 MHz, Chloroform-*d*) 173.9 (C7), 155.4 (C3), 78.0 (C2), 61.8 (C9), 46.7 (C5), 32.3 (C8), 28.5 (C1), 18.9 (C6).
LRMS (ESI): Calc. for $[M+Na]^+$ $C_{10}H_{20}N_2O_4Na^+$ = 255.1, Obs. = 255.1.

***tert*-Butyl *(S)*-(4-methyl-3-oxopent-4-en-2-yl)carbamate**

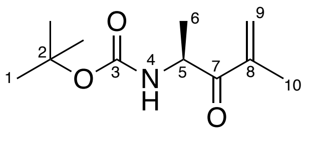

Under an atmosphere of argon to a stirred solution of **26** (200 mg, 863 µmol) in THF (3.5 mL) at 0 °C was added isopropenylmagnesium bromide in THF (500 mM, 3.6 mL, 1.8 mmol)

dropwise. The reaction was stirred at 0 °C for 6 hours, then poured into saturated ammonium chloride solution (5 mL) and ice (ca. 5 mL). When ice had melted, pH was adjusted to 7 with HCl (6 M). Solution was extracted with ethyl acetate (3 × 30 mL). Organic extracts were combined and washed with saturated $NaHCO_3$ (2 × 10 mL), brine (2 × 10 mL) and dried ($MgSO_4$). Solvent was removed in *vacuo* to yield the desired product **27** as a pale yellow oil (113 mg, 530 µmol, 61%).[17]

$\delta_H$ (500 MHz, Chloroform-*d*) 6.04 (1H, s, H9), 5.89 (1H, s, H9), 5.38 (1H, br. d, *J* 5.5, H4), 5.07-4.99 (1H, m, H5), 1.91 (3H, s, H10), 1.44 (9H, s, H1), 1.31 (3H, d, *J* 7.0, H6).

$\delta_C$ (125 MHz, Chloroform-*d*) 201.2 (C7), 155.2 (C3), 141.9 (C8), 126.4 (C9), 79.8 (C2), 50.3 (C5), 28.5 (C1), 20.4 (C6), 18.0 (C10).

LRMS (ESI): Calc. for $[M+H]^+$ $C_{11}H_{19}NO_3Na^+$ = 236.1, Obs. = 236.1.

##### *(S)*-4-methyl-3-oxopent-4-en-2-ammonium trifluoroacetate

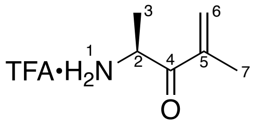

To a stirred solution of **27** (97.3 mg, 456 µmol) in DCM (8 mL) was added TFA (2 mL) dropwise. After half an hour the solution was diluted with DCM (3 × 20 mL), removing solvent in *vacuo* successively. The resulting crude oil was washed successively with petroleum ether (3 × 5 mL), and residual solvent removed in *vacuo* from the immiscible oil to give the desired product **28** as an orange oil (92.7 mg, 408 µmol, 89%).

$\delta_H$ (500 MHz, Methanol-*d*$_4$) 6.17 (1H, s, H6), 6.13 (1H, br. d, *J* 1.0, H6), 4.83 (1H, q, *J* 7.5, H2), 1.93 (3H, s, H7), 1.49, (3H, d, *J* 7.0, H3).

$\delta_C$ (125 MHz, Methanol-*d*$_4$) 198.5 (C4), 142.3 (C5), 129.2 (C6), 52.0 (C2), 18.0 (C3), 17.6 (C7).

HRMS (ESI): Calc. for $[M+H]^+$ $C_6H_{12}NO^+$ = 114.0913, Obs. = 114.0913.

##### *N*-*((S)*-3-hydroxy-1-*(((S)*-4-methyl-3-oxopent-4-en-2-yl)amino)-1-oxopropan-2-yl)butyramide (46)

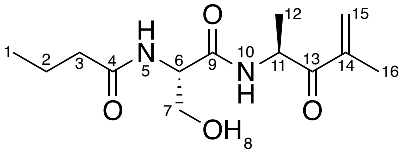

Under an atmosphere of argon to a stirred solution of HOBt (72.6 mg, 537 µmol) in THF (4.8 mL) at 0 °C was added *N*-butanoyl-L-serine **29** (39.0 mg, 223 µmol) in THF (390 µL), **28** (50.4 mg, 222 µmol) in THF (500 µL), triethylamine (130 µL, 933 µmol) and EDC (85.2 mg, 444 µmol). The reaction was warmed to room temperature and stirred for 24 hours. Reaction mixture was filtered, and solvent removed in *vacuo*. The resulting residue was purified by silica gel chromatography (gradient of 5% methanol in ethyl acetate to 10% methanol in ethyl acetate) to yield the desired product **24** as a pale yellow solid (28.4 mg, 105 µmol, 47%).

$\delta_H$ (500 MHz, Methanol-$d_4$) 6.12 (1H, s, H15), 5.94 (1H, br d, *J* 1.5, H15), 5.21 (1H, q, *J* 7., H11), 4.45 (1H, t, *J* 5.5, H6), 3.76 (2H, d, *J* 5.5, H7), 2.26 (2H, dt, *J* 7.5, 1.5, H3), 1.88 (3H, s, H16), 1.66 (2H, sext, *J* 7.5, H2), 1.32 (3H, d, *J* 7.0, H12), 0.96 (3H, t, *J* 7.5, H1).

$\delta_C$ (125 MHz, Methanol-$d_4$) 201.9 (C13), 176.3 (C4), 171.9 (C9), 143.3 (C14), 126.9 (C15), 63.1 (C7), 56.7 (C6), 50.7 (C11), 38.7 (C3), 20.2 (C2), 18.7 (C12), 18.0 (C16), 14.0 (C1).

HRMS (ESI): Calc. for $[M+Na]^+$ $C_{13}H_{22}N_2O_4Na^+$ = 293.1472, Obs. = 293.1467.

#### 1.3 Biochemical procedures

##### 1.3.1 Materials and instrumentation

All medium, buffer and assay components were purchased from Sigma Aldrich, Thermo Scientific, Difco, and Becton Dickinson Microbiology unless otherwise stated. Polymerases, DNA loading dye and markers, Sybr safe, KLD enzyme mix were purchased from New England Biolabs or Thermo Scientific unless otherwise stated. Monarch® Spin Plasmid Miniprep Kit and Monarch® Spin DNA gel extraction kit were purchased from New England Biolabs.

Media was autoclaved at 121 °C and 1 bar for 20 minutes. Minisart syringe filters (Sartorius) were used for sterile filtering. Bacterial cultures were incubated in New Brunswick Scientific Innova shaking incubators and or in Thermo Scientific Heraeus static incubators. PCR reactions were carried out on an Eppendorf Mastercycler nexus gradient thermocycler. SDS-PAGE and agarose gels were run using a Bio-Rad PowerPac Basic connected to Bio-Rad Mini-PROTEAN Tetra System or Bio-Rad SubCell Gel tanks respectively. $OD_{600}$ measurements were taken using a WPA Biowave CO8000 Cell Density Meter. Large scale cell cultures were pelleted using a Hitachi CR22N High-Speed Refrigerated Centrifuge. Cell lysis was conducted using a Constant Systems Ltd TS-Series Cabinet cell disruptor (One-Shot mode, 20 KPSI).

Samples from enzyme assays were analysed on a C18 column using either the Bruker Compact mass spectrometer (HR-MS) or a Bruker amazon speed ETD mass spectrometer (LR-MS) and a gradient from 5 100% acetonitrile in water each supplemented with 0.1% formic acid at a flow rate of 0.2 mL/min.

##### 1.3.2 Bacterial strains and plasmids

| **Strain** | **Source** |
|---|---|
| *E. coli* BL21 DE3* | Invitrogen |
| *E. coli* NEB5α | New England Biolabs |

| **Plasmid** | **Resistance** | **Source** |
|---|---|---|
| pET24a-His8_G2K_EpnF | Kan | Epoch Life Sciences |

**His8_G2K_EpnF**
<u>MKHHHHHHHHGGLVPRGSHGS</u>VSDSKSVNLFHRHGVPSFLEGIYQGRFEWDMISNFVAQ
DSADEKAGDAAVERLTDILRNRVNPTAVDATRELPEGLLEELRRTGFLNLQDSPDIGGLGLS
SYNTFRVVQAAASWSVPVALVLGIQTAVGSGTYLRALPPGELHSYVEQRLLDGIISGSADTE
PAGASNSARRTRAVPTDDGEAFLLTGEKIHIGNGPIADIVTVSAMLDEDGQDRPRLFFVETS
DPGFSIRSRHEFMGVKGFPNAALVLDGVRVPRERMIVEVDPDTEVRITAELTMAVVRGRLHL
ITAPSLAISKLCLEWSRNFVNRRTIDGRPLGAREEIQHMVSSTMADVFAIQALAEWSLLPADQ
PDLGLNVAFEQNVTKAISAEICWRAADRTMDLLAGEGFETAPSKARRGAPALPLERFYRDA

RNLRISGGVSFLLQFWAARMSQFTYYGPDHAGQSAALSQDGGQQPCTDGLDPLNADHLR
FAAAETRRLGAACQKFAADHPAPGLYEHQHRLIAFSRIADEILTMSVVLAKAARLHHEGRTEA
QDLAAIYCAHARDRLAALWRQAEPVTAGPDHAAVSDAWLSGDDTYASLITGVITDVPPTADT
HPGKR

##### 1.3.3 PCR primers

**pET24a-His8_G2K_EpnF_S158A**
*Site directed mutagenesis*
Forward: CATTTCTGGCGCAGCTGACACCG
Reverse: ATACCATCCAGCAGACGTT
Template – pET24a-His8_G2K_EpnF

**pET24a-His8_G2K_EpnF_R283K**
*Site directed mutagenesis*
Forward: TGTTCGTGGTAAACTGCACCTGATC
Reverse: ACTGCCATAGTCAGTTCAG
Template - pET24a-His8_G2K_EpnF

**pET24a-His8_G2K_EpnF_R283A**
*Site directed mutagenesis*
Forward: TGTTCGTGGTGCACTGCACCTGATC
Reverse: ACTGCCATAGTCAGTTCAG
Template - pET24a-His8_G2K_EpnF

**pET24a-His8_G2K_EpnF_S416A**
*Site directed mutagenesis*
Forward: TCTGCGCATCGCAGGTGGCGTGT
Reverse: TTGCGTGCATCACGGTAG
Template - pET24a-His8_G2K_EpnF

##### 1.3.4 Site-directed mutagenesis

All EpnF mutants were produced using Q5 site-directed mutagenesis (NEB), with the wild-type construct as a template and the primers detailed in Section S1.3.3. Gel purified DNA resulting from site directed mutagenesis PCR reactions was circularised using KLD enzyme mix (NEB) in the following reaction: DNA (1 µL), 2x KLD reaction buffer (5 µL), 10x KLD enzyme master mix (0.5 µL) and d.$H_2O$ (3.5 µL). The reaction mixture was incubated at ambient temperature for 5 mins, prior to chemical transformation with chemically competent *E. coli* cells and resulting plasmids sequenced (Eurofins Genomics) to verify the success of mutagenesis.

##### 1.3.5 Protein overproduction and purification

Chemically competent BL21 (DE3) cells were transformed by heat-shock with the relevant plasmid and grown overnight on LB agar containing the appropriate antibiotic. 10 mL of LB was then inoculated with a single colony of *E. coli* and incubated overnight (37 °C, 180 rpm). 3 L flasks containing 1 L LB, supplemented with the appropriate antibiotic, were inoculated with the seed cultures and incubated (37 °C, 180 rpm) until reaching an $OD_{600}$ of ~0.8. The cultures were cooled to 15 °C and IPTG (100 µM for epoxyketone synthases and 250 µM for other proteins) and riboflavin (only for epoxyketone synthases, 500 µM) were added and the cultures incubated overnight (15 °C, 180 rpm). Cells were harvested by centrifugation (4000 rpm, 40 minutes, 4 °C) and resuspended in wash buffer (20 mL/L of culture). The resulting cell suspensions were lysed by high pressure cell disruption (Constant Systems) and centrifuged (22000 rpm, 40 minutes, 4 °C) and the supernatant filtered (0.45 µm). Filtered lysate was loaded onto a 1 mL HisTrap™ Nickel FastFlow column and wash buffer (15 mL) was passed through the column. Proteins were then eluted from the column with elution buffers (5 mL, 50 mM), (3 mL, 100 mM, 200 mM, 300 mM) and 5 mL (500 mM). Elution fractions were analysed by SDS-PAGE and fractions containing the desired protein were concentrated to a volume of <2.5 mL using a Vivaspin centrifugal concentrator (10 kDa, 30 kDa, or 50 kDa cutoffs) and buffer-exchanged into storage buffer using a PD-10 desalting column (Cytiva) as per the manufacturer's procedure. The resulting elution was transferred into a fresh Vivaspin centrifugal concentrator and washed once more with storage buffer (5 mL) then concentrated for storage at -80 °C after flash freezing with liquid $N_2$.

Buffers

**Nickel (II) affinity washing buffer:** 100 mM NaCl, 20 mM Tris-HCl, 20 mM imidazole, 10% glycerol, pH 8.0.

**Nickel (II) affinity elution buffer:** 100 mM NaCl, 20 mM Tris-HCl, 50-, 100-, 200-, 300- and 500-mM imidazole, 10% glycerol, pH 8.0.

**Storage buffer**: 100 mM NaCl, 20 mM Tris-HCl, 10% glycerol, pH 8.0.

###### 1.3.6 UHPLC-ESI-Q-TOF-MS Analysis of Intact Proteins

All intact protein mass spectrometry analyses were conducted on a Bruker MaXis II ESI-Q-TOF-MS connected to a Dionex 3000 RS UHPLC fitted with an ACE C4-300 RP column (100 x 2.1 mm, 5 µm, 30 °C). The column was eluted with a linear gradient of 5 - 100% MeCN containing 0.1% formic acid over 30 min. The mass spectrometer was operated in positive ion mode with a scan range of 200 - 3000 m/z. Source conditions were: end plate offset at -500 V; capillary at -4500 V; nebulizer gas (N2) at 1.8 bar; dry gas (N2) at 9.0 L min−1 ; dry temperature at 200 °C. Ion transfer conditions were: ion funnel RF at 400 Vpp; multiple RF at 200 Vpp; quadrupole low mass at 200 m/z; collision energy at 8.0 eV; collision RF at 2000 Vpp; transfer time at 110.0 μs; pre-pulse storage time at 10.0 μs.

##### 1.3.7 Coupled PLE/EpnF assays

Coupled PLE/EpnF assays were prepared with a reaction volume of 100 μL in storage buffer and incubated for 3 hours at 30 °C. A typical reaction contained 1 mM substrate, 5 μM PLE, 50 μM epoxyketone synthase, and 2 mM FAD, with a negative control set up with the same composition without epoxyketone synthase. Assay products were analysed using either the Bruker Compact mass spectrometer (HR-MS) or a Bruker amazon speed ETD mass spectrometer (LR-MS).

For LR-MS analysis, the assays were quenched by addition of 400 μL acetonitrile + 0.1% formic acid then centrifuged (13,000 rpm, 10 minutes, 4 °C), the supernatant filtered (0.2 μm) and analysed in positive mode LC-MS.

For HR-MS analysis the assays were quenched with an equal volume of HPLC grade $H_2O$ containing formic acid (0.1%) and centrifuged (13,000 rpm, 10 minutes, 4 °C). In parallel, a C18 10 μL ZipTip® was washed with acetonitrile (3x) and equilibrated in HPLC grade $H_2O$ (5x). Molecules were then adsorbed from the assay supernatant onto the ZipTip® by aspirating (10x) and washed with HPLC grade $H_2O$ (0.1% formic acid) by aspiration (5x) before elution into 75 μL of acetonitrile:water (1:1) for LC-MS analysis. Care was taken not to introduce bubbles to the C18 matrix at any time.

EICs from LR-LC-MS analysis were prepared with a ±0.15 m/z tolerance, EICs from HR-LC-MS analysis were prepared with a ±0.01 m/z tolerance.

##### 1.3.8 CD analysis

Far-UV circular dichroism spectra were recorded on a JASCO J-1500 spectropolarimeter using a 0.01 mm pathlength quartz cuvette over 260–180 nm, with a 0.2 nm data pitch and 9 accumulations per spectrum. Protein spectra were baseline-corrected by subtraction of corresponding buffer blanks acquired between acquisitions of protein samples under identical conditions. Dictionary of secondary structure in proteins (DSSP) analysis was performed using the inbuilt CDSSTR method and reference set 4.[18–20]

#### 1.4 In silico methods

##### 1.4.1 Phylogenetic analysis

Multiple sequence alignments and phylogenetic analysis of biosynthetic protein sequences were generated using Multiple Sequence Comparison by Log- Expectation (MUSCLE) by performing an alignment with default parameters.[21]

For phylogenetic analysis of ACADs, ACOXs and epoxyketone synthases, RaxML_NG was used to perform a heuristic search of 20 maximum-likilihood trees to identify the best-scoring topology. The tree-drawing used a fixed empirical substitution matrix (LG), empirical aminoacid frequencies from alignment (F), and 8 discrete GAMMA categories (G8). This was followed by bootstrapping with 100 replicates to determine the confidence in the tree topology.[22]

##### 1.4.2 Protein sequences

All protein sequences used were retrieved from either the NCBI, UniProt, or PDB databases.

SCAD-Bacillus_subtilis (WP_000545520.1), SCAD-Pseudomonas_aeruginosa (NP_251242.1), SCAD-Rhodococcus_jostii (WP_011595336.1), SCAD-Streptomyces_avermitilis (WP_010988001.1), SCAD-Streptomyces_griseus (WP_012381543.1), SCAD-Myxococcus_xanthus (WP_011553809.1), SCAD-Sus_scrofa (NP_999063.1), SCAD-Homo_sapiens (2VIG), SCAD-Rattus_norvegicus (1JQI), MCAD-Pseudomonas_aeruginosa (CAL5248463.1), MCAD-Rhodococcus_jostii (WP_009478133.1), MCAD-Streptomyces_avermitilis (WP_010986431.1), MCAD-Streptomyces_griseus (WP_012380636.1), MCAD-Myxococcus_xanthus (WP_011555872.1), MCAD-Sus_scrofa (NP_999204.1), MCAD-Homo_sapiens (NP_000007.1), MCAD-Rattus_norvegicus (NP_058682.2), LCAD-Myxococcus_xanthus (WP_011553103.1), LCAD-Pseudomonas_aeruginosa (NP_253125.1), LCAD-Rhodococcus_jostii (WP_011593682.1), LCAD-Streptomyces_avermitilis (WP_010986684.1), LCAD-Streptomyces_griseus (WP_012378229.1), LCAD-Sus_scrofa (NP_999062.1), LCAD-Homo_sapiens (NP_001599.1), VLCAD-Streptomyces_avermitilis (WP_010984003.1), VLCAD-Streptomyces_griseus (WP_012378801.1), VLCAD-Homo_sapiens (3B96), VLCAD-Rattus_norvegicus (NP_037023.1), VLCAD-Bacillus_rossius (XP_063218815.1), ACOX-Homo_sapiens (spO15254), ACOX-Saccharomyces_cerevisiae (sp_P13711), ACOX-Candida_tropicalis (sp_P06598.3), ACOX-Yarrowia_lipolytica (sp_O74936.1), ACOX-Arabidopsis_thaliana (sp_O65202.1), ACOX-Bos_taurus (sp_Q3SZP5.1), TmcF (CUX96953), EpnF (WP_402041890.1), tryptopeptin_EKS-Streptomyces_sparsogenes (WP_362202621.1), tryptopeptin_EKS_Streptomyces_maeda (Unpublished data), EpxF

(AHB38499.1), MynC (WP_002626007.1), landepoxcin_EKS (AKA59435.1), clarepoxcin_EKS (AKA59449.1), MatG (WP_019634557.1), SnaO (WP_013019281.1)

##### 1.4.3 AlphaFold modelling and MD simulations

The EpnF dimer used in MD simulations was predicted using ColabFold (implemented locally with localcolabfold (https://github.com/YoshitakaMo/localcolabfold)) using Alphafold2 weights.[23,24] Fully oxidised FAD was placed by structural alignment with homologous proteins using BLAST in ChimeraX.[25] Clashes were eliminated by energy minimisation in Chimera.[26] In simulation 1 the N-acyl-dipeptidyl β-ketoacid substrate was placed manually in VR in the active site of one monomer and the active site of the other monomer was left without the substrate. In simulation 2 the FAD molecules in the active site of one monomer were modified to the fully reduced, dihydrogenated form, $FADH_2$ and to $FADH_2O_2$ in the active site of the other monomer. Enone intermediate molecules were placed manually in VR in both active sites. In simulation 3 one monomer was loaded with $FADH_2O_2$ and enone intermediate and other was loaded with $FADH^-$ and enone intermediate.

Molecular dynamics (MD) simulations were performed using the AMBER ff19SB force field.[27] All the ligands were parametrised using ANTECHAMBER and GAFF force field with bcc charge model.[28–30] The charge states of the amino acids at pH 7.4 were evaluated using the H++ webserver.[31] The resulting structure was neutralised with $Na^+$ ions and solvated with OPCBOX water, such that no atom belonging to the complex was less than 10 Å from any box edge, using the LEaP module.[32] Additional $Na^+$ and $Cl^-$ ions were added to obtain 150 mM final salt concentration. MD heating, equilibration, and production steps were performed using the GPU accelerated AMBER software on an HPC cluster equipped with RTX6000 graphics cards or local workstation equipped with RTX2080Ti graphics cards.[33] Simulation used the SHAKE algorithm to constrain all protein bonds involving a hydrogen atom; a 2.0 fs time-step was used in these simulations.[34] Long-range electrostatics were calculated using the Particle Mesh Ewald (PME) method with a 12.0 Å cut-off.[35] PME was used for nonbonded interactions. In all simulations, the Langevin thermostat ($\gamma$ = 2.0 $ps^{-1}$) was used to maintain temperature control.[36] The solvated protein was then equilibrated by carrying out a short minimization, 50 ps of heating and 50 ps of density equilibration with weak restraints on the protein followed by 500 ps of constant pressure equilibration at 300 K. After a two-step minimization process, in which solvent molecules were allowed to relax before the entire system was minimized, the system was slowly heated to 300 K over 0.1 ns in a canonical ensemble (NVT) simulation, then equilibrated for 2 ns by performing isothermal-isobaric (NPT) simulations at 300 K using a Berendsen barostat.[37] For each simulation set up 100 ns production using classic approach (with no acceleration) run were performed with simulation frames written every 20 ps for

analysis. For all simulations, long-range electrostatics were calculated using the PME method with a 8.0 Å cutoff.
A combination of CCPTRAJ, Chimera and ChimeraX were used for analysis of the trajectories, ChimeraX was used throughout the course to prepare and visualize the structures.[26,38,39]

Additional protein models of SnaO, MatG and EpnF were generated using AlphaFold3.[40]

#### 2 Coupled PLE/EpnF assay LC-MS chromatograms

##### 2.1.1 LC-MS analysis of substrates bearing various $R^2$ functional groups

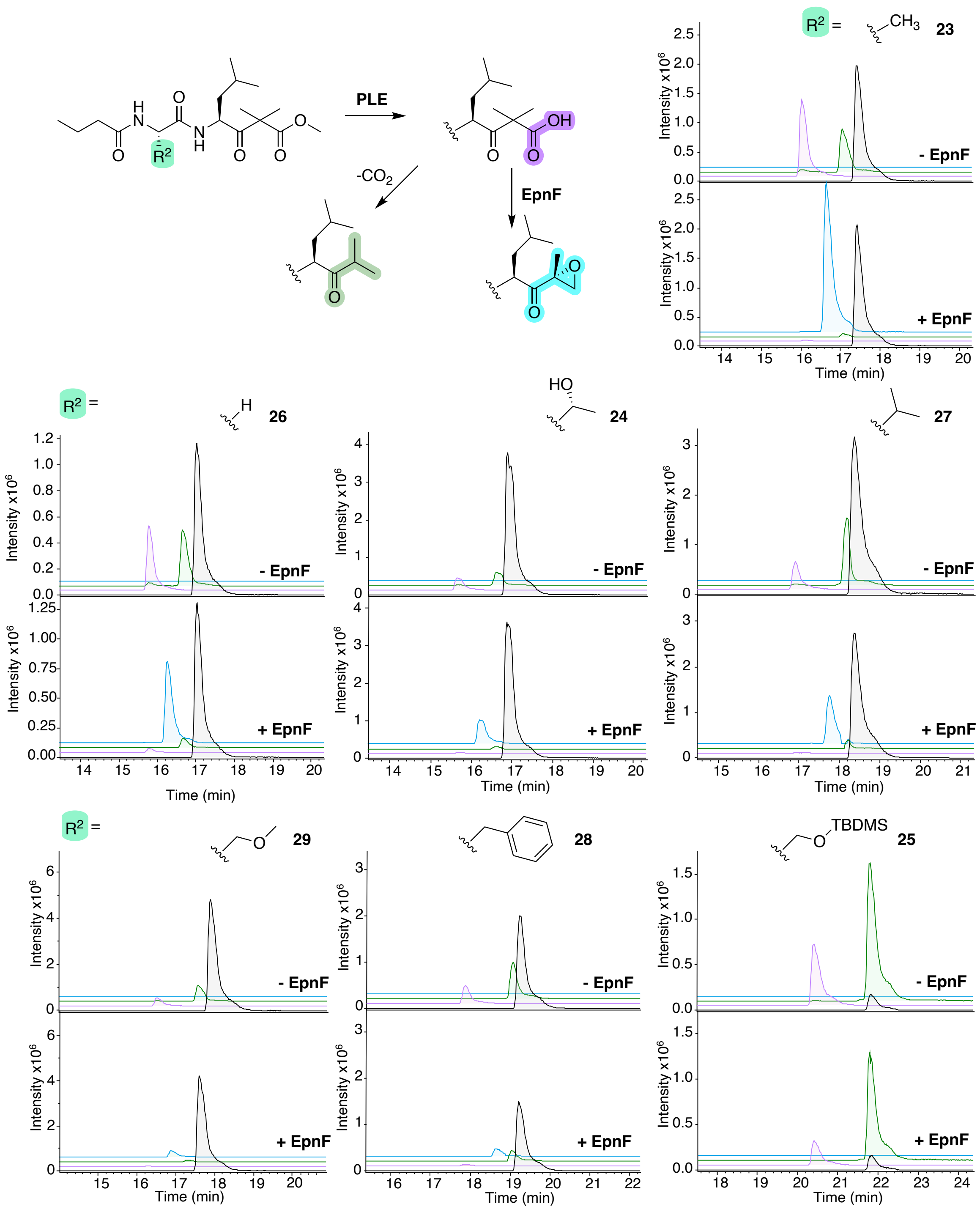

*Figure S1: LC-MS analysis of coupled enzyme assays (1:200 PLE to substrate) with $R^2$-functionalised substrates. EICs for m/z corresponding to both $[M+H]^+$ and $[M+Na]^+$ for the methyl ester starting material (grey), β-keto-acid hydrolysis product (purple), spontaneous decarboxylation product (green) and epoxyketone (cyan) from the control reactions without EpnF and PLE/EpnF reactions.*

##### 2.1.2 LC-MS analysis of substrates bearing various $R^1$ functional groups

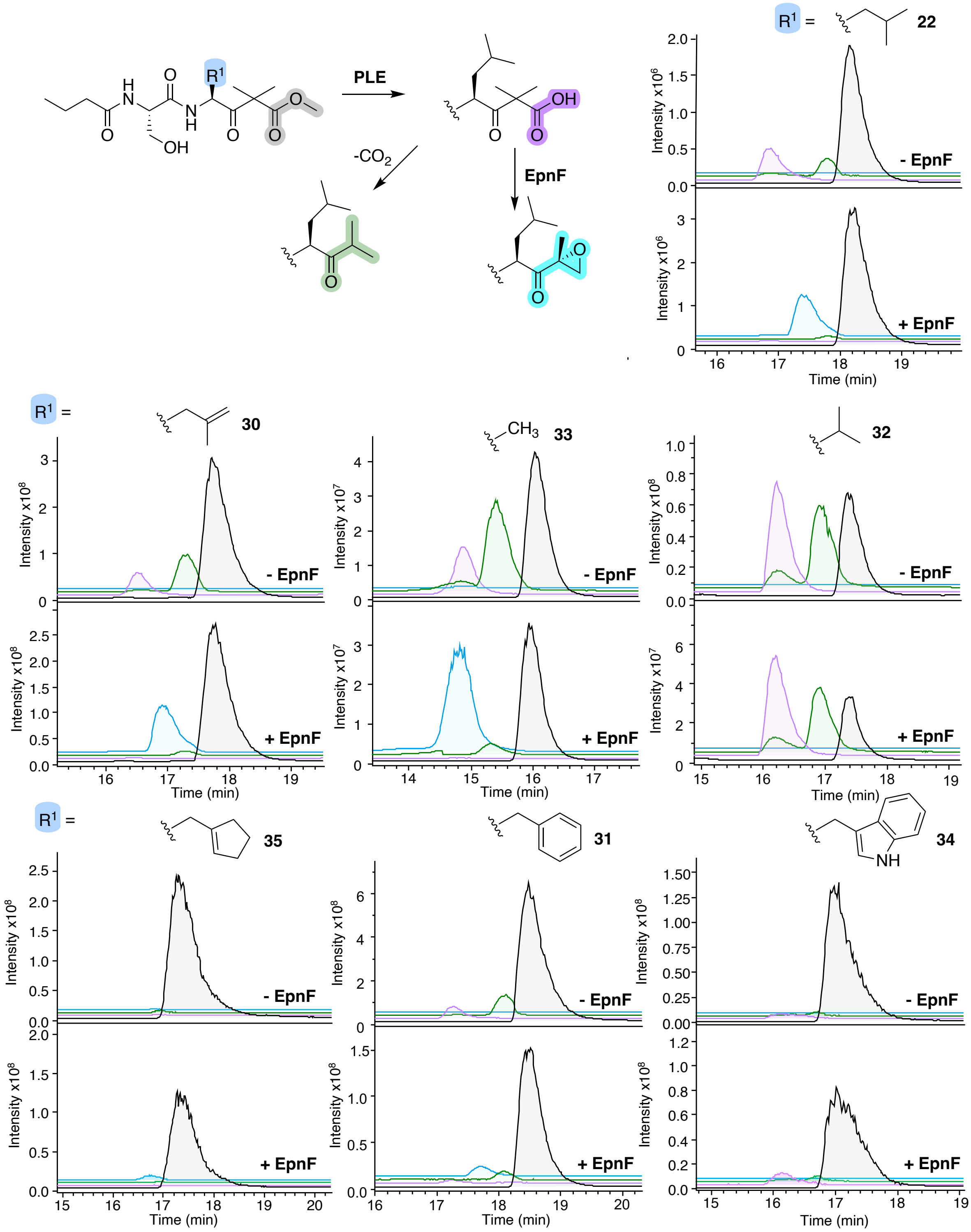

*Figure S2: LC-MS analysis of coupled enzyme assays (1:10 PLE to substrate) with $R^1$-functionalised substrates. EICs for m/z corresponding to both $[M+H]^+$ and $[M+Na]^+$ for the methyl ester starting material (grey), β-keto-acid hydrolysis product (purple), spontaneous decarboxylation product (green) and epoxyketone (cyan) from the control reactions without EpnF and PLE/EpnF reactions.*

##### 2.1.3 LC-MS analysis of substrates bearing various $R^3$ functional groups

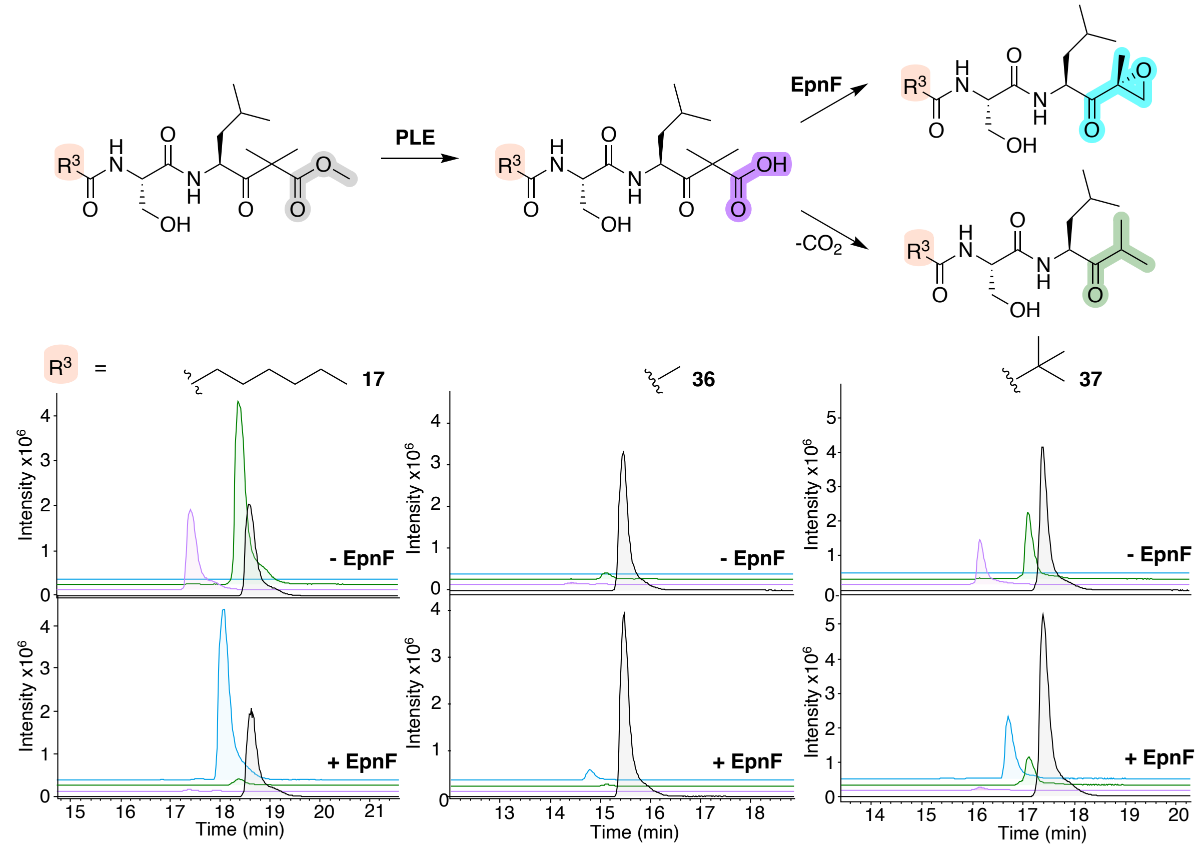

*Figure S3: LC-MS analysis of coupled enzyme assays (1:200 PLE:substrate) with $R^3$-functionalised substrates. Possible reaction products formed in coupled enzyme assays (top). EICs for m/z corresponding to both $[M+H]^+$ and $[M+Na]^+$ for the methyl ester starting material (grey), β-keto-acid hydrolysis product (purple), spontaneous decarboxylation product (green) and epoxyketone (cyan) from the control reactions without EpnF and PLE/EpnF reactions (bottom).*

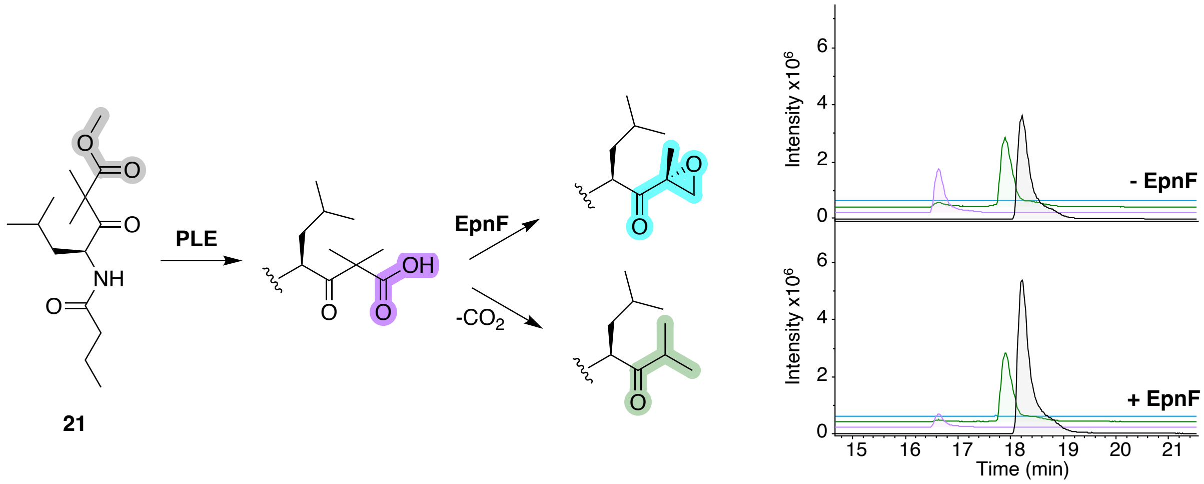

*Figure S4: EICs for m/z corresponding to both $[M+H]^+$ and $[M+Na]^+$ for the methyl ester starting material* ***21*** *(grey), β-keto-acid hydrolysis product (purple), spontaneous decarboxylation product (green) and epoxyketone (cyan) from the control reactions without EpnF and PLE/EpnF reactions (bottom).*

##### 2.1.4 LC-MS analysis of substrates bearing various $R^4$ functional groups

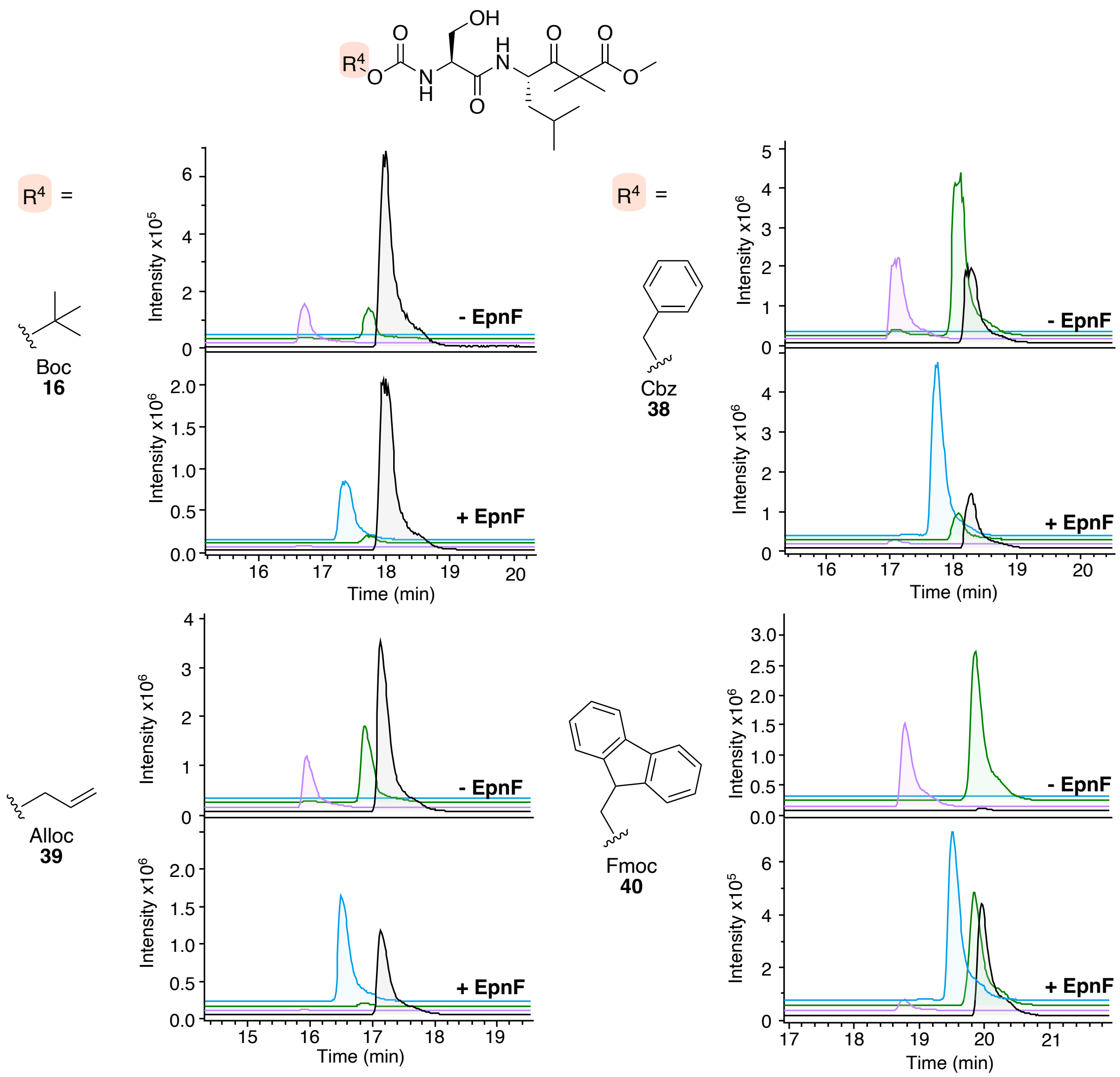

*Figure S5: LC-MS analysis of coupled PLE/EpnF assays (1:200 PLE to substrate) with N-protected α-dimethyl-β-keto-ester substrates. EICs for m/z corresponding to both [M+H]+ and [M+Na]+ for the methyl ester starting material (grey), β-keto-acid (purple), spontaneous decarboxylation product (green) and epoxyketone (cyan) from the control reactions without EpnF and PLE/EpnF reactions.*

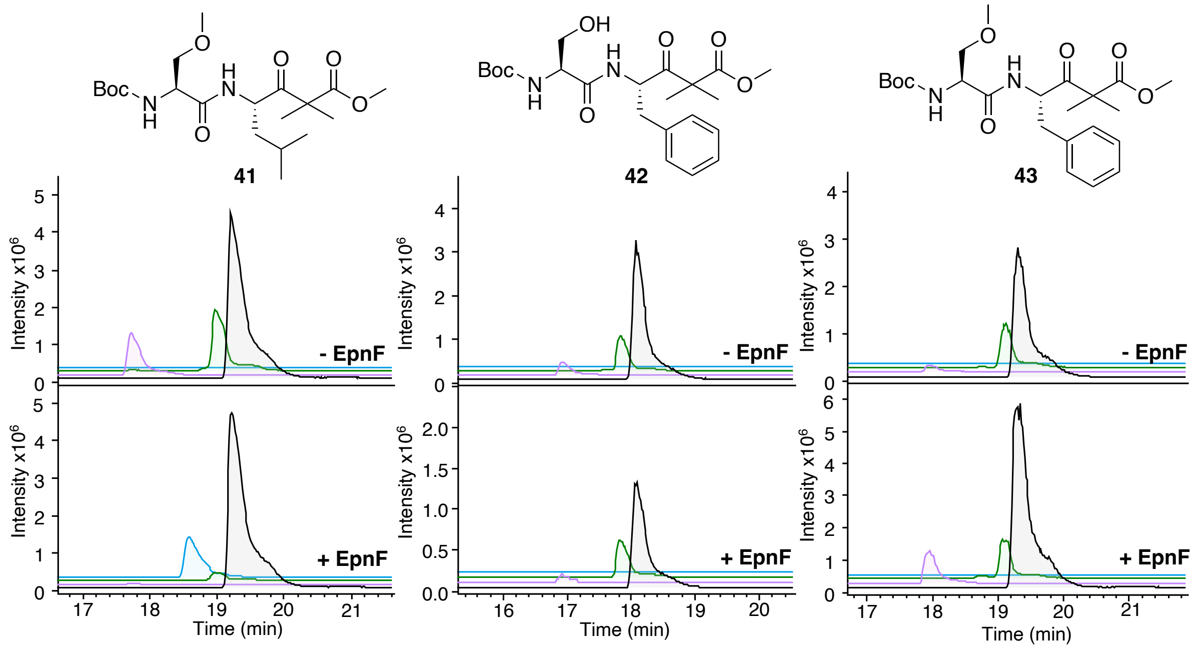

*Figure S6: LC-MS analysis of coupled PLE/EpnF assays (1:200 PLE to substrate) with Oprozomib-related N-protected α-dimethyl-β-keto-ester substrates. EICs for m/z corresponding to both [M+H]+ and [M+Na]+ for the methyl ester starting material (grey), β-keto-acid (purple), spontaneous decarboxylation product (green) and epoxyketone (cyan) from the control reactions without EpnF and PLE/EpnF reactions.*

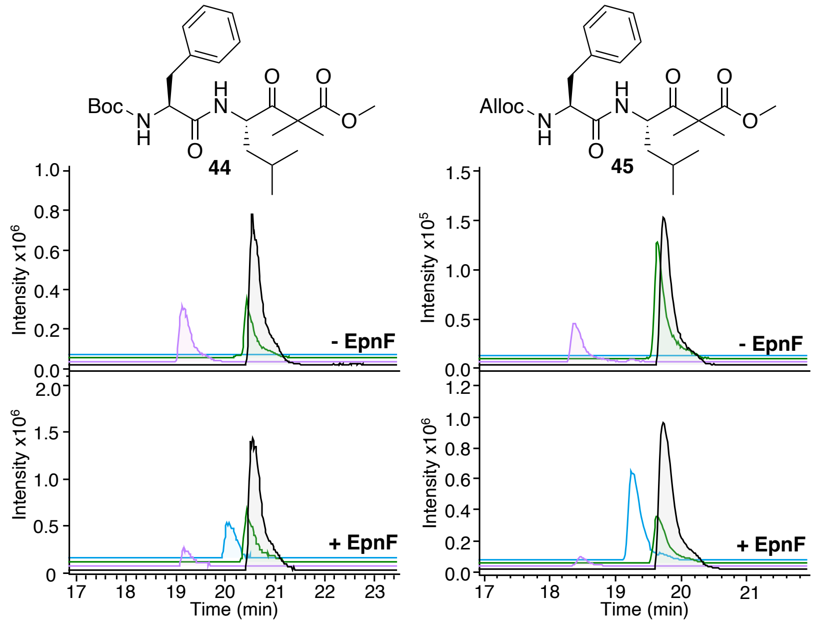

*Figure S7: LC-MS analysis of coupled PLE/EpnF assays (1:200 PLE to substrate) with Carfilzomib-related N-protected α-dimethyl-β-keto-ester substrates. EICs for m/z corresponding to both [M+H]+ and [M+Na]+ for the methyl ester starting material (grey), β-keto-acid (purple), spontaneous decarboxylation product (green) and epoxyketone (cyan) from the control reactions without EpnF and PLE/EpnF reactions.*

##### 2.1.5 LC-MS comparisons of an enone authentic standard with PLE/EpnF reaction products

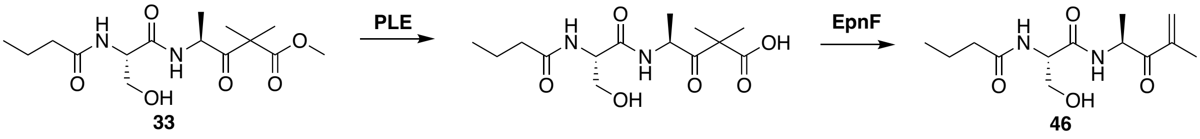

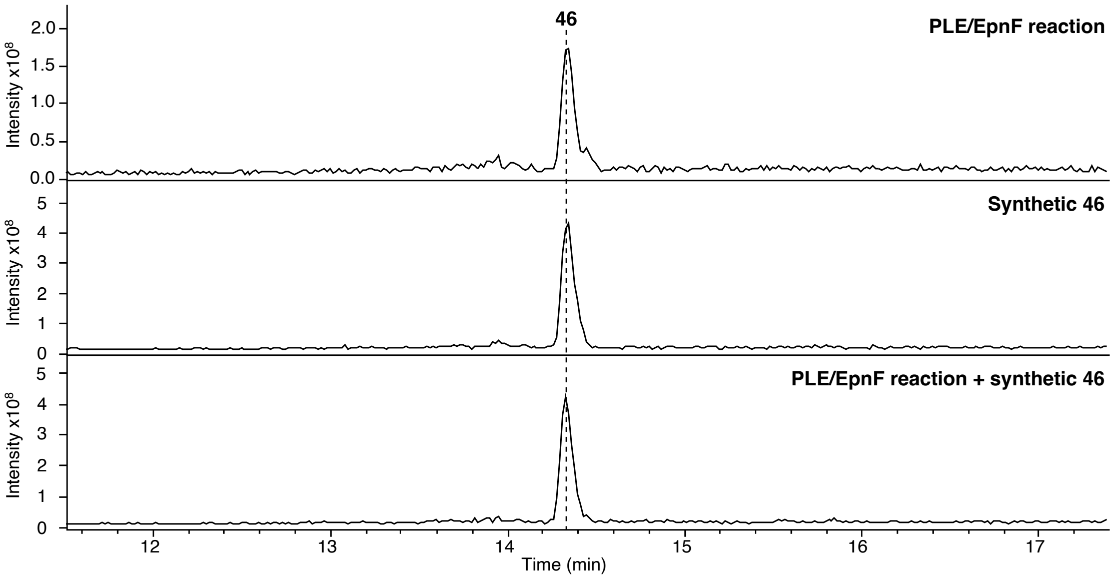

Figure S8: *EICs for [M + Na]$^+$ for **46** from UHPLC-ESI-Q-TOF MS analyses. Supernatant from the reaction with PLE, EpnF and **33** (top trace), authentic standard **46** (middle trace), combined supernatant and authentic standard **46** (bottom trace).*

### 3 HR-MS analysis of chemoenzymatically-derived epoxyketones

#### 3.1.1 R[1]-functionalised epoxyketones

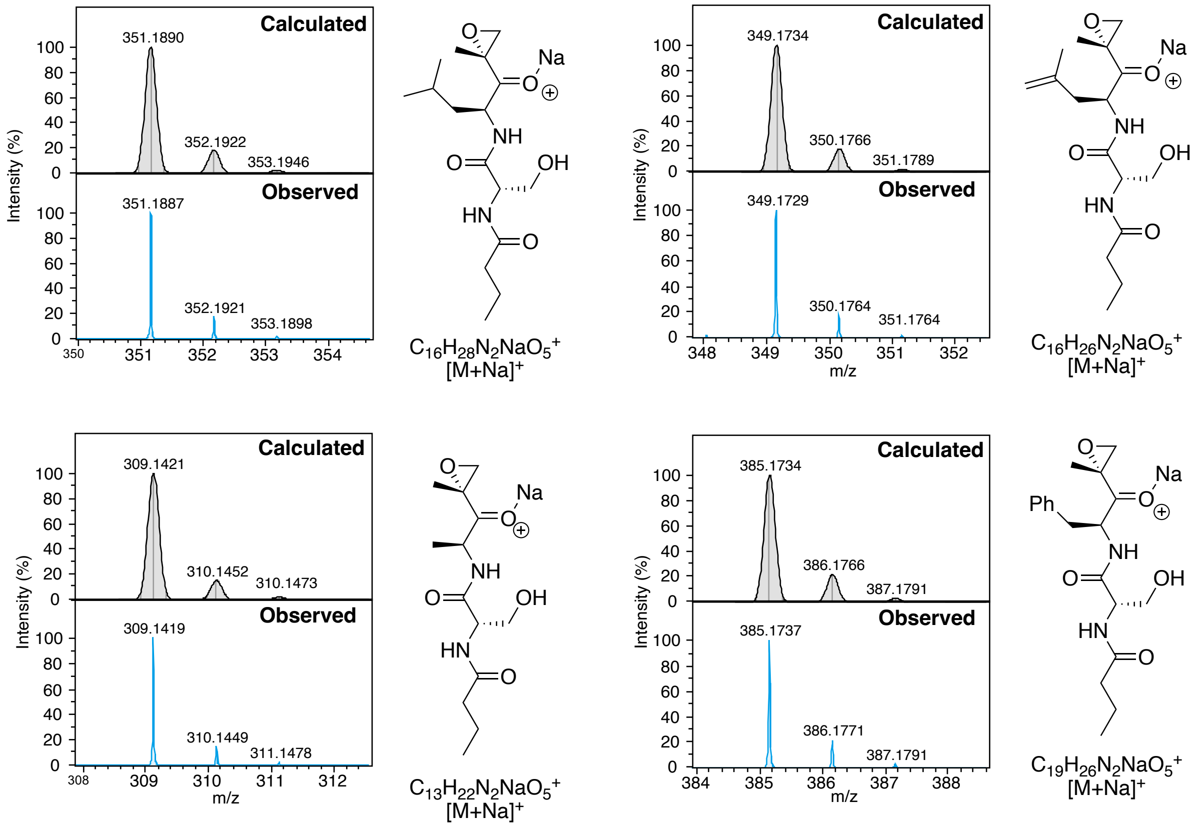

*Figure S9: Simulated mass spectra (top panels) and observed HR-MS spectra (bottom panels) for R[1]-functionalised epoxyketone species generated in coupled enzyme assays.*

##### 3.1.2 $R^2$-functionalised epoxyketones

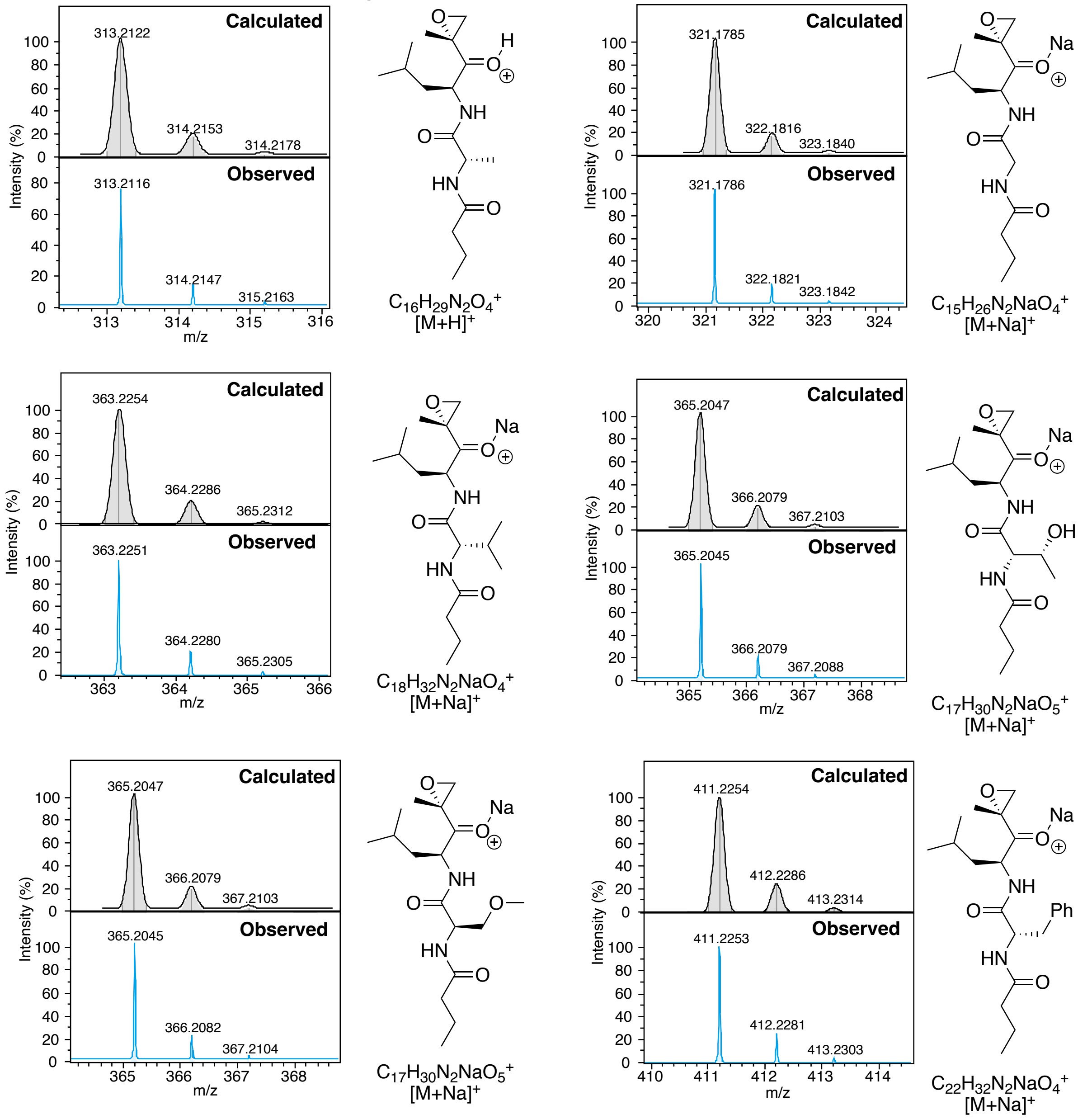

*Figure S10: Simulated mass spectra (top panels) and observed HR-MS spectra (bottom panels) for $R^2$-functionalised epoxyketone species generated in coupled enzyme assays.*

##### 3.1.3 R³-functionalised epoxyketones

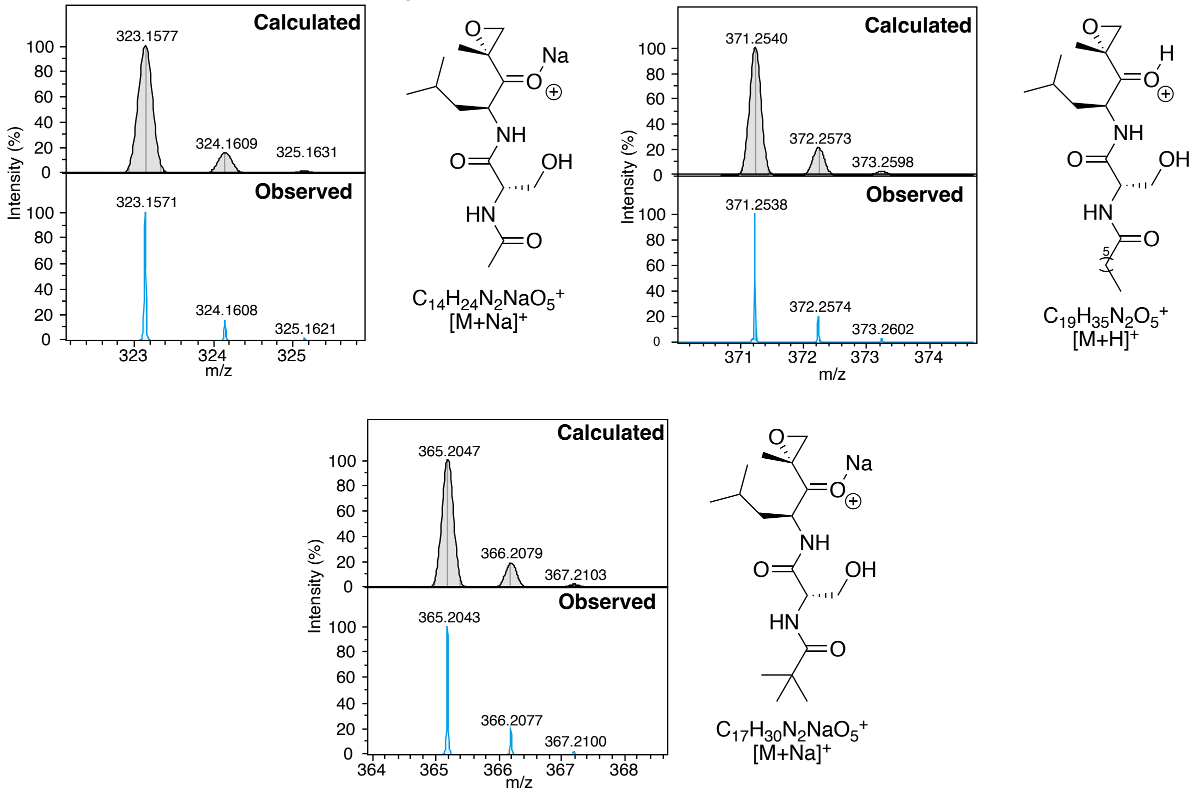

*Figure S11: Simulated mass spectra (top panels) and observed HR-MS spectra (bottom panels) for R³-functionalised epoxyketone species generated in coupled enzyme assays.*

##### 3.1.4 R⁴-functionalised dipeptidyl epoxyketones

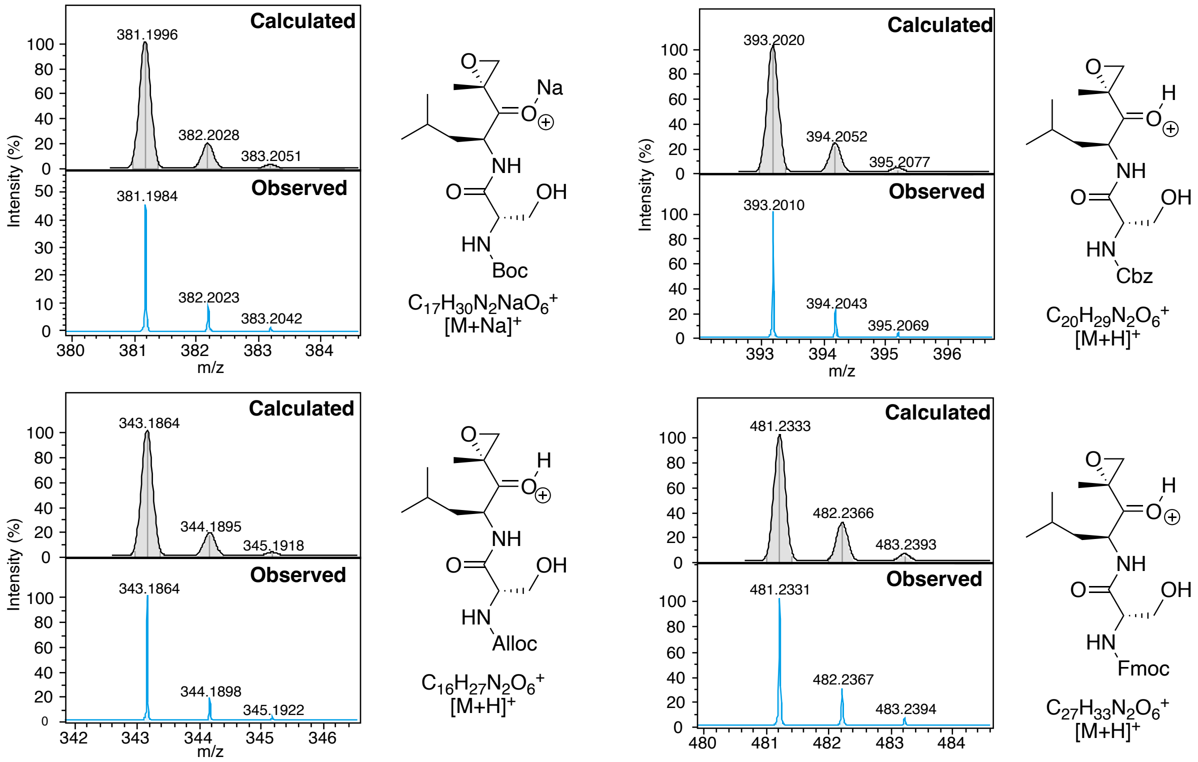

*Figure S12: Simulated mass spectra (top panels) and observed HR-MS spectra (bottom panels) for N-protected epoxyketone species generated in coupled enzyme assays.*

##### 3.1.5 Oprozomib-related epoxyketone

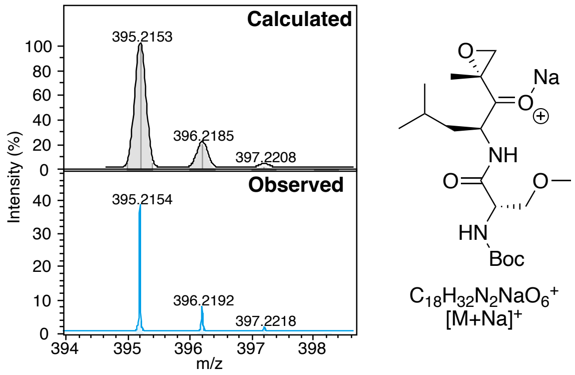

*Figure S13: Simulated mass spectra (top panels) and observed HR-MS spectra (bottom panels) for N-Boc, R²-hydroxymethyl epoxyketone species generated in coupled enzyme assays.*

##### 3.1.6 Dipeptidyl Carfilzomib fragment epoxyketones

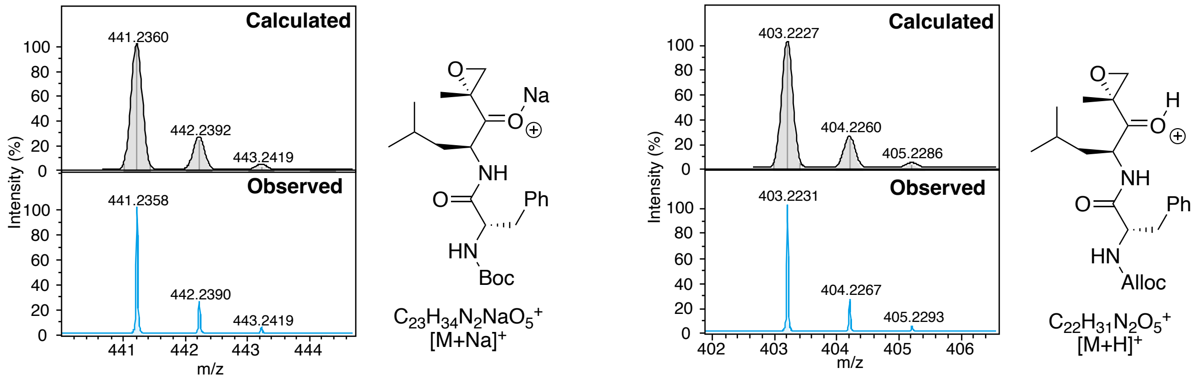

*Figure S14: Simulated mass spectra (top panels) and observed HR-MS spectra (bottom panels) for N-protected epoxyketone species which could act as a late-stage intermediate in Carfilzomib synthesis, generated in coupled enzyme assays.*

### 4 Protein biochemistry and bioinformatic analysis

#### 4.1 Phylogenetic analysis of epoxyketone synthases, ACADs and ACOs

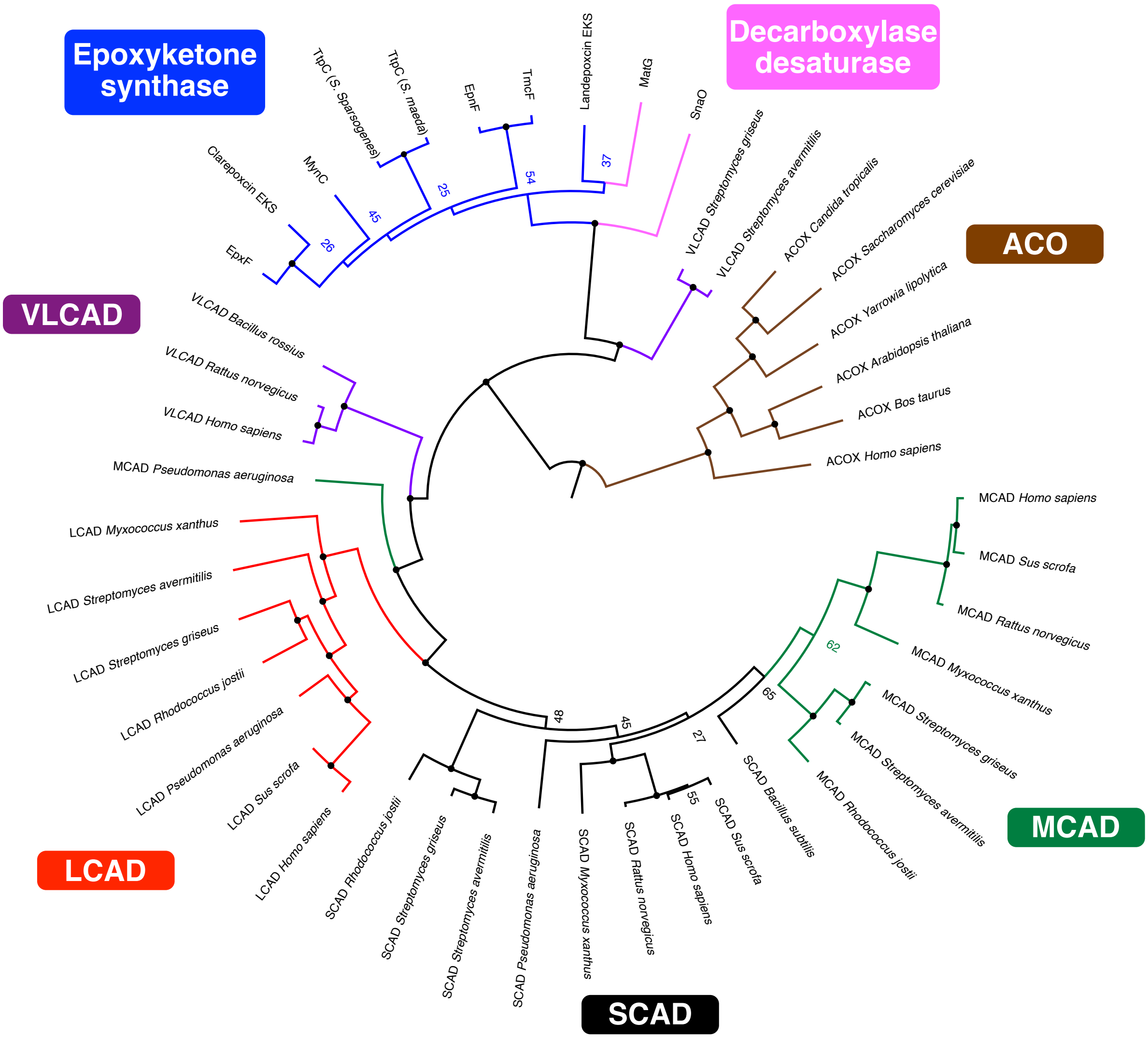

*Figure S15: Phylogenetic analysis of SCADs, MCADs, LCADs, VLCADs, ACOs, decarboxylase-desaturases and epoxyketone synthases. Branch points with a bootstrap value of greater than 70 are marked by a circle*

##### 4.2 Structural comparisons of VLCAD crystal structure with EpnF AlphaFold 3 model

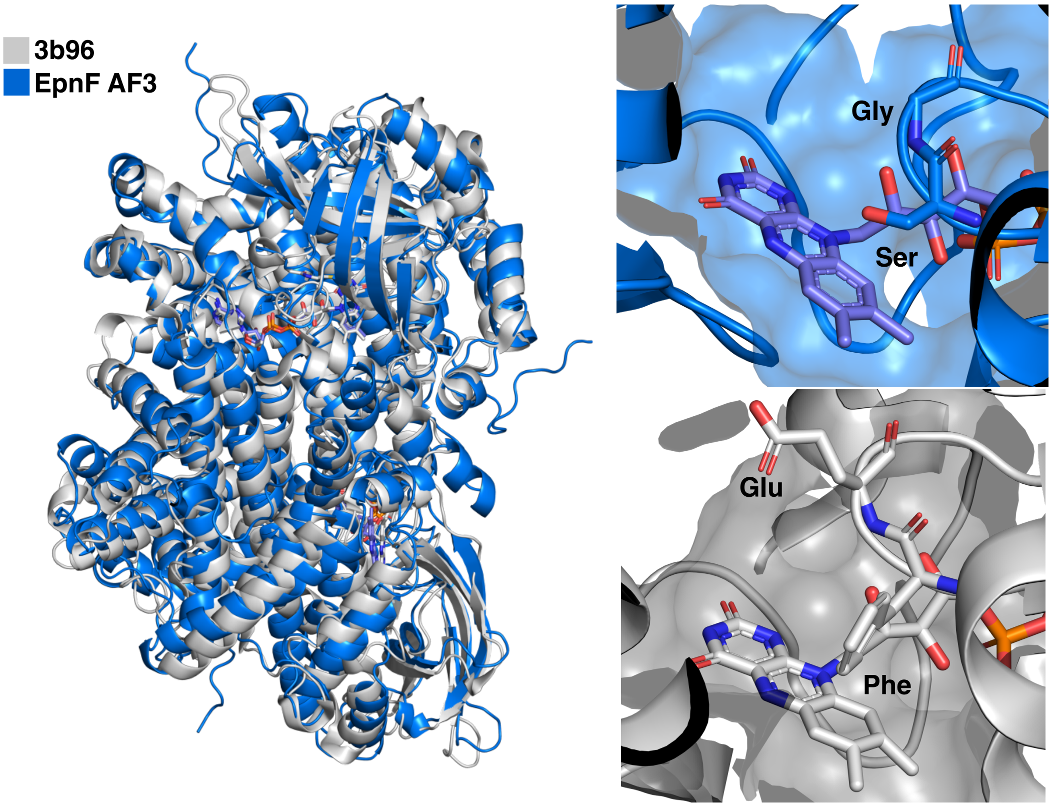

*Figure S16: Structural overlay of a VLCAD crystal structure (PDB: 3b96) with an AlphaFold 3 model of EpnF. RMSD over 651 atoms 2.266 Å*

#### 4.3 Oxyanion hole-like environment around the EpnF flavin cofactor

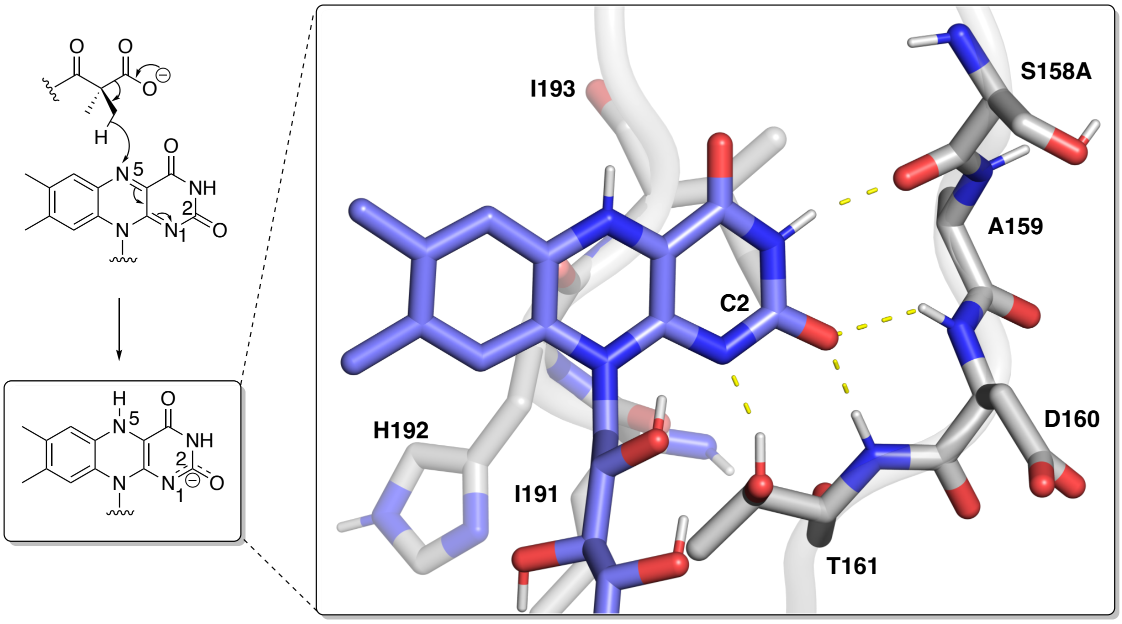

*Figure S17: Structure of the proposed oxyanion-like environment in EpnF stabilising a negative charge at the C2 oxygen atom from the modelling of an anionic FADH⁻ with an EpnF AlphaFold 2.0 model.*

#### 4.4 Plot of distance between the C4a-peroxyflavin peroxide and enone substrate in MD simulations

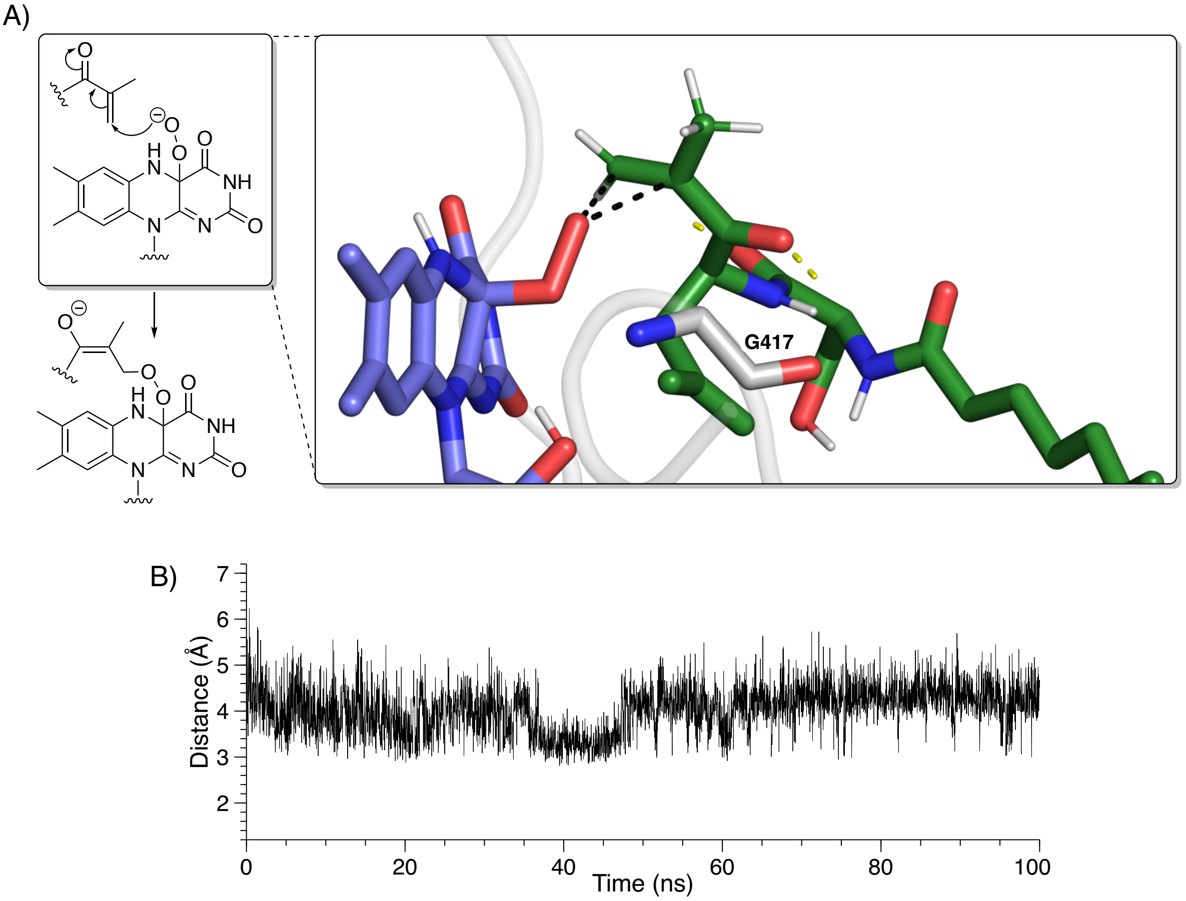

*Figure S18: A) Representative MD simulation frame highlighting binding of the α,β-unsaturated ketone intermediate positioned for epoxidation by C4a-peroxyflavin, B) plot of the calculated distance between the β-carbon of the enone and the distal oxygen of the flavin–C4a–peroxide throughout the MD simulation.*

#### 4.5 SDS-PAGE and intact protein mass spectrometry analyses of purified EpnF mutants

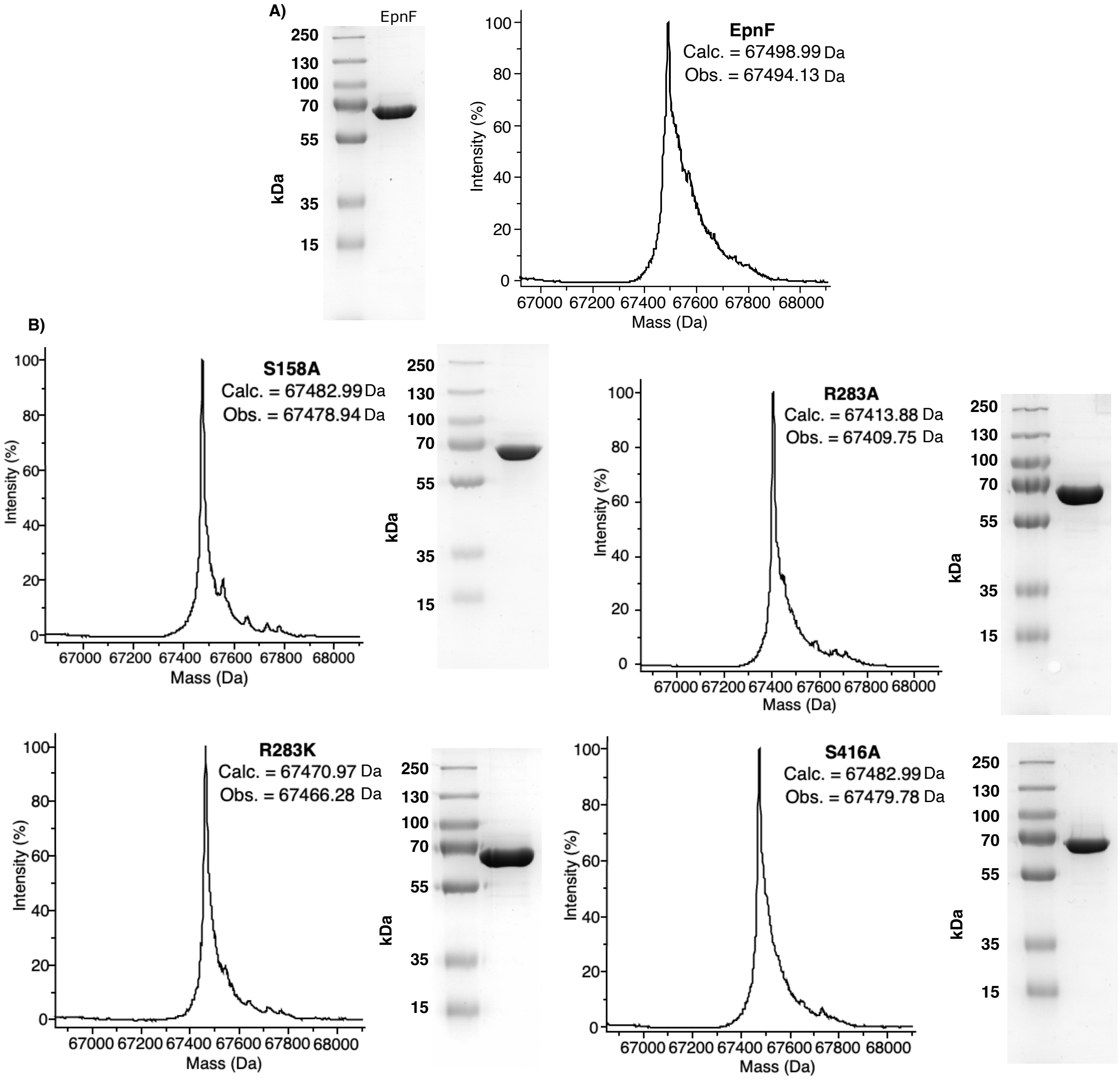

*Figure S19: A) Intact protein mass spectra and SDS-PAGE analysis of purified His$_8$-EpnF, B) Intact protein mass spectra and SDS-PAGE analysis of purified His$_8$-EpnF mutants.*

#### 4.6 CD analysis of purified EpnF mutants

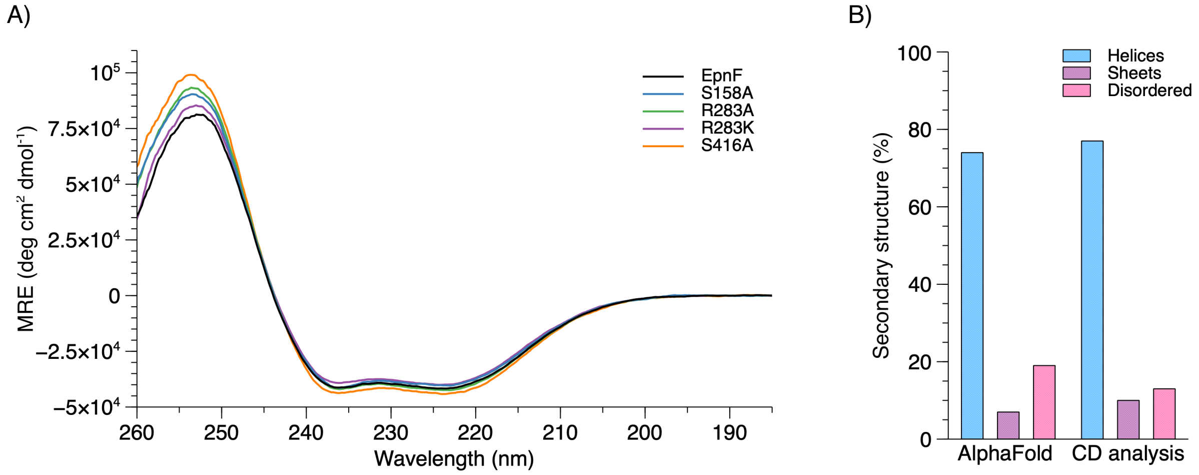

*Figure S20: A) Normalised CD spectra of EpnF and EpnF mutants, B) Comparison of DSSP-predicted secondary structure for the EpnF AlphaFold 2.0 model and Dichroweb-predicted secondary structure from CD analysis of EpnF.*[18]

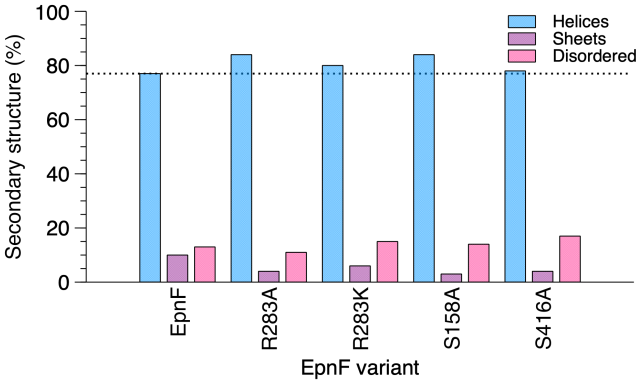

*Figure S21: Dichroweb-predicted secondary structure of EpnF mutants from CD analysis, the 77% helical content of wildtype EpnF is marked by a dotted line.*

#### 4.7 LC-MS analyses of reaction products in assays with EpnF mutants

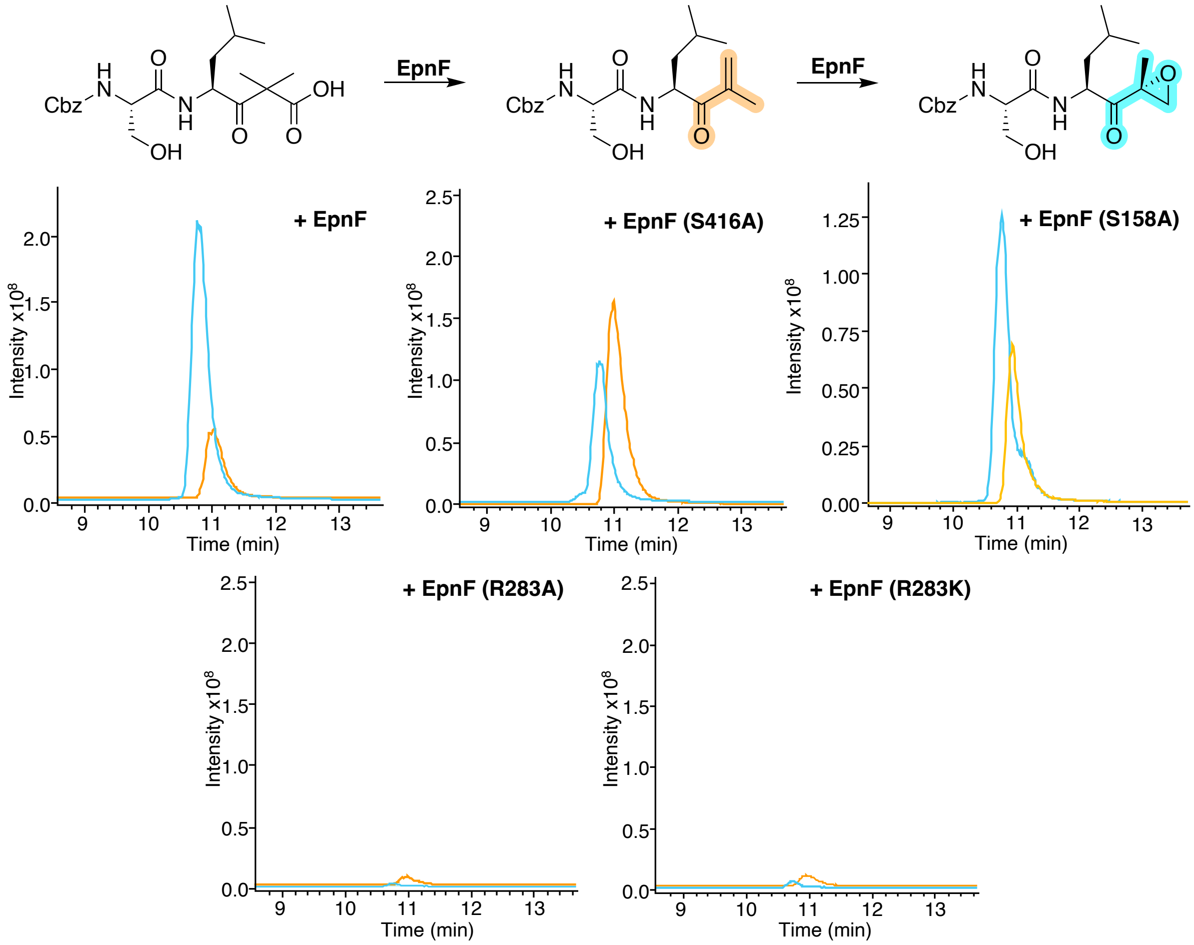

*Figure S22: LC-MS analysis of coupled PLE/EpnF assays (1:200 PLE to substrate) with CBz-protected substrate and EpnF mutants. EICs for m/z corresponding to both [M+H]+ and [M+Na]+ for the α,β unsaturated ketone (orange) and epoxyketone (cyan) from PLE/EpnF assays.*

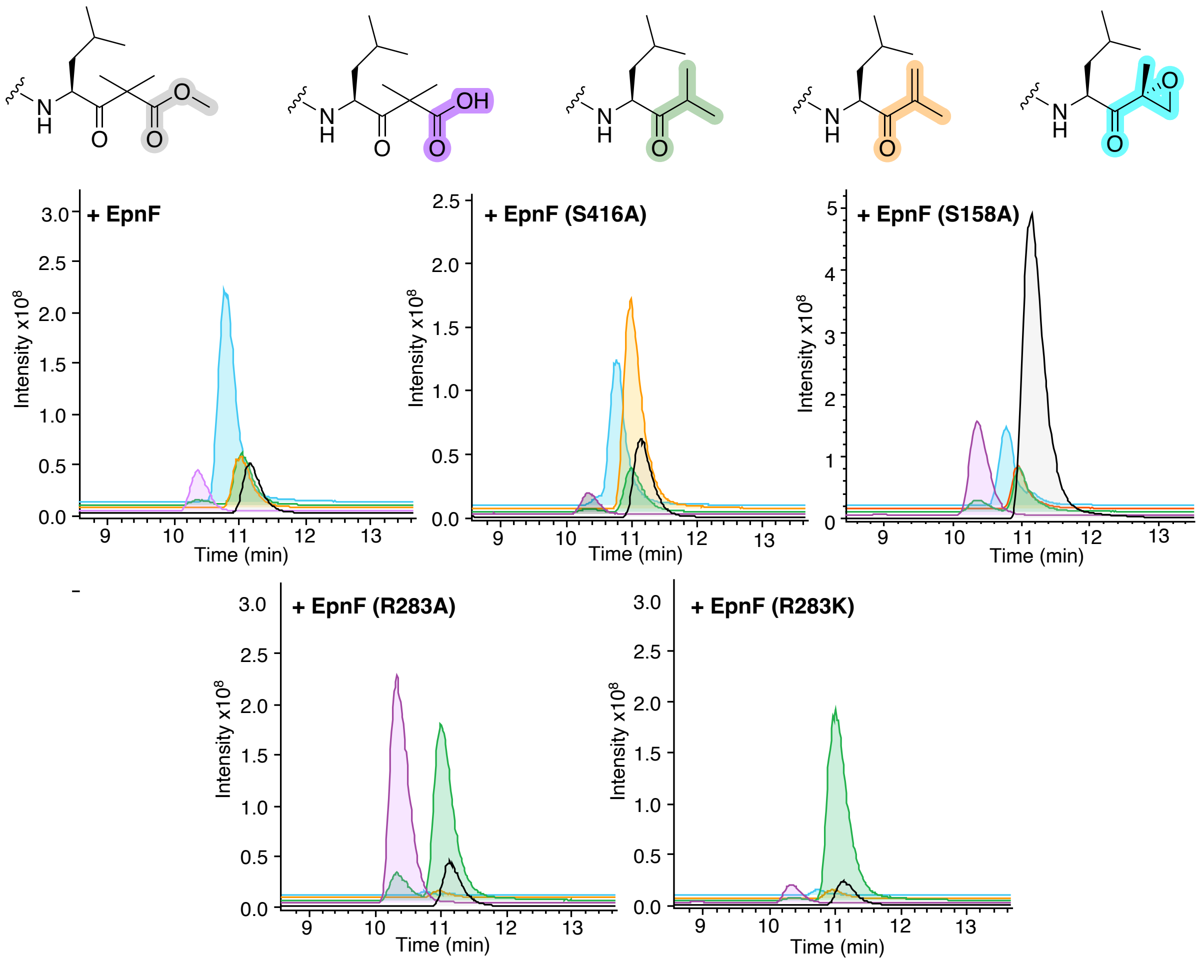

*Figure S23: LC-MS analysis of coupled enzyme assays (1:200 PLE to substrate) with Cbz-protected substrate and EpnF mutants. EICs for m/z corresponding to both $[M+H]^+$ and $[M+Na]^+$ for the methyl ester starting material (grey), β-keto-acid hydrolysis product (purple), spontaneous decarboxylation product (green), enone intermediate (orange) and epoxyketone (cyan) from PLE/EpnF reactions.*

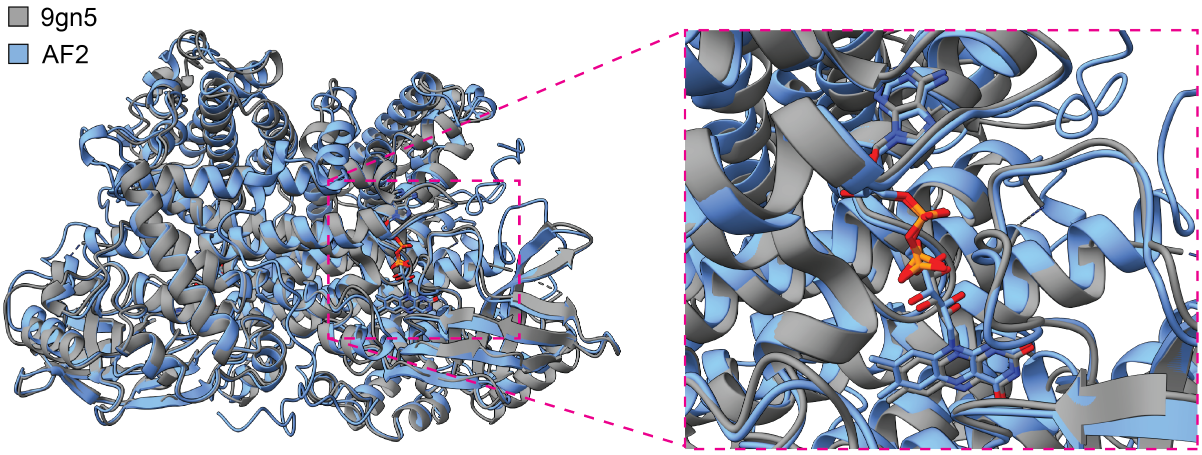

*Figure S24: Structural overlay of the EpxF crystal structure (PDB: 9gn5) with an AlphaFold 2 model of EpnF. RMSD over 483 pruned atoms 0.814 Å, over all 526 pairs 1.688 Å.*

### 5 NMR spectra of synthetic molecules

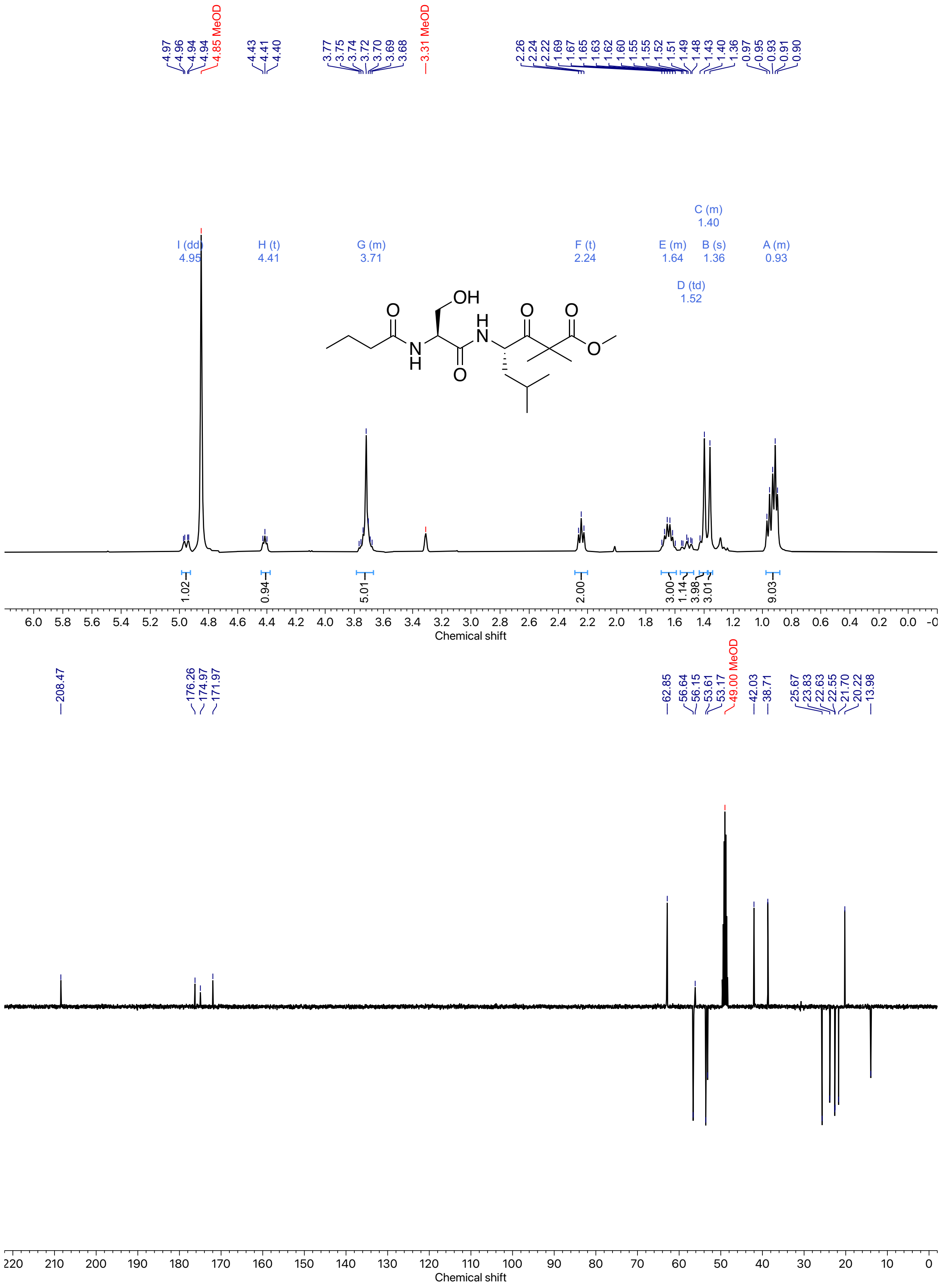

*Figure 25: $^{1}H$ and $^{13}C$ NMR spectra of **22** (Methanol-$d_4$, 400 MHz).*

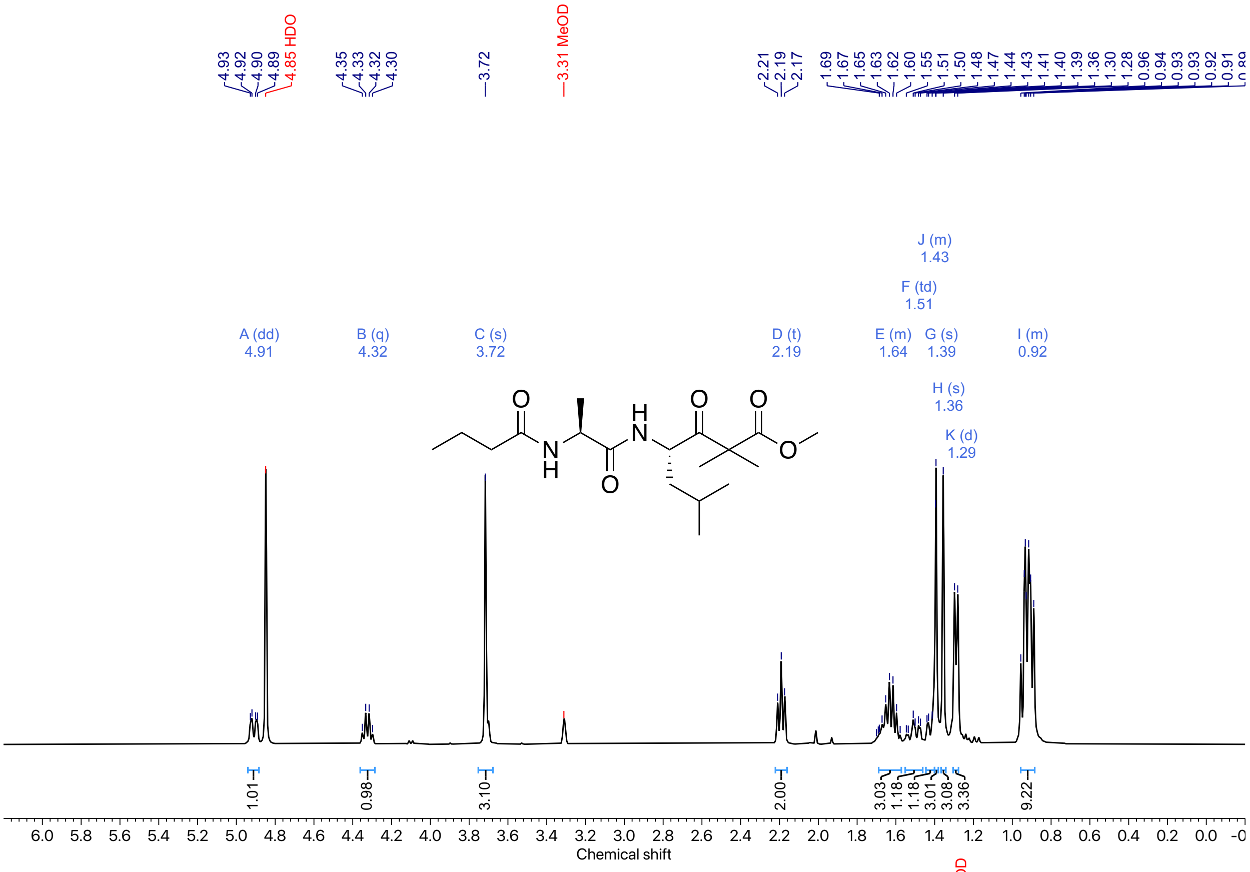

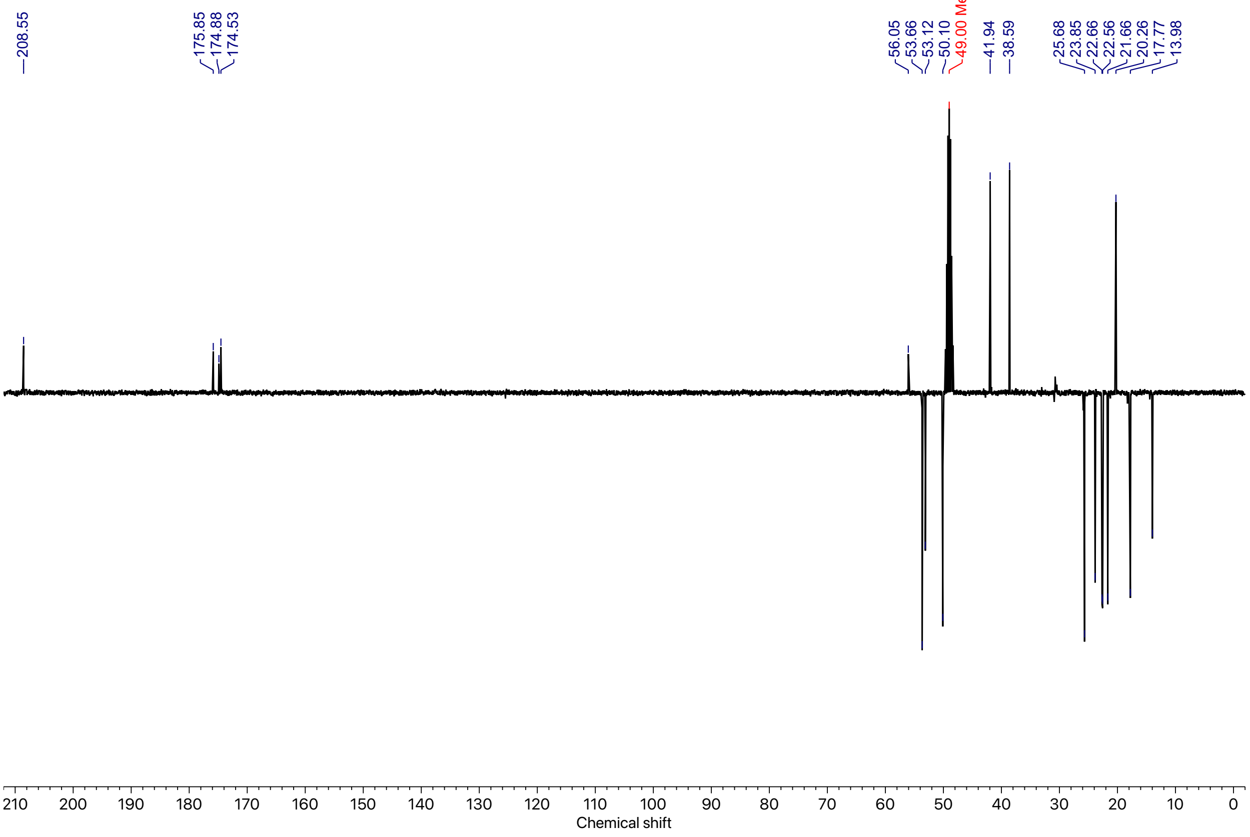

*Figure 26: $^{1}H$ and $^{13}C$ NMR spectra of **23** (Methanol-$d_4$, 400 MHz).*

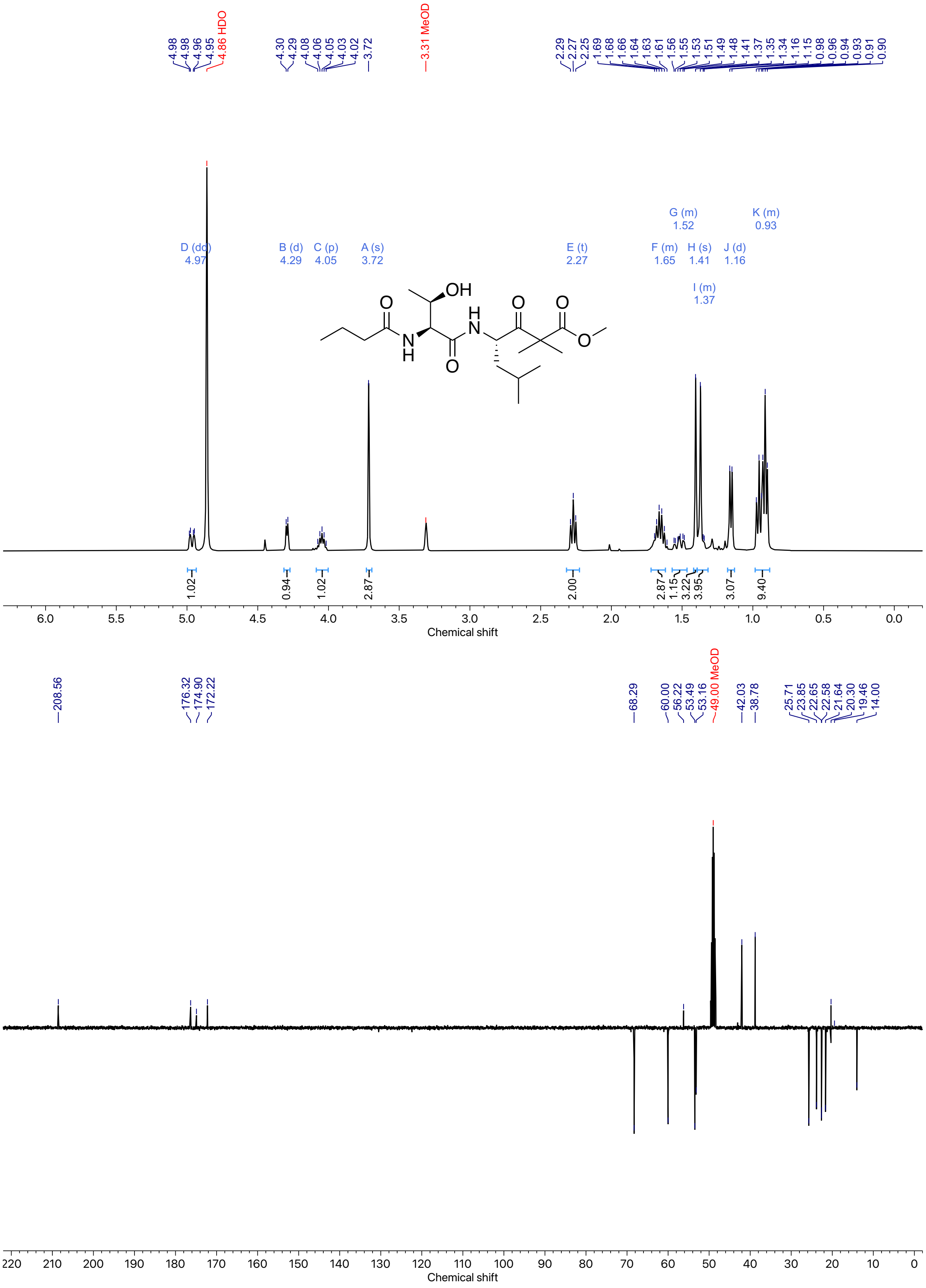

*Figure 27: $^{1}H$ and $^{13}C$ NMR spectra of **24** (Methanol-$d_4$, 400 MHz).*

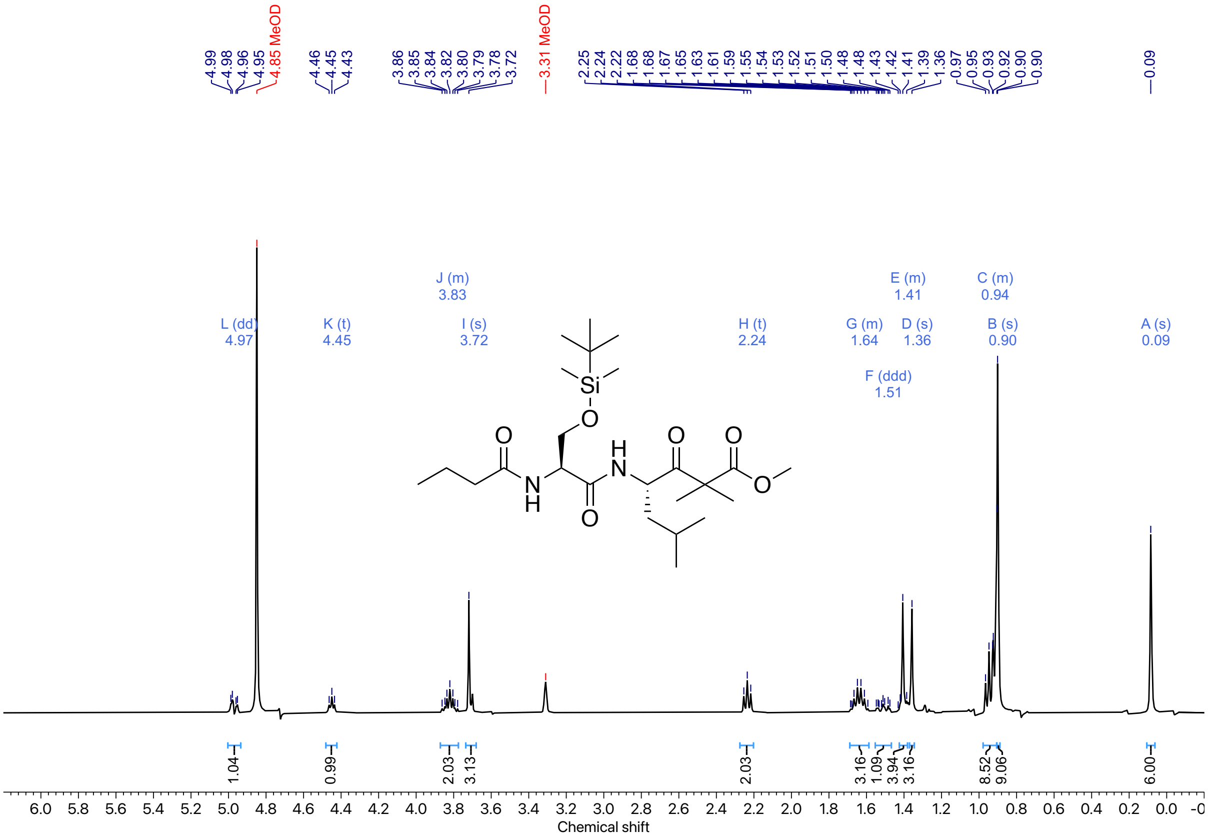

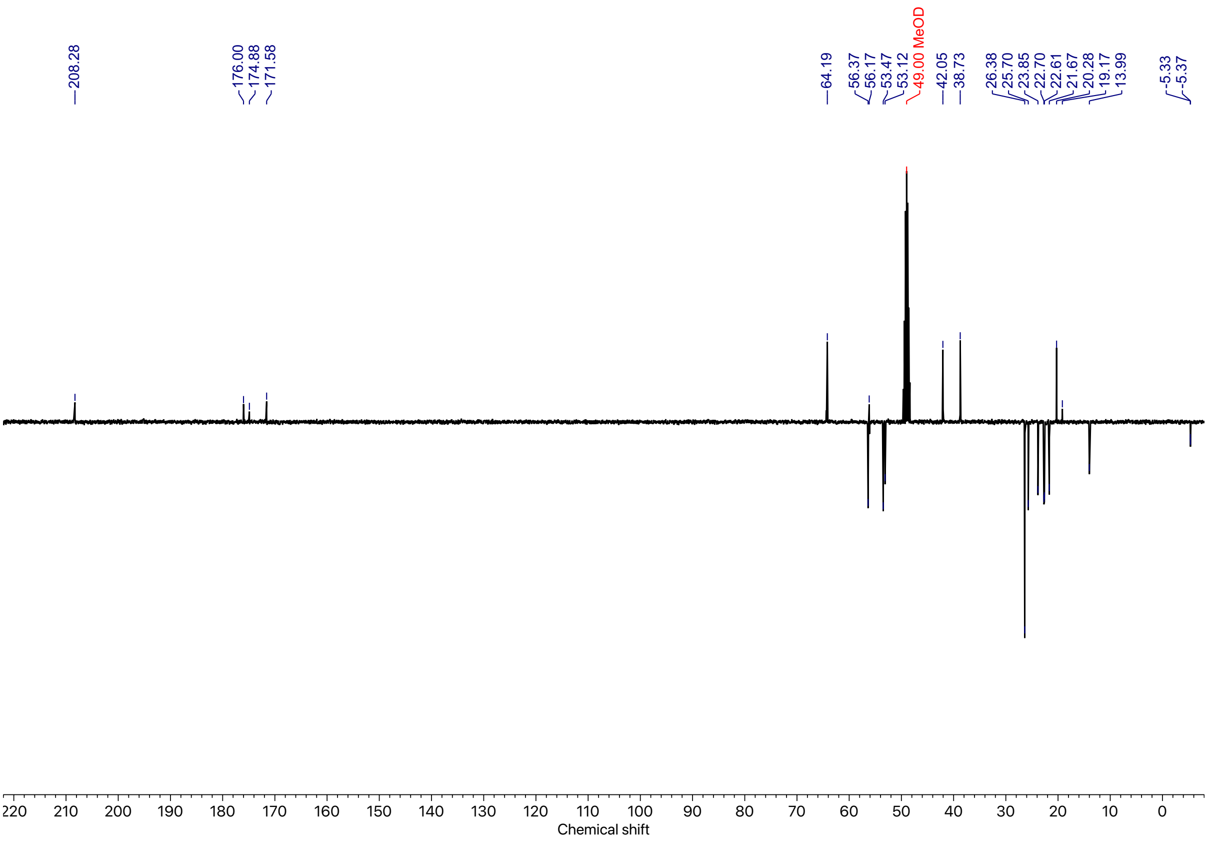

*Figure 28: [1]H and [13]C NMR spectra of **25** (Methanol-d4, 400 MHz).*

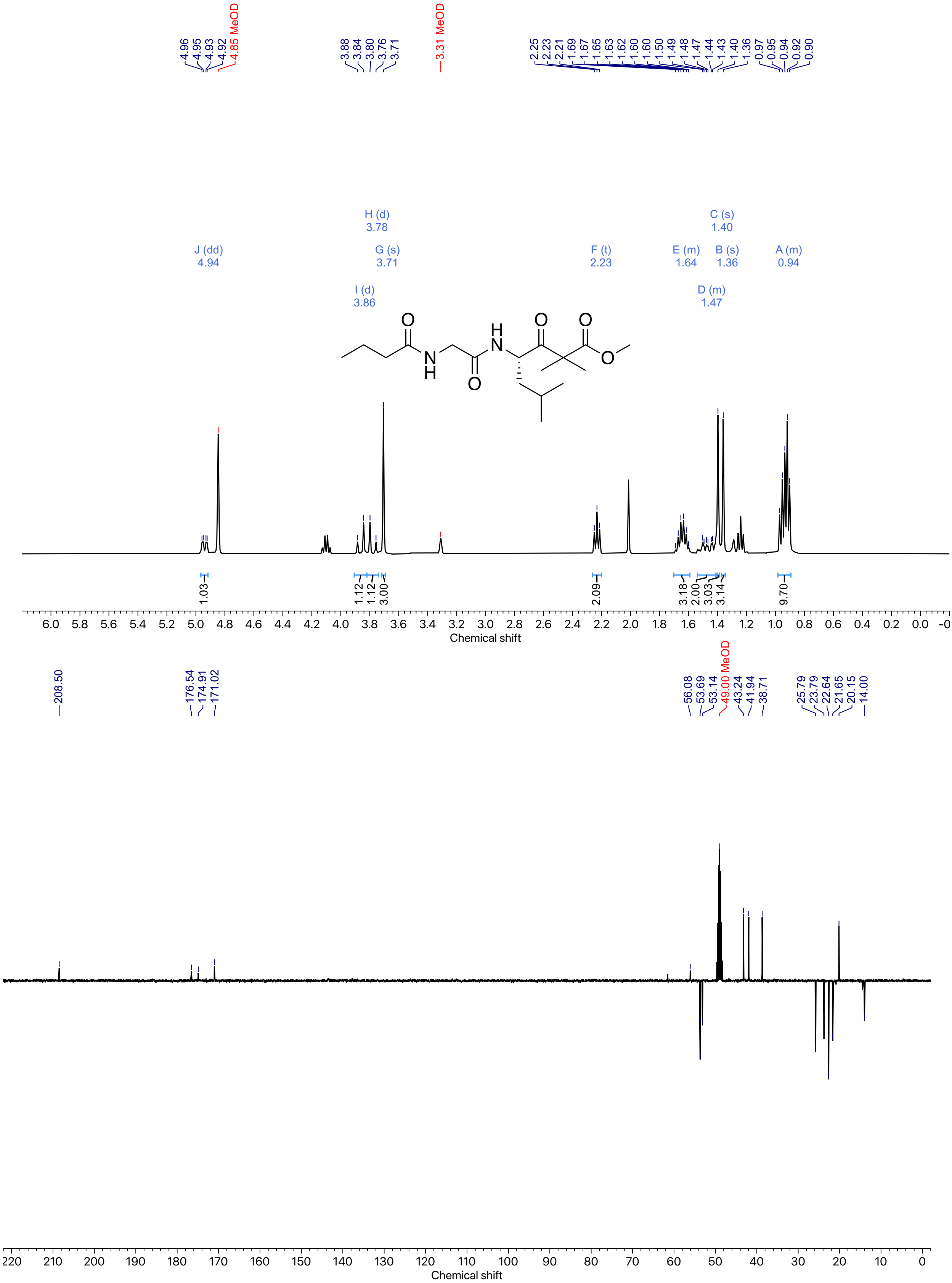

*Figure 29: [1]H and [13]C NMR spectra of **26** (Methanol-d$_4$, 400 MHz).*

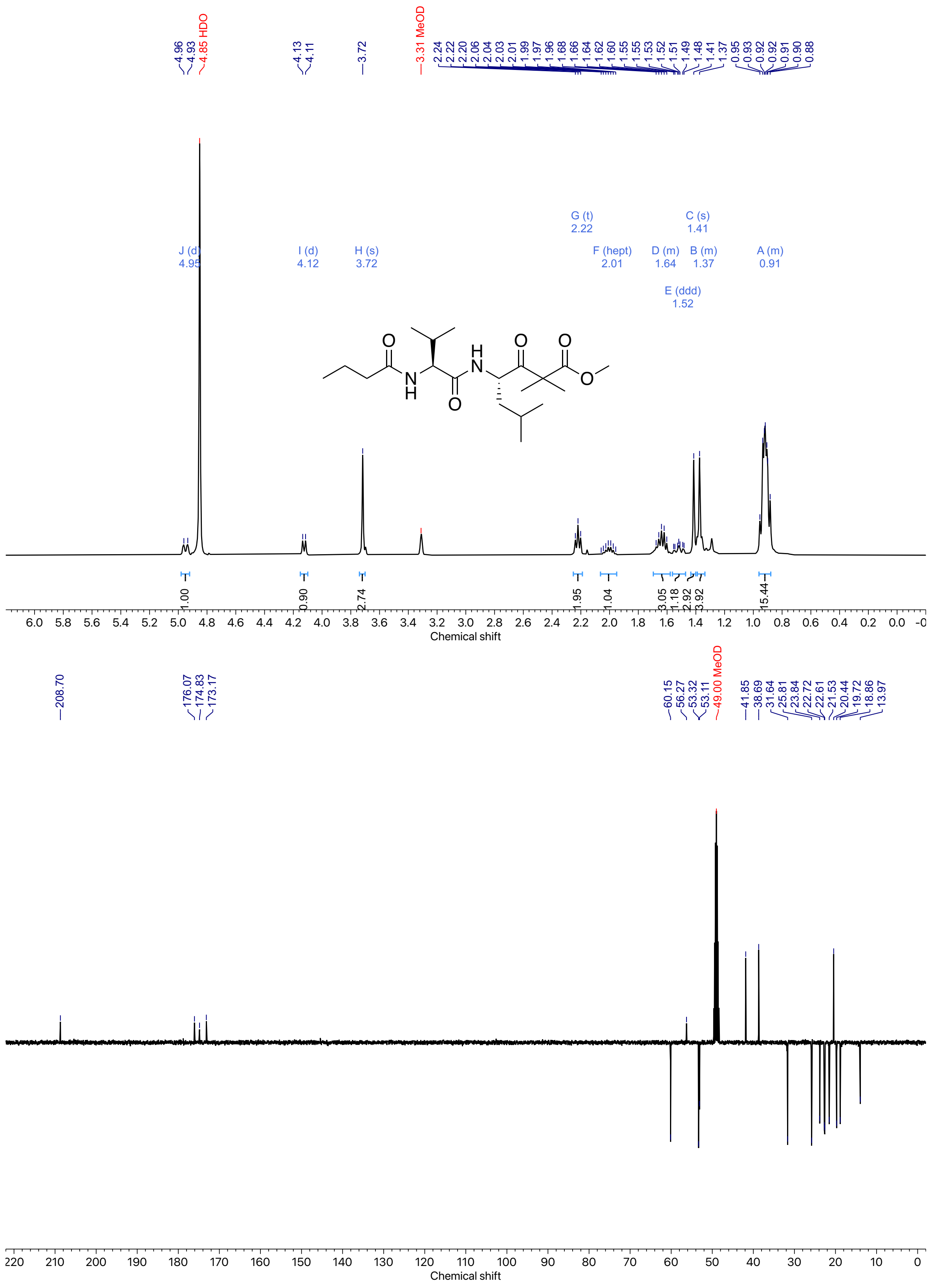

Figure 30: $^{1}H$ and $^{13}C$ NMR spectra of **27** (Methanol-d$_4$, 400 MHz).

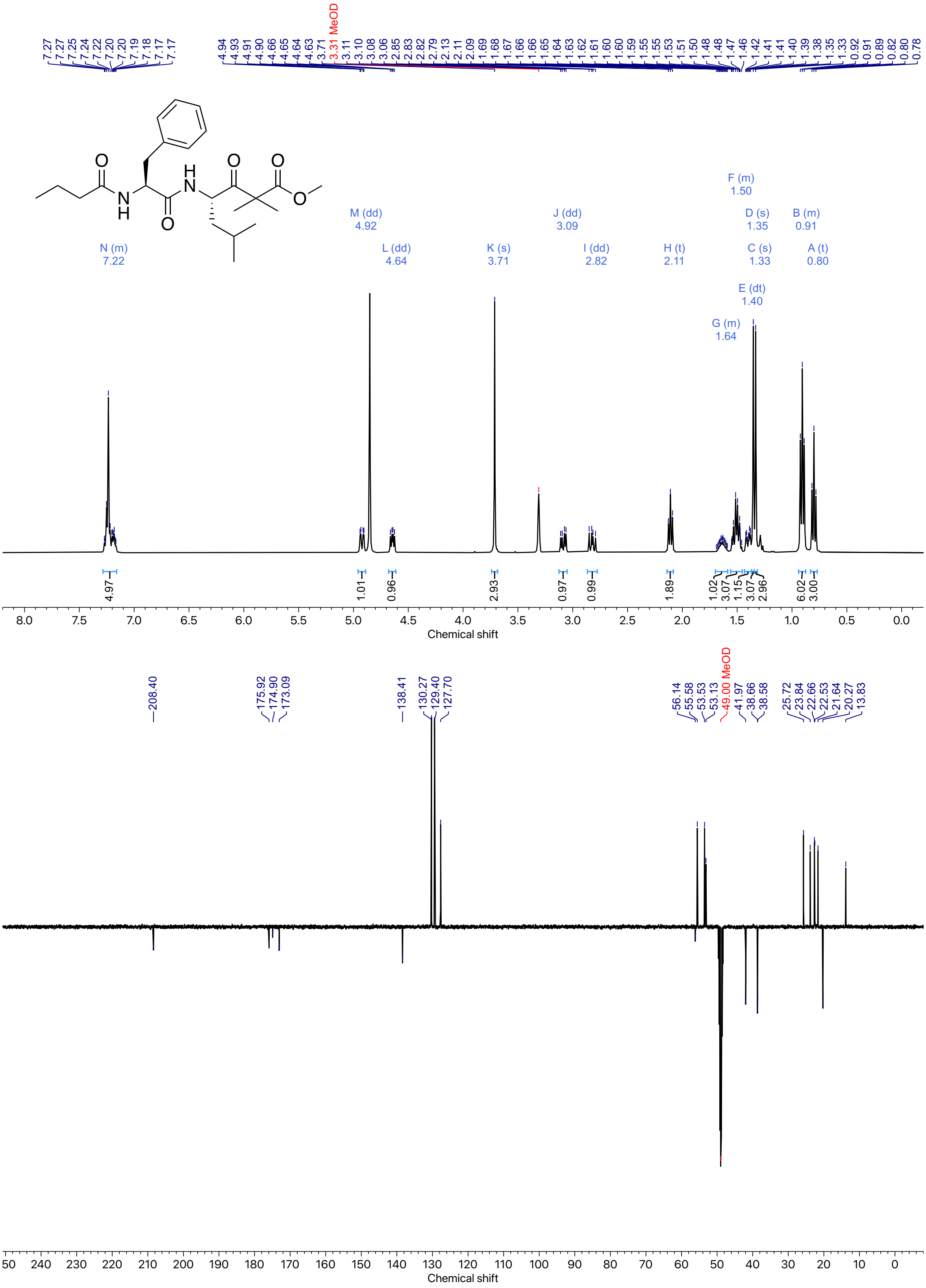

*Figure 31: [1]H and [13]C NMR spectra of* ***28*** *(Methanol-d$_4$, 400 MHz).*

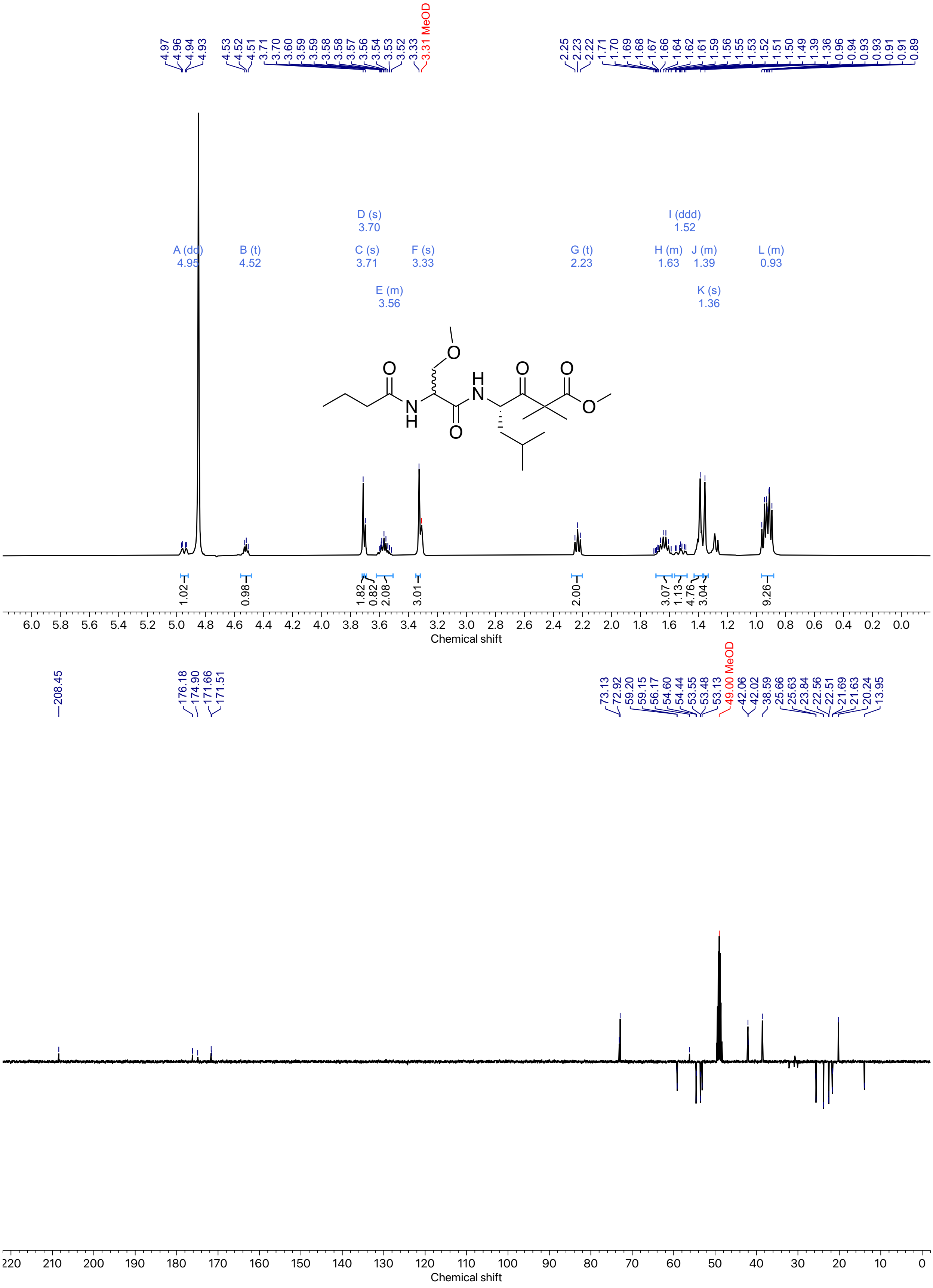

*Figure 32: [1]H and [13]C NMR spectra of **29** (Methanol-d₄, 400 MHz).*

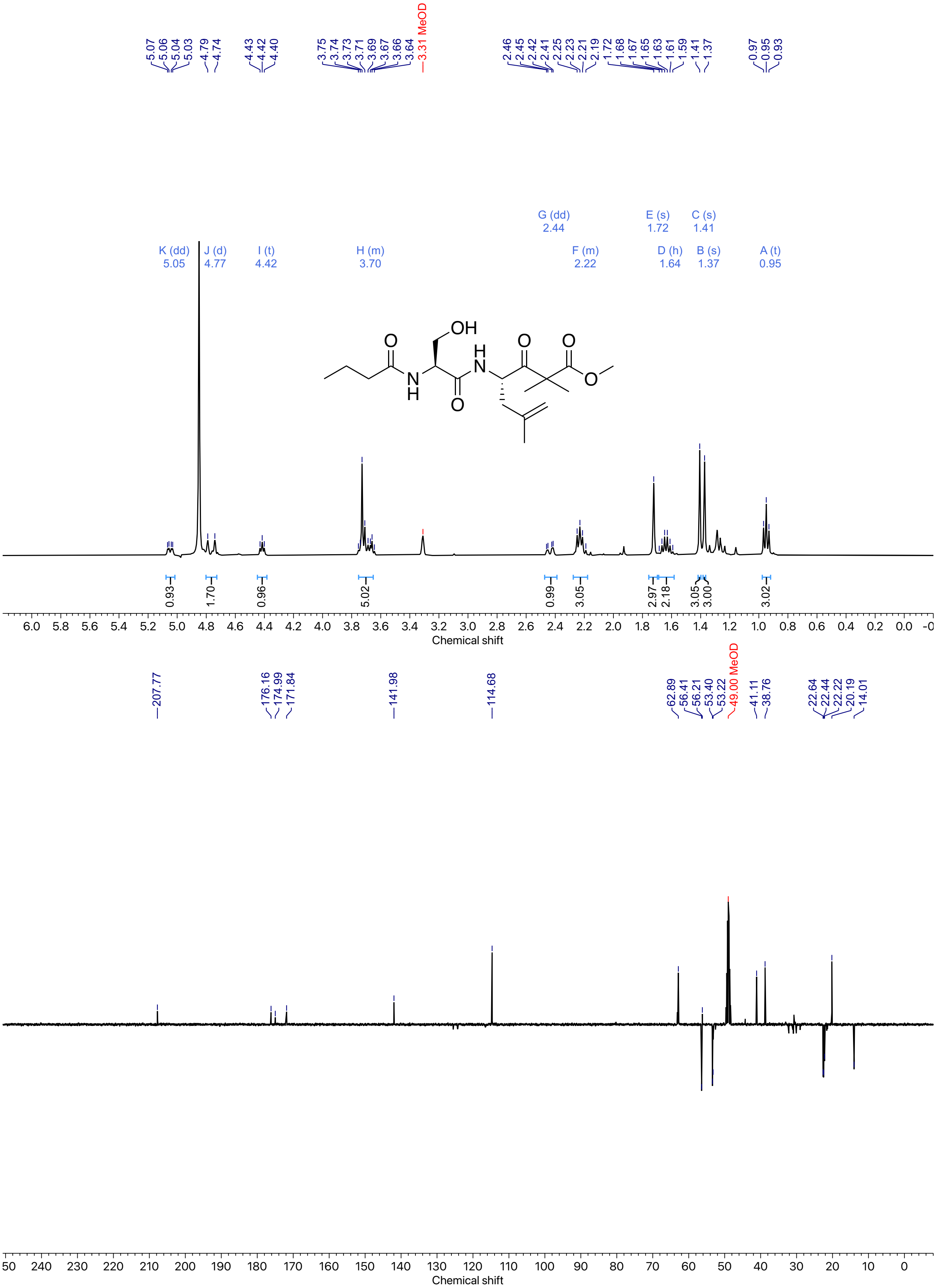

*Figure 33: [1]H and [13]C NMR spectra of **30** (Methanol-$d_4$, 400 MHz).*

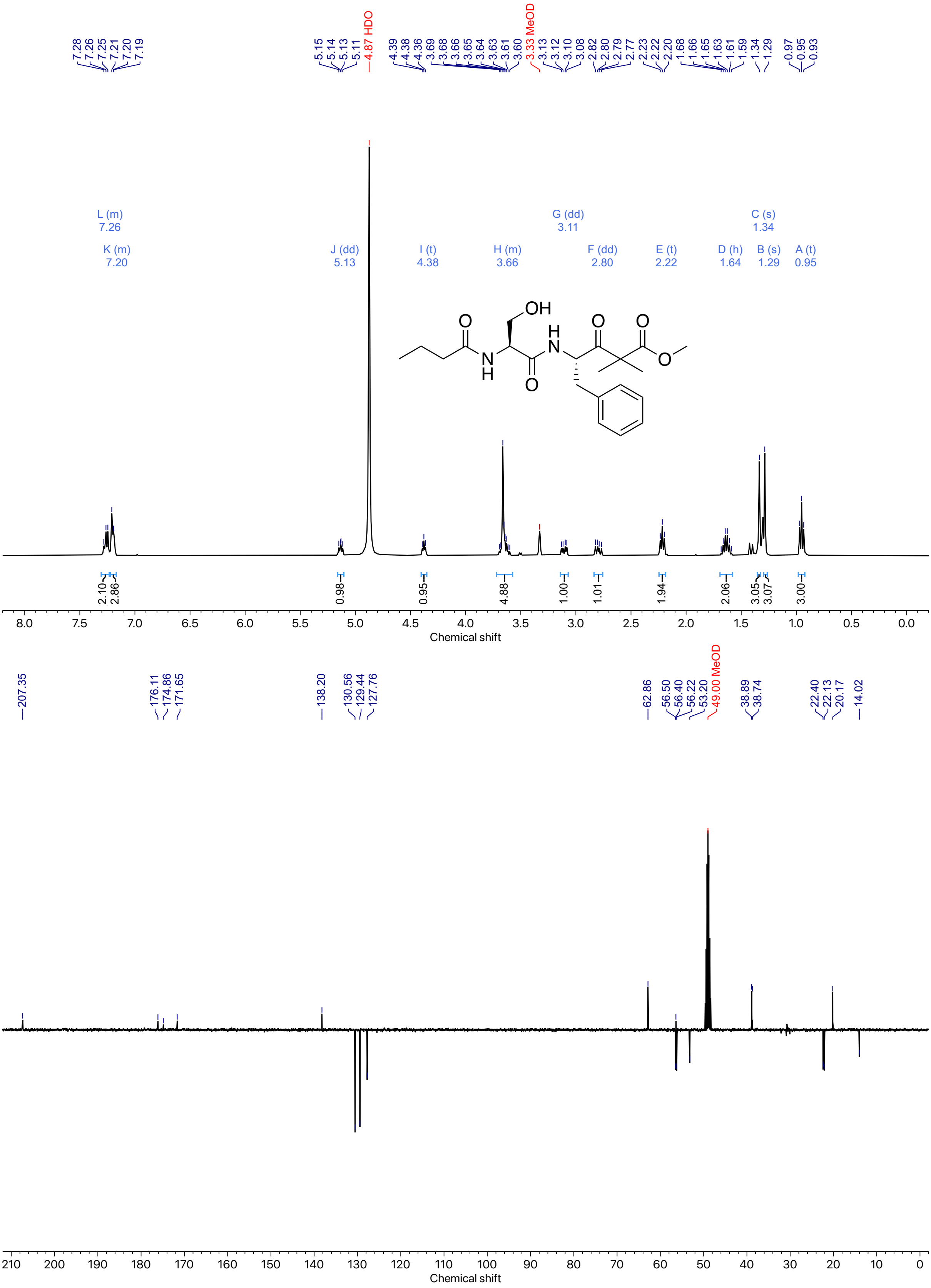

*Figure 34: ¹H and ¹³C NMR spectra of **31** (Methanol-d₄, 400 MHz).*

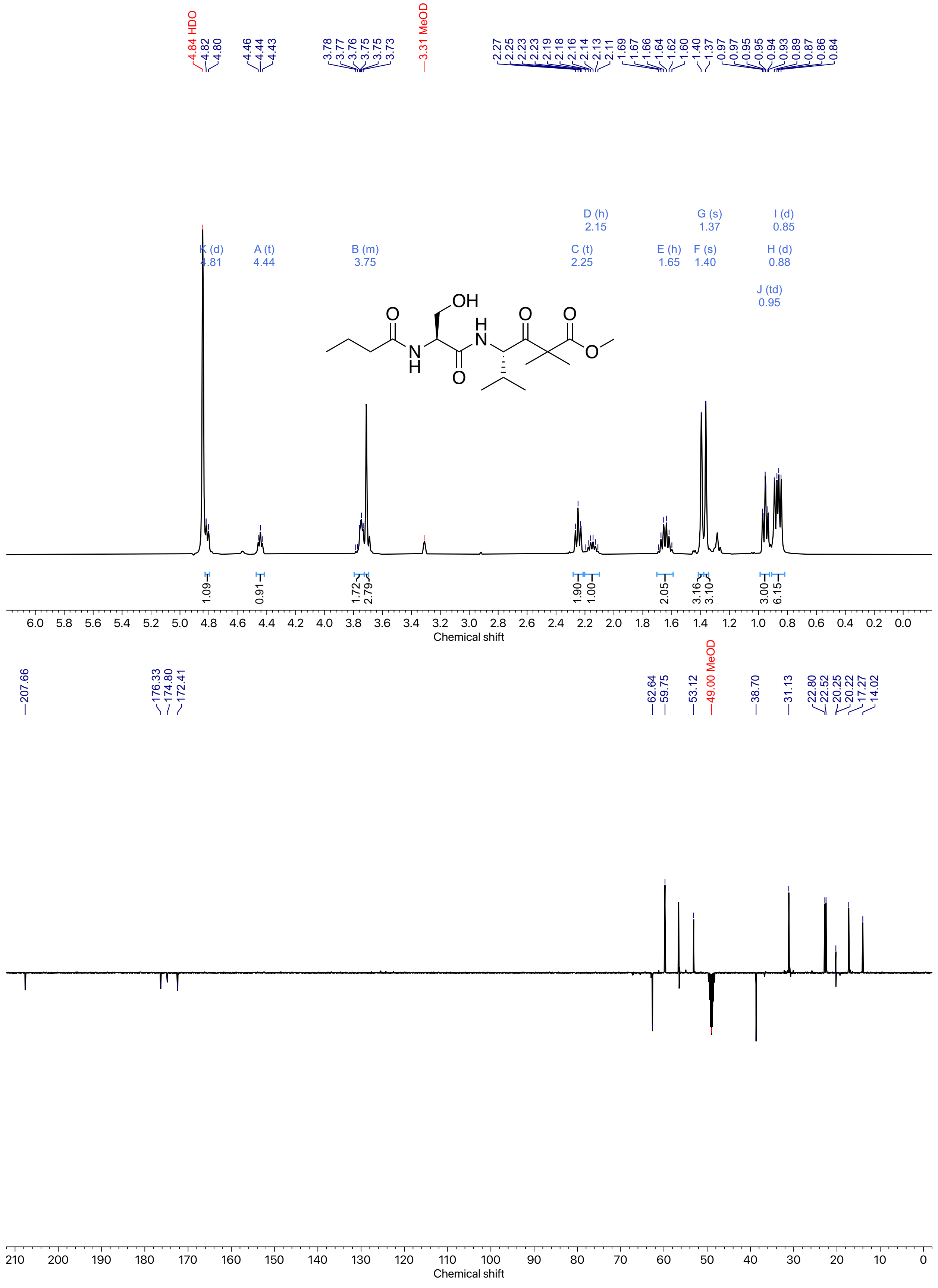

*Figure 35: $^{1}H$ and $^{13}C$ NMR spectra of **32** (Methanol-$d_4$, 400 MHz).*

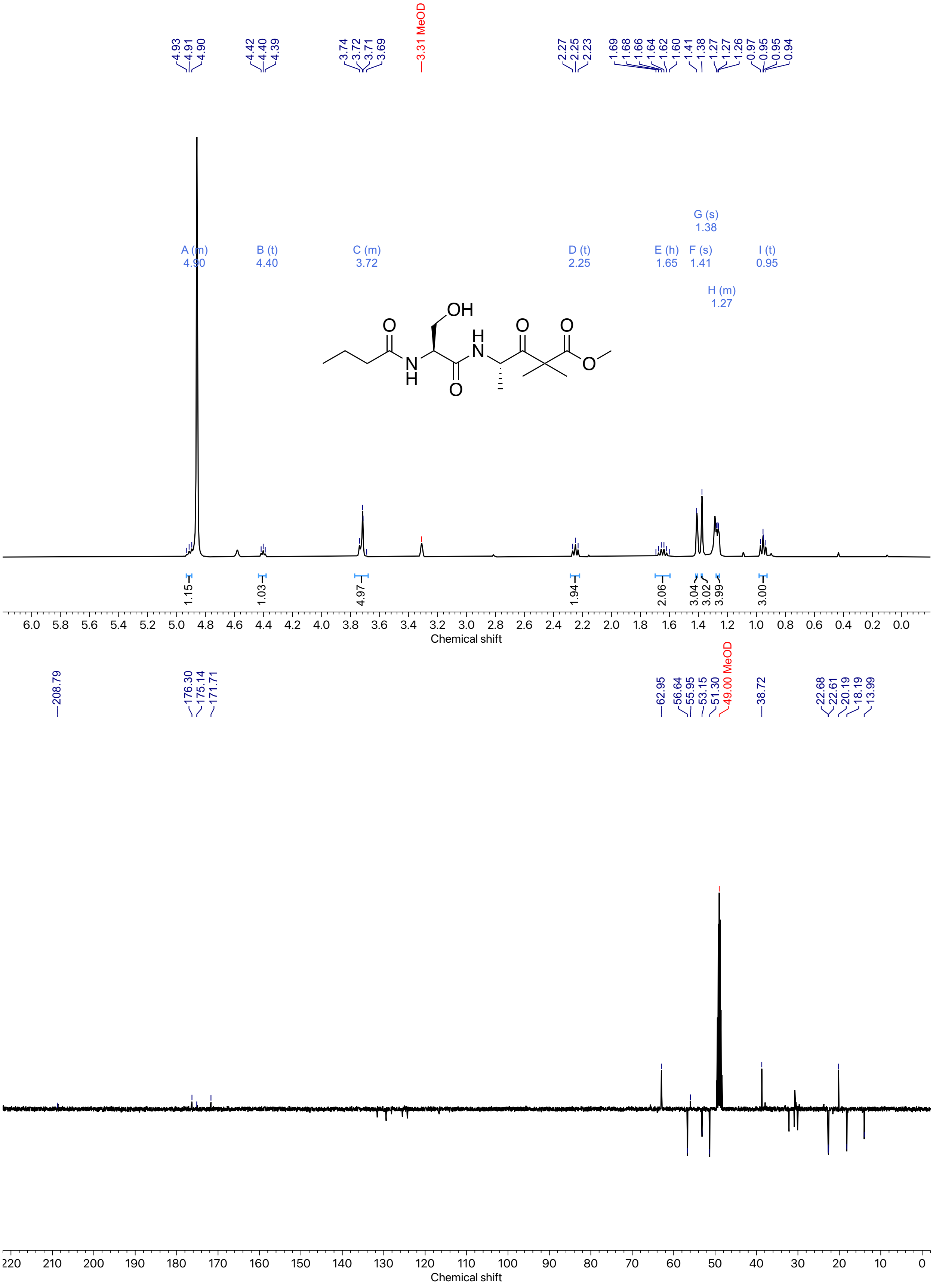

*Figure 36: $^1H$ and $^{13}C$ NMR spectra of **33** (Methanol-d$_4$, 400 MHz).*

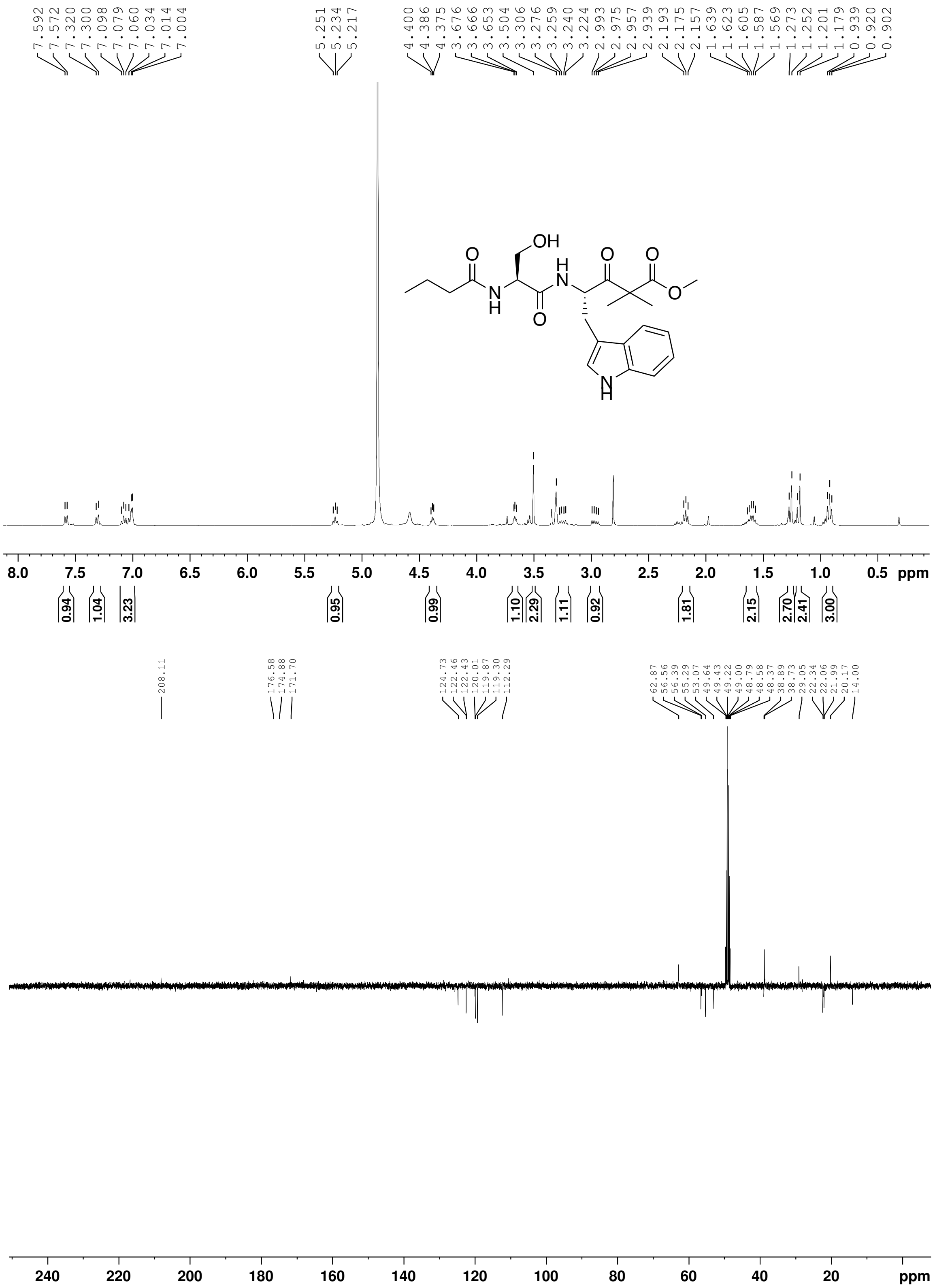

*Figure 37: ¹H and ¹³C NMR spectra of **34** (Methanol-$d_4$, 400 MHz). ¹³C peaks for quaternary carbons were assigned using HMBC analysis.*

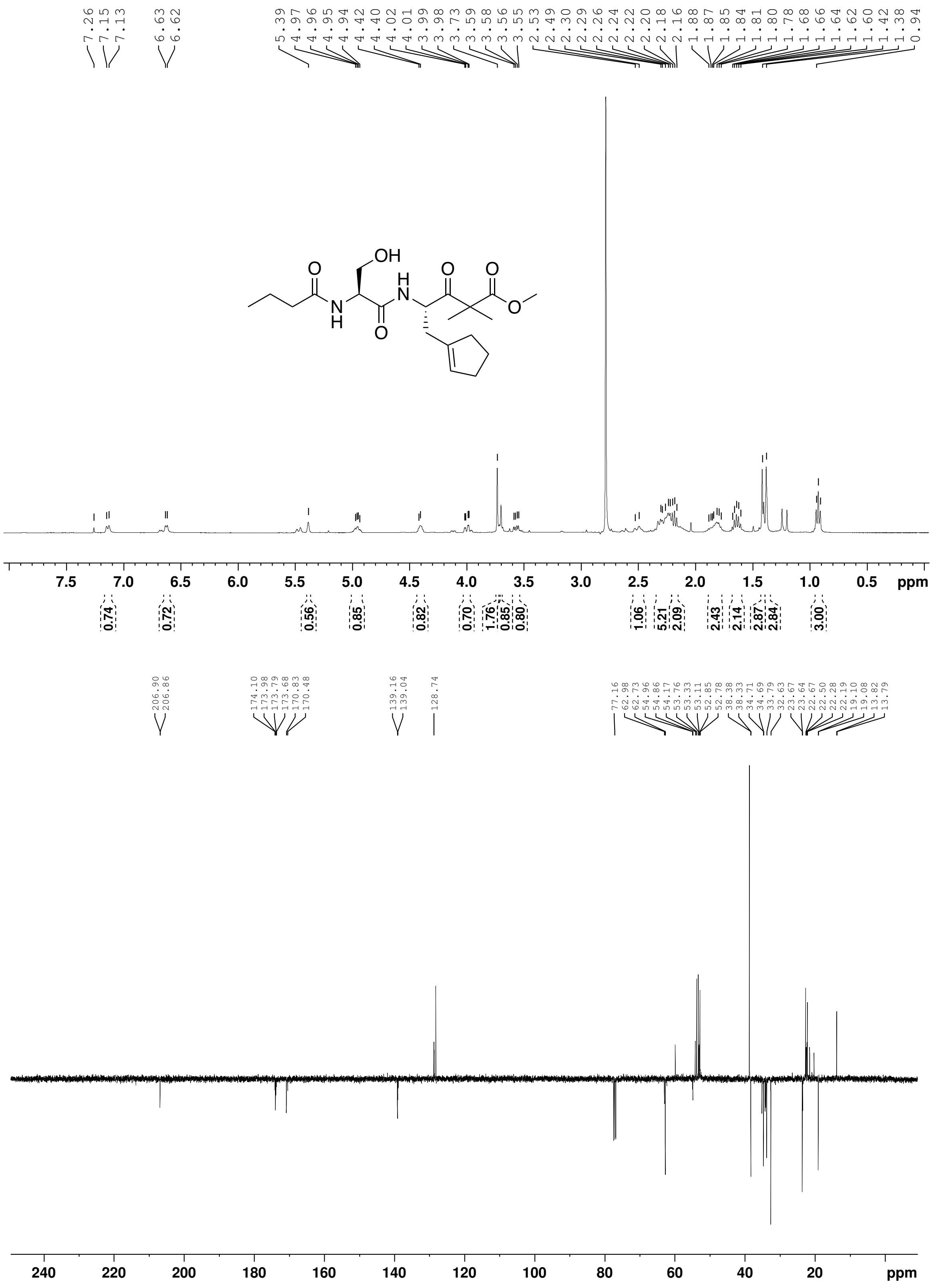

*Figure 38: $^{1}H$ and $^{13}C$ NMR spectra of **35** (Chloroform-d, 400 MHz).*

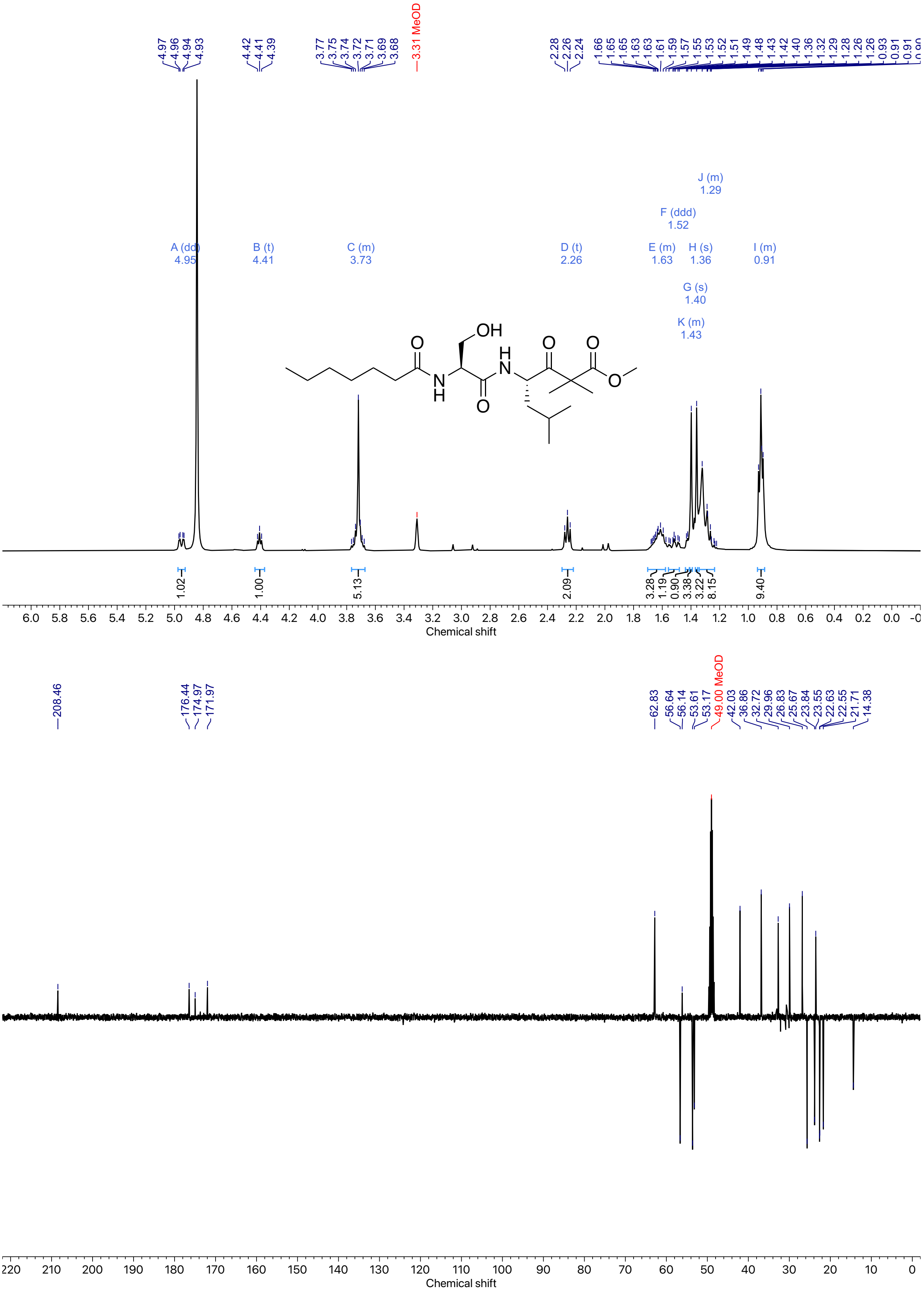

*Figure 39: $^{1}H$ and $^{13}C$ NMR spectra of **17** (Methanol-$d_4$, 400 MHz).*

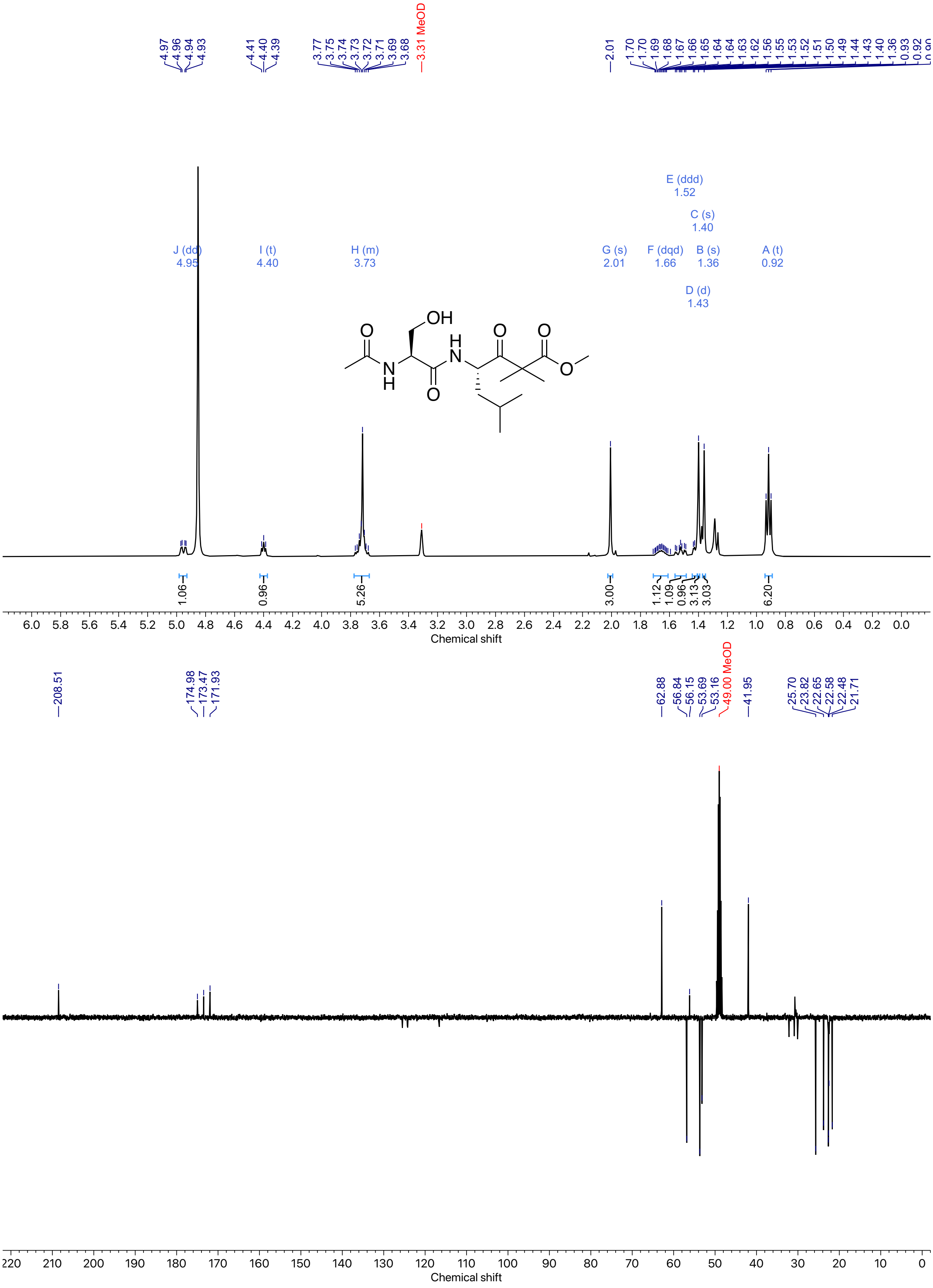

*Figure 40: [1]H and [13]C NMR spectra of **36** (Methanol-d₄, 400 MHz).*

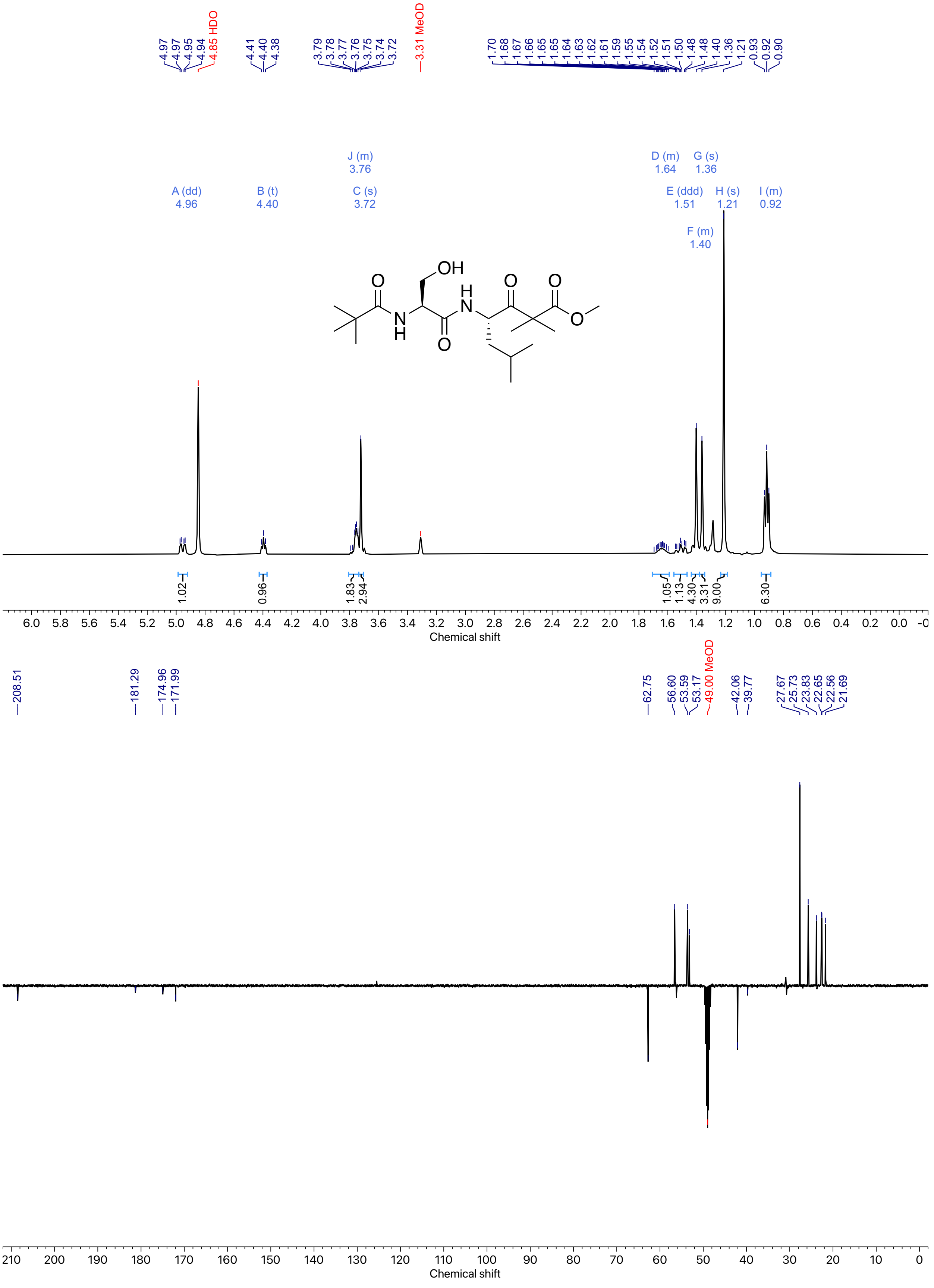

*Figure 41: $^{1}H$ and $^{13}C$ NMR spectra of **37** (Methanol-$d_4$, 400 MHz).*

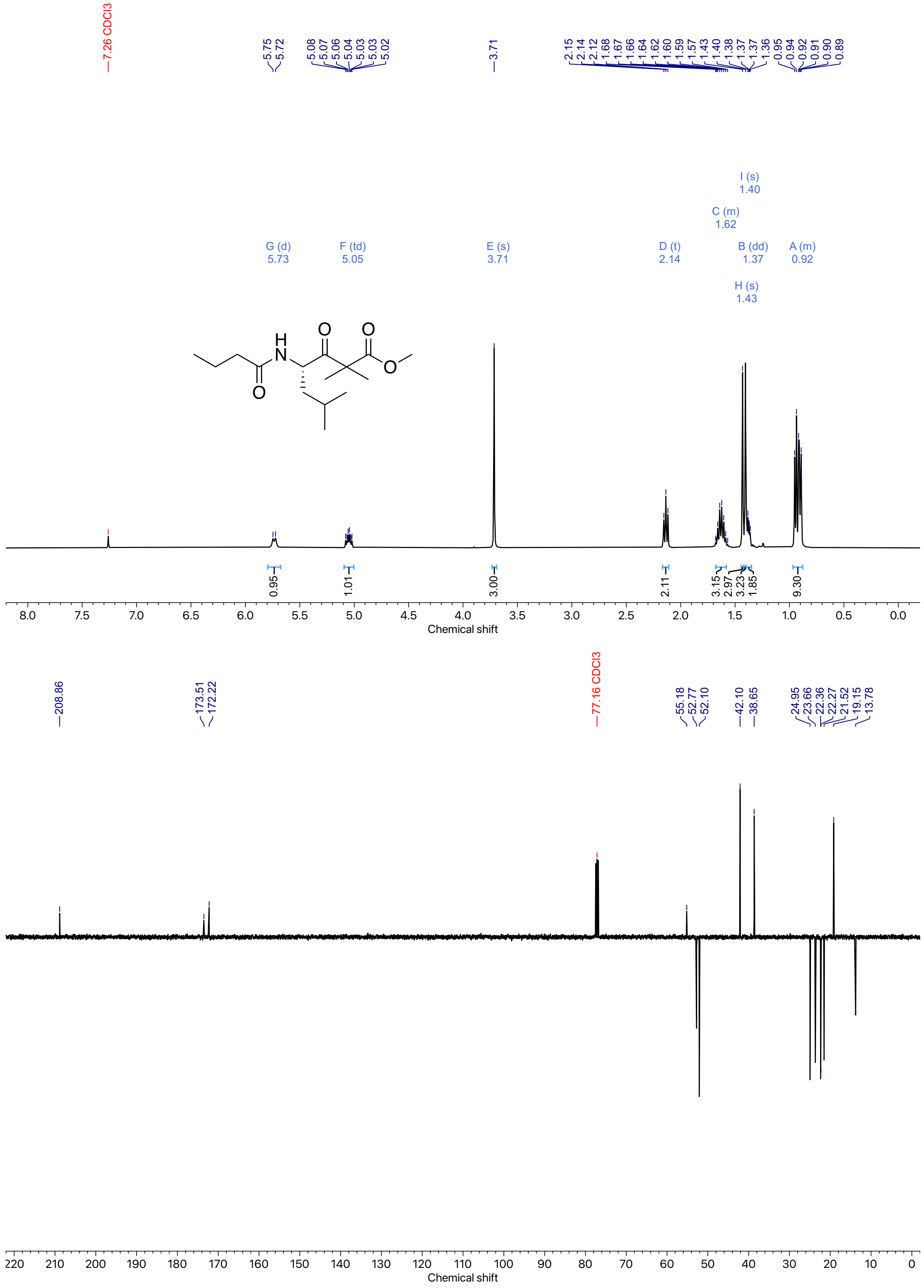

*Figure 42: [1]H and [13]C NMR spectra of **21** (Chloroform-d, 400 MHz).*

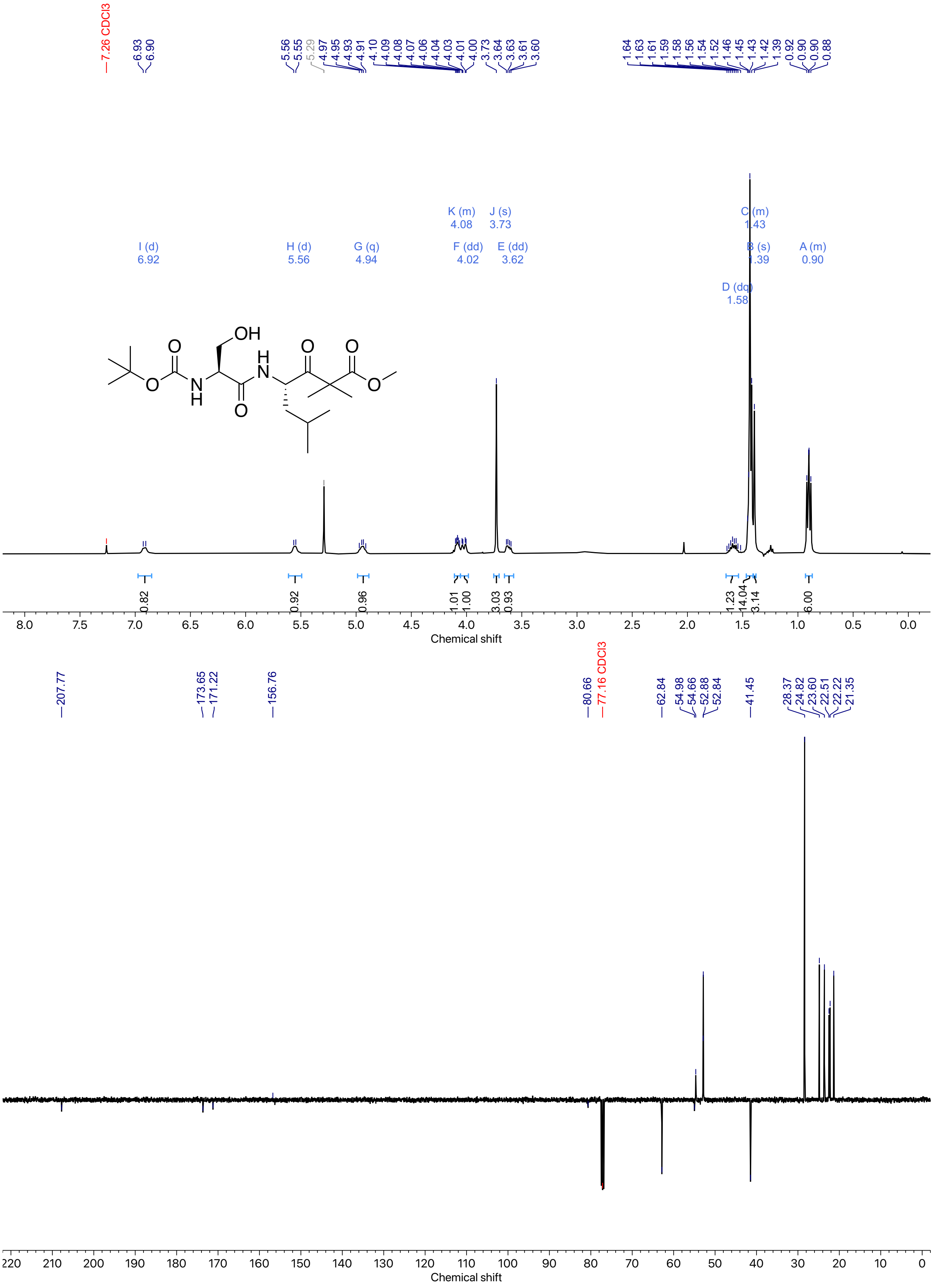

*Figure 43: $^{1}H$ and $^{13}C$ NMR spectra of **16** (Chloroform-d, 400 MHz).*

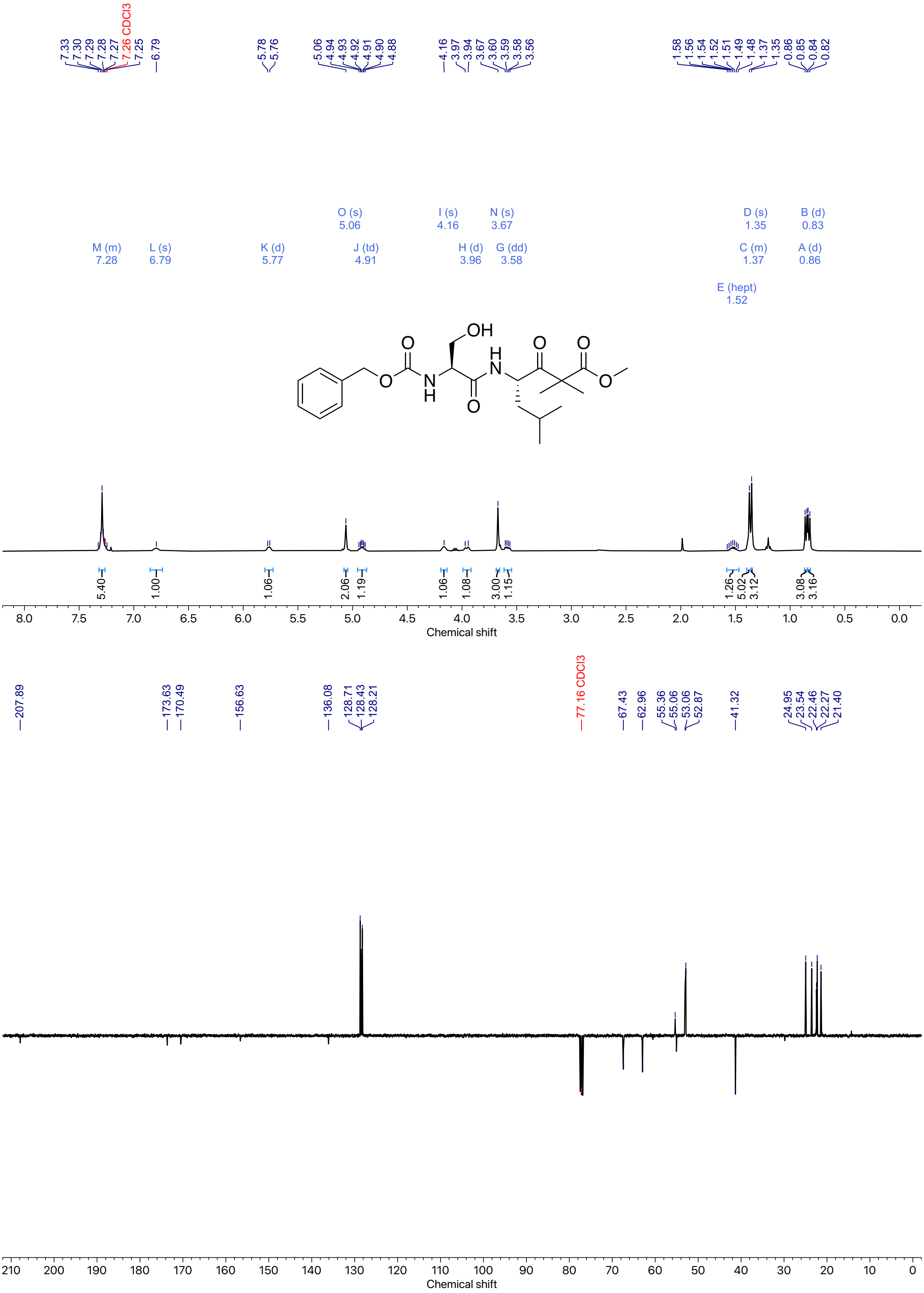

*Figure 44: [1]H and [13]C NMR spectra of **38** (Chloroform-d, 400 MHz).*

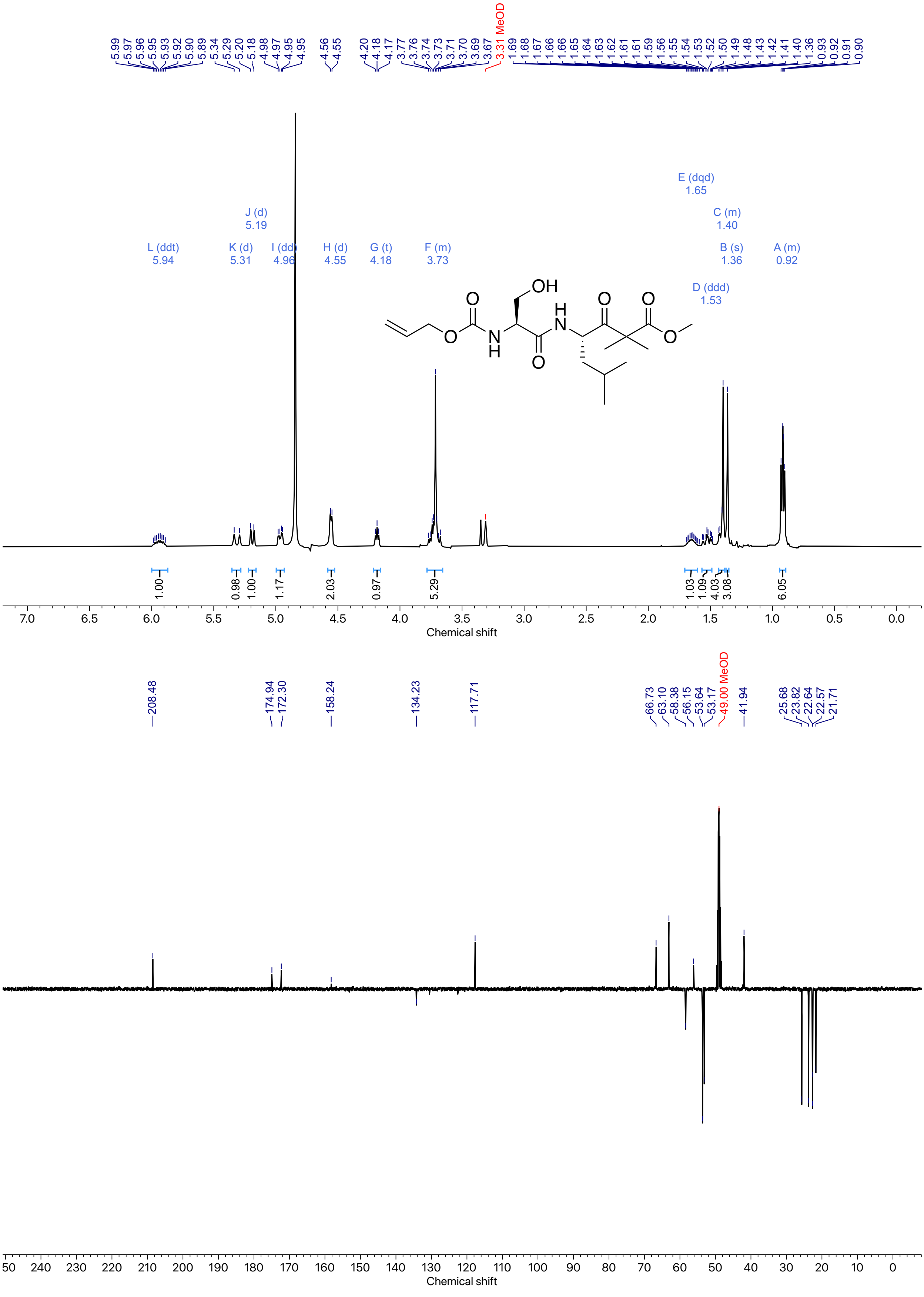

*Figure 45: $^{1}H$ and $^{13}C$ NMR spectra of **39** (Methanol-d$_4$, 400 MHz).*

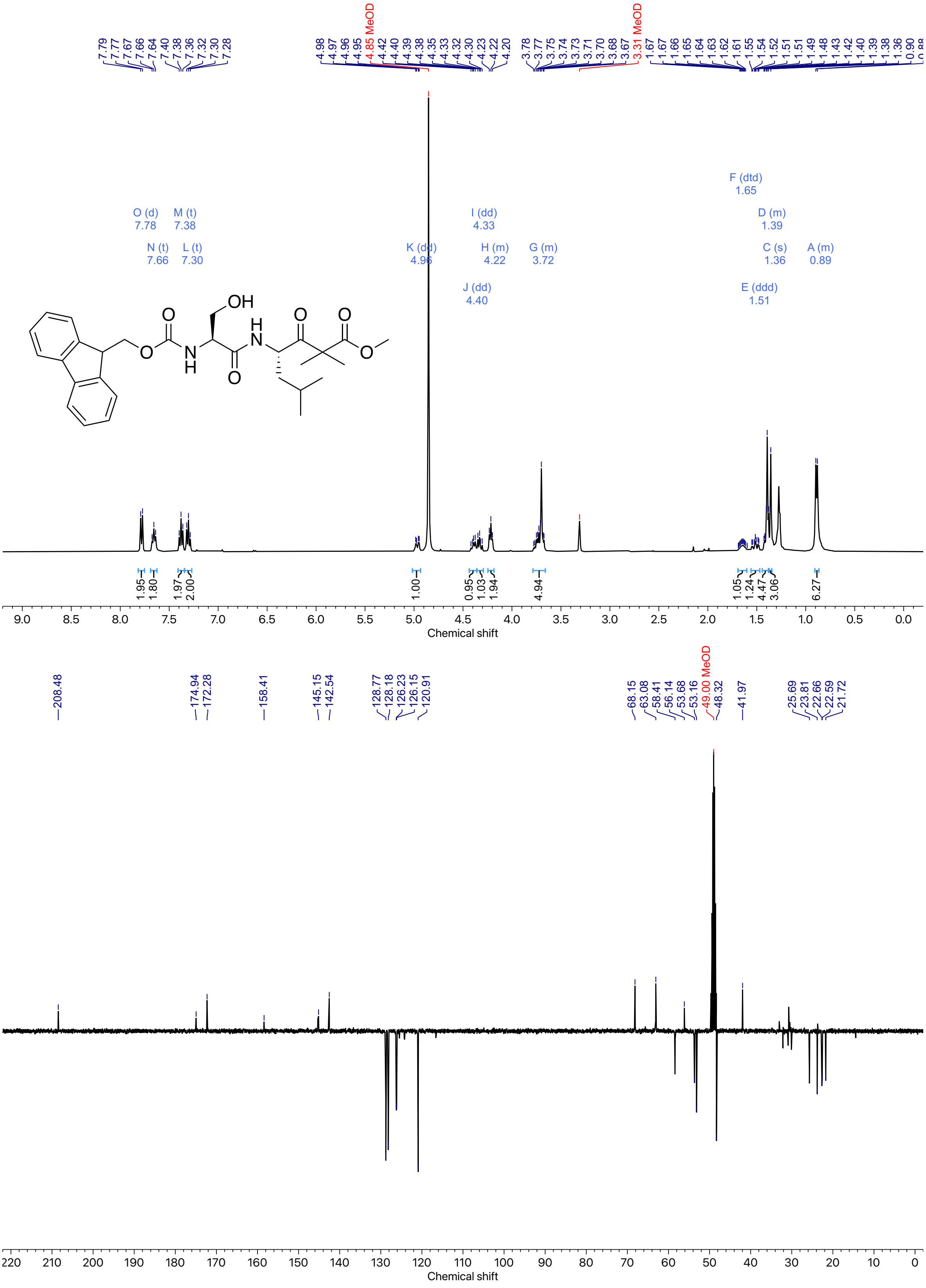

*Figure 46: [1]H and [13]C NMR spectra of **40** (Methanol-d$_4$, 400 MHz).*

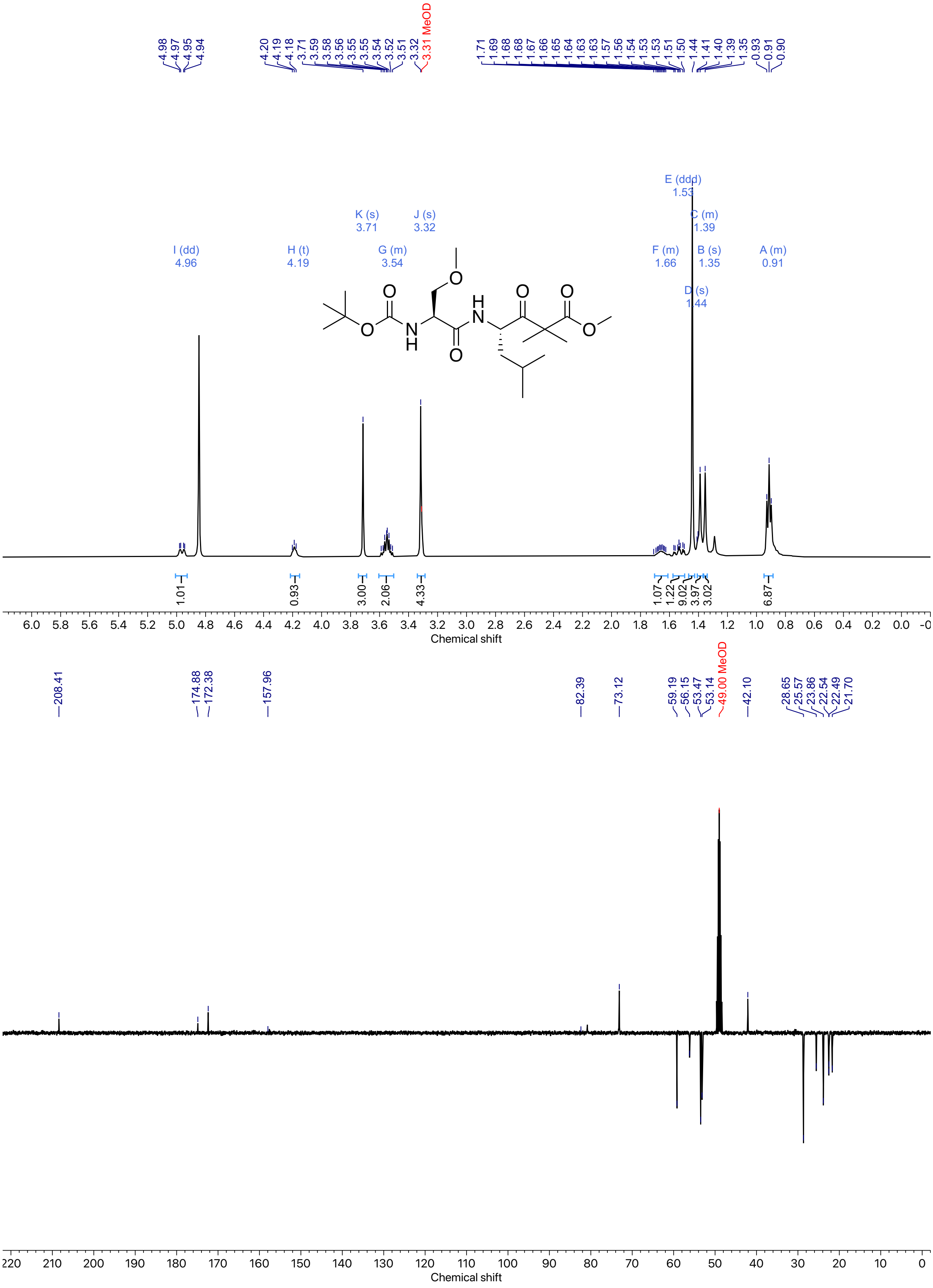

*Figure 47: [1]H and [13]C NMR spectra of **41** (Methanol-d$_4$, 400 MHz).*

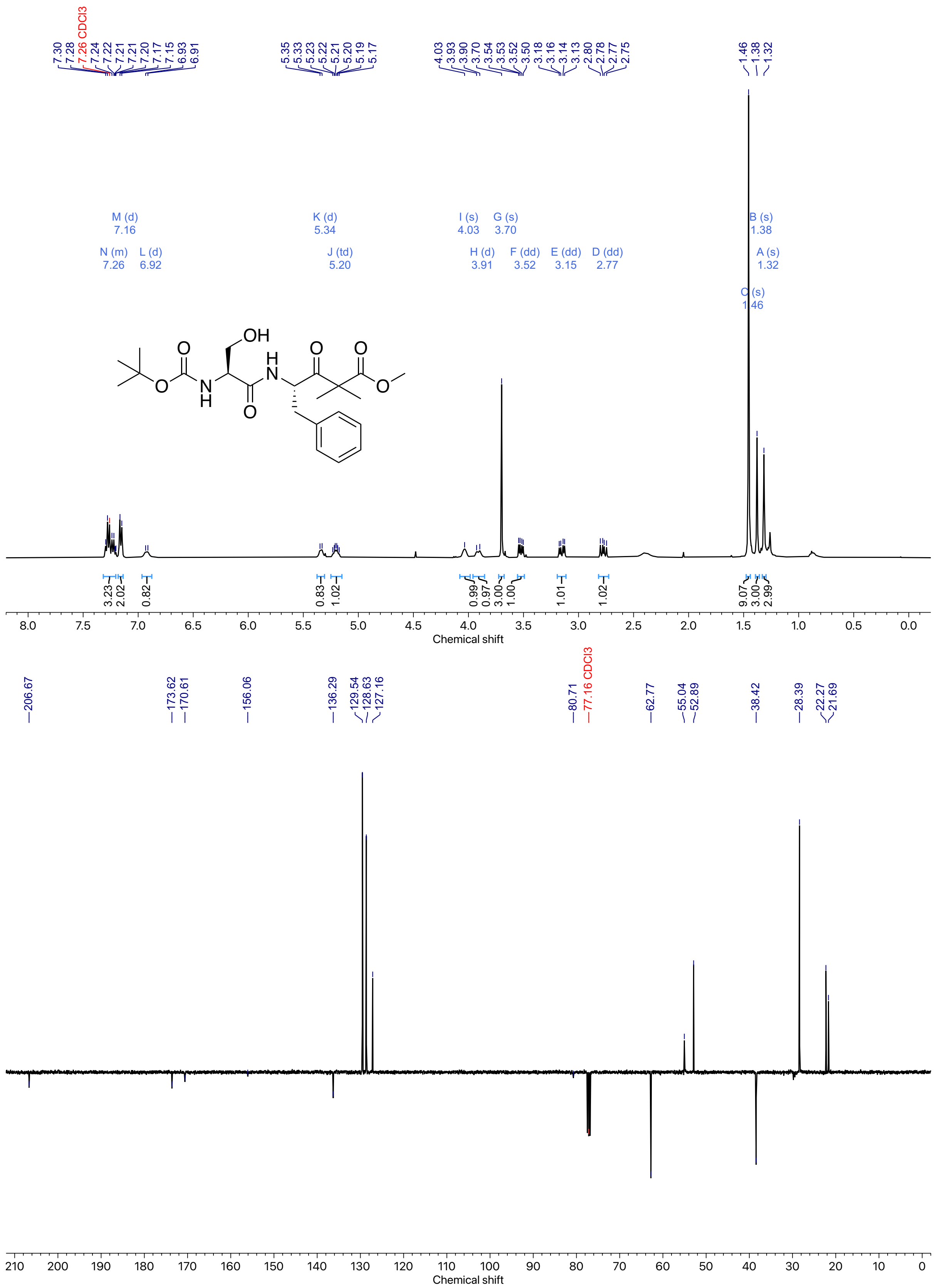

*Figure 48: $^{1}H$ and $^{13}C$ NMR spectra of **42** (Chloroform-d, 400 MHz).*

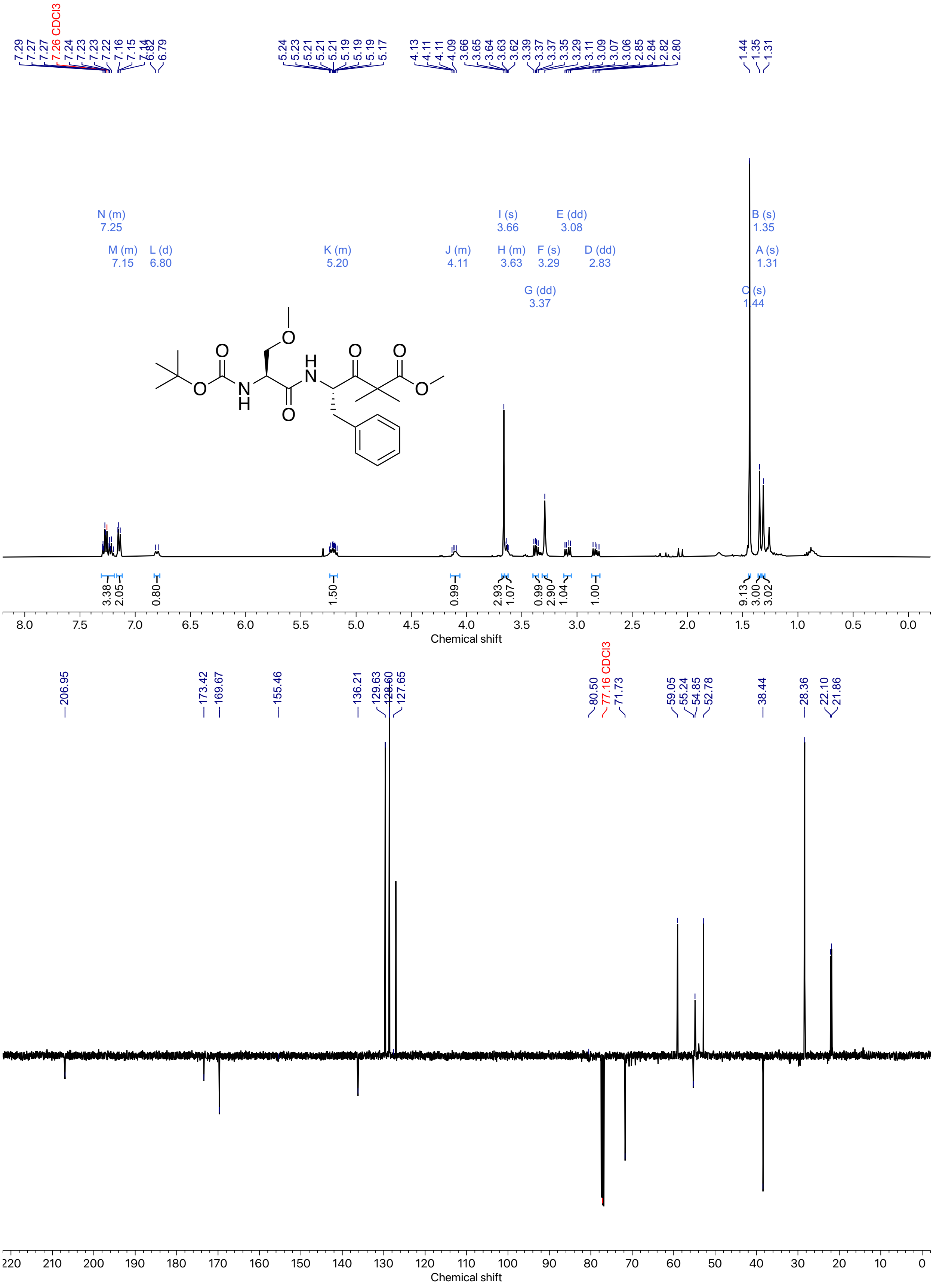

*Figure 49: $^{1}H$ and $^{13}C$ NMR spectra of **43** (Chloroform-d, 400 MHz).*

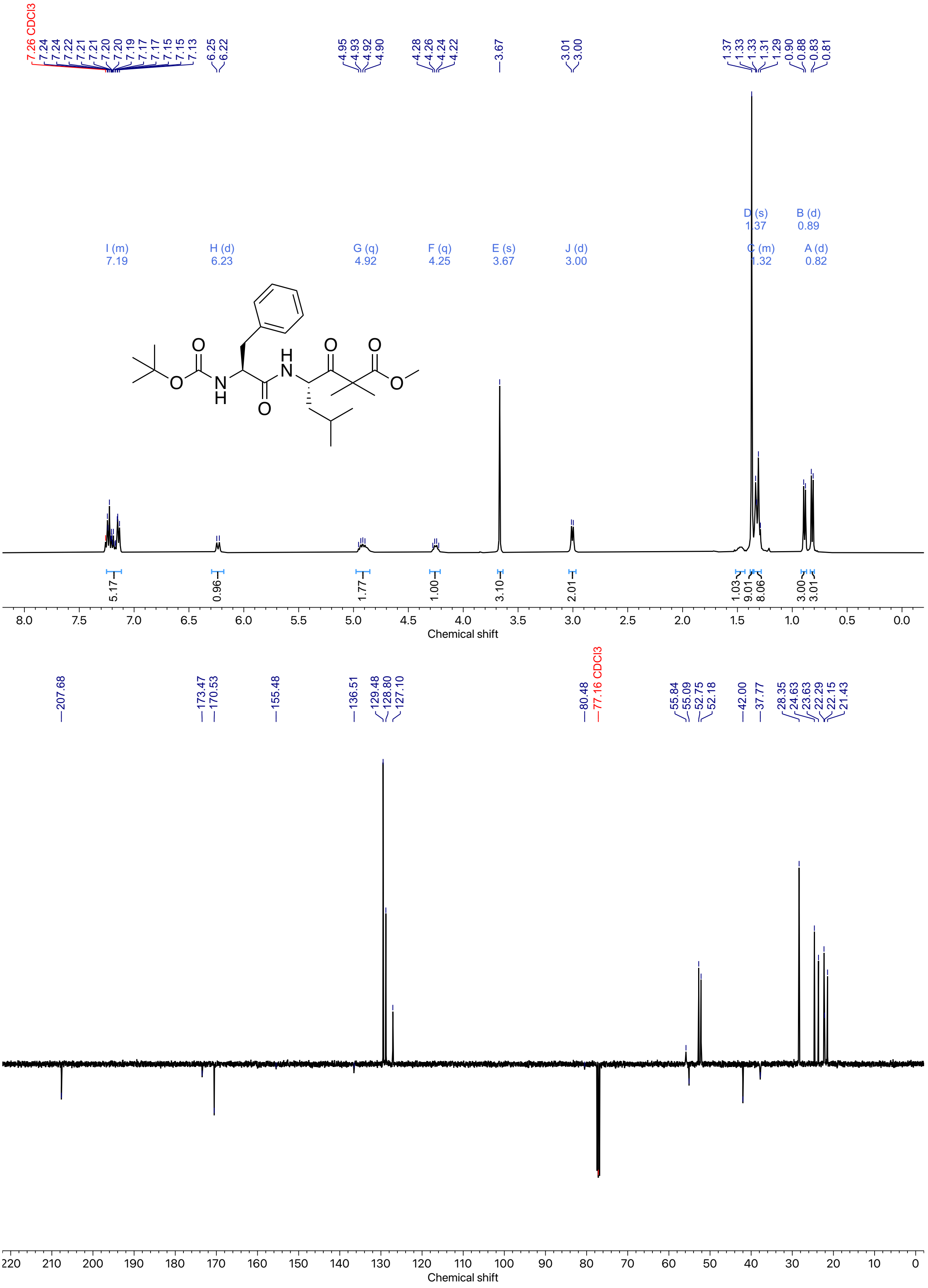

*Figure 50: $^1$H and $^{13}$C NMR spectra of **44** (Chloroform-d, 400 MHz).*

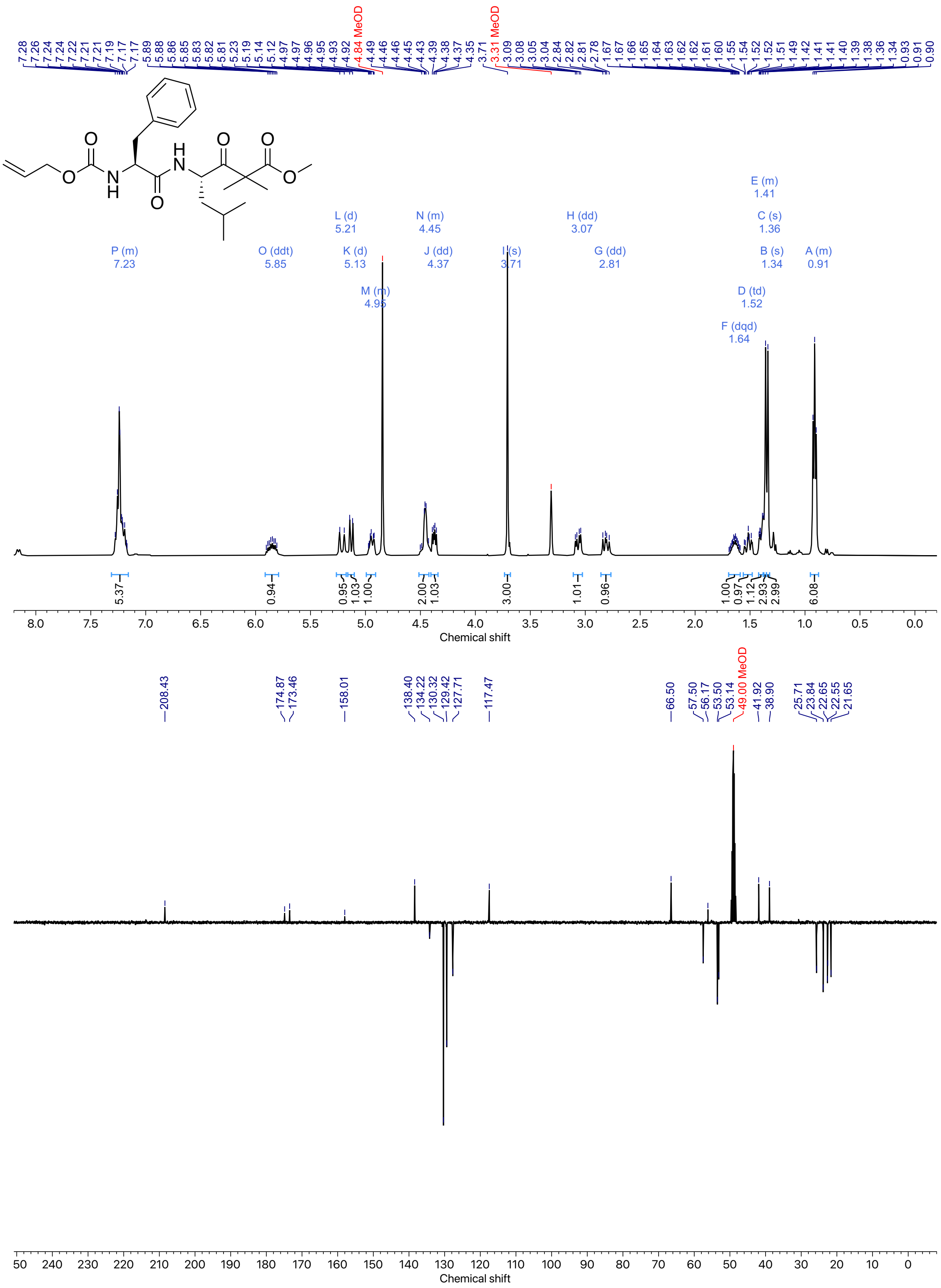

*Figure 51: $^{1}H$ and $^{13}C$ NMR spectra of **45** (Methanol-$d_4$, 400 MHz).*

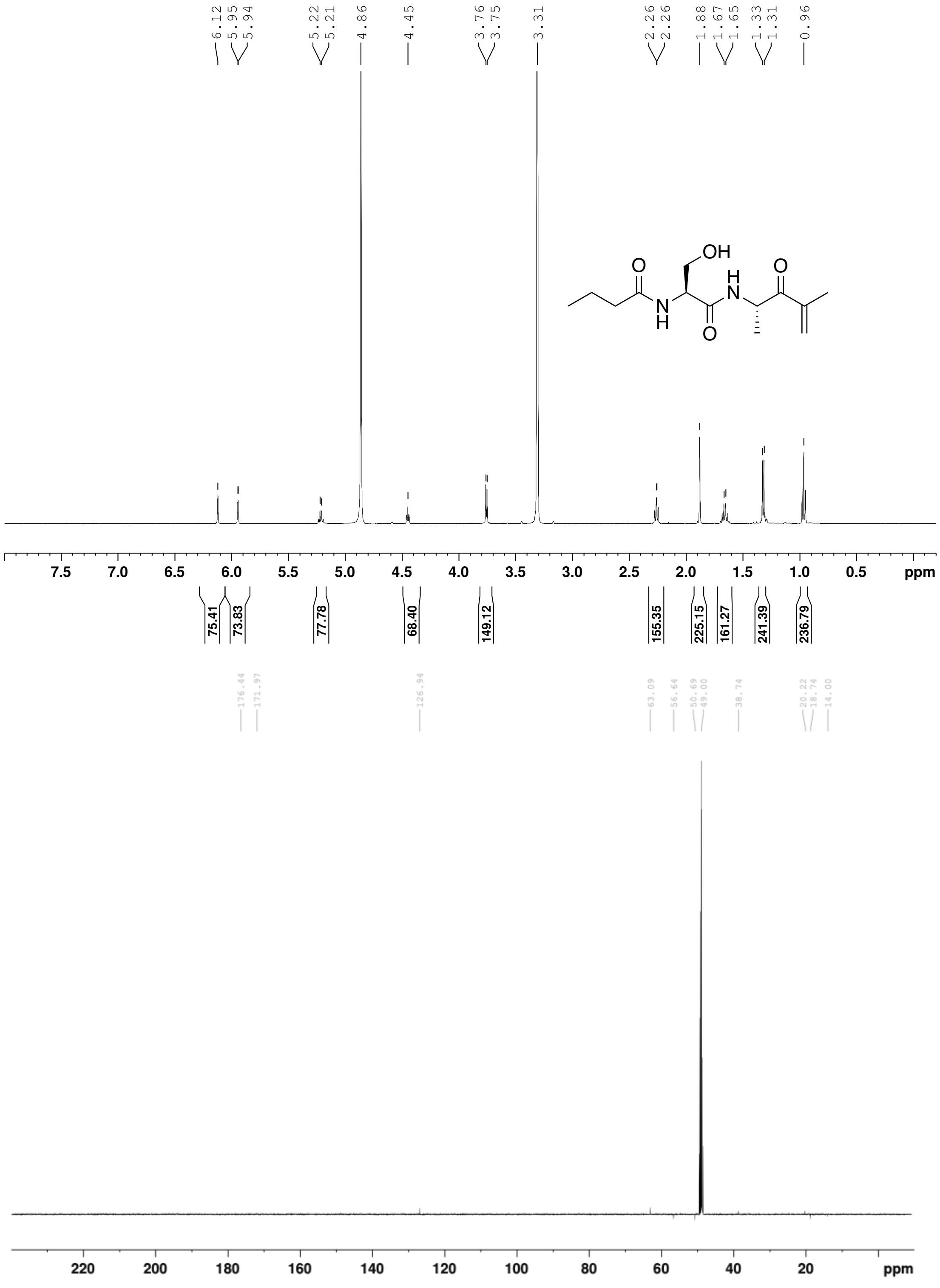

*Figure 52: $^{1}H$ and $^{13}C$ NMR spectra of **46** (Methanol-d$_4$, 500 MHz). $^{13}C$ peaks for quaternary carbons were assigned using HMBC analysis.*

(20) Sreerama, N.; Woody, R. W. Estimation of Protein Secondary Structure from Circular Dichroism Spectra: Comparison of CONTIN, SELCON, and CDSSTR Methods with an

Expanded Reference Set. *Anal. Biochem.* **2000**, *287* (2), 252–260. https://doi.org/10.1006/abio.2000.4880.

(21) Madeira, F.; Madhusoodanan, N.; Lee, J.; Eusebi, A.; Niewielska, A.; Tivey, A. R. N.; Lopez, R.; Butcher, S. The EMBL-EBI Job Dispatcher Sequence Analysis Tools Framework in 2024. *Nucleic Acids Res.* **2024**, *52* (W1), W521–W525. https://doi.org/10.1093/nar/gkae241.

(22) Kozlov, A. M.; Darriba, D.; Flouri, T.; Morel, B.; Stamatakis, A. RAxML-NG: A Fast, Scalable and User-Friendly Tool for Maximum Likelihood Phylogenetic Inference. *Bioinformatics* **2019**, *35* (21), 4453–4455. https://doi.org/10.1093/bioinformatics/btz305.

(23) Jumper, J.; Evans, R.; Pritzel, A.; Green, T.; Figurnov, M.; Ronneberger, O.; Tunyasuvunakool, K.; Bates, R.; Žídek, A.; Potapenko, A.; Bridgland, A.; Meyer, C.; Kohl, S. A. A.; Ballard, A. J.; Cowie, A.; Romera-Paredes, B.; Nikolov, S.; Jain, R.; Adler, J.; Back, T.; Petersen, S.; Reiman, D.; Clancy, E.; Zielinski, M.; Steinegger, M.; Pacholska, M.; Berghammer, T.; Bodenstein, S.; Silver, D.; Vinyals, O.; Senior, A. W.; Kavukcuoglu, K.; Kohli, P.; Hassabis, D. Highly Accurate Protein Structure Prediction with AlphaFold. *Nature* **2021**, *596* (7873), 583–589. https://doi.org/10.1038/s41586-021-03819-2.

(32) C. E. A. F. Schafmeister, W. S. Ross and V. Romanovski. LEaP, 1995.

(33) D.A. Case, H.M. Aktulga, K. Belfon, I.Y. Ben-Shalom, J.T. Berryman, S.R. Brozell, F.S. Carvahol, D.S. Cerutti, T.E. Cheatham, III, G.A. Cisneros, V.W.D. Cruzeiro, T.A. Darden, N. Forouzesh, M. Ghazimirsaeed, G. Giambaşu, T. Giese, M.K. Gilson, H. Gohlke, A.W. Goetz, J. Harris, Z. Huang, S. Izadi, S.A. Izmailov, K. Kasavajhala, M.C. Kaymak, I. Kolossv\'a ry, A. Kovalenko, T. Kurtzman, T.S. Lee, P. Li, Z. Li, C. Lin, J. Liu, T. Luchko, R. Luo, M. Machado, M. Manathunga, K.M. Merz, Y. Miao, O. Mikhailovskii, G. Monard, H. Nguyen, K.A. O'Hearn, A. Onufriev, F. Pan, S. Pantano, A. Rahnamoun, D.R. Roe, A. Roitberg, C. Sagui, S. Schott-Verdugo, A. Shajan, J. Shen, C.L. Simmerling, N.R. Skrynnikov, J. Smith, J. Swails, R.C. Walker, J. Wang, J. Wang, X. Wu, Y. Wu, Y. Xiong, Y. Xue, D.M. York, C. Zhao, Q. Zhu, and P.A. Kollman. Amber, 2025.

(40) Abramson, J.; Adler, J.; Dunger, J.; Evans, R.; Green, T.; Pritzel, A.; Ronneberger, O.; Willmore, L.; Ballard, A. J.; Bambrick, J.; Bodenstein, S. W.; Evans, D. A.; Hung, C.-C.; O'Neill, M.; Reiman, D.; Tunyasuvunakool, K.; Wu, Z.; Žemgulytė, A.; Arvaniti, E.; Beattie, C.; Bertolli, O.; Bridgland, A.; Cherepanov, A.; Congreve, M.; Cowen-Rivers, A. I.; Cowie, A.; Figurnov, M.; Fuchs, F. B.; Gladman, H.; Jain, R.; Khan, Y. A.; Low, C. M. R.; Perlin, K.; Potapenko, A.; Savy, P.; Singh, S.; Stecula, A.; Thillaisundaram, A.; Tong, C.; Yakneen, S.; Zhong, E. D.; Zielinski, M.; Žídek, A.; Bapst, V.; Kohli, P.; Jaderberg, M.; Hassabis, D.; Jumper, J. M. Accurate Structure Prediction of Biomolecular Interactions with AlphaFold 3. *Nature* **2024**, *630* (8016), 493–500. https://doi.org/10.1038/s41586-024-07487-w.
